# Novel naphthyridones targeting Pannexin 1 for colitis management

**DOI:** 10.1101/2024.09.15.613164

**Authors:** Wen-Yun Hsueh, Yi-Ling Wu, Meng-Tzu Weng, Shin-Yun Liu, Jascinta P Santavanond, Yi-Chung Liu, Ching-I Lin, Cheng-Nong Lai, Yi-Ru Lu, Jing Yin Hsu, Hong-Yu Gao, Jinq-Chyi Lee, Shu-Chen Wei, Ping-Chiang Lyu, Ivan K H Poon, Hsing-Pang Hsieh, Yu-Hsin Chiu

## Abstract

Pannexin 1 (PANX1) forms cell-surface channels capable of releasing signaling metabolites for diverse patho-physiological processes. While inhibiting dysregulated PANX1 is proposed as a therapeutic strategy for many pathological conditions, including inflammatory bowel disease (IBD), low efficacy or poor specificity of classical PANX1 inhibitors introduces uncertainty for their applications in basic and translational research. Here, we performed hit-to-lead optimization and identified a naphthyridone, compound **12**, as a new PANX1 inhibitor with an IC_50_ of 0.73 μM that does not affect pannexin-homologous LRRC8/SWELL1 channels. Using structure-activity relationship analysis, mutagenesis, cell thermal shift assays, and molecular docking, we revealed that compound **12** directly engages PANX1 Trp74 residue. Using a dextran sodium sulfate mouse model of IBD, we found that compound **12** markedly reduced colitis severity, highlighting new PANX1 inhibitors as a proof-of-concept treatment for IBD. These data describe the mechanism of action for a new PANX1 inhibitor, identify the binding site for future drug design, and present a targeted strategy for treating IBD.

**Teaser:** A specific PANX1 inhibitor presents a proof-of-concept treatment for inflammatory bowel disease.

## Introduction

Every living organism faces daily life-threatening challenges, such as infection, toxins, mechanical injury, or chronic inflammation. In damaged tissues, dead or dying cells can present damage-associated molecular patterns (DAMPs) to trigger a series of responses, including recruitment, proliferation, and activation of circulating or tissue-resident immune cells (*1*). Particularly, ATP is a well-recognized DAMP that can incite inflammatory responses during pathogen infection or tissue injury (*2*) via engaging various purinergic receptors (*3*).

Pannexin 1 (PANX1) constitutes plasma membrane ion channels expressed in a wide variety of vertebrate tissues (*4*). PANX1 channels are well-known for their ability to release ATP and other intracellular metabolites ≤1 kDa, such as glutamate and spermidine (*5, 6*). These metabolites can initiate various receptor signaling pathways in an autocrine or paracrine manner. For example, activated PANX1 channels in apoptotic cells can release ATP and UTP as “find-me” signals to recruit phagocytes for expedited removal of apoptotic corpse via P2Y2 receptor pathways (*7*). On the other hand, aberrant activation of PANX1 channels has also been implicated in diverse disease states, including hypertension (*8*), diabetes (*9*), seizure (*10*), chronic neuropathic pain (*11*), ischemia-reperfusion injury (*12*), as well as inflammatory bowel disease (*13*). Thus, pharmacological inhibition of PANX1 channels is considered as a therapeutic strategy for multiple disease conditions (*14*).

Inflammatory bowel disease (IBD), comprising Crohn’s disease (CD) and ulcerative colitis (UC), is characterized by chronic and relapsing inflammation in the gastrointestinal tract. Over the years, IBD patients represent 5% of the population in the Western countries, and the prevalence of IBDs is rising in the Asian-Pacific countries (*15*). Multiple treatments are available for treating IBD, including corticosteroids, anti-inflammatory reagents or immunosuppressives. However, many patients relapse or show intolerance to these agents (*16*), emphasizing the need to develop alternative and personalized therapeutics. While the emergence of biologic therapies, such as anti-TNF antibodies, presents an encouraging breakthrough in IBD treatment, higher annual cost of the biologics is considerable for patients and increases healthcare spending given high prevalence of IBDs globally (*15*).

A previous study reported that ^10^Panx or a purinergic P2X7 receptor antagonist significantly decreased the crypt damage in tumor necrosis factor alpha (TNFα) and IL-1β induced colitis of human colonic mucosa strip (*17*). Consistent with this, PANX1 inhibition using ^10^Panx or probenecid could protect against the loss of myenteric neurons in mouse models of colitis (*13*). Therefore, PANX1 inhibitors represent an alternative therapeutic approach for IBD and the associated intestinal dysmotility. However, classical PANX1 inhibitors, such as carbenoxolone (CBX) (*18*), probenecid (*19*), mefloquine (*20*), and ^10^Panx (*21*), have non-specific effects on many topologically-or functionally-related channels and receptors (*22*), limiting their applications in basic and translational research.

To address these pharmacological limitations of existing PANX1 blockers, we previously undertook an unbiased screening campaign and discovered trovafloxacin (Trovan) as a new PANX1 inhibitor that did not affect activity of topologically-similar PANX2 or Connexin 43 (Cx43) (hemi)channels (*23*). Trovan is likely a direct PANX1 inhibitor since it can reduce PANX1 single-channel activity in cell-free, inside-out membrane patches. Later studies also demonstrated that inhibition of PANX1 channels using Trovan improved the outcome of traumatic brain injury or chronic neuropathic pain in mice, highlighting Trovan as a potential therapeutic strategy for inflammatory disorders (*11, 24*). However, Trovan has been withdrawn from clinical use since early 2000s due to reported incidents of idiosyncratic hepatotoxicity (*25*).

In this study, we employed the hit-to-lead optimization approach and identified a new naphthyridone, compound **12**, as a potent PANX1 inhibitor that does not have non-specific effects on the volume-regulated anionic LRRC8/SWELL1 channels. We used a number of orthogonal methods, including whole-cell voltage-clamp recordings, fluorescent dye-uptake assays, ATP release assays, and apoptotic body formation assays, to comprehensively assess inhibitory effects of new inhibitors on PANX1 channels in heterologous and endogenous systems. By combining a structure-activity relationship (SAR) analysis of varying naphthyridone analogues with mutagenesis analysis of PANX1 channels, we uncovered the mechanism of action for naphthyridone-mediated PANX1 inhibition, including the critical moieties responsible for PANX1 inhibition and the amino acid residues for direct interactions with the inhibitor. We also examined compound **12** as a proof-of-concept approach for treating inflammatory bowel disease by using a dextran sodium sulfate (DSS)-induced mouse model of colitis, further underscoring the potentials of new PANX1 inhibitors in basic and translational research.

## Results

### New PANX1 inhibitors are identified from the newly synthesized naphthyridones

By screening a library of pharmacologically active compounds, a previous study identified trovafloxacin (Trovan), a fourth-generation fluoroquinolone antibiotic, as a PANX1 inhibitor that did not affect the activity of PANX2 or Cx43 (*23*). Additionally, other fluoroquinolones, such as difloxacin or tosufloxacin demonstrated moderate PANX1 inhibitory efficacy, in contrast to ciprofloxacin or levofloxacin, which showed no PANX1 inhibitory effect, indicating the presence of functional moieties of Trovan critical for PANX1 inhibition potency. Trovan consists of a naphthyridone core structure, linked to 2,4-difluorophenyl group at N1 position, carboxylic acid at C3 position, fluoro substituent at C6 position, and cyclopropylamine-fused pyrrolidine at C7 position (**Fig. 1A**). Therefore, to generate specific PANX1 inhibitors of higher efficacy, we introduced 14, 6, or 5 different substituents at N1, C3 and C7 positions of naphthyridone scaffold, respectively, and investigated the structure-activity relationship for a total of 27 different naphthyridone analogues on PANX1 activity (**Fig. 1A**). The general synthetic scheme for all the analogues is shown in **Fig. 1B**. The synthesis procedures of individual compounds, along with their structures and purity verifications through nuclear magnetic resonance (NMR), mass spectrometry, and high-performance liquid chromatography (HPLC), are detailed in the **Supplementary Text** and **Figs. S1-S2**.

**Figure 1.**
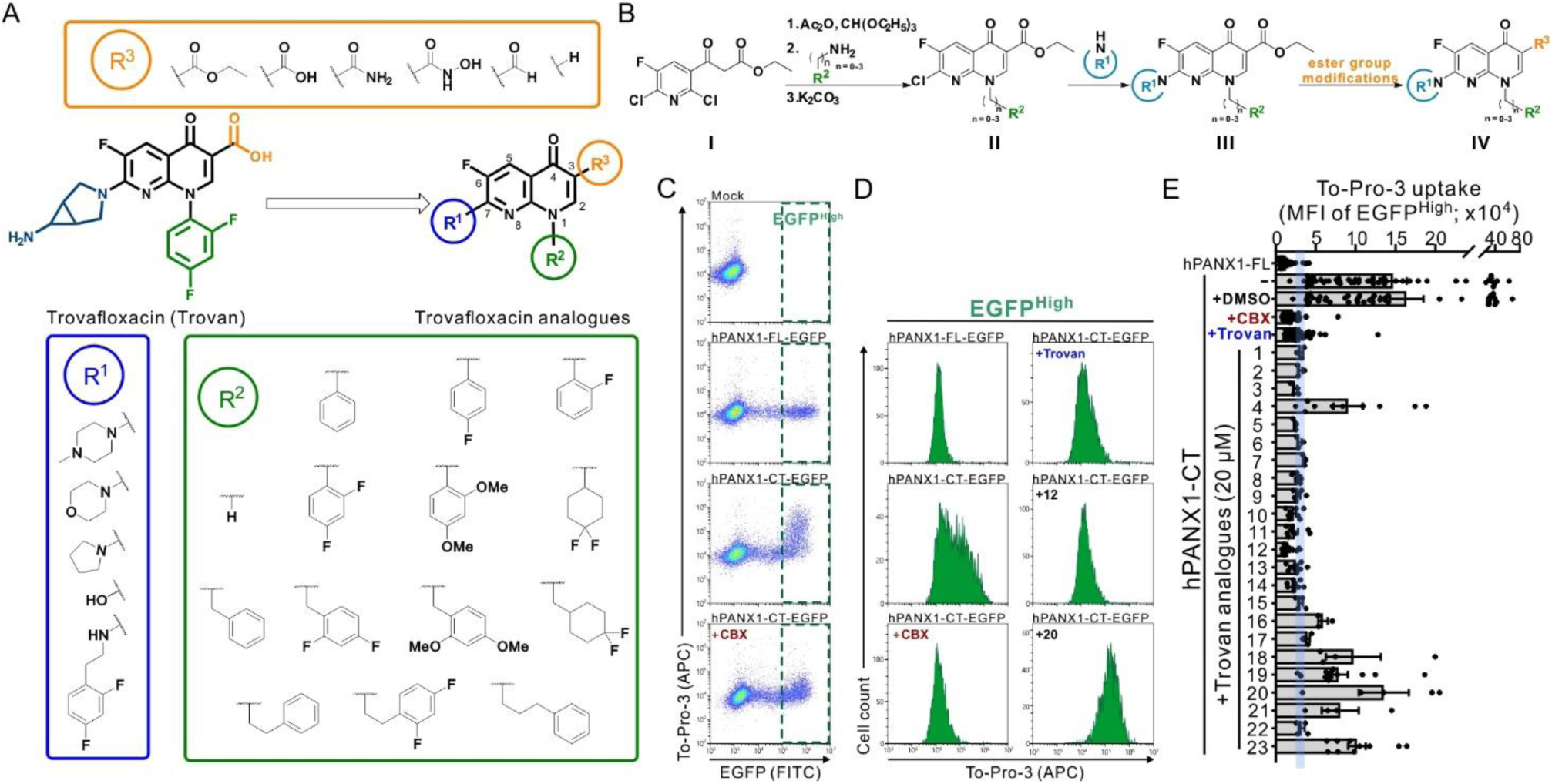
Identification of new PANX1 inhibitors from the newly synthesized naphthyridone analogues. **(A)** Twenty-seven naphthyridone analogues were synthesized by substituting 2,4-difluorophenyl group at N1 position, carboxylic acid at C3 position, and cyclopropylamine-fused pyrrolidine at C7 position of Trovan using varying R^1^, R^2^, or R^3^ groups. **(B)** A general synthesis scheme for naphthyridone analogues. Ethyl 3-oxo-3-(pyridin-3-yl)propanoate **I** was reacted with triethyl orthoformate in the presence of acetic anhydride to form ethyl α-(ethoxymethylene)-propanoate. Amines or anilines listed as R^2^ (from **A**) were sequentially added for condensation to generate enamine intermediates. Addition of K2CO3 to the intermediates led to the core naphthyridone structure **II** through nucleophilic aromatic substitution (SNAr). Various amines specified as R^1^ (from **A**) were then reacted with the chloro group at the C7 position to produce compounds **III**. Finally, we modified the ester group of **III** at the C3 position to yield compounds **IV**. **(C)** Exemplary flow cytometry analyses of To-Pro-3 uptake in HEK293T cells expressing EGFP-tagged, full-length (FL) or C-terminally-truncated (CT) human PANX1 (hPANX1). The green dashed lines demarcate cells with high EGFP signal intensity (EGFP^High^) used for dye uptake analysis. **(D)** Representative histograms showing the To-Pro-3 uptake obtained from the EGFP^High^ cells expressing hPANX1-FL-EGFP or hPANX1-CT-EGFP, including after treatment with carbenoxolone (CBX, 50 μM), Trovan (20 μM) or two naphthyridone analogues, compounds **12** and **20** (20 μM). **(E)** Bar graph showing the mean fluorescence intensity (MFI) of To-Pro-3 uptake obtained from EGFP^High^ cells expressing hPANX1-FL-EGFP or hPANX1-CT-EGFP, with or without application of CBX (50 μM), Trovan (20 μM) or naphthyridone analogues (20 μM). Cyan shaded area indicates 95% confidence interval of Trovan-mediated effects on To-Pro-3 MFI.

Given that the basally-silent human PANX1 can be activated by removing the C-terminal tails, as indicated by the increased permeation to fluorescent dyes (e.g., To-Pro-3 or Yo-Pro-1) (*6, 7, 23, 26*), we employed To-Pro-3 uptake assays to determine whether the newly-synthesized compounds can inhibit PANX1-mediated dye uptake. We transiently transfected EGFP-tagged, C-terminally truncated human PANX1 (hPANX1-CT-EGFP) in HEK293T cells and assessed the inhibitory effects of various naphthyridone analogues on To-Pro-3 uptake using flow cytometry (**Fig. 1C** and **Fig. S3)**. We compared the mean fluorescence intensity (MFI) of To-Pro-3 in the cells showing high MFI of EGFP (EGFP^High^) as the cell populations with a higher level of PANX1 expression (**Fig. 1C** and **D**). Similar to the previous reports (*7, 23, 27*), the wild-type, full-length human PANX1 (hPANX1-FL-EGFP) displayed a minimal To-Pro-3 uptake, in contrast to the C-terminally-truncated human PANX1 (hPANX1-CT-EGFP), which demonstrated a significantly higher To-Pro-3 MFI that was reduced by applying carbenoxolone (CBX; 50 μM) or trovafloxacin (Trovan; 20 μM) (**Fig. 1E**). The baseline PANX1-mediated To-Pro-3 uptake was determined as the difference of To-Pro-3 MFI from cells expressing hPANX1-CT-EGFP relative to control cells expressing hPANX1-FL-EGFP, with the level of PANX1 inhibition by naphthyridone analogues expressed as a percentage of that baseline To-Pro-3 uptake (**Table 1**).

**Table 1:**
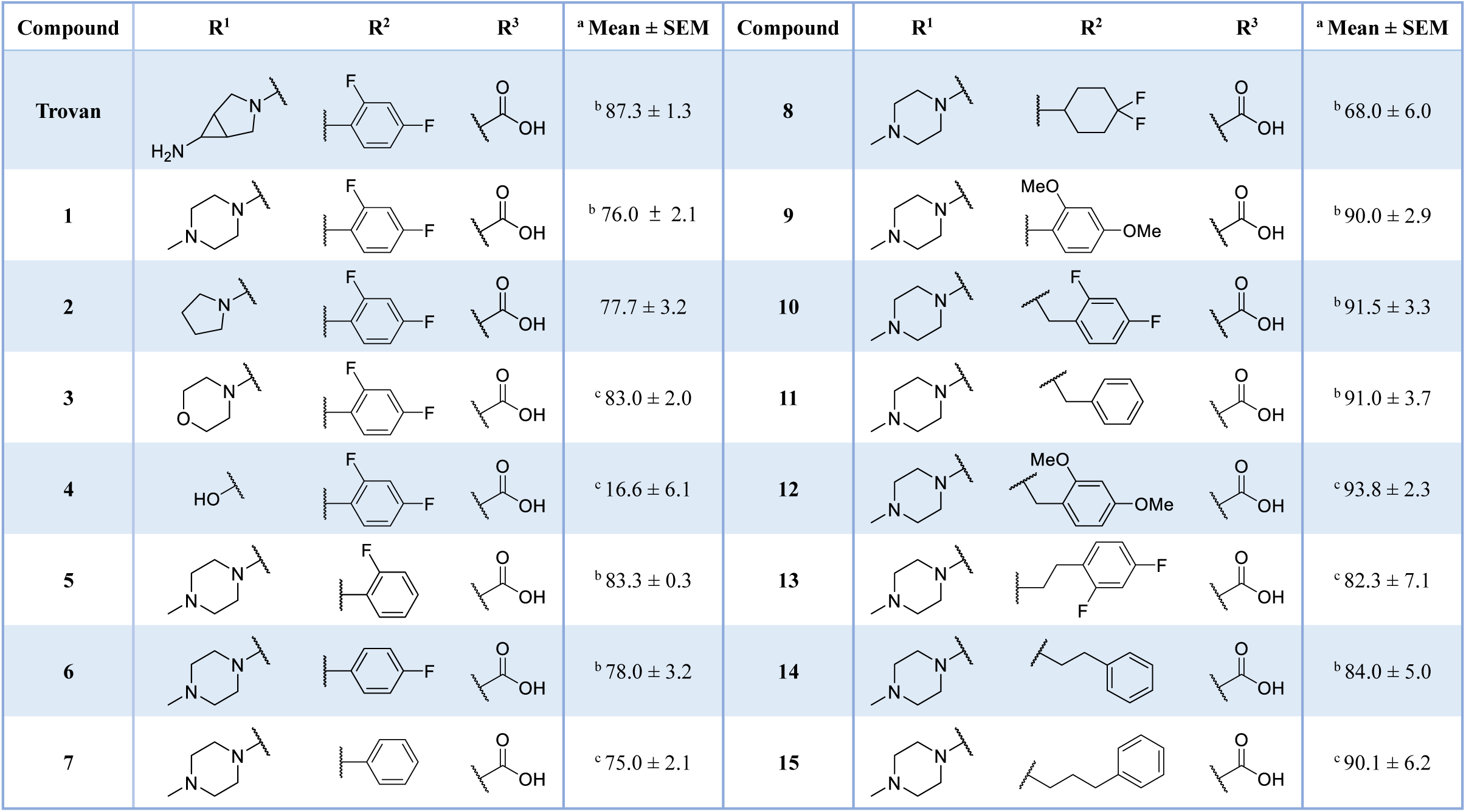
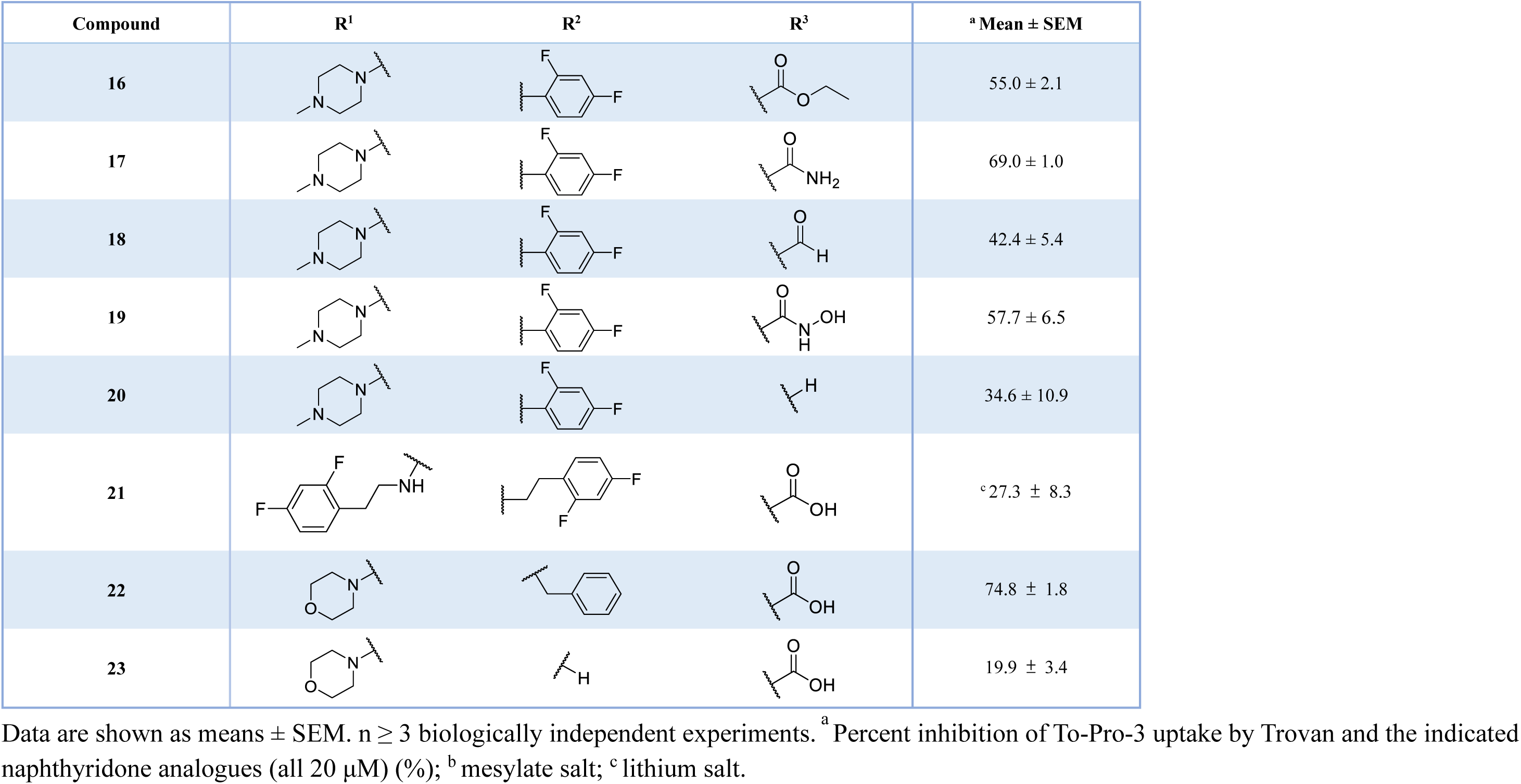
Inhibition of PANX1-dependent To-Pro-3 uptake by Trovan and analogues.

To reduce potential hepatotoxicity presumably resulting from the transformation of cyclopropylamine-fused pyrrolidine to an unstable, unsaturated α-β aldehyde at C7 position of Trovan (*28*), we simplified the fused-ring with morpholine (**1**), methylpiperazine (**2**), pyrrolidine (**3**) or a smaller hydroxyl group (**4**). By examining different naphthyridone analogues at a concentration of 20 μM using To-Pro-3 uptake assays, we found that compounds **1** (76.0 ± 2.1%), **2** (77.7±3.2%), and **3** (83.0±2.0%) showed inhibition of dye uptake comparable to Trovan (87.3 ± 1.3%), whereas compound **4** was less effective (16.6 ± 6.1%) (**Table 1**). This suggests that a sizable group at R^1^ position is required for a substantial inhibition of PANX1 activity. Additionally, compounds **5** (83.3 ± 0.3%), **6** (78.0 ± 3.2%), **7** (75.0 ± 2.1%), and **8** (68.0 ± 6.0%) also showed inhibition of To-Pro-3 uptake similar to Trovan, despite the differences in the arrangement of fluoro atoms on a phenyl or cyclohexyl at R^2^ position (**Table 1**). By replacing the difluorophenyl group at N1 position of Trovan with an electron-donating dimethoxy group, we found that compound **9** showed a substantially enhanced inhibition of To-Pro-3 uptake (90.0 ± 2.9%) compared with other analogues (**Table 1**).

By comparing compounds with different R^3^ groups, we also found that compounds **16** (55.0 ± 2.1%), **17** (69.0 ± 1.0%), **18** (42.4 ± 5.4%), and **20** (34.6 ± 10.9%) demonstrated a lower percent inhibition of PANX1-mediated To-Pro-3 uptake than compound **1 (Table 1**). We also noted a higher percent inhibition by compound **1** (76.0 ± 2.1%) than compound **19** (57.7 ± 6.5%), in which R^3^ substituent is hydroxamic acid, a bioisostere of carboxylic acid. These results indicated that the carboxylic acid at R^3^ substituent plays a critical role for PANX1 inhibition. Moreover, by comparing different naphthyridone analogues with the R^2^ substituents comprising varying numbers of carbons (ranging from 1 to 3) extended from the naphthyridone core, we noted an increase in inhibition of To-Pro-3 uptake for compounds **10** (91.5 ± 3.3%), **11** (91.0 ± 3.7%), **12** (93.8 ± 2.3%), **13** (82.3 ± 7.1%), **14** (84.0 ± 5.0%), and **15** (90.1 ± 6.2%), when compared to Trovan or compound **1**, which does not present a carbon extension. Among these, compound **12** possesses a one-carbon extension and electron donating dimethoxy group at R^2^ position, and represents the most effective analogue based on inhibition of PANX1-dependent To-Pro-3 uptake (**Table 1**). Moreover, given that compounds **22** (74.8 ± 1.8%) and **23** (19.9 ± 3.4%) showed a lower percent inhibition of To-Pro-3 uptake than compound **11** (91.0 ± 3.7%), our data further demonstrated that methylpiperazine at R^1^ position presented better PANX1 inhibitory effect than other substituents (e.g., morpholine) but that the presence of a spatial occupying R^2^ substituent remain important for PANX1 inhibition.

### Compound 12 showed a superior inhibitory efficacy on PANX1 channels than other naphthyridone analogues

To further examine the dose-dependent effect of compound **12** on PANX1 channel activities, we carried out the whole-cell voltage-clamp recordings in HEK293T cells expressing hPANX1(TEV)-EGFP, an GFP-tagged PANX1 construct in which the caspase cleavage site (IKMDVVD) was replaced by the TEV protease recognition sequence (ENLYFQG) (*26, 29*). Consistent with previous reports (*26*), cleavage of PANX1 C-terminal tails in cells co-expressing TEV protease (TEVp) resulted in outwardly rectifying currents sensitive to CBX (50 μM; **Fig. 2A**). We found that bath application of compound **12** dose-dependently reduced whole-cell PANX1 currents across a broad range of membrane potentials (ranging from -130 mV to +80 mV; **Fig. 2A**). Like Trovan (*23*), inhibition of PANX1 by compound **12** was voltage-dependent, with greater percent inhibition at negative membrane potentials (e.g., -50 mV) than at positive membrane potentials (e.g., +80 mV) (**Fig. 2A** and **B**). By analyzing the percent inhibition of PANX1 whole-cell currents at -50 mV, we found that compound **12** showed an IC_50_ of ∼0.73 μM (**Fig. 2C**), lower than the IC_50_ of Trovan (∼4 μM) as previously reported (*23*).

**Figure 2.**
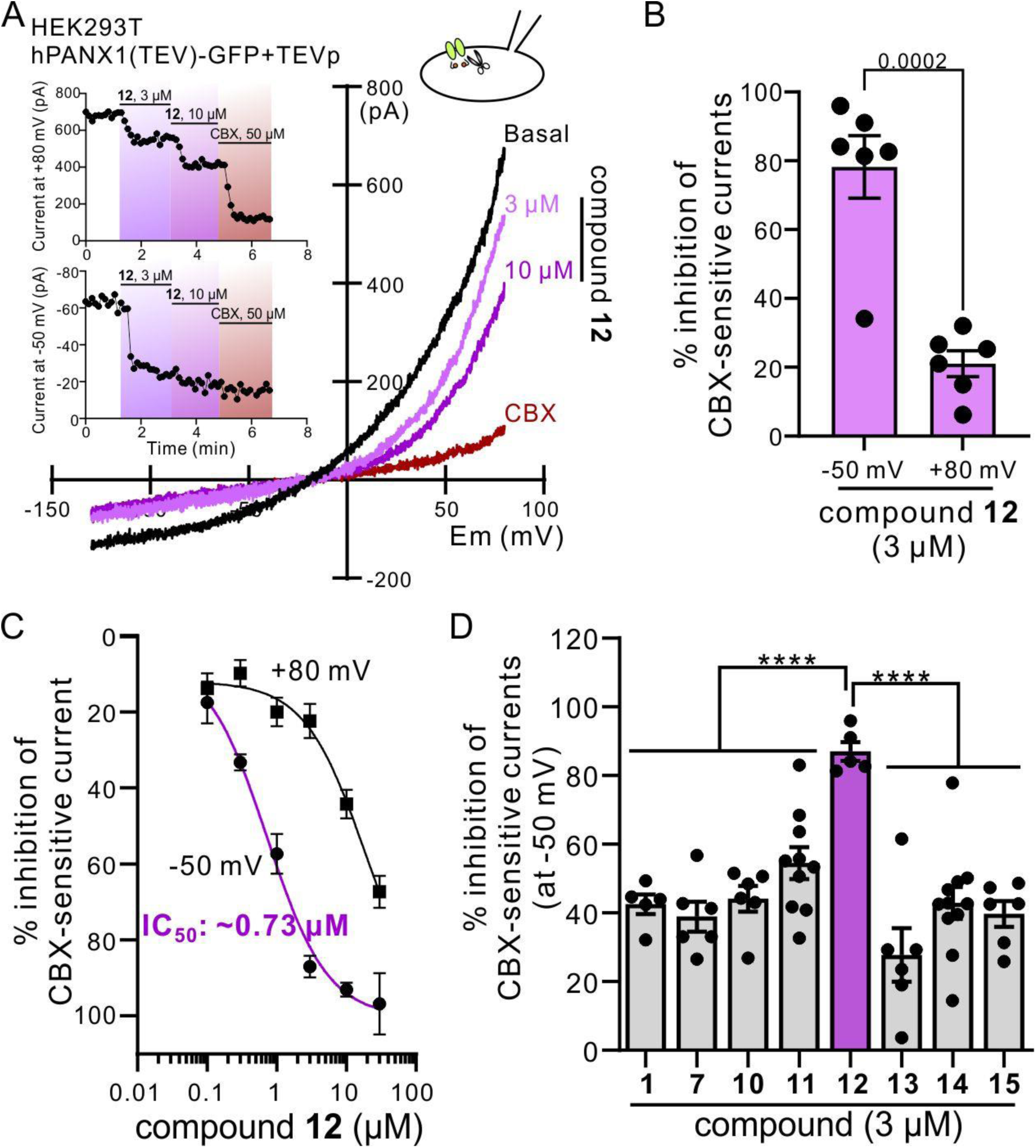
Compound 12 demonstrated a dose-dependent inhibition on PANX1 currents. **(A)** Exemplar whole-cell current-voltage relationships obtained from a HEK293T cell expressing hPANX1(TEV)-EGFP and TEV protease (TEVp). The basal current (black) was reduced by bath application of compound **12** (3 or 10 μM) or CBX (50 μM). Inset: Corresponding time course of whole-cell current amplitudes obtained at either +80 mV (upper) or -50 mV (lower). No CBX-sensitive current was detected in HEK293T cells only expressing hPANX1(TEV)-EGFP. **(B)** Compound **12** showed a voltage-dependent inhibition on the PANX1 currents, where the percent inhibition of PANX1 currents at -50 mV is significantly greater than that at +80 mV. n = 6 individual cells. *P* = 0.0002 by two-tailed, unpaired t test. **(C)** IC_50_ of compound **12** is ∼0.73 μM based on whole-cell recordings at -50 mV (n=5∼6 cells). **(D)** At 3 μM, compound **12** showed a significantly higher percent inhibition of PANX1 currents at -50 mV than other compounds. n= 5∼11 cells per group. ****: *P* ≤ 0.0001 using one-way ANOVA with Bonferroni test. All data are shown as means ± SEM.

Inhibition of PANX1 currents by compound **12** at -50 mV was nearly complete at 3 μM, with little inhibitory effect on currents obtained at +80 mV. Therefore, we compared inhibition by other select naphthyridone analogues on PANX1 currents at 3 μM and -50 mV. We found that compound **12** demonstrated a significantly higher percent inhibition on PANX1 currents (87.0 ± 2.8%; **Fig. 2D**) than other analogues that showed similar inhibition of To-Pro-3 uptake (**Table 1**). It is also worth noting that compound **11** (54.5 ± 4.6%) with a phenyl group and one extended carbon at R^2^ position showed greater inhibition than compounds **7** (38.9 ± 4.3%), **14** (42.9 ± 4.7%), and **15** (39.7 ± 3.7%) with a phenyl group and 0, 2, and 3 carbon extension at R^2^ position, respectively (**Table 2**). Similarly, compound **10** (44.1 ± 3.7%) with a 2,4-difluorophenyl group and one extended carbon at R^2^ position also demonstrated a higher percent inhibition on PANX1 currents than compounds **1** (38.9 ± 4.3%) and **13** (27.7 ± 7.8%) at a concentration of 3 μM (**Table 2**). These data indicate that an aromatic ring and one extended carbon at R^2^ position could lead to an enhanced inhibitory efficacy against PANX1 channels.

**Table 2:**
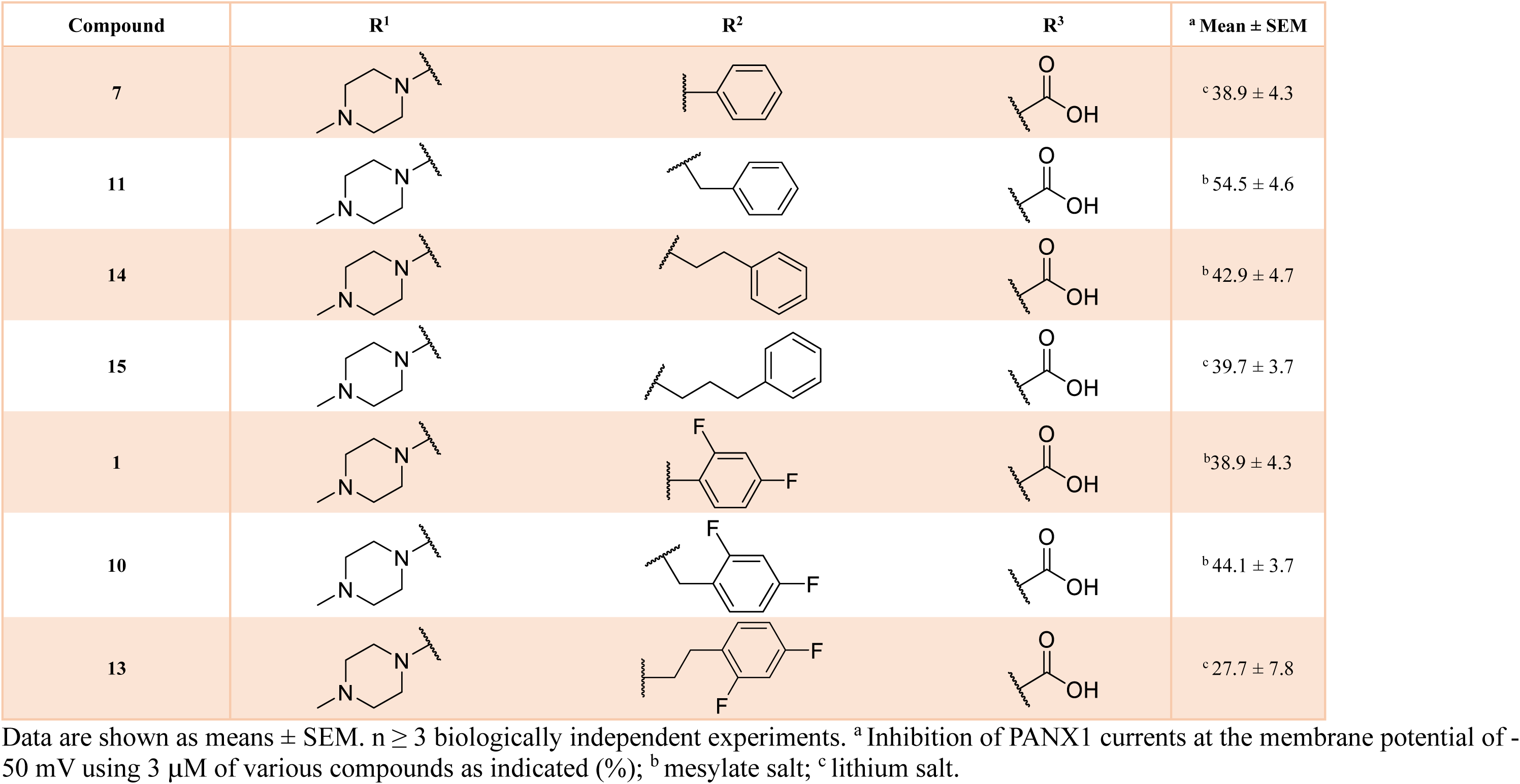
Percent inhibitions of PANX1 currents at -50 mV among different compounds.

To further examine the SAR of compound **12** as a PANX1 inhibitor, we performed To-Pro-3 uptake assays and found that compound **24** with a saturated 4,4-difluorocyclohexyl group and one extended carbon at R^2^ position was less potent in hindering PANX1-mediated To-Pro-3 uptake than compound **12** (**Table 3**; **12**: 76.2 ± 6.0% at 2.5 μM and 48.1 ± 4.9% at 1 μM *cf.* **24**: 60.3 ± 6.6% at 2.5 μM and 15.3 ± 10.8% at 1 μM). Moreover, by exchanging the carboxylic acid at R^3^ position with an ester, we consistently found that compounds with a carboxylic acid at R^3^ position showed superior inhibitory efficacy on PANX1-mediated To-Pro-3 uptake (**Table 3**). For example, compounds **24**, **9**, and **12** showed greater percent inhibition than **25**, **26**, and **27**, respectively, regardless of varying substituents at R^2^ position. This finding suggests that the ability of carboxylic acid as hydrogen bondings may contribute to the enhanced inhibitory potency of compound **12** on PANX1 channels. We also observed that compound **12** had a higher percent inhibition of To-Pro-3 uptake than compound **11** (**Table 3**; **12**: 76.2 ± 6.0% at 2.5 μM and 48.1 ± 4.9% at 1 μM *cf.* **11**: 50.5 ± 0.9% at 2.5 μM and 23.6 ± 2.7% at 1 μM), implicating that the dimethoxy group with electron-donating ability at R^2^ position of compound **12** might be also accounted for its enhanced efficacy. Furthermore, compounds **24** and **12** also rendered a higher percent inhibition of To-Pro-3 uptake than compounds **8** and **9**, respectively (**Table 3**), further supporting the hypothesis that extending one carbon chain at R^2^ position improved the inhibition efficacy regardless of the presence or absence of an aromatic ring.

**Table 3:**
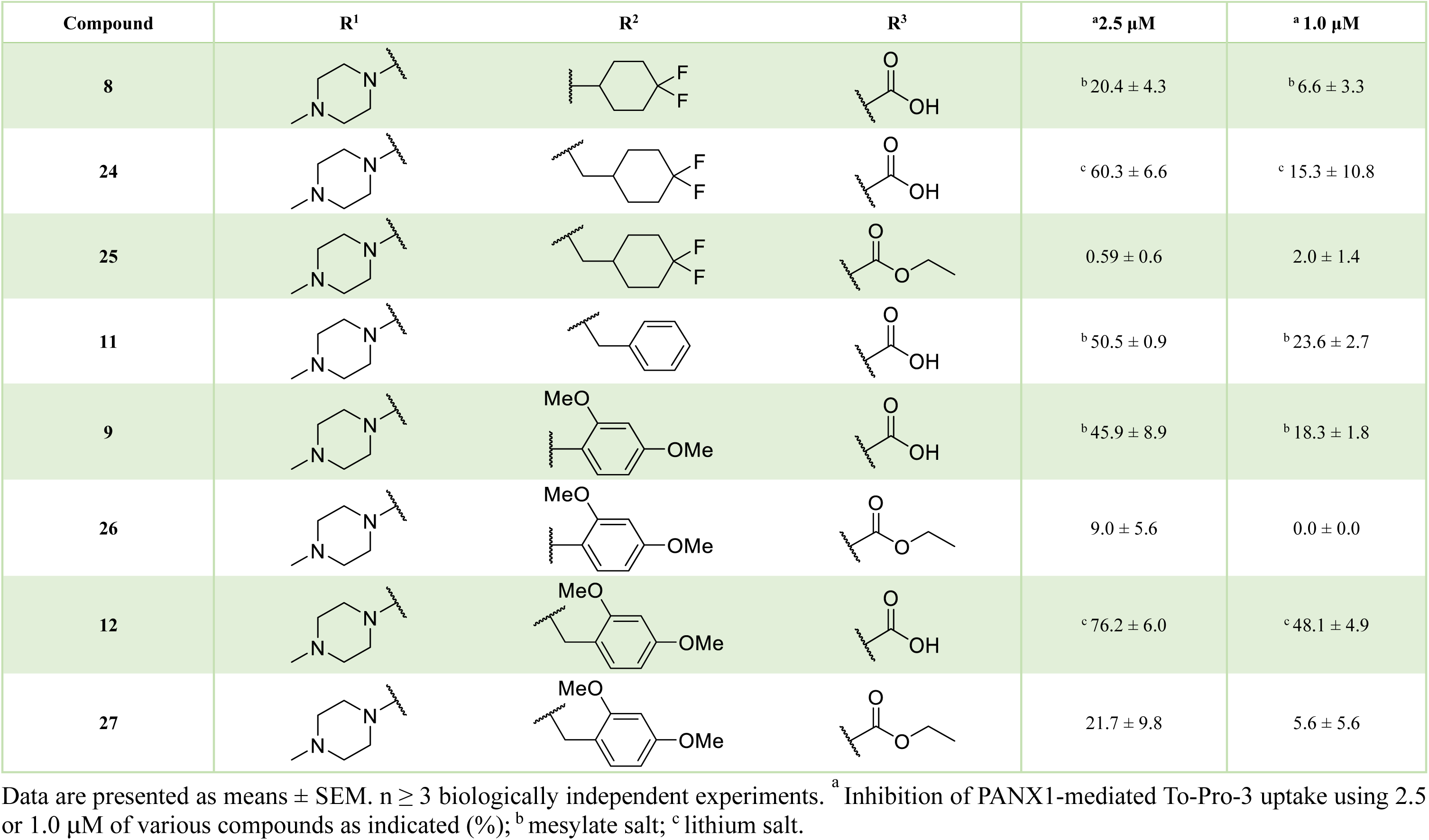
Inhibition of PANX1-mediated To-Pro-3 uptake by lower concentrations of naphthyridone analogues.

### Extracellular Trp74 of hPANX1 is a critical residue for compound 12-mediated PANX1 inhibition

By using mutagenesis and cryogenic electron microscopic (cryo-EM) analyses, previous studies suggested that the first extracellular loop of PANX1, particularly Trp74 residue, could be the binding site on PANX1 for carbenoxolone and probenecid (*30, 31*). To examine whether Trp74 also served as an interacting site for compound **12**, we performed whole-cell recording in HEK293T cells expressing a mutant hPANX1(TEV), where Trp74 residue was replaced by alanine (hPANX1(TEV)-W74A) (**Fig. 3A-D**). In cells co-expressing hPANX1(TEV)-W74A and TEVp, we observed an outwardly rectifying whole-cell current with a current-voltage (I-V) relationship similar to that of wildtype hPANX1(TEV) activated by TEVp-mediated C-tail cleavage (**Fig. 3A** *cf.* **Fig. 2A**). Cells were first superfused with compound **12** (20 μM), washed, and then exposed to CBX (50 μM) (**Fig. 3B**). As previously reported (*30, 31*), CBX did not inhibit hPANX1(TEV)-W74A currents, consistent with the proposed role for Trp74 in binding CBX (**Fig. 3A-C**). Interstingly, PANX1 currents were likewise unaffected by compound **12** at -50 mV (**Fig. 3A,B,D**) with a negligible, albeit significant reduction in the whole-cell current density found at +80 mV (**Fig. 3D**, upper). These data suggest that Trp74 could also be a critical PANX1 residue interacting with compound **12**.

**Figure 3.**
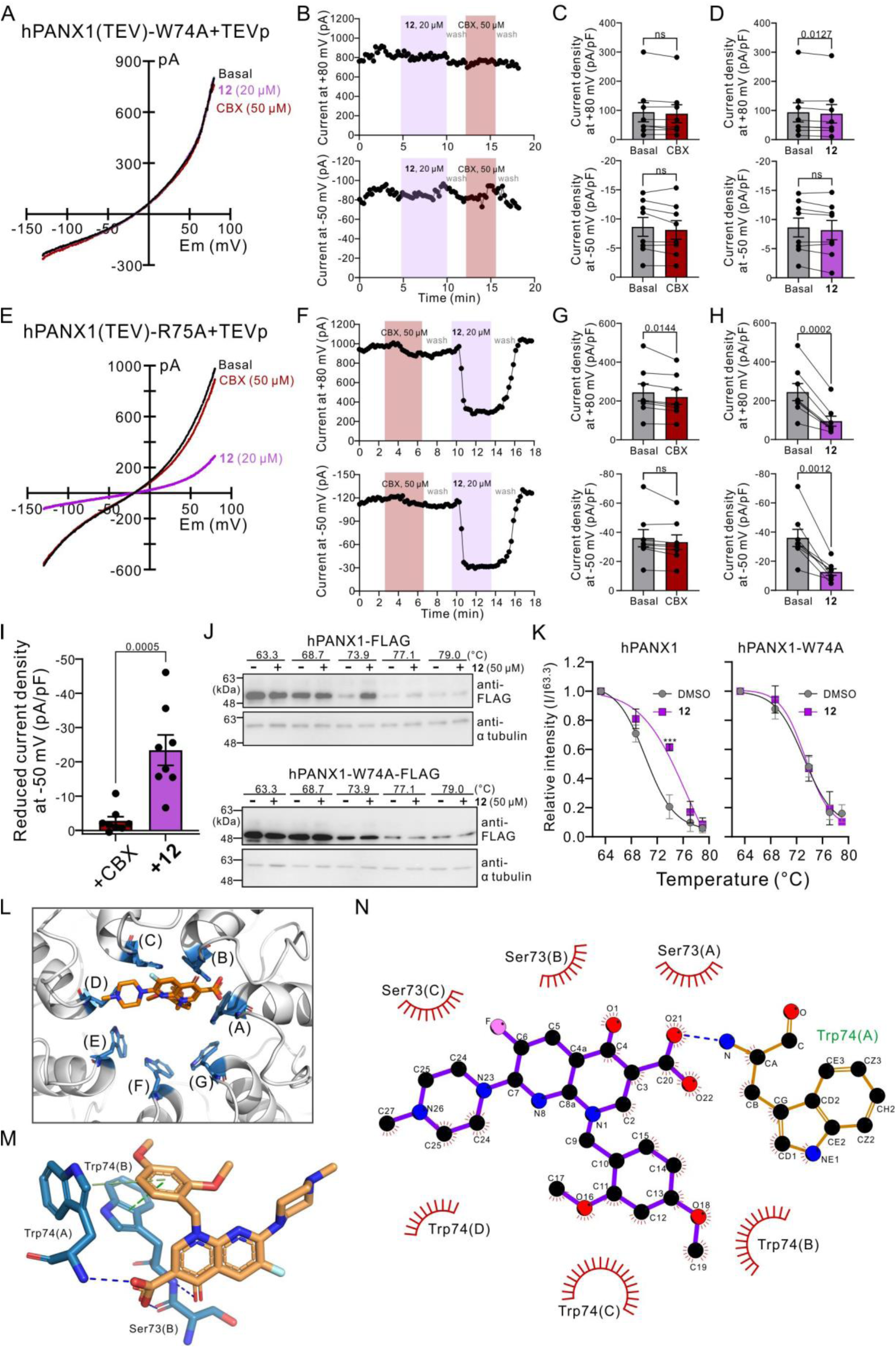
Trp74 is a critical residue for PANX1-compound 12 interactions. **(A)** Representative whole-cell I-V relationships of HEK293T cells co-expressing hPANX1(TEV)-W74A and TEVp, under basal conditions and in the presence of CBX (50 μM) or compound **12** (20 μM). **(B)** Corresponding time series (from **A**) of whole-cell current amplitudes at +80 mV (upper) and -50 mV (lower). **(C,D)** Summary results (n=8 cells) of whole-cell current density before or after application of CBX **(C)** or compound **12 (D)**. *P* values derived from two-tailed, unpaired t test are indicated (n=8 cells). ns: not significant. **(E)** Representative whole-cell I-V relationships of a HEK293T cell co-expressing hPANX1(TEV)-R75A and TEVp, with or without the presence of CBX (50 μM), compound **12** (20 μM). **(F)** Corresponding time series (from **E**) of whole-cell current amplitudes at +80 mV (upper) and -50 mV (lower). **(G,H)** Summary results (n=8 cells) of whole-cell current density before or after application of CBX **(G)** or compound **12 (H)**. **(I)** Inhibition of TEVp-activated hPANX1(TEV)-R75A by compound **12** was greater than by CBX. Data are presented as means ± SEM. *P* values derived from two-tailed, unpaired t tests are as indicated (n=8 cells). ns: not significant. **(J)** Representative immunoblots of cell thermal shift assays (CETSA) obtained from HEK293T cells expressing either hPANX1-FLAG (upper) or hPANX1-W74A-FLAG (lower), with or without compound **12** (50 μM), at indicated temperatures. Αnti-α-tubulin was used as a loading control. **(K)** Grouped results showing the intensity of immunoreactive signals from hPANX1 (left) or hPANX1-W74A (right) at different temperatures, relative to that at 63.3℃. ***: *P*≤0.001 using two-way ANOVA with Bonferroni’s test (n=3 biologically independent experiments). Curves represent the fitted results using a sigmoidal model. **(L)** Diagram presents the extracellular view of a top-ranked PANX1-compound **12** docking model proposed by using PyRx program. Trp74 residues from 7 PANX1 subunits (A-G) are highlighted in blue. **(M)** Three hydrogen bonds (blue dashed lines) and 2 π-stacking (green dashed lines) are predicted by using Protein-Ligand Interaction Profiler (PLIP). **(N)** Schematics show one hydrogen bond (blue dashed lines) and several hydrophobic interactions (eyebrow-like icons) between PANX1 and compound **12** predicted by using LigPlot+.

Recent cryo-EM studies have suggested that a halo of Trp74 residues presents the narrowest constriction of channel permeation vestibule and that the neighboring Arg75 residue may serve as a selectivity filter (*31, 32*), rendering an anion-favoring permeation profile of PANX1 channels (*6, 30*). Additionally, cryo-EM maps also suggested that a π-cation interaction between Trp74 and Arg75 residues from two adjacent PANX1 subunits serves to orient the Trp74 side chains (*31, 33*). Using whole-cell recordings, we found that HEK293T cells co-expressing TEVp and a mutant hPANX1(TEV) with an Arg75-to-Ala75 substitution (hPANX1(TEV)-R75A) showed a I-V relationship that rectifies weakly in both the outward and inward directions, distinct from the outwardly rectifying I-V relationship of wildtype hPANX1(TEV) channels (Fig. **3E** *cf.* **Fig. 2A**). For these TEVp-activated hPANX1(TEV)-R75A channels, bath application of CBX modestly reduced currents at +80 mV with no effect at -50 mV (**Fig. 3E-G**). This is consistent with previous reports suggesting that Arg75 residues might also be involved in CBX-PANX1 interactions (*30, 31*). On the other hand, bath application of compound **12** markedly reduced currents mediated by these C-tail-cleaved hPANX1(TEV)-R75A channels across a broad range of the membrane potentials (**Fig. 3E,F,H**). That, unlike the virtual loss of CBX sensitivity, not only was channel inhibition by compound **12** retained in these hPANX1(TEV)-R75A channels but it now extended to positive potentials and outward currents. Note also that hPANX1(TEV)-R75A demonstrated a significantly greater sensitivity to compound **12** than CBX (**Fig. 3I**). These results together suggest that Trp74 residue of PANX1, but not Arg75, could be pivotal for the interaction between PANX1 and compound **12**.

We further carried out the cellular thermal shift assay (CETSA) (*34, 35*) to examine whether Trp74 residues of PANX1 are crucial for engaging compound **12** in a physiological environment. HEK293T cells expressing FLAG-tagged wildtype hPANX1, hPANX1-W74A, or hPANX1-R75A were treated with compound **12** (50 μM), heated at different temperatures, and lyzed, and the protein samples were separated by using SDS-PAGE (**Fig. 3J; Fig. S4**). We found that compound **12**-exposed wildtype hPANX1 showed a stronger immunoblotting signal intensity at 73.9℃ than those without compound **12** treatment (**Fig. 3J**, upper), indicating an increase in thermal stability of hPANX1 upon compound **12** treatment. However, exposure of hPANX1-W74A to compound **12** produced no noticeable change in signal intensity at any of the temperatures tested (**Fig. 3J**, lower). Meanwhile, exposure of hPANX1-R75A to compound **12** produced a stronger signal intensity at 68.7℃ and 73.9℃ (**Fig. S4**), similar to the wildtype hPANX1 and consistent with the preserved inhibition by compound **12** of the R75A channel. By fitting the relative signal intensity of hPANX1 at various temeraptures using a sigmoidal model, we found that compound **12** increased thermal stability of wildtype hPANX1, raising its apparent aggregation temperature (T_agg_) (*34*) from 70.3℃ to 77.0℃ (**Fig. 3K, left**), without affecting the thermal stability of hPANX1-W74A (**Fig. 3K, right**; T_agg_ of DMSO: 73.2℃; T_agg_ of compound **12**: 73.4℃). Thus, both electrophysiological and biochemical evidence support Trp74 as an essential residue for PANX1 to engage compound **12**.

Moreover, we performed a molecular docking analysis using PyRx software and AutoDock Vina program (*36*) to evaluate potential binding conformations of compound **12** and PANX1 based on cryo-EM strucutres retrieved from the Protein Data Bank (PDB; 6WBI) (*31*). Three-dimentional (3D) diagrams of the predicted models were recreated using PyMol software (**Fig. 3L**). We also used the Protein-Ligand Interaction Profiler (PLIP) (*37*) and LigPlot+ (*38*) to visualize 3D (**Fig. 3M; Movie S1**) and 2D (**Fig. 3N**) diagrams of compound **12** interactions with PANX1. The top-ranked model predicted the formation of 3 hydrogen bondings and 2 π-stacking interactions (**Fig. 3M**), as well as multiple hydrophobic interactions (**Fig. 3N**), between compound **12** and PANX1. Particularly, carboxylic acid at R^3^ position of compound **12** was suggested to form two hydrogen bonds with Trp74 and Ser73 from the neighboring subunit (**Fig. 3M**, blue lines). The presence of another hydrogen bond was predicted between the naphthyridone core of compound **12** and a Ser73 residue (**Fig. 3M**). Additionally, two π-stacking interactions were proposed between the indole groups of two adjacent Trp74 residues and the dimethoxybenzyl group at R^2^ position of compound **12** in either parallel or perpendicular fashion (**Fig. 3M**, green dashlines). The proposed model is aligned with our experimental data showing that Trp74 residue of PANX1 indeed represents a critical binding site of compound **12** and that dimethoxybenzyl group at R^2^ position and carboxylic acid at R^3^ position profoundly contribute to the enhanced PANX1 inhibitory efficacy of compound **12**.

### Compound 12 selectively inhibits PANX1 without affecting the homologous LRRC8 channels

Pannexins share the amino acid sequence homology and transmembrane topology with LRRC8 (LRRC8A∼E)(*39*), which constitutes the volume-regulated anion channels (VRACs) activated by a lowered ionic strength (*40–42*). In light of the similar structural features between PANX1 and LRRC8A channels revealed by recent cryo-EM analyses (*32, 43*), we examined a potential off-target effect of compound **12** on LRRC8/SWELL1 channels. We found that wild-type HEK293T cells take up the negatively-charged sulfo-Cy5 dye when exposed to a hypotonic solution (125 mOsm), indicating an activation of VRACs (**Fig. 4A**). Additionally, we observed a similarly increased sulfo-Cy5 uptake in either Cas9-expressing (**Fig. 4B**) or PANX1-deleted (**Fig. 4C; Fig. S5**) HEK293T cells stimulated by hypotonic solutions, indicating that PANX1 is not responsible for this uptake of sulfo-Cy5. Note that neither Trovan nor compound **12** inhibited the hypotonicity-induced sulfo-Cy5 uptake whereas it was blocked by two physicochemically distinct VRAC inhibitors, carbenoxolone (CBX) and dicoumarol (Dic) (*41, 44*) (**Fig. 4A-C**). Note also that dicoumarol does not inhibit To-Pro-3 uptake mediated by the C-tail-cleaved PANX1 channels (**Fig. S6**). Furthermore, by using HEK293T cells expressing an iodide-sensing yellow fluorescence protein (YFP-H148Q/I152L) (*45*), we demonstrated that the hypotonicity-induced influx of inorganic iodide was deminished by CBX or dicoumarol, but not by Trovan or compound **12** (**Fig. 4D**). These results indicate that Trovan and compound **12** render no off-target effect on the PANX1-related LRRC8/SWELL1 channels and that PANX1 is not involved in the cell volume regulation following changes in osmolarity.

**Figure 4:**
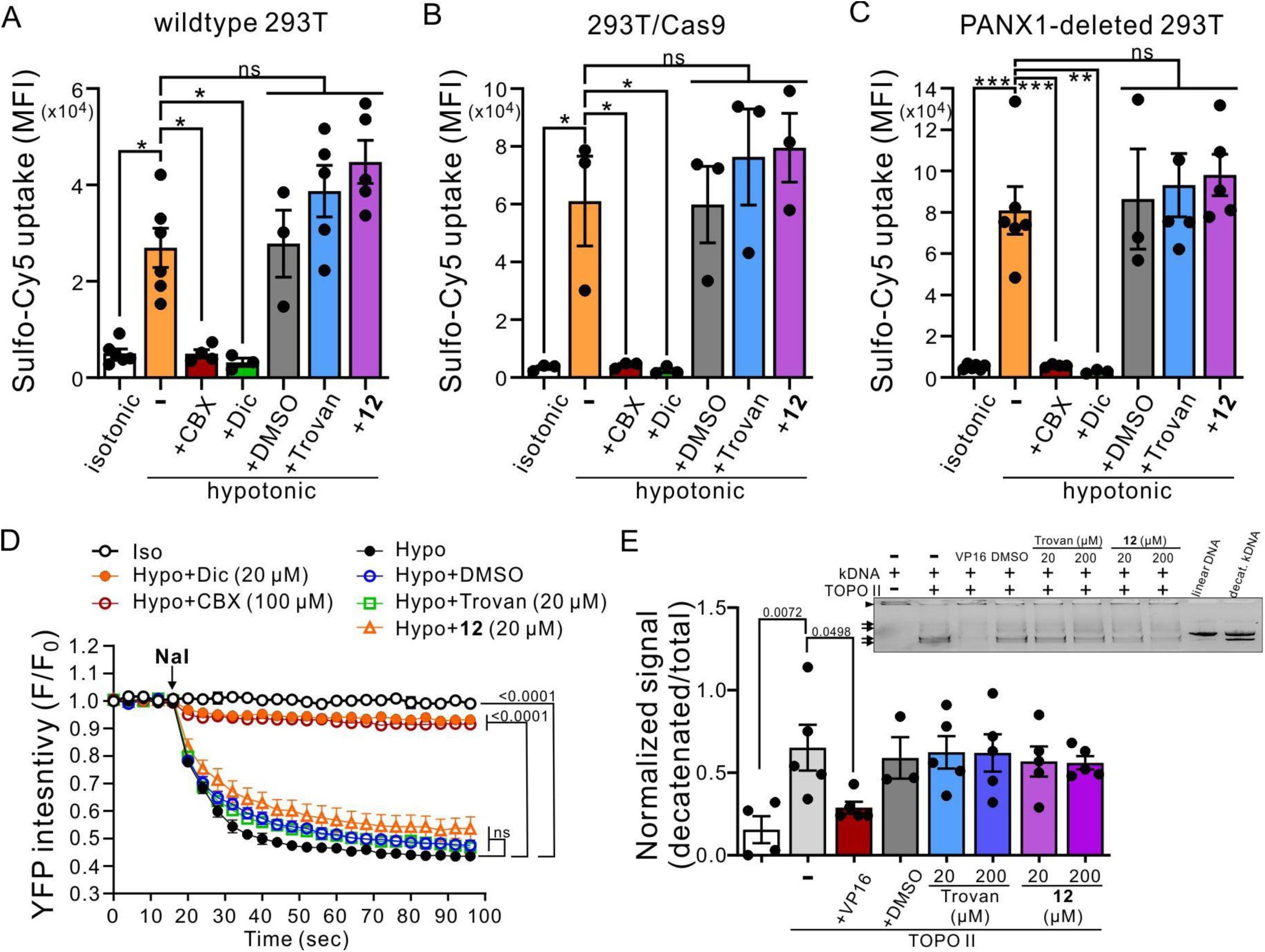
Compound 12 does not inhibit LRRC8/SWELL1 channels or human topoisomerase II. **(A-C)** Hypotonic (125 mOsm)-induced sulfo-Cy5 uptake was assessed in wild-type **(A)**, Cas9-expressing **(B)** or PANX1-deleted **(C)** HEK293T cells and was reduced by treatments of CBX (50 μM) or dicoumarol (Dic; 20 μM), but not Trovan or compound **12** (20 μM). *: *P* ≤ 0.05, **: *P* ≤ 0.01, ***: *P* ≤ 0.001, ns: not significant by one-way ANOVA followed by Bonferroni’s test (n=3∼6 independent experiments). **(D)** Averaged YFP signal intensity of HEK293T cells expressing YFP-H148Q/I152L in isotonic (Iso; 310 mOsm) or hypotonic (Hypo; 125 mOsm) solutions in response to NaI (10 mM), with or without a pre-incubation with dicoumarol (Dic), CBX, Trovan or compound **12**. *P* values derived from two-way ANOVA with Dunnett’s test are shown (n=4 biologically independent experiments). **(E)** Normalized intensity of decatenated DNA showing that a topoisomerase inhibitor, VP16 (0.1 mM), but neither Trovan nor compound **12** (20 or 200 μM), reduced signal intensity of decatenated DNA as an indication of inhibition on human topoisomerase II. *P* values derived from one-way ANOVA with Dunnett’s test (n=3∼5 independent experiments) were indicated. Inset: Exemplar results of topoisomerase II assay. Arrowhead: catenated DNA. Arrows: decatenated DNA.

While Trovan was previously used clinically as an antibiotic, it was withdrawn from the market due to incidents of serious hepatotoxicity (*25*). The hypothesis regarding why Trovan leads to idiosyncratic drug-induced liver injury is that Trovan non-specifically inhibits topoisomerase II in the liver (*46*). Therefore, we carried out topoisomerase assays and found that both Trovan and compound **12** have negligible effects on the activity of human topoisomerase II *in vitro* at concentrations of either 20 μM or 200 μM, as indicated by a similar level of decatenated DNA in the presence or absence of Trovan or compound **12** (**Fig. 4E**). Therefore, the newly-synthesized compound **12** represents a specific PANX1 inhibitor with a largely enhanced potency compared with Trovan and various other analogues.

### Compound 12 effectively inhibits native PANX1 channels

Previous studies have shown that caspase 3/7-mediated cleavage of PANX1 C-termini during apoptosis can lead to irreversibly activated channels capable of permeating organic fluorescent dyes (e.g., Yo-Pro-1 and To-Pro-3) and mediating ATP release (*7, 23*). To examine whether compound **12** can inhibit PANX1 channels in a native condition, we performed Yo-Pro-1 uptake assays in apoptotic Jurkat cells, with or without endogenously expressed PANX1 channels (**Fig. 5A-C**). From both wildtype and Cas9-expressing Jurkat cells, we found that UV irradiation resulted in a higher mean fluorescence intensity (MFI) of Yo-Pro-1 in apoptotic cells (Annexin V^+^/7-AAD^-^) than in the live cells (Annexin V^-^/7-AAD^-^) without UV exposure (**Fig. 5A** and **B**; **Fig. S7**). We also found that compound **12** (20 μM) significantly dampened Yo-Pro-1 uptake in apoptotic wild-type and Cas9-expressing Jurkat cells to a similar level of two different PANX1 inhibitors, CBX (50 μM) or Trovan (20 μM) (**Fig. 5A** and **B**), demonstrating that compound **12** can indeed inhibit native caspase-activated PANX1 channels. Meanwhile, we found no increase in Yo-Pro-1 uptake in apoptotic Jurkat cells from which PANX1 had been deleted by using CRISPR-Cas9 techniques (*47*) (**Fig. 5C**), even though we found comparable levels of cell death in wildtype, Cas9-expressing, and PANX1-deleted Jurkat cells following UV irradiation (**Fig. 5D-F**). In the PANX1-deleted Jurkat cells, compound **12** showed no additional effect on the Yo-Pro-1 uptake (**Fig. 5C**). These results not only confirmed that Yo-Pro-1 uptake of apoptotic Jurkat cells is mediated by PANX1 channels (*7*) but also indicate that compound **12** can effectively and specifically inhibit the activity of native PANX1 channels.

**Figure 5.**
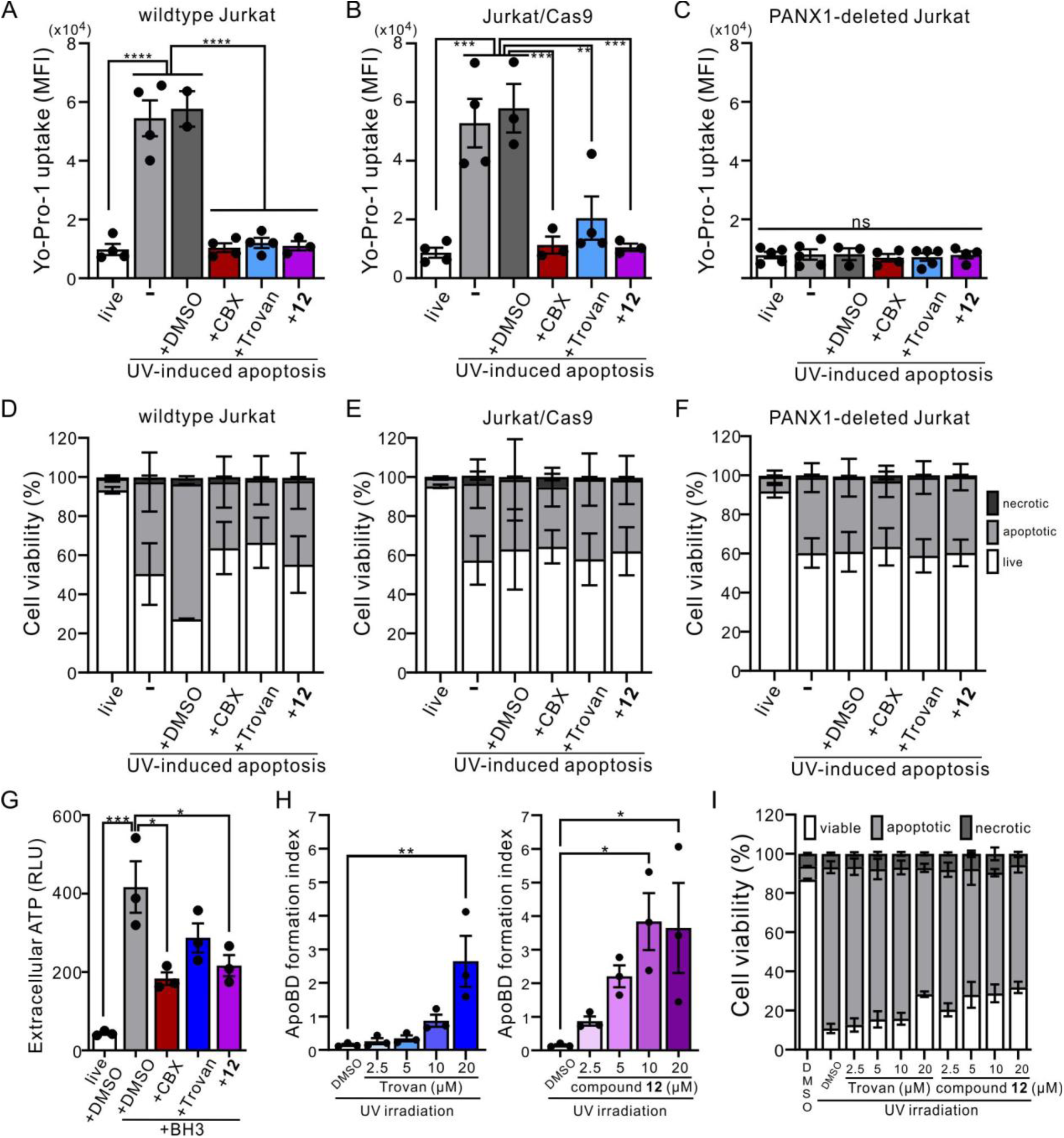
Compound 12 inhibits the activity of native PANX1 channels in the apoptotic Jurkat cells. **(A-C)** Flow cytometry analyses showing the averaged mean fluorescence intensity (MFI) of Yo-Pro-1 uptake in live or UV-induced (100 mJ cm^-2^) apoptotic wildtype **(A)**, Cas9-expressing **(B)**, or PANX1-deleted **(C)** Jurkat cells, with or without treatements of vehicle (DMSO), CBX (50 μM), Trovan (20 μM), or compound **12** (20 μM) (n=2∼4 independent experiments). **: *P* ≤ 0.01, ***: *P* ≤ 0.001, ****: *P* ≤ 0.0001, ns: not statistically significant by using one-way ANOVA with Bonferroni post hoc test. **(D-F)** Percentage of viable (Annexin V^-^/7-AAD^-^), apoptotic (Annexin V^+^/7-AAD^-^) and necrotic (7-AAD^+^) cells obtained from the flow cytometry analyses of wildtype **(D)**, Cas9-expressing **(E)**, or PANX1-deleted **(F)** Jurkat cells with or without UV-irradiation, DMSO, CBX, Trovan or compound **12** as shown in **A-C** (n=2∼4 independent experiments). **(G)** Averaged results of extracellular ATP levels (relative light unit; RLU) obtained from wildtype Jurkat cells in the presence or absence of BH3-mimetic ABT-737 (5 μM)/S63845 (10 μM), with or without applying CBX (500 μM), Trovan (20 μM), or compound **12** (20 μM) for 2 hours. *: P ≤ 0.05, ***: P ≤ 0.001, ns: not significant by one-way ANOVA followed by Bonferroni’s test (n=3 independent experiments). **(H)** Formation of apoptotic body (ApoBD) observed from UV-irradiated (150 mJ cm^-2^) Jurkat cells in the presence or absence of Trovan (2.5 ∼ 20 μM; left) or compound **12** (2.5 ∼ 20 μM; right). *: *P* ≤ 0.05, **: *P* ≤ 0.01 using one-way ANOVA followed by Dunnett’s test (n=3 independent experiments). **(I)** Percentages of viable, apoptotic and necrotic cells, as determined based on Annexin V-FITC/To-Pro-3 staining using a previously described electronic gating strategy (62), were obtained from the flow cytometry analyses of wildtype Jurkat cells exposed to UV as shown in **H** (n=3 independent experiments). Each dot represents data from a biological replicate of experiments. All histograms are preseted as means ± SEM.

Furthermore, by examining extracellular ATP concentration following apoptosis induction, we demonstrated that compound **12** (20 μM) significantly reduced the ATP released from BH3-treated Jurkat cells, similar to CBX (500 μM) (**Fig. 5G**). Consistent with our earlier reports showing that inhibiting PANX1 using Trovan or raptinal can increase the formation of large extracellular vesicles called apoptotic bodies (ApoBDs) (*23, 29*), we observed a dose-dependent effect of compound **12** on ApoBDs formation (**Fig. 5H**). While both Trovan and compound **12** markedly promoted ApoBD formation at 20 μM, we found that this effect was already prominent only with compound **12** at 10 μM (**Fig. 5H**) even as the levels of cell death were comparable across different groups (**Fig. 5I**) at all concentrations. These data indicate that compound **12** is a superior PANX1 inhibitor than the structurally similar Trovan.

### Compound 12 alleviated colitis severity of DSS-treated mice

Activated PANX1 forms a well-recognized conduit for the release of many intracellular metabolites, including ATP (*5, 6*). ATP is a well-known danger associated molecular patterns (DAMPs), and its extracellular concentration is elevated during tissue damage or chronic inflammations (*48*). Therefore, inhibition of activated PANX1 channels has been proposed as a therapeutic strategy for diverse pathological conditions in nervous, cardiovascular, and gastrointestinal systems (*12, 14*). Given that PANX1 inhibition using probenecid or ^10^Panx peptides improved colitis symptoms in animal models (*13, 17*), we examined the potential of compound **12** as an alternative pharmacological approach for treating colitis (**Fig. 6A**).

**Figure 6:**
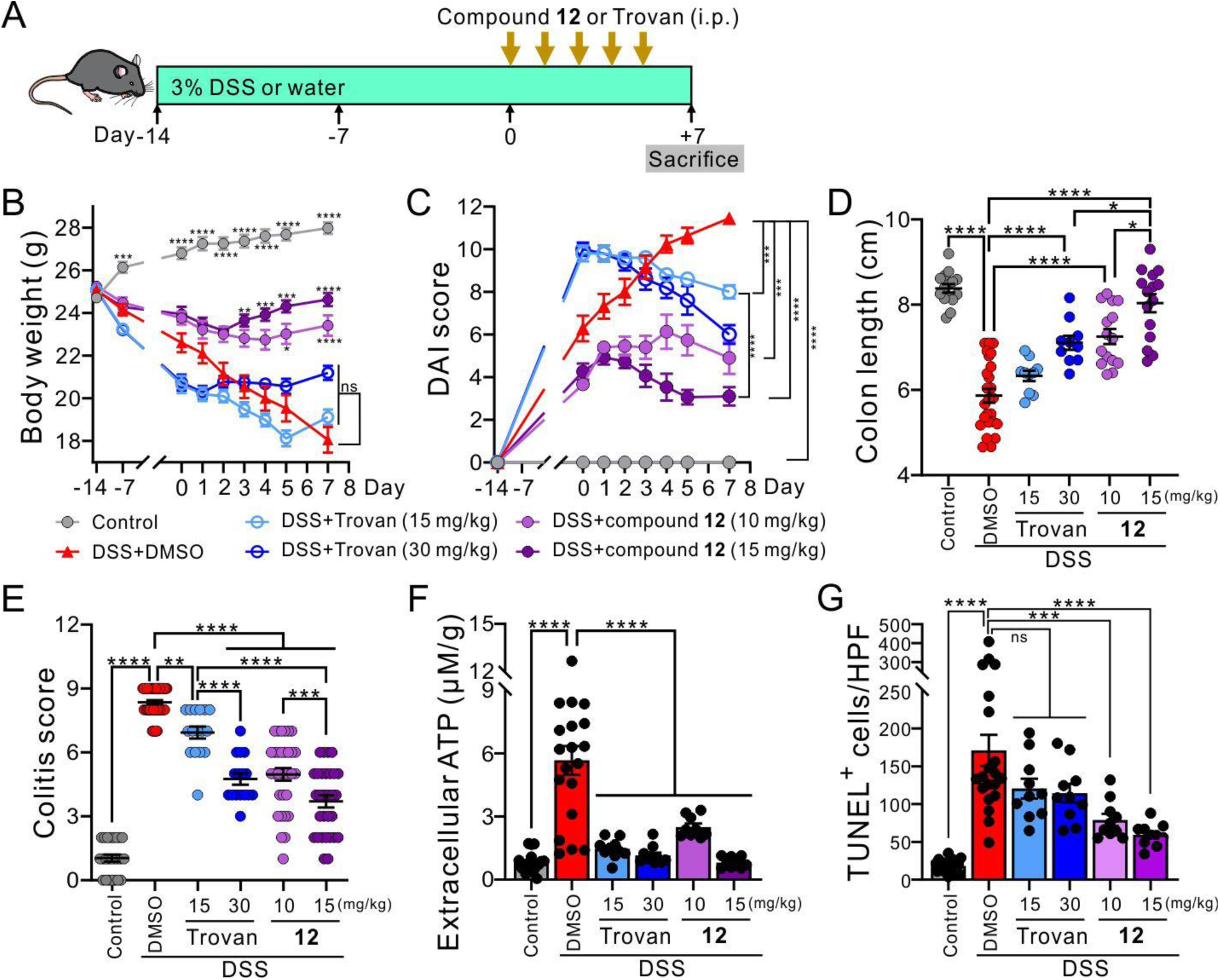
Compound 12 mitigated colitis symptoms and pathology in DSS-treated mice. **(A)** Schematic diagram illustrates the experimental design of dextran sodium sulfate (DSS)-induced colitis mice. C57Bl/6 mice were fed with regular drinking water (control) or 3% DSS-containing water for 14 days before intraperitoneal injections of compound **12** or Trovan for an additional of 5 consecutive days. Mice were sacrificed on the 7^th^ day after drug administration for macroscopic, biochemical or histological analyses. **(B)** Averaged body weights of control or DSS-fed mice, with or without treatments of DMSO (vehicle control), Trovan (15 or 30 mg/kg) or compound **12** (10 or 15 mg/kg). **(C)** Grouped data showing the disease activity index (DAI) of control and DSS-induced colitis mice, with or without Trovan or compound **12** treatments. **(D)** Grouped results of colon lengths from control or colitis mice treated with DMSO, Trovan or compound **12** as indicated. Each dot represents an individual colon sample. **(E)** Grouped results of the colitis scores from mice of the indicated treatments. Each dot represents the colitis score of an individual mouse. **(F)** Extracellular concentration of ATP measured from *ex vivo* mouse colon samples of the indicated treatments. Each dot represents the data from an individual mouse. **(G)** Groups results showing the averaged numbers of TUNEL^+^ cells within each high-power field (HPF) of colon samples from mice of the indicated treatments. Each dot represents the data of an individual HPF. Averaged results are shown as means ± SEM from 10∼25 mice per group. *: *P* ≤ 0.05, **: *P* ≤ 0.01, ***: *P* ≤ 0.001, ****: *P* ≤ 0.0001, ns: not significant using one-way **(D-G)** or two-way **(B,C)** ANOVA followed by Bonferroni’s test between the indicated groups.

Compared with the control mice consuming regular drinking water, C57Bl/6 mice consuming 3% dextran sodium sulfate (DSS) in drinking water for 14 days showed clinically relevant symptoms of colitis, including loss of body weight (**Fig. 6B**), worsened disease activity index (DAI; **Fig. 6C**), and shortened colon lengths (**Fig. 6D** and **Fig. S8A**). We found that daily intraperitoneal injections (i.p.) of compound **12** (10 or 15 mg/kg), but not Trovan (15 or 30 mg/kg), for 5 consecutive days significantly reversed DSS-induced weight loss (**Fig. 6B**). We also observed that both Trovan and compound **12** reduced the DAI scores of colitis mice in a dose-dependent manner 7 days after drug administration, yet compound **12**-treated mice had a larger decrease of DAI scores than Trovan-treated mice at the same concentration (i.e., 15 mg/kg) (**Fig. 6C**), suggesting an enhanced therapeutic potential of compound **12**. Moreover, while both Trovan and compound **12** treatments lengthened the colon of DSS-fed mice in a dose-dependent fashion, compound **12** at either 10 or 15 mg/kg was enough to restore the normal colon length, whereas a higher dose of Trovan (30 mg/kg) was needed (**Fig. 6D** and **Fig. S8A**).

To further evaluate the severity of colon inflammation in the DSS-induced colitis mice, we performed hematoxylin and eosin (H&E) staining and found that DSS-treated mice exhibited increased leukocyte infiltration and submucosal inflammation, along with goblet cell depletion (**Fig. S8B**), with an increased colitis score compared with the control mice (**Fig. 6E**), suggesting a worsened local inflammation. While both Trovan and compound **12** dose-dependently reduced the colitis scores in DSS-treated mice, compound **12**-treated mice showed significantly lowered colitis scores than Trovan-treated mice at 15 mg/kg (**Fig. 6E**), suggesting that the newly identified PANX1 inhibitors might represent an alternative, proof-of-principle therapeutic approach to reduce colitis severity. We also examined the extracellular ATP concentration of the colon samples collected from different groups of mice and found that treatments with either Trovan or compound **12** significantly reduced the concentration of extracellular ATP in the colitis mice (**Fig. 6F**). Moreover, by carrying out terminal deoxynucleotidyl transferase dUTP nick end labeling (TUNEL) assays, we found that compound-**12**-treated colitis mice had fewer TUNEL-positive (TUNEL^+^) cells than vehicle-treated colitis mice (**Fig. 6G** and **Fig. S8C**), indicating that compound **12** might ease colitis symptoms by reducing cell death. Of note, even at the higher dose of compound **12** (15 mg/kg), we did not find any noticeable alterations on the microscopic structures of tissues in healthy mice (**Fig. S9**), indicating compound **12** could be well-tolerated *in vivo*.

## Discussion

In this study, we identified a naphthyridone, compound **12** (lithium 1-(2,4-dimethoxybenzyl)-6-fluoro-7-(4-methylpiperazin-1-yl)-4-oxo-1,4-dihydro-1,8-naphthyridine-3-carboxylate), among 27 naphthyridone analogues as a new PANX1 inhibitor with a superior IC_50_ (∼0.73 μM) than the commonly used inhibitors (e.g., probenecid or ^10^Panx) (*14, 22*). By combining SAR analysis, mutagenesis, CETSA and molecular docking, we also demonstrated that compound **12** likely inhibits PANX1 channels through a direct interaction with Trp74 residue at the first extracellular loop, which may involve hydrogen bonding and π-stacking with carboxylic acid at R^3^ position and 2,4-dimethoxylbenzyl group at R^2^ position of the compound, respectively. Given that compound **12** showed no effects on the activity of pannexin-homologous LRRC8/SWELL1 channels (*39*) or human Topoisomerase II, another potential off-target proposed for Trovan (*46*), it represents the most specific and potent PANX1 inhibitor with marked inhibitory effects on PANX1 currents and permeation of large organic molecules (i.e., ATP, To-Pro-3, and Yo-Pro-1).

Despite tremendous efforts to identify new PANX1 inhibitors, few are specific and effective (*14*). Carbenoxolone inhibits PANX1 at low micromolar levels, but it also affects connexins, P2X7 receptors, and LRRC8/SWELL1 channels (*22*). Mefloquine is more potent yet interferes with LRRC8/SWELL1 channels and adenosine receptors (*22*), which may cause adverse neurological effects (*20*). A higher concentration of probenecid is required to reduce PANX1 activity (*19*), which also affects the activity of TRPV2 channels (*49*) and organic acid transporters (*50*). Besides small chemicals, the PANX1-mimetic ^10^Panx peptides, comprising 10 amino acid residues from the first extracellular loop of PANX1, has been extensively employed in research to investigate the roles of PANX1 in asthma, seizure, morphine withdrawal syndrome, and breast cancer metastasis (*10, 51–53*). However, ^10^Panx may simply form a steric block at the channel pore and could also non-specifically inhibit connexins at a similar concentration (*54*). The presence of aforementioned limitations underscores the importance for careful interpretation of experimental results with those inhibitors and highlights the need of developing more specific and potent PANX1 inhibitors for basic and translational studies.

For instance, there is an ongoing debate regarding whether PANX1 channels can be activated by mechanical stretch or hypotonic swelling (*55*). However, by using sulfo-Cy5 uptake and YFP quenching assays, we demonstrated that genetically deleting or pharmacologically inhibiting PANX1 channels does not affect the hypo-osmolarity-induced activity of volume regulated anion channels (VRACs), indicating that PANX1 channels are insensitive to changes in cell volume. This emphasizes the need to reassess PANX1 inhibitors originally identified in hypotonic conditions, such as the recently reported macrocyclic peptides (*56*). It also highlights the advantage in utilizing Trovan or compound **12** as PANX1 inhibitors to prevent misinterpreting the roles between PANX1 and LRRC8/SWELL1 channels, given their similar pharmacological and electrophysiological properties (*55*).

A recent study reported a quinoline-based new PANX1 inhibitor with an IC_50_ of 1.5 μM (*57*). By performing electrophysiological recording using *Xenopus* oocytes expressing PANX1 channels, that study found a gradually reduced percent inhibition of PANX1 currents with a stepwise extension of methylene linker between N1 position of quinoline and the phenyl ring. Consistently, our SAR analysis also suggested that one-carbon extension at N1 position yielded an improved PANX1 inhibition potency. However, it is worth noting that in the present study, compound **12** features a 2,4-dimethoxy substituent on N1 aromatic ring, which may lead to a relatively electron-donating characteristics rather than an electron-withdrawing 4-carboxyl group used in the previous report (*57*). The variation in the inhibitory potency between two compounds might be explained by the differences in their ability to form π-stacking interactions with Trp74 residues of PANX1. Therefore, it would be worth exploring whether introducing a rotational barrier of 2,4-dimethoxy substituent in compound **12**, such as adding a methyl group to the carbon extended from N1 position, could alter the PANX1 inhibitory potency of the compound.

Given that Trp74-to-Ala74 substitution (hPANX1-W74A) led to compound-**12**-insensitive PANX1 currents (**Fig. 3**) and the loss of compound-**12**-dependent thermal shift in PANX1 proteins (**Fig. 3**), Trp74 represents a pivotal residue responsible for the interaction between PANX1 channels and compound **12**. Intriguingly, Trp74 has also been suggested as a binding site for CBX (*30*), which is chemical-physically distinct from compound **12**. Thus, a common inhibitory mechanism could be shared by multiple PANX1 inhibitors. While it is appealing to envision inhibitors as a pore plug since Trp74 residue represents the narrowest constriction of PANX1 permeation pathway, previous single-channel analyses reported a reduced open probability with a preserved unitary conductance of PANX1 channels upon exposure to CBX or Trovan (*23, 27*), which rather implicates an allosteric mechanism for inhibition. Therefore, Trp74 and the nearby amino acid residues may constitute a region associated with the still unidentified gating mechanism of PANX1 channels. Additionally, while CETSA results support a direct interaction between PANX1 and compound **12**, the binding stoichiometry and actual occupancy of compound **12** remain to be determined by structural analyses using PANX1-compound-**12** complexes of high-resolution. It should be also noted that in the current study we only evaluated the effect of new inhibitors on the cleavage-activated PANX1 channels. Whether compound **12** can also inhibit PANX1 channels activated by different reversible mechanisms (*12*) requires further examination.

By performing electrophysiological recordings, we noted that the C-terminally cleaved hPANX1-R75A channels mediated a whole-cell current with weak outward and inward rectification, by contrast with the outwardly rectifying profile of wild type channels. In addition, there was no obvious shift in the reversal potential compared with wild type channels (**Fig. 2A** *cf.* **Fig. 3E**). While Arg75 likely contributes to the anionic preference of PANX1 channels (*31, 33*), the unchanged reversal potential in R75A-mutated channels suggests that C-terminal-cleaved hPANX1 channels are relatively non-selective, consistent with the previous report (*6*). Meanwhile, we also found that compound **12** demonstrated a more prominent inhibition on currents of the C-terminally-cleaved hPANX1 at negative potentials than positive potentials, as also observed with Trovan (*23*). However, this voltage-dependent inhibition was unexpectedly abolished in the C-terminally-cleaved hPANX1-R75A channels, where compound **12** significantly reduced the currents at both positive and negative potentials (**Fig. 3E-F**, and **3H**). Multiple cryo-EM maps have suggested that the inter-subunit π-cation interaction between Trp74 and Arg75 from the nearby subunit poses a rigid entrance surrounded with an anion-selective filter comprising 7 positively charged Arg75 residues (*31, 33*). Therefore, it is possible that the augmented inwardly rectifying currents observed in hPANX1-R75A channels and its diminished voltage-dependent inhibition by compound **12** might be attributed to a more flexible positioning of Trp74 residues at PANX1 extracellular entrance, where Trp74 no longer can curb the ionic flux across the plasma membrane in a certain direction. Additionally, this rectifying property of wild type PANX1 channels likely involves an intrinsic mechanism, rather than unidentified soluble agents (*58*), because the same voltage-dependent effect was found in both whole-cell current and the unitary conductance obtained from the inside-out membrane patches with a constant superfusion of recording solutions (*55*).

PANX1 is a well-recognized plasma membrane ion channel that can release signaling metabolites, including ATP (*12*). While ATP and purinergic receptors, particularly P2X7 receptors, have been implicated in the inflammatory bowel disease (IBD) (*59*), the role of PANX1 in IBD awaits clarification. A previous study reported that ^10^Panx peptides or P2X7 receptor antagonists significantly decreased the crypt damage in tumor necrosis factor alpha (TNFα) and IL-1β induce colitis of human colonic mucosa strip (*17*). Application of ^10^Panx also reportedly prevented the tight junction (TJ) protein zonula occludens 1 (ZO 1) diminishment and attenuated the increase of paracellular permeability in cytokine-induced colitis (*17*). Furthermore, inhibition of PANX1 channels using probenecid was also found to protect against the loss of myenteric neurons in multiple colitis mouse models (*13*). Our proof-of-concept examination in the DSS mouse model of colitis supports the hypothesis that PANX1 could be a potential therapeutic target for treating IBD, warranting further explorations on PANX1 inhibitors as a novel therapeutic strategy to treat IBD and to ameliorate IBD-associated dysmotility. Besides IBD, effects of compound **12** on other PANX1-related disorders, such as chronic neuropathic pain (*11*), traumatic brain injury (*24*), and ischemia-reperfusion injuries (*60*), remain to be investigated. Finally, while the current study focuses on identifying new PANX1 inhibitors, further research to develop PANX1 activators is essential to advance our understandings regarding the roles of PANX1 channels in their native environment.

## Material & Methods

### Reagents

Carbenoxolone and trovafloxacin were purchased from Sigma-Aldrich. Dicoumarol was obtained from Selleck Chemicals. To-Pro-3 and Yo-Pro-1 were purchased from Thermo-Fisher Scientific. Sulfo-Cyanine5 carboxylic acid (sulfo-Cy5) was purchased from Lumiprobe. Annexin V-FITC and annexin-V binding buffer were purchased from BD Bioscience. Annexin V-EV450/7-AAD apoptosis detection kit was obtained from Elabscience. ABT-737 and S63845 were purchased from Jomar Life Research. Dextran sodium sulfate (DSS; molecular weight 36,000–50,000) was purchased from MP Biomedicals.

### General procedures for chemical synthesis

All commercial chemicals and solvents are of reagent grade and were used without further purification unless otherwise stated. All reactions were carried out under dry nitrogen or argon atmosphere and were monitored for completion by thin-layer chromatography (TLC) using Merck 60 F254 silica gel glass-backed plates or aluminum plates, which were detected visually under UV irradiation (254 nm). Flash column chromatography was carried out using silica gel (Silicycle SiliaFlash P60, R12030B, 230−400 mesh or Merck Grade 9385, 230−400 mesh). Structures of synthesized compounds were verified by using NMR and spectra of ^1^H and ^13^C can be found in **Fig. S1**. ^1^H and ^13^C NMR spectra were recorded with Bruker 400 or 600 MHz AVANCE III spectrometers. Data for NMR spectra were analyzed with Mnova software (Mestrelab Research). Chemical shift (δ) was reported in ppm and referenced to solvent residual signals as follows: DMSO-*d_6_* at 2.49 ppm, chloroform-*d* at 7.26 ppm, methanol-*d*_4_ at 3.31 ppm for ^1^H NMR; DMSO-*d_6_* at 39.5 ppm, chloroform-*d* at 77.0 ppm, methanol-*d*_4_ at 49.0 ppm for ^13^C NMR. Splitting patterns are indicated as follows: s = singlet; d = doublet; t = triplet; q = quartet; quin = quintet; dd = doublet of doublets; dt = doublet of triplets; td = triplet of doublets; ddd = doublet of doublets of doublets; br = broad; m = multiplet. Coupling constants (*J*) were given in Hertz (Hz). Low-resolution mass spectra (LRMS) data were measured with Agilent MSD-1100 ESI-MS/MS system or Agilent Infinity II 1290 LC/MS (ESI) systems. High-resolution mass spectra (HRMS) data were measured with a Varian 901-MS FT-ICR HPLC/MS-MS (ESI) system. Purity of the final compounds was determined using a high-performance liquid chromatography (HPLC) system (Hitachi 2000 series) equipped with a C_18_ column (Agilent ZORBAX Eclipse XDB-C_18_ 5 μm. 4.6 mm × 150 mm) and operating at 25 °C. The injection volume of each sample in dimethyl sulfoxide (DMSO) was 20 μl. The flow rate of the mobile phases was 0.5 ml per minute. The elution was carried out using acetonitrile as mobile phase A, and water containing 0.1% formic acid + 2 mmol NH_4_OAc as mobile phase B. Elution conditions: at 0-minute, 10% phase A + 90% phase B; at 25-minute, 90% phase A + 10% phase B; at 30-minute, 90% phase A + 10% phase B; at 30.5-minute, 10% phase A + 90% phase B; and at 37-minute, 10% phase A + 90% phase B. Peaks were detected at 254 nm. The mesylate salt compounds were prepared using neutral form (1.0 equiv) in methanol and CH_2_Cl_2_ co-solvent. The mixture was added methanesulfonic acid (1.0 equiv) and stirred at r.t. for 12 hours. After the solvent was evaporated, the residue was washed with methanol and collected the solid via filtration to give the mesylate salt compounds.

### Plasmids and side-directed mutagenesis

Plasmids encoding C-terminally FLAG-tagged human Pannexin 1 (hPANX1, hPANX1-FL-EGFP, hPANX1-CT-EGFP, and hPANX1(TEV)-EGFP) and Tobacco Etch Virus protease (TEVp) were previously described (*26, 27, 29*). To generate the C-terminally FLAG-tagged hPANX1-W74A, hPANX1-R75A, hPANX1(TEV)-W74A-EGFP, and hPANX1(TEV)-R75A-EGPF, we performed site-directed mutagenesis on the FLAG-tagged hPANX1 or hPANX1(TEV)-EGFP using PROTECH-HP™ high performance *Pfu* DNA polymerase (Protech Technology) and the following primers: 5’-GTTCTTTCTCCGCTCGTCAGGCTGCCTTTGTGG-3’ / 5’-GGCAGCCTGACGAGCGGAGAAAGAACTT-3’ (for W74A) and 5’-CTTTCTCCTGGGCTCAGGCTGCCTTTGTGGATTCATATTG-3’ / 5’-CAAAGGCAGCCTGAGCCCAGGAGAAAGAACTTGGAGAG-3’ (for R75A). pcDNA3.1 Hygro EYFP H148Q/I152L was a gift from Dr. Peter Haggie (Addgene plasmid #25874)(*45*). All plasmids were verified by using Sanger’s sequencing.

### Cell culture and transfection

HEK293T cells (passage 7–20, ATCC) were maintained at 37 °C with humidified air containing 5% CO2 in Dulbecco’s Modified Eagle Medium (DMEM, high glucose, Gibco) containing 10% fetal bovine serum (FBS, Gibco), penicillin, streptomycin, and sodium pyruvate. Human Jurkat T cells (E6.1, ATCC) were cultured in RPMI 1640 (Gibco) supplemented with 10% (v/v) fetal calf serum (Gibco), 50 IU/ml penicillin and 50 μg/ml streptomycin. Cas9-stably-expressed and PANX1-deleted Jurkat cells were described previously (*47*). To generate PANX1-deleted HEK293T cells, HEK293T cells stably expressing Cas9 and EGFP (GeneCopoeia; #SL502) were transfected using Lipofectamine 3000 and pLX-2 plasmids encoding two PANX1 sgRNAs described in the previous study (*47*), and colonies of transfected cells were selected using 1 μg/mL puromycin and 200 μg/mL Zeocin. Deletion of PANX1 was verified by using immunoblotting (**Fig. S5**). To generate cells expressing iodide-sensing YFP (*45*), following a transient transfection using Lipofectamine 3000, colonies of HEK293T cells stably expressing EYFP H148Q/I152L were selected by using culture media containing hygromycin B (125 μg/ml). For whole-cell recordings, HEK293T cells were transiently co-transfected with plasmids encoding TEV protease and plasmids encoding hPANX1(TEV)-EGFP, hPANX1(TEV)-W74A-EGFP, or hPANX1(TEV)-R75A using Lipofectamine 3000 following manufacturers’ manual. Cells were replated onto poly-L-lysine-coated coverslips 16-18 hours after transfection.

### Dye-uptake assays and flow cytometry

We used To-Pro-3 uptake assays to examine whether varying naphthyridone analogues can inhibit the activity of C-terminally-truncated PANX1 channels. HEK293T cells were transiently transfected with hPANX1-FL-EGFP or hPANX1-CT-EGFP, collected, and washed once with ice-cold phosphate-buffered saline (PBS; 137 mM NaCl, 2.7 mM KCl, 1.8 mM KH2PO4, 10 mM Na2HPO4, pH 7.4). To test effects of different analogues, 150 µl of cell solution (∼5×10^5^ cells) was added into 150 µl of PBS containing 2× concentrated analogues, followed by a 10-minute incubation at ambient temperature before 3 µl of 0.1 mM To-Pro-3 was added into the cell-analogues mixture for another 10-minute incubation at ambient temperature.

To assess the effects of different naphthyridone analogues on the activity of endogenous PANX1 channels, Jurkat T cells (3×10^6^ cells/ml in PBS) were collected and spread on a Petri dish before exposed to UV radiation (100 mJ/cm^2^) using CL-1000 ultraviolet crosslinker (Analytik Jena), followed by a 1-hour incubation at 37 °C. A mixture of 150 µl UV-irradiated cells (5×10^5^ cells) and 150 µl PBS containing 2× concentrated naphthyridone analogues was incubated at ambient temperature for 10 minutes before 3 µl of 0.1 mM Yo-Pro-1 was added into the mixture for another 10-minute incubation at ambient temperature. Yo-Pro-1-stained cells were collected, and a mixture of annexin V/7-AAD (3 µl annexin V-EV450 and 3 µl 7-AAD in 300 µl annexin V binding buffer, Elabscience) was added into cell-analogues mixture, followed by an incubation at room temperature for 15 minutes before flow cytometry analysis.

To determine whether Trovan or compound **12** affected the activity of LRRC8/SWELL1 channels, we performed sulfo-Cy5 uptake assays in wildtype, Cas9-expressing, or PANX1-deleted HEK293T cells. 24 hours after seeding 2×10^4^ HEK293T cells per well of a 24-well plate, cells were washed once using PBS before a 10-minute incubation with isotonic (310 mOsm; 140 NaCl, 5 KCl, 20 HEPES in mM, pH 7.4) or hypotonic (125 mOsm; 5 KCl, 20 HEPES, and 90 mannitol in mM) solutions at 37 °C. Sulfo-Cy5 at a final concentration of 1 µM was then mixed with the cells for 30 minutes at 37 °C before the cells were trypsinzed and analyzed by using flow cytometry (CytoFLEX, Beckman Coulter).

All the samples were placed on ice before analyzed by using CytoFLEX flow cytometry and CytExpert 2.0 software (Beckman Coulter).

### Whole-cell voltage-clamp recordings

We performed whole-cell recordings at room temperature to examine whether compound **12** can inhibit ionic currents of C-terminally-cleaved PANX1 channels in HEK293T cells. Following transient transfections, whole-cell currents of PANX1-expressing, EGFP-positive HEK293T cells were recorded using a bath solution containing 140 mM NaCl, 3 mM KCl, 2 mM MgCl2, 2 mM CaCl2, 10 mM HEPES, and 10 mM glucose (pH 7.3; ∼300 mOsm) and an pipette solution composed of 100 mM CsMeSO4, 30 mM TEACl, 4 mM NaCl, 1 mM MgCl2, 0.5 mM CaCl2, 10 mM HEPES, 10 mM EGTA, 3 mM ATP-Mg, and 0.3 mM GTP-Tris (pH 7.3; ∼290 mOsm). Micropipettes of 3–5 MΩ were pulled from thin-walled, fire-polished borosilicate glass capillaries (Harvard Apparatus) using a MP-500 RWD Micropipette Puller (RWD). Axopatch 200B amplifier controlled by pCLAMP11 software (Molecular Devices) and Digidata 1550B digitizer (Molecular Devices) was used to apply ramp voltage-clamp commands (from -130 mV to +80 mV) at 7-second intervals. CBX-sensitive current was taken as the difference in current at -50 mV or +80 mV before and after CBX application and was normalized to cell capacitance as a measure of current density.

### Hypertonicity-induced YFP quenching

HEK293T cells stably expressing EYFP H148Q/I152L were seeded at a density of 6×10^4^ cells per well in black opaque 96-well plates. The culture media was removed after 24 hours, and the cells were washed once with PBS before the addition of 90 µl of isotonic (310 mOsm) or hypotonic (125 mOsm) solution for 10 minutes at ambient temperature, with or without the presence of CBX, dicoumarol, Trovan or compound **12**. The fluorescence intensity of YFP (excitation: 497-15; emission:545-20) was assessed at 3-second intervals by using a CLARIOstar Plus multimode microplate reader (BMG Labtech) before and after the auto-injection of 10 µl NaI (100 mM).

### Topoisomerase II assay

We used Topoisomease II assay kit (TopoGEN) to examine whether Trovan or compound **12** inhibited the activity of human Topoisomerase II *in vitro*. In brief, catenated DNA (kDNA) of 80 ng and 1U human Topoisomerase II were incubated with or without VP16 (0.1 mM), CBX (50 µM), Trovan, or compound **12** (20 or 200 µM) for 30 minutes at 37 °C, and then 2 µl of 10% sodium dodecyl sulfate (SDS) and 2 µl of protease K (50 µM) were added. Samples were then incubated for another 15 minutes at 37 °C, followed by electrophoretic separation on 1% agarose gels with GelRed® nucleic acid gel stain (Biotium) at 200 V for 15 minutes at 4 °C. Signals of catenated and decatenated DNA were visualized using ChemiDoc™ Imaging Systems (BioRad) and quantified by using ImageJ software (ver. 15.4).

### ATP release assay of apoptotic Jurkat T cells

To induced apoptosis, Jurkat T cells in 1% BSA/RPMI media were seeded into a 96-well plate at 1.5×10^5^ cells per well and treated with BH3 mimetics, ABT-737 (5 μM) and S63845 (10 μM), in the presence or absence of Trovan (20 μM) or compound **12** (20 μM) at 37°C. Levels of extracellular ATP were measured every 10 mins for 4 hours by using a luciferase activity assay kit (RealTime-Glo™ Extracellular ATP Assay, Promega) according to manufacturer’s instructions. Luminescent intensity (relative light unit; RLU) was determined by using a multimode microplate reader (CLARIOstar Plus; BMG Labtech).

### Apoptotic body (ApoBD) formation index

To induce apoptosis by UV irradiation, Jurkat T cells in 1% BSA/RPMI media were exposed to 150 mJ/cm^2^ of irradiation using the Stratagene UV Stratalinker 1800 (Agilent Technologies) and incubated for 4 hours at 37°C. Cell viability and ApoBD formation following UV-mediated apoptosis induction and PANX1 inhibitor treatments were determined by flow cytometry using the BD FACSCanto II flow cytometer (BD Bioscience) as previously described (*61, 62*). All flow cytometry data was analyzed with FlowJo software (ver. 10.8.0).

### Cellular thermal shift assay (CETSA) and immunoblotting analysis

All procedures were modified based on previous studies (*34, 35*). In brief, after 24 hr of transfection, HEK293T cells expressing varying FLAG-tagged hPANX1 constructs were trypsinized, washed once using PBS, and resuspended in PBS at a density of 1×10^7^ cells per ml. For each temperature, 50 μl of resuspended cells were mixed and incubated with 50 μl of PBS, with or without compound **12** (50 μM) at 37 °C in humidified air composed of 5% CO2 for 1 hour. Cells were then exposed to varying temperatures (63.3, 68.7, 73.9, 77.1, and 79 °C) for 3 minutes using a PCR amplifier (T100™ Thermal Cycler, BioRad), followed by adding NP-40 to a final concentration of 2% (w/v). The samples were snap frozen using liquid nitrogen and quickly thawed 3 times, followed by a centrifugation at 20,500 ×*g* for 20 minutes at 4 °C.

The supernatants were mixed with 5× Laemmli buffer (62.5% glycerol, 12.5% SDS, 0.5% bromophenol blue, 25% fresh 2-mercaptoenthanol in 30 mM Tris-HCl, pH 6.8) and separated by SDS-PAGE. The protein samples were then transferred onto 0.2 µm PVDF membranes (Pall Corporation) and blocked with 5% non-fat dry milk dissolved in a Tris-based buffer (10 mM Tris, 150 mM NaCl, and 0.1% Tween 20, pH 7.4) at room temperature for 1 hour. Human PANX1 proteins were detected by incubating with THE™ DYKDDDK tag antibody (GenScript; #A00187; 1:1000) at 4 °C for 16-18 hours. An antibody to α-tubulin (GeneTex; GTX112141; 1:20,000) was used as a loading control for CETSA. Horseradish peroxidase (HRP)-linked secondary antibodies (Jackson ImmunoResearch, Inc.; 115-035-003 or 111-035-003; 1:5000), T-Pro LumiLong Plus Chemiluminescence Detection kit (T-Pro Biotechnology), and ChemiDoc™ Imaging Systems (BioRad) were used to visualize immunoreactive signals. The intensity of immunoreactive signals at different temperatures was quantified using ImageJ software (ver. 15.4) and was normalized to that at 63.3°C. Averaged results were fitted using a sigmoidal model in GraphPad Prism (ver. 10) to obtain the apparent aggregation temperatures (T_agg_) (*34*), with and without compound **12**.

### PANX1-compound 12 docking analysis

Molecular docking was performed to investigate the binding mode of compound **12** to a PANX1 channel. Structure of compound **12** was created from the Reaxys database (Elsevier, NL). The cryo-EM structure of human PANX1 channel (PDB: 6WBI) (*31, 63*) was used to predict the binding status of compound **12** by molecular docking analysis with RyRx/AutoDock VINA (ver. 0.9.8). A grid box space with dimensions of 24.37 Å x 24.04 Å x 30.10 Å (x-, y-, and z-axes) near the extracellular entrance was generated using AutoGrid embedded in PyRx/AutoDock Vina (*36, 64*). The exhaustiveness parameter was 50, and the number of modes was 11. The docking results were analyzed according to the predicted binding mode and corresponding binding affinity (kcal/mol). Docked complexes were visualized and analyzed using the PyMOL Molecular Graphics System (Schrödinger, LLC; ver. 2.4), LigPlot+ 2.2.8 program (https://www.ebi.ac.uk/thornton-srv/software/LigPlus/)(38) and the Protein-Ligand Interaction Profiler (PLIP) (*37*).

### Dextran sodium sulfate (DSS)-induced colitis mouse model

Male C57BL/6J mice aged 7–10 weeks were obtained from the National Laboratory Animal Center in Taipei, Taiwan. All mice were kept in specific pathogen-free environments with controlled temperature (23±2 °C) and a 12:12-hour light-dark cycle. The experimental procedures were approved by the Laboratory Animal Care Committee of National Taiwan University College of Medicine (202308118RINB).

We induced colitis in mice by using dextran sodium sulfate (DSS) as previously described (*65*). While both male and female mice can develop DSS-induced colitis symptoms, male mice were found to be highly sensitive to DSS induction and can develop more aggressive symptoms than female mice (*66*). Therefore, we used only male mice for this proof-of-principle study assessing the therapeutic potentials for the newly developed PANX1 inhibitors. For the DSS-treated groups, mice were fed with 3% DSS (MP Biomedicals; Molecular weight 36,000– 50,000) in their drinking water for 21 days (**Fig. 6A**). Starting on the 15^th^ day of DSS-treatment, the mice received daily intraperitoneal injections of either DMSO (5%; vehicle control), Trovan (15 or 30 mg/kg), or compound **12** (10 or 15 mg/kg) for 5 consecutive days. Mice were euthanized at the end of the treatment period on day 21^st^, and their colons were collected for further analyses.

Body weight and disease activity index (DAI) of mice were monitored over the course of treatments. DAI was evaluated based on criteria of percent weight loss, intestinal bleeding (i.e., no blood, occult blood (hemoccult+), or gross blood), and stool consistency (i.e., normal stools indicate well-formed pellets, pasty and loose stools indicate semi-formed stools, and diarrhea indicates liquid stools) and was scored according to **Table 4** as previously described (*67*).

**Table 4.** Scoring of the DAI.

| Score | Weight loss (%) | Stool consistency | Bleeding |
| --- | --- | --- | --- |
| 0 | None | Normal | Normal |
| 1 | 0-10 |  |  |
| 2 | 10-15 | Loose stool | Hemoccult + |
| 3 | 15-20 |  |  |
| 4 | >20% | Diarrhea | Gross bleeding |

After mice were euthanized, the colonic tissues were collected and stained by using hematoxylin and eosin (H&E) for further histological severity grading by a pathologist who was blinded to the experimental conditions. The colitis scores were recorded based on the following criteria as previously reported (*68*): a blinded scorer assessed the sections for goblet cell depletion, leukocyte infiltration, and submucosal inflammation using a point scale from 0 to 3, where 0 indicates no pathology and 3 represents the most severe pathology. Three equally weighted subscores were summed, resulting in a combined histological colitis severity score ranging from 0 to 9. Hematoxylin and eosin (H&E) staining were conducted as previously described (*69*). Heart, liver, kidney, lung, colon and spleen tissues from different groups (healthy control, DMSO-treated, and PANX1 inhibitor-treated) were fixed in 10% formalin, embedded in paraffin, and sectioned to a thickness of 4 µm. These sections were then stained with H&E and examined under an optical microscope. Images were captured and digitized by using Nikon Eclipse Ni and NIS Elements BR 4.30.02.

### Terminal deoxynucleotidyl transferase dUTP nick end labeling (TUNEL)

Apoptosis cells in distal colon tissues were assessed utilizing the TUNEL method. Paraffin-embedded tissue sections of 4 µm thick from the distal colon were processed following the manufacturer’s protocol for the TUNEL assay kit (Abcam, #ab206386). Images of 10 randomly selected fields per slide were captured at 400X magnification. Quantification was performed by counting the number of TUNEL-positive cells in the field using ImageJ (NIH, Bethesda, MD, USA), and its implemented image processing version Fiji (https://imagej.net/) (*70*).

### ATP release assay of *ex vivo* colon samples

To examine whether PANX1 inhibitors can reduce ATP release from the *ex vivo* colon tissues, we measured the extracellular concentration of ATP by using a commercial ATP assay kit (Elabscience, #E-BC-F002) following the manufacturer’s instruction. Colon samples were incubated with a 10-fold volume (w/v) of phosphate-buffered saline (PBS) for 1 hour at 37 °C, and then 100 μl of supernatant was collected and mixed with 100 μl of the substrate solution. After a constant shaking in the dark, the luminescence intensity of the mixture was measured by using a multifunctional microplate reader (Hidex Sense, Hidex), and the data was normalized to the sample weight (g).

### Statistical analysis

All data were presented as means ± SEM. Statistical analyses were performed using GraphPad Prism (v.10). Statistical significance was determined by two-tailed, unpaired t test, one-way or two-way ANOVA, followed by the Bonferroni’s or Dunnett’s multiple comparisons test, as described in figure legends, with *P* ≤ 0.05 deemed significance significant.

## Acknowledgments

The authors appreciate the technical support from the Structural Proteomics and Pharmaceutical Application Service at the TMBD Bioinformatics Core (http://www.tbi.org.tw), which is funded by the Bioinformatics Core Facility for Biotechnology and Pharmaceuticals. The authors would also like to thank Dr. Hui-Chun Cheng (National Tsing Hua University, Taiwan) and Dr. Douglas Bayliss (University of Virginia, U.S.A.) for the helpful discussion.

## Author contributions

W.-Y.H, Y.-L.W, H.-P.H. and Y.-H.C. developed concept and designed experiments. W.-Y.H, J.-C.L and H.-P.H designed chemical synthesis procedures, and W.-Y.H synthesized and validated all chemical compounds. Y.-L.W carried out mutagenesis, topoisomerase II assays, Yo-Pro-1 and sulfo-Cy5 uptake assays. Y.-L.W and Y.-R.L performed To-Pro-3 uptake assays. Y.-L.W and J.Y.H. performed YFP quenching assays. Y.-L.W. and C.-N.L. performed CETSA. Y.-H.C. performed electrophysiological experiments. M.-T.W, S.-Y-L., and C.-I.L. carried out the experiments of DSS-induced colitis mice. J.P.S. performed ATP release assays and ApoBD formation assays using the apoptotic Jurkat cells. Y.-C.L. carried out molecular docking and the associated data visualization. H.-Y.G generated PANX1-deleted HEK293T cells. Y.-H.C., H.-P.H., M.-T.W., I.K.H.P., P.-C.L, S.-C.W. and J.-C.L supervised the experiments. Y.-H.C., W.-Y.H, Y.-L.W, and H.-P.H. conceptualized and wrote the manuscript, and all authors edited and commented on the manuscript.

## Data and materials availability

All data and materials supporting the findings of this study are available within the paper and its Supplementary Materials and from the corresponding authors on reasonable request.

