## Supplementary materials for "Novel naphthyridones targeting Pannexin 1 for colitis management"

**This PDF file includes:**

Supplementary Text: synthesis procedures for compounds **1–27**, **29–29l**, **30a–30b**, **31a–31l**, **32a–32b**, **33–34**.

Figs. S1 to S9

Movie S1

References (1 to 2)

**Other Supplementary Materials for this manuscript include the following:**

Movie S1

### Supplementary Text

#### Scheme 1. Synthesis of 7-substituted 1-(2,4-difluorophenyl) fluoroquinolones[a]

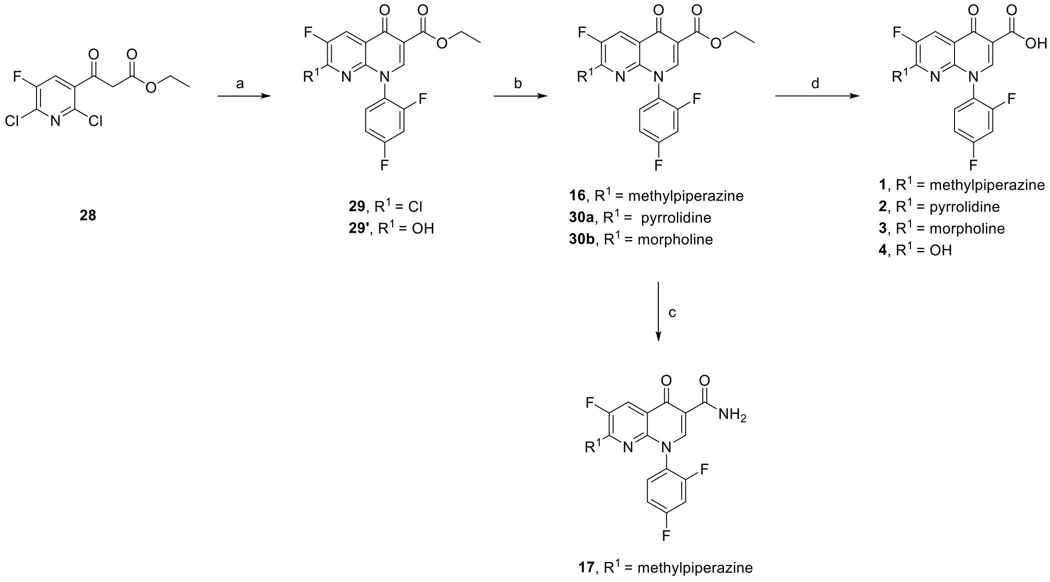


[a]Reagents and conditions: (a) (i) $Ac_2O$, $CH(OC_2H_5)_3$, 130 °C, 1 h; (ii) 2,4-difluoroaniline, $CH_2Cl_2$, r.t., 2 h; (iii) $K_2CO_3$, DMF, 90 °C, 78% (3 steps yield); (b) $R^1H$, DMF, 90 °C, 60-80%; (c) $NH_3$ in methanol, reflux in sealed tube, 18 h, 41%; (d) LiOH, ethanol, 60 °C, 3 h, then HCl, 40-60%.

#### Scheme 2. Synthesis of 1- or 7-substituted fluoroquinolones[a]

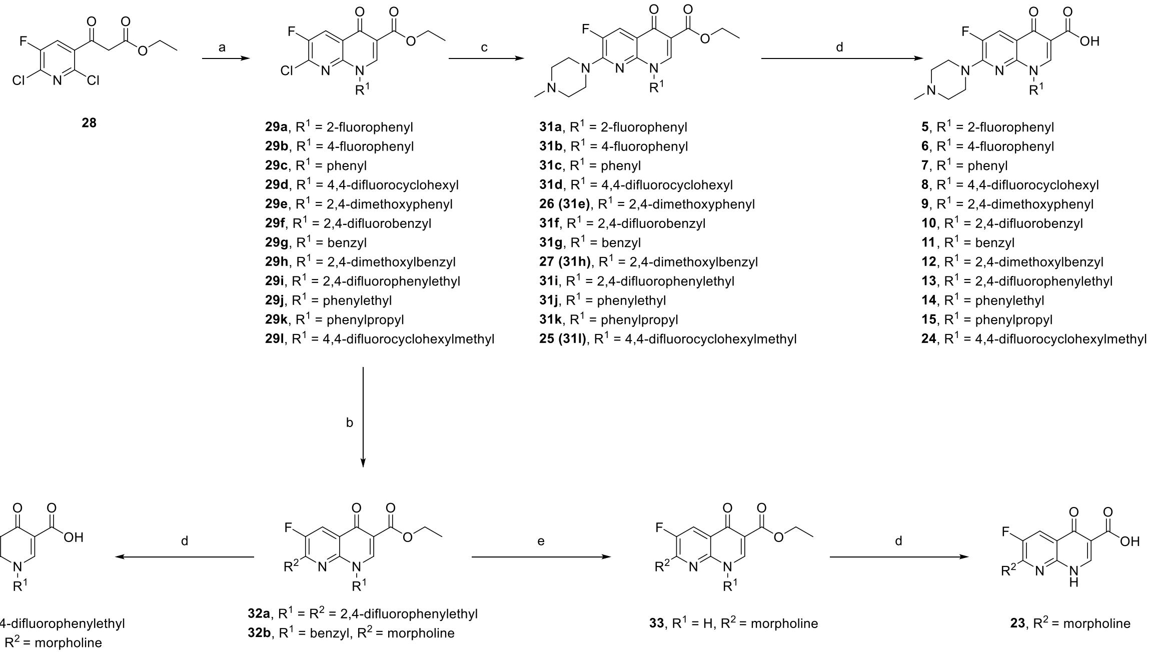


[a]Reagents and conditions: (a) (i) $Ac_2O$, $CH(OC_2H_5)_3$, 130 °C, 1 h; (ii) $NH_2R^1$, $CH_2Cl_2$, r.t., 2 h; (iii) $K_2CO_3$, DMF, 90 °C, 50-80% (3 steps yield); (b) $NH_2R^2$, DMF, 90 °C, 4% or 73%; (c) 1-methylpiperazine, DMF, 90 °C, 50-80%; (d) LiOH, ethanol, 60 °C, 3 h, then HCl, 40-60%. (e) 10 mol% Pd, cat. 12 N HCl, $H_2$, ethanol, r.t., 18 h, 30%.

**Scheme 3. Synthesis of 3-substituted 1-(2,4-difluorophenyl)-7-methylpiperazine fluoroquinolones[a]**

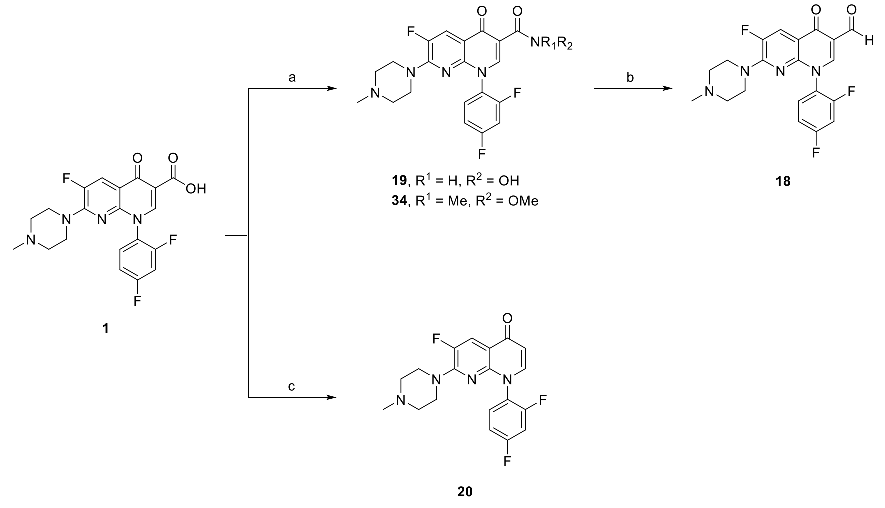


[a]Reagents and conditions: (a) EtOCOCl, $NHR^1R^2$ • HCl, $Et_3N$, $CH_2Cl_2$, 0 °C to r.t., 18 h, 41-50%. (b) DIBAL in toluene, $CH_2Cl_2$, -50 °C, 6 h, 6% (c) KCN, DMSO, 160 °C, 8 h, 50%.

### Chemistry

As described in Scheme 1 and 2, syntheses of 1- or 7-substituted fluoroquinolones **1**–**17** and **21**–**27** were prepared according to the previous reports (*1, 2*). The reaction of **28** with triethyl orthoformate, corresponding amine and cyclization afforded 6-chloro fluoroquinolones **29**, **29'** and **29a-l**. Nucleophilic aromatic substitution ($S_NAr$) of corresponding chloro derivatives with selected amine gave ethyl esters derivatives **16**, **30a**–**b**, **31a-l** and **32a**–**b**. Deprotection of benzyl group in **32b** with palladium on carbon and catalytically HCl gave **33**. Hydrolysis of ethyl ester group yield carbonyl derivatives **1**–**15** and **21**–**24**. Amide **17** was synthesized from ethyl esters derivatives **16** with $NH_3$ in methanol.

As depicted in Scheme 3, syntheses of 3-substituted 1-(2,4-difluorophenyl)-7-methylpiperazine fluoroquinolones were prepared from **1**. Activation of carboxylic acid in **1** with ethyl chloroformate and reacted with corresponding amine yield **19** and **34**. Reduction the Weinreb amide in **34** with DIBAL gave aldehyde **18**. Decarboxylation of **1** with KCN gave **20**.

### Experimental Section

**General Methods for Chemistry.** All commercial chemicals and solvents are of reagent grade and were used without further purification unless otherwise stated. All reactions were carried out under dry nitrogen or argon atmosphere and were monitored for completion by thin-layer chromatography (TLC) using Merck 60 F254 silica gel glass-backed plates or aluminum plates, which were detected visually under UV irradiation (254 nm). Flash column chromatography was carried out using silica gel (Silicycle SiliaFlash P60, R12030B, 230−400 mesh or Merck Grade 9385, 230−400 mesh). Structures of synthesized compounds were verified by using NMR and spectra of 1H and 13C can be found in Fig. S1. $^1H$ and $^{13}C$ NMR spectra were recorded with Bruker 400 or 600 MHz AVANCE III spectrometers. Data for NMR spectra were analyzed with Mnova software (Mestrelab Research). Chemical shift (δ) was reported in ppm and referenced to solvent residual signals as follows: DMSO-$d_6$ at 2.49 ppm, chloroform-*d* at 7.26 ppm, methanol-$d_4$ at 3.31 ppm for $^1H$ NMR; DMSO-$d_6$ at 39.5 ppm, chloroform-*d* at 77.0 ppm, methanol-$d_4$ at 49.0 ppm for $^{13}C$ NMR. Splitting patterns are indicated as follows: s = singlet; d = doublet; t = triplet; q = quartet; quin = quintet; dd = doublet of doublets; dt = doublet of triplets; td = triplet of doublets; ddd = doublet of doublets of doublets; br = broad; m = multiplet. Coupling constants (*J*) were given in Hertz (Hz). Low-resolution mass spectra (LRMS) data were measured with Agilent MSD-1100 ESI-MS/MS system or Agilent Infinity II 1290 LC/MS (ESI) systems. High-resolution mass spectra (HRMS) data were measured with a Varian 901-MS FT-ICR HPLC/MS-MS (ESI) system. Purity of the final compounds was determined using a high-performance liquid chromatography (HPLC) system (Hitachi 2000 series) equipped with a $C_{18}$ column (Agilent ZORBAX Eclipse XDB-$C_{18}$ 5 μm. 4.6 mm × 150 mm) and operating at 25 °C. The injection volume of each sample in dimethyl sulfoxide (DMSO) was 20 μl. The flow rate of the mobile phases was 0.5 ml per minute. The elution was carried out using acetonitrile as mobile phase A, and water containing 0.1% formic acid + 2 mmol NH4OAc as mobile phase B. Elution conditions: at 0-minute, 10% phase A + 90% phase B; at 25-minute, 90% phase A + 10% phase B; at 30-minute, 90% phase A + 10% phase B; at 30.5-minute, 10% phase A + 90% phase B; and at 37-minute, 10% phase A + 90% phase B. Peaks were detected at 254 nm. The mesylate salt compounds were prepared using neutral form (1.0 equiv.) in methanol and $CH_2Cl_2$ co-solvent. The mixture was added methanesulfonic acid (1.0 equiv.) and stirred at r.t. for 12 hours. After the solvent was evaporated, the residue was washed with methanol and collected the precipitation via filtration to give the mesylate salt compounds.

**Representative procedure A: ethyl 7-chloro-1-(2,4-difluorophenyl)-6-fluoro-4-oxo-1,4-dihydro-1,8-naphthyridine-3-carboxylate (29)**

To a solution of triethyl orthoformate (4.5 mL, 27.1 mmol, 1.5 equiv.) and acetic anhydride (15.0 mL, 158.7 mmol, 8.8 equiv.) was added ethyl 3-(2,6-dichloro-5-fluoropyridin-3-yl)-3-oxopropanoate (**28**) (5.0 g, 17.9 mmol, 1.0 equiv.). The mixture was stirred at 130 °C for 1 h. After the solvent was evaporated, the residue was dissolved in $CH_2Cl_2$ (30 mL). To a resulting solution was added 2,4-difluoroaniline (2.0 mL, 19.6 mmol, 1.1 equiv.) and stirred at r.t. for 1 h. After the solvent was evaporated, DMF (35 mL) and $K_2CO_3$ (3.2 g, 23.2 mmol, 1.3 equiv.) were added to the residue followed by stirring at 90 °C for 50 min. The mixture was poured into water (450 mL) and the precipitation was collected via filtration to give **29** in 78% yield. This compound was directly used in the next step without further purification. $^1H$ NMR (400 MHz, chloroform-*d*) δ 8.55 (s, 1H), 8.48 (d, *J* = 7.2 Hz, 1H), 7.47–7.39 (m, 1H), 7.16–7.06 (m, 2H), 4.41 (q, *J* = 7.1 Hz, 2H), 1.40 (t, *J* = 7.1 Hz, 3H). LRMS (ESI) *m/z:* 383.1 $[M+H]^+$.

**Ethyl 1-(2,4-difluorophenyl)-6-fluoro-7-hydroxy-4-oxo-1,4-dihydro-1,8-naphthyridine-3-carboxylate (29')**

Evaporating the filtrate in the last step of representative procedure A of **29** give **29'** in 13% yield. This compound was directly used in the next step without further purification. $^1H$ NMR (600 MHz, chloroform-*d*) δ 8.63 (d, *J* = 8.1 Hz, 1H), 8.56 (s, 1H), 7.46–7.40 (m, 1H), 7.14–7.05 (m, 2H), 4.41 (q, *J* = 7.1 Hz, 2H), 1.41 (t, *J* = 7.1 Hz, 3H). LRMS (ESI) *m/z:* 365.1 $[M+H]^+$.

**Representative procedure B: ethyl 1-(2,4-difluorophenyl)-6-fluoro-7-(4-methylpiperazin-1-yl)-4-oxo-1,4-dihydro-1,8-naphthyridine-3-carboxylate (16)**

To a solution of **29** (1.5 g, 3.9 mmol, 1.0 equiv.) in DMF (4.0 mL) was added $Et_3N$ (820 μL, 5.9 mmol, 1.5 equiv.) and 1-methylpiperazine (565 μL, 5.1 mmol, 1.3 equiv.). The mixture was stirred at 90 °C for 1 h. After the solvent was evaporated, the residue extracted with $CH_2Cl_2$ and $H_2O$. The organic layer was dried over $MgSO_4$ and concentrated under reduced pressure to give **16** in 96% yield. This compound was directly used in the next step without further purification. $^1$H NMR (600 MHz, chloroform-*d*) δ 8.38 (s, 1H), 8.08 (d, *J* = 13.2 Hz, 1H), 7.42 (ddd, *J* = 8.4, 8.4, 5.7 Hz, 1H), 7.09−7.00 (m, 2H), 4.35 (q, *J* = 7.1 Hz, 2H), 3.61−3.47 (m, 4H), 2.46−2.36 (m, 4H), 2.28 (s, 3H), 1.36 (t, *J* = 7.1 Hz, 3H). $^{13}$C NMR (101 MHz, chloroform-*d*) δ 173.7, 165.0, 163.0 (dd, *J* = 253.0, 11.1 Hz), 158.0 (dd, *J* = 255.3, 12.5 Hz), 149.6 (d, *J* = 9.3 Hz), 147.4, 147.2 (d, *J* = 257.7 Hz), 144.9, 130.2 (d, *J* = 10.1 Hz), 124.6 (dd, *J* = 13.1, 4.3 Hz), 121.3 (d, *J* = 22.4 Hz), 115.5 (d, *J* = 3.0 Hz), 112.9, 112.1 (dd, *J* = 22.7, 3.8 Hz), 104.9 (dd, *J* = 26.7, 23.3 Hz), 61.1, 54.6, 46.5 (d, *J* = 7.8 Hz), 45.8, 14.4. HRMS (ESI) calcd for [$C_{22}H_{21}F_3N_4O_3 + H^+$]: 447.1644; found: 447.1644. Purity: 97.3%.

**Ethyl 1-(2,4-difluorophenyl)-6-fluoro-4-oxo-7-(pyrrolidin-1-yl)-1,4-dihydro-1,8-naphthyridine-3-carboxylate (30a)**

Following representative procedure B, **30a** was prepared in 73% yield from **29** (200 mg, 0.52 mmol). $^1$H NMR (600 MHz, DMSO-$d_6$) δ 8.46 (s, 1H), 7.86 (d, *J* = 12.8 Hz, 1H), 7.74 (ddd, *J* = 8.7, 8.7, 6.0 Hz, 1H), 7.58–7.52 (m, 1H), 7.29 (ddd, *J* = 8.7, 8.7, 2.6 Hz, 1H), 4.19 (q, *J* = 7.1 Hz, 2H), 3.32 (br, 4H), 1.85–1.70 (m, 4H), 1.24 (t, *J* = 7.1 Hz, 3H). LRMS (ESI) *m/z:* 418.1 [M+H]$^+$.

**Ethyl 1-(2,4-difluorophenyl)-6-fluoro-7-morpholino-4-oxo-1,4-dihydro-1,8-naphthyridine-3-carboxylate (30b)**

Following representative procedure B, **30b** was prepared in 95% yield from **29** (1.0 g, 2.61 mmol). $^1$H NMR (600 MHz, chloroform-*d*) δ 8.42 (s, 1H), 8.16 (d, *J* = 13.4 Hz, 1H), 7.41 (ddd, *J* = 8.6, 8.6, 5.6 Hz, 1H), 7.15–6.97 (m, 2H), 4.39 (q, *J* = 7.1 Hz, 2H), 3.71–3.67 (m, 4H), 3.53–3.49 (m, 4H), 1.40 (t, *J* = 7.1 Hz, 3H). LRMS (ESI) *m/z:* 434.1 [M+H]$^+$.

**Representative procedure C for neutral form: 1-(2,4-difluorophenyl)-6-fluoro-7-(4-methylpiperazin-1-yl)-4-oxo-1,4-dihydro-1,8-naphthyridine-3-carboxylic acid (1)**

To a solution of **16** (2.0 g, 4.5 mmol, 1.0 equiv.) in ethanol (45 mL) was added lithium hydroxide (0.21 g, 9.0 mmol, 2.0 equiv.) in $H_2O$ (45 mL). The mixture was stirred at 60 °C for 5 h. After the solvent was evaporated, the residue was neutralized using HCl to give neutral form. Using general mesylate salt methods give **1•MsOH** in 73% yield. $^1$H NMR (600 MHz, DMSO-$d_6$) δ 9.92 (s, 1H), 8.91 (s, 1H), 8.26 (d, *J* = 13.0 Hz, 1H), 7.82 (ddd, *J* = 8.7, 8.7, 6.0 Hz, 1H), 7.63–7.56 (m, 1H), 7.35 (ddd, *J* = 8.3, 8.3, 2.8 Hz, 1H), 4.21–4.12 (m, 2H), 3.50–3.39 (m, 2H), 3.37–3.27 (m, 2H), 3.10–2.97 (m, 2H), 2.77 (s, 3H), 2.38 (s, 3H). $^{13}$C NMR (151 MHz, DMSO-$d_6$) δ 177.3, 165.2, 162.7 (dd, *J* = 249.6, 11.6 Hz), 157.2 (dd, *J* = 252.2, 13.4 Hz), 149.6 (d, *J* = 9.4 Hz), 148.8, 147.3 (d, *J* = 259.5 Hz), 145.4, 131.1 (d, *J* = 10.3 Hz), 123.8 (d, *J* = 13.0 Hz), 120.3 (d, *J* = 22.2 Hz), 113.1, 112.5 (d, *J* = 23.8 Hz), 109.3, 105.0 (dd, *J* = 24.5, 24.5 Hz), 51.7, 43.6, 43.5, 42.3. HRMS (ESI) calcd for [$C_{20}H_{17}F_3N_4O_3 - H^+$]: 417.1174; found: 417.1172. Purity: 99.3%.

**1-(2,4-Difluorophenyl)-6-fluoro-4-oxo-7-(pyrrolidin-1-yl)-1,4-dihydro-1,8-naphthyridine-3-carboxylic acid (2)**

Following representative procedure C, **2** was prepared in 99% yield from **30a** (60 mg, 0.14 mmol). $^1$H NMR (400 MHz, DMSO-$d_6$) δ 15.19 (s, 1H), 8.77 (s, 1H), 7.98 (d, *J* = 12.4 Hz, 1H), 7.78 (ddd, *J* = 8.5, 8.5, 6.2 Hz, 1H), 7.58 (ddd, *J* = 10.3, 8.8, 2.8 Hz, 1H), 7.38–7.26 (m, 1H), 3.70 (s, 2H), 3.15 (s, 2H), 1.82 (s, 4H). $^{13}$C NMR (101 MHz, DMSO-$d_6$) δ 176.9, 165.5, 162.5 (dd, *J* = 249.7, 11.7 Hz), 157.3 (d, *J* = 253.2, 13.6 Hz), 148.7 (d, *J* = 12.7 Hz), 147.7, 146.4, 146.1 (d, *J* = 259.6 Hz), 130.8 (d, *J* = 10.4 Hz), 124.2 (dd, *J* = 12.9, 3.9 Hz), 117.3 (d, *J* = 20.6 Hz), 112.2 (dd, *J* = 22.6, 3.5 Hz), 110.1 (d, *J* = 3.3 Hz),

108.7, 104.9 (d, $J$ = 24.5 Hz), 104.7 (d, $J$ = 24.4 Hz), 48.46. HRMS (ESI) calcd for [$C_{19}H_{14}F_3N_3O_3 - H^+$]: 388.0909; found: 388.0909. Purity: 99.5%.

**Lithium 1-(2,4-difluorophenyl)-6-fluoro-7-morpholino-4-oxo-1,4-dihydro-1,8-naphthyridine-3-carboxylic acid (3)**

Following representative procedure C but without neutralization, **3** was prepared in 81% yield from **30b** (770 mg, 1.77 mmol). $^1$H NMR (600 MHz, DMSO-$d_6$) δ 14.94 (s, 1H), 8.85 (s, 1H), 8.16 (d, $J$ = 13.2 Hz, 1H), 7.80 (ddd, $J$ = 8.6, 8.6, 6.0 Hz, 1H), 7.59 (ddd, $J$ = 10.9, 8.4, 2.7 Hz, 1H), 7.36–7.30 (m, 1H), 3.61–3.56 (m, 4H), 3.55–3.50 (m, 4H). $^{13}$C NMR (151 MHz, DMSO-$d_6$) δ 177.1, 165.2, 162.6 (dd, $J$ = 251.0, 11.3 Hz), 157.2 (dd, $J$ = 252.7, 13.2 Hz), 149.8 (d, $J$ = 7.2 Hz), 148.3, 147.0 (d, $J$ = 260.0 Hz), 145.6, 130.9 (d, $J$ = 10.0 Hz), 123.9 (d, $J$ = 10.8 Hz), 119.6 (d, $J$ = 22.4 Hz), 112.3 (d, $J$ = 23.2 Hz), 112.0, 109.0, 104.8 (dd, $J$ = 25.5, 25.5 Hz), 65.7, 46.9 (d, $J$ = 7.8 Hz). HRMS (ESI) calcd for [$C_{19}H_{14}F_3N_3O_4 + Na^+$]: 428.0834; found: 428.0839. Purity: 99.8%.

**Lithium 1-(2,4-difluorophenyl)-6-fluoro-7-hydroxy-4-oxo-1,4-dihydro-1,8-naphthyridine-3-carboxylate (4)**

Following representative procedure C but without neutralization, **4** was prepared in 59% yield from **29'** (409 mg, 1.12 mmol).$^1$H NMR (600 MHz, DMSO-$d_6$) δ 8.23 (s, 1H), 7.57 (ddd, $J$ = 8.6, 8.6, 6.1 Hz, 1H), 7.51–7.44 (m, 2H), 7.26–7.20 (m, 1H). $^{13}$C NMR (151 MHz, DMSO-$d_6$) δ 176.2, 167.5, 163.0 (d, $J$ = 16.4 Hz), 161.8 (dd, $J$ = 247.7, 11.5 Hz), 157.5 (dd, $J$ = 251.1, 13.2 Hz), 150.4 (d, $J$ = 253.5 Hz), 148.9, 145.1, 131.1 (d, $J$ = 10.0 Hz), 126.3 (d, $J$ = 13.0 Hz), 116.3, 112.9 (d, $J$ = 19.3 Hz), 111.9 (d, $J$ = 22.4 Hz), 107.7, 104.8 (dd, $J$ = 25.5, 25.5 Hz). HRMS (ESI) calcd for [$C_{15}H_7F_3N_2O_4 - H^+$]: 335.0279; found: 335.0278. Purity: 99.9%.

**1-(2,4-Difluorophenyl)-6-fluoro-7-(4-methylpiperazin-1-yl)-4-oxo-1,4-dihydro-1,8-naphthyridine-3-carboxamide (17)**

To **16** (0.3 g, 4.5 mmol, 0.6 equiv.) in sealed tube was added 7M $NH_3$ in methanol (10 mL) and stirred at 110°C for 18 h. After the solvent was evaporated, the residue was washed with methanol and collected the precipitation via filtration to give **17** in 94% yield. $^1$H NMR (400 MHz, DMSO-$d_6$) δ 9.12 (d, $J$ = 4.4 Hz, 1H), 8.60 (s, 1H), 8.07 (d, $J$ = 13.6 Hz, 1H), 7.79 (ddd, $J$ = 8.7, 8.7, 6.0 Hz, 1H), 7.64 (d, $J$ = 4.4 Hz, 1H), 7.59 (ddd, $J$ = 10.5, 9.0, 2.7 Hz, 1H), 7.36–7.28 (m, 1H), 3.52–3.43 (m, 4H), 2.31–2.25 (m, 4H), 2.13 (s, 3H). $^{13}$C NMR (151 MHz, DMSO-$d_6$) δ 175.2, 164.8, 162.4 (dd, $J$ = 248.9, 11.5 Hz), 157.4 (dd, $J$ = 251.9, 13.2 Hz), 149.3 (d, $J$ = 9.3 Hz), 146.9, 146.7 (d, $J$ = 257.5 Hz), 145.0, 130.9 (d, $J$ = 10.5 Hz), 124.4 (dd, $J$ = 12.9, 3.2 Hz), 120.2 (d, $J$ = 22.3 Hz), 113.5 (d, $J$ = 1.8 Hz), 112.9, 112.3 (dd, $J$ = 22.7, 1.8 Hz), 104.8 (dd, $J$ = 26.3, 24.9 Hz), 54.1, 46.4 (d, $J$ = 7.6 Hz), 45.5. HRMS (ESI) calcd for [$C_{20}H_{18}F_3N_5O_2 + Na^+$]: 440.1310; found: 440.1311. Purity: 87.7%.

**Ethyl 7-chloro-6-fluoro-1-(2-fluorophenyl)-4-oxo-1,4-dihydro-1,8-naphthyridine-3-carboxylate (29a)**

Following representative procedure A, **29a** was prepared in 90% yield from **28** (3.0 g, 10.7 mmol). $^1$H NMR (600 MHz, chloroform-$d$) δ 8.59 (s, 1H), 8.49 (d, $J$ = 7.2 Hz, 1H), 7.62–7.56 (m, 1H), 7.46–7.42 (m, 1H), 7.41–7.36 (m, 1H), 7.36–7.32 (m, 1H), 4.41 (q, $J$ = 7.1 Hz, 2H), 1.40 (t, $J$ = 7.1 Hz, 3H). LRMS (ESI) *m/z:* 365.0 [M+H]$^+$.

**Ethyl 7-chloro-6-fluoro-1-(4-fluorophenyl)-4-oxo-1,4-dihydro-1,8-naphthyridine-3-carboxylate (29b)**

Following representative procedure A, **29b** was prepared in 20% yield from **28** (2.0 g, 7.14 mmol). $^1$H NMR (600 MHz, chloroform-*d*) δ 8.63 (s, 1H), 8.50 (d, *J* = 7.2 Hz, 1H), 7.43–7.38 (m, 2H), 7.30–7.26 (m, 2H), 4.41 (q, *J* = 7.1 Hz, 2H), 1.40 (t, *J* = 7.1 Hz, 3H). LRMS (ESI) *m/z:* 365.0 $[M+H]^+$.

**Ethyl 7-chloro-6-fluoro-4-oxo-1-phenyl-1,4-dihydro-1,8-naphthyridine-3-carboxylate (29c)**

Following representative procedure A, **29c** was prepared in 64% yield) from **28** (1.5 g, 5.35 mmol). $^1$H NMR (600 MHz, chloroform-*d*) δ 8.67 (s, 1H), 8.50 (d, *J* = 7.3 Hz, 1H), 7.62–7.54 (m, 3H), 7.44–7.40 (m, 2H), 4.41 (q, *J* = 7.1 Hz, 2H), 1.40 (d, *J* = 7.1 Hz, 3H). LRMS (ESI) *m/z:* 347.0 $[M+H]^+$.

**Ethyl 7-chloro-1-(4,4-difluorocyclohexyl)-6-fluoro-4-oxo-1,4-dihydro-1,8-naphthyridine-3-carboxylate (29d)**

Following representative procedure A, **29d** was prepared in 42% yield from **28** (1.0 g, 3.57 mmol). $^1$H NMR (600 MHz, chloroform-*d*) δ 8.64 (s, 1H), 8.49 (d, *J* = 7.3 Hz, 1H), 5.48–5.37 (m, 1H), 4.42 (q, *J* = 7.1 Hz, 2H), 2.42–2.32 (m, 2H), 2.16–2.04 (m, 6H), 1.42 (t, *J* = 7.1 Hz, 3H). LRMS (ESI) *m/z:* 389.1 $[M+H]^+$.

**Ethyl 7-chloro-1-(2,4-dimethoxyphenyl)-6-fluoro-4-oxo-1,4-dihydro-1,8-naphthyridine-3-carboxylate (29e)**

Following representative procedure A, **29e** was prepared in 74% yield from **28** (1.5 g, 5.35 mmol). $^1$H NMR (400 MHz, chloroform-*d*) δ 8.53 (s, 1H), 8.47 (d, *J* = 7.4 Hz, 1H), 7.24–7.19 (m, 1H), 6.65–6.59 (m, 2H), 4.39 (q, *J* = 7.1 Hz, 2H), 3.90 (s, 3H), 3.72 (s, 3H), 1.39 (t, *J* = 7.1 Hz, 3H). LRMS (ESI) *m/z:* 407.1 $[M+H]^+$.

**Ethyl 7-chloro-1-(2,4-difluorobenzyl)-6-fluoro-4-oxo-1,4-dihydro-1,8-naphthyridine-3-carboxylate (29f)**

Following representative procedure A, **29f** was prepared in 85% yield from **28** (1.5 g, 5.35 mmol). $^1$H NMR (400 MHz, chloroform-*d*) δ 8.76 (d, *J* = 1.4 Hz, 1H), 8.44 (d, *J* = 7.3 Hz, 1H), 7.63–7.43 (m, 1H), 6.94–6.70 (m, 2H), 5.51 (d, *J* = 1.4 Hz, 2H), 4.40 (q, *J* = 7.1 Hz, 2H), 1.41 (t, *J* = 7.1 Hz, 3H). LRMS (ESI) *m/z:* 397.0 $[M+H]^+$.

**Ethyl 1-benzyl-7-chloro-6-fluoro-4-oxo-1,4-dihydro-1,8-naphthyridine-3-carboxylate (29g)**

Following representative procedure A, **29g** was prepared in 94% yield from **28** (1.5 g, 5.35 mmol). $^1$H NMR (600 MHz, chloroform-*d*) δ 8.68 (s, 1H), 8.45 (d, *J* = 7.6 Hz, 1H), 7.40–7.32 (m, 5H), 5.54 (s, 2H), 4.39 (q, *J* = 7.1 Hz, 2H), 1.40 (t, *J* = 7.1 Hz, 3H). LRMS (ESI) *m/z:* 361.1 $[M+H]^+$.

**Ethyl 7-chloro-1-(2,4-dimethoxybenzyl)-6-fluoro-4-oxo-1,4-dihydro-1,8-naphthyridine-3-carboxylate (29h)**

Following representative procedure A, **29h** was prepared in 71% yield from **28** (2.0 g, 7.14 mmol). $^1$H NMR (600 MHz, chloroform-*d*) δ 8.91 (s, 1H), 8.42 (d, *J* = 7.3 Hz, 1H), 7.50 (d, *J* = 8.4 Hz, 1H), 6.48 (dd, *J* = 8.4, 2.4 Hz, 1H), 6.45 (d, *J* = 2.4 Hz, 1H), 5.42 (s, 2H), 4.39 (q, *J* = 7.1 Hz, 2H), 3.84 (s, 3H), 3.79 (s, 3H), 1.41 (t, *J* = 7.1 Hz, 3H). LRMS (ESI) *m/z:* 421.1 $[M+H]^+$.

**Ethyl 7-chloro-1-(2,4-difluorophenethyl)-6-fluoro-4-oxo-1,4-dihydro-1,8-naphthyridine-3-carboxylate (29i)**

Following representative procedure A, **29i** was prepared in 71% yield from **28** (1.5 g, 5.35 mmol). $^{1}$H NMR (600 MHz, chloroform-*d*) δ 8.45 (d, *J* = 7.4 Hz, 1H), 8.28 (s, 1H), 6.99 (td, *J* = 8.2, 6.1 Hz, 1H), 6.83–6.70 (m, 2H), 4.58 (t, *J* = 6.9 Hz, 2H), 4.35 (q, *J* = 7.2 Hz, 2H), 3.18 (t, *J* = 6.9 Hz, 2H), 1.37 (t, *J* = 7.2 Hz, 3H). LRMS (ESI) *m/z:* 411.1 $[M+H]^+$.

**Ethyl 7-chloro-6-fluoro-4-oxo-1-phenethyl-1,4-dihydro-1,8-naphthyridine-3-carboxylate (29j)**

Following representative procedure A, **29j** was prepared in 32% yield from **28** (1.0 g, 3.57 mmol). $^{1}$H NMR (600 MHz, chloroform-*d*) δ 8.46 (d, *J* = 7.3 Hz, 1H), 8.25 (s, 1H), 7.31–7.27 (m, 2H), 7.25–7.22 (m, 1H), 7.12–7.09 (m, 2H), 4.59 (t, *J* = 7.1 Hz, 2H), 4.34 (q, *J* = 7.1 Hz, 2H), 3.15 (t, *J* = 7.1 Hz, 2H), 1.36 (t, *J* = 7.1 Hz, 3H). LRMS (ESI) *m/z:* 375.1 $[M+H]^+$.

**Ethyl 7-chloro-6-fluoro-4-oxo-1-(3-phenylpropyl)-1,4-dihydro-1,8-naphthyridine-3-carboxylate (29k)**

Following representative procedure A, **29k** was prepared in 14% yield from **28** (1.5 g, 5.35 mmol). $^{1}$H NMR (400 MHz, chloroform-*d*) δ 8.51 (s, 1H), 8.45 (d, *J* = 7.4 Hz, 1H), 7.34–7.27 (m, 2H), 7.24–7.17 (m, 3H), 4.45–4.33 (m, 4H), 2.74 (t, *J* = 7.4 Hz, 2H), 2.26 (dq, *J* = 8.5, 7.4 Hz, 2H), 1.42 (t, *J* = 7.1 Hz, 3H). LRMS (ESI) *m/z:* 389.1 $[M+H]^+$.

**Ethyl 7-chloro-1-((4,4-difluorocyclohexyl)methyl)-6-fluoro-4-oxo-1,4-dihydro-1,8-naphthyridine-3-carboxylate (29l)**

Following representative procedure A, **29l** was prepared in 86% yield from **28** (1.5 g, 5.35 mmol). $^{1}$H NMR (400 MHz, chloroform-*d*) δ 8.51 (s, 1H), 8.47 (d, *J* = 7.3 Hz, 1H), 4.41 (q, *J* = 7.1 Hz, 2H), 4.26 (d, *J* = 7.3 Hz, 2H), 2.26–1.94 (m, 3H), 1.80–1.66 (m, 4H), 1.52–1.45 (m, 2H), 1.42 (t, *J* = 7.1 Hz, 3H). LRMS (ESI) *m/z:* 403.1 $[M+H]^+$.

**Ethyl 6-fluoro-1-(2-fluorophenyl)-7-(4-methylpiperazin-1-yl)-4-oxo-1,4-dihydro-1,8-naphthyridine-3-carboxylate (31a)**

Following representative procedure B, **31a** was prepared in 89% yield from **29a** (500 mg, 1.37 mmol). $^{1}$H NMR (400 MHz, chloroform-*d*) δ 8.48 (s, 1H), 8.19 (d, *J* = 13.3 Hz, 1H), 7.61–7.51 (m, 1H), 7.45–7.39 (m, 1H), 7.38–7.33 (m, 1H), 7.33–7.27 (m, 1H), 4.41 (q, *J* = 7.1 Hz, 2H), 3.75–3.60 (m, 4H), 2.70–2.50 (s, 4H), 2.42 (s, 3H), 1.41 (t, *J* = 7.1 Hz, 3H). LRMS (ESI) *m/z:* 429.1 $[M+H]^+$.

**Ethyl 6-fluoro-1-(4-fluorophenyl)-7-(4-methylpiperazin-1-yl)-4-oxo-1,4-dihydro-1,8-naphthyridine-3-carboxylate (31b)**

Following representative procedure B, **31b** was prepared in 70% yield from **29b** (300 mg, 0.82 mmol). $^{1}$H NMR (400 MHz, chloroform-*d*) δ 8.48 (s, 1H), 8.14 (d, *J* = 13.4 Hz, 1H), 7.40–7.34 (m, 2H), 7.25–7.18 (m, 2H), 4.38 (q, *J* = 7.1 Hz, 2H), 3.62–3.56 (m, 4H), 2.49–2.42 (m, 4H), 2.33 (s, 3H), 1.41 (t, *J* = 7.1 Hz, 3H). LRMS (ESI) *m/z:* 429.2 $[M+H]^+$.

**Ethyl 6-fluoro-7-(4-methylpiperazin-1-yl)-4-oxo-1-phenyl-1,4-dihydro-1,8-naphthyridine-3-carboxylate (31c)**

Following representative procedure B, **31c** was prepared in 81% yield from **29c** (500 mg, 1.44 mmol). $^{1}$H NMR (400 MHz, chloroform-*d*) δ 8.54 (s, 1H), 8.18 (d, *J* = 13.3 Hz, 1H), 7.57–7.47 (m, 3H), 7.41–7.35 (m, 2H), 4.38 (q, *J* = 7.1 Hz, 2H), 3.80–3.55 (s, 4H), 2.76–2.55 (br, 4H), 2.50–2.35 (s, 3H), 1.39 (t, *J* = 7.1 Hz, 3H). LRMS (ESI) *m/z:* 411.2 $[M+H]^+$.

**Ethyl 1-(4,4-difluorocyclohexyl)-6-fluoro-7-(4-methylpiperazin-1-yl)-4-oxo-1,4-dihydro-1,8-naphthyridine-3-carboxylate (31d)**

Following representative procedure B, **31d** was prepared in 46% yield from **29d** (350 mg, 0.90 mmol). $^{1}$H NMR (600 MHz, chloroform-*d*) δ 8.47 (s, 1H), 8.13 (d, *J* = 13.8 Hz, 1H), 5.11 (s, 1H), 4.38 (q, *J* = 7.2 Hz, 2H), 3.83–3.74 (m, 4H), 2.66–2.55 (m, 4H), 2.41–2.29 (s, 5H), 2.21 – 2.05 (m, 4H), 2.04–1.88 (m, 2H), 1.39 (t, *J* = 7.2 Hz, 3H). LRMS (ESI) *m/z:* 453.2 $[M+H]^+$.

**Ethyl 1-(2,4-dimethoxyphenyl)-6-fluoro-7-(4-methylpiperazin-1-yl)-4-oxo-1,4-dihydro-1,8-naphthyridine-3-carboxylate (31e; 26 in the main text)**

Following representative procedure B, **31e** was prepared in 99% yield from **29e** (500 mg, 1.23 mmol). $^{1}$H NMR (600 MHz, chloroform-*d*) δ 8.37 (s, 1H), 8.11 (d, *J* = 13.4 Hz, 1H), 7.20–7.14 (m, 1H), 6.59–6.52 (m, 2H), 4.35 (q, *J* = 7.1 Hz, 2H), 3.88 (s, 3H), 3.69 (s, 3H), 3.57–3.47 (m, 4H), 2.44–2.34 (m, 4H), 2.28 (s, 3H), 1.37 (t, *J* = 7.1 Hz, 3H). $^{13}$C NMR (151 MHz, chloroform-*d*) δ 174.1, 165. 6, 161.4, 155.7, 149.4 (d, *J* = 9.0 Hz), 149.2, 147.1 (d, *J* = 257.1 Hz), 145.7, 129.3, 122.5, 121.2 (d, *J* = 21.9 Hz), 116.0, 111.8, 104.1, 99.1, 60.8, 55.7, 55.7, 54.6, 46.4 (d, *J* = 7.7 Hz), 45.9, 14.4. LRMS (ESI) *m/z:* 471.2 $[M+H]^+$.

**Ethyl 1-(2,4-difluorobenzyl)-6-fluoro-7-(4-methylpiperazin-1-yl)-4-oxo-1,4-dihydro-1,8-naphthyridine-3-carboxylate (31f)**

Following representative procedure B, **31f** was prepared in 71% yield from **29f** (500 mg, 1.26 mmol). $^{1}$H NMR (400 MHz, chloroform-*d*) δ 8.54 (d, *J* = 0.7 Hz, 1H), 8.12 (d, *J* = 13.4 Hz, 1H), 7.09 (ddd, *J* = 8.5, 8.5, 6.1 Hz, 1H), 6.94–6.75 (m, 2H), 5.44 (s, 3H), 4.38 (q, *J* = 7.1 Hz, 2H), 3.83–3.57 (m, 4H), 2.56–2.42 (m, 4H), 2.33 (s, 3H), 1.40 (t, *J* = 7.1 Hz, 3H). LRMS (ESI) *m/z:* 461.2$[M+H]^+$.

**Ethyl 1-benzyl-6-fluoro-7-(4-methylpiperazin-1-yl)-4-oxo-1,4-dihydro-1,8-naphthyridine-3-carboxylate (31g)**

Following representative procedure B, **31g** was prepared in 76% yield from **29g** (500 mg, 1.38 mmol). $^{1}$H NMR (600 MHz, chloroform-*d*) δ 8.53 (s, 1H), 8.13 (d, *J* = 13.5 Hz, 1H), 7.37–7.29 (m, 3H), 7.21–7.18 (m, 2H), 5.44 (s, 2H), 4.38 (q, *J* = 7.1 Hz, 2H), 3.84–3.56 (m, 4H), 2.56–2.38 (m, 4H), 2.30 (s, 3H), 1.39 (t, *J* = 7.1 Hz, 3H). LRMS (ESI) *m/z:* 425.2 $[M+H]^+$.

**Ethyl 1-(2,4-dimethoxybenzyl)-6-fluoro-7-(4-methylpiperazin-1-yl)-4-oxo-1,4-dihydro-1,8-naphthyridine-3-carboxylate (31h; 27 in the main text)**

Following representative procedure B, **31h** was prepared in 99% yield from **29h** (1.00 g, 2.37 mmol). $^{1}$H NMR (600 MHz, chloroform-*d*) δ 8.64 (s, 1H), 8.11 (d, *J* = 13.4 Hz, 1H), 7.13 (d, *J* = 8.4 Hz, 1H), 6.46 (d, *J* = 2.3 Hz, 1H), 6.40 (dd, *J* = 8.4, 2.3 Hz, 1H), 5.34 (s, 2H), 4.35 (q, *J* = 7.1 Hz, 2H), 3.82 (s, 3H), 3.81–3.79 (m, 4H), 3.78 (s, 3H), 2.61–2.55 (m, 4H), 2.38 (s, 3H), 1.38 (t, *J* = 7.1 Hz, 3H). $^{13}$C NMR (151 MHz, chloroform-*d*) δ 173.7, 165.7, 161.3, 158.7, 149.8 (d, *J* = 9.0 Hz), 148.7, 147.3 (d, *J* = 257.0 Hz), 144.7, 131.0, 121.5 (d, *J* = 21.7 Hz), 116.8, 115.8, 111.0, 104.3, 98.7, 60.7, 55.4, 55.4, 54.8, 49.6, 47.0 (d, *J* = 7.5 Hz), 46.0, 14.4. LRMS (ESI) *m/z:* 485.2 $[M+H]^+$.

**Ethyl 1-(2,4-difluorophenethyl)-6-fluoro-7-(4-methylpiperazin-1-yl)-4-oxo-1,4-dihydro-1,8-naphthyridine-3-carboxylate (31i)**

Following representative procedure B, **31i** was prepared in 60% yield from **29i** (300 mg, 0.73 mmol). $^1$H NMR (600 MHz, chloroform-*d*) δ 8.30 (s, 1H), 8.21 (d, *J* = 12.7 Hz, 1H), 7.08–7.01 (m, 1H), 6.90–6.78 (m, 2H), 4.63–4.44 (m, 4H), 4.37 (q, *J* = 7.1 Hz, 2H), 4.04–3.96(m, 2H), 3.65–3.57 (m, 2H), 3.20–3.10 (m, 4H), 2.90 (d, *J* = 4.6 Hz, 3H), 1.38 (t, *J* = 7.1 Hz, 3H). LRMS (ESI) *m/z:* 475.2 $[M+H]^+$.

**Ethyl 6-fluoro-7-(4-methylpiperazin-1-yl)-4-oxo-1-phenethyl-1,4-dihydro-1,8-naphthyridine-3-carboxylate (31j)**

Following representative procedure B, **31j** was prepared in 73% yield from **29j** (340 mg, 0.90 mmol). $^1$H NMR (600 MHz, chloroform-*d*) δ 8.17 (s, 1H), 8.13 (d, *J* = 13.4 Hz, 1H), 7.30–7.27 (m, 2H), 7.25–7.22 (m, 1H), 7.09–7.04 (m, 2H), 4.45 (t, *J* = 7.4 Hz, 2H), 4.32 (q, *J* = 7.1 Hz, 2H), 3.91–3.82 (m, 4H), 3.11 (t, *J* = 7.4 Hz, 2H), 2.74–2.63 (m, 4H), 2.44 (s, 3H), 1.36 (t, *J* = 7.1 Hz, 3H). LRMS (ESI) *m/z:* 439.2 $[M+H]^+$.

**Ethyl 6-fluoro-7-(4-methylpiperazin-1-yl)-4-oxo-1-(3-phenylpropyl)-1,4-dihydro-1,8-naphthyridine-3-carboxylate (31k)**

Following representative procedure B, **31k** was prepared in 76% yield from **29k** (250 mg, 0.64 mmol). $^1$H NMR (400 MHz, chloroform-*d*) δ 8.38 (s, 1H), 8.10 (d, *J* = 13.5 Hz, 1H), 7.34–7.27 (m, 2H), 7.25–7.19 (m, 1H), 7.18–7.14 (m, 2H), 4.37 (q, *J* = 7.1 Hz, 2H), 4.29–4.19 (m, 2H), 3.77–3.66 (m, 4H), 2.70 (t, *J* = 7.4 Hz, 2H), 2.61–2.54 (m, 4H), 2.39 (s, 3H), 2.24–2.12 (m, 2H), 1.40 (t, *J* = 7.1 Hz, 3H). LRMS (ESI) *m/z:* 453.2 $[M+H]^+$.

**Ethyl 1-((4,4-difluorocyclohexyl)methyl)-6-fluoro-7-(4-methylpiperazin-1-yl)-4-oxo-1,4-dihydro-1,8-naphthyridine-3-carboxylate (31l; 25 in the main text)**

Following representative procedure B, **31l** was prepared in 77% yield from **29l** (1.0 g, 2.48 mmol). $^1$H NMR (600 MHz, chloroform-*d*) δ 8.34 (s, 1H), 8.10 (d, *J* = 13.8 Hz, 1H), 4.36 (q, *J* = 7.2 Hz, 2H), 4.13 (d, *J* = 7.2 Hz, 2H), 3.78 (t, *J* = 4.9 Hz, 4H), 2.56 (t, *J* = 4.9 Hz, 4H), 2.35 (s, 3H), 2.18 – 2.07 (m, 2H), 2.05 – 1.96 (m, 1H), 1.75 –1.57 (m, 4H), 1.45 – 1.40 (m, 2H), 1.38 (t, *J* = 7.2 Hz, 3H). $^{13}$C NMR (151 MHz, chloroform-*d*) δ 173.5, 165.6, 149.8 (d, *J* = 9.0 Hz),147.8, 147.2 (d, *J* = 257.4 Hz), 144.5, 122.9 (dd, *J* = 239.9, 239.9 Hz), 121.6 (d, *J* = 21.8 Hz), 116.6, 111.6, 61.0, 56.3, 54.8, 46.9 (d, *J* = 7.8 Hz), 46.0, 36.0, 33.0 (dd, *J* = 24.4, 24.4 Hz), 26.7 (d, *J* = 9.8 Hz), 14.4. LRMS (ESI) *m/z:* 467.3 $[M+H]^+$.

**Ethyl 1-(2,4-difluorophenethyl)-7-((2,4-difluorophenethyl)amino)-6-fluoro-4-oxo-1,4-dihydro-1,8-naphthyridine-3-carboxylate (32a)**

Following representative procedure B, **32a** was a side product prepared in 4% yield from **29i** (1.5 g, 5.35 mmol). $^1$H NMR (600 MHz, chloroform-*d*) δ 8.21 (s, 1H), 8.08 (d, *J* = 10.5 Hz, 1H), 7.16 (ddd, *J* = 8.4, 8.4, 6.4 Hz, 1H), 6.99–6.93 (m, 1H), 6.88–6.72 (m, 4H), 5.56 (s, 1H), 4.52 (t, *J* = 7.0 Hz, 2H), 4.35 (q, *J* = 7.1 Hz, 2H), 3.84 (td, *J* = 6.8, 6.9 Hz, 2H), 3.16 (t, *J* = 7.2 Hz, 2H), 3.03 (t, *J* = 7.2 Hz, 2H), 1.38 (t, *J* = 7.1 Hz, 3H). LRMS (ESI) *m/z:* 532.2 $[M+H]^+$.

**Ethyl 1-benzyl-6-fluoro-7-morpholino-4-oxo-1,4-dihydro-1,8-naphthyridine-3-carboxylate (32b)**

Following representative procedure B, **32b** was prepared in 73% yield from **29g** (500 mg, 1.38 mmol). $^1$H NMR (400 MHz, chloroform-*d*) δ 8.53 (s, 1H), 8.16 (d, *J* = 13.4 Hz, 1H), 7.39–7.27 (m, 3H), 7.21–

7.15 (m, 2H), 5.44 (s, 2H), 4.38 (q, *J* = 7.1 Hz, 2H), 3.78–3.71 (m, 4H), 3.70–3.64 (m, 4H), 1.40 (t, *J* = 7.1 Hz, 3H). LRMS (ESI) *m/z:* 412.2 $[M+H]^+$.

**Ethyl 6-fluoro-7-morpholino-4-oxo-1,4-dihydro-1,8-naphthyridine-3-carboxylate (33)**

To a solution of **32b** (0.2 g, 0.5 mmol, 1.0 equiv.) in ethanol and $CH_2Cl_2$ (1:1) co-solvent (3 mL) was added palladium on activated charcoal (0.06 g, 0.5 mmol, 0.1 equiv.), 12 N HCl (0.3 ml) and stirred under $H_2$ at r.t. for 3 h. The resulting mixture filtered through a pad of Celite and extracted with $CH_2Cl_2$ and $H_2O$. The organic layer was dried over $MgSO_4$ and concentrated under reduced pressure. The residue was washed sequentially with acetonitrile and $CHCl_3$. The precipitation was collected via filtration to give **33** in 30% yield. $^1$H NMR (600 MHz, DMSO-$d_6$) δ 12.36 (s, 1H), 8.30 (s, 1H), 7.90 (d, *J* = 13.6 Hz, 1H), 4.18 (q, *J* = 7.1 Hz, 2H), 3.75–3.70 (m, 4H), 3.69–3.65 (m, 4H), 1.25 (t, *J* = 7.1 Hz, 3H). LRMS (ESI) *m/z:* 322.1 $[M+H]^+$.

**6-Fluoro-1-(2-fluorophenyl)-7-(4-methylpiperazin-1-yl)-4-oxo-1,4-dihydro-1,8-naphthyridine-3-carboxylic acid (5)**

Following representative procedure C and using general mesylate salt methods, **5•MsOH** was prepared in 67% yield from **31a** (250 mg, 0.58 mmol). $^1$H NMR (400 MHz, DMSO-$d_6$) δ 14.82 (s, 1H), 9.74 (s, 1H), 8.89 (s, 1H), 8.30 (d, *J* = 13.0 Hz, 1H), 7.77–7.70 (m, 1H), 7.70–7.62 (m, 1H), 7.56–7.48 (m, 1H), 7.48–7.41 (m, 1H), 4.13 (s, 2H), 3.33 (s, 4H), 3.02 (s, 2H), 2.74 (s, 3H), 2.29 (s, 3H). $^{13}$C NMR (151 MHz, DMSO-$d_6$) δ 177.1, 165.1, 156.7 (d, *J* = 250.3 Hz), 149.5 (d, *J* = 9.3 Hz), 148.5, 147.2 (d, *J* = 259.6 Hz), 145.2, 132.1 (d, *J* = 7.6 Hz), 129.6, 127.0 (d, *J* = 12.4 Hz), 125.3, 120.2 (d, *J* = 21.7 Hz), 116.2 (d, *J* = 19.3 Hz), 113.1, 109.1, 51.6, 43.5, 43.5, 42.2. HRMS (ESI) calcd for $[C_{20}H_{18}F_2N_4O_3 + H^+]$: 401.1425; found: 401.1415. Purity: 95.8%.

**6-Fluoro-1-(4-fluorophenyl)-7-(4-methylpiperazin-1-yl)-4-oxo-1,4-dihydro-1,8-naphthyridine-3-carboxylic acid (6)**

Following representative procedure C and using general mesylate salt methods, **6•MsOH** was prepared in 60% yield from **31b** (100 mg, 0.23 mmol). $^1$H NMR (400 MHz, DMSO-$d_6$) δ 9.85 (s, 1H), 8.75 (s, 1H), 8.27 (d, *J* = 13.0 Hz, 1H), 7.74–7.63 (m, 2H), 7.53–7.37 (m, 2H), 4.21–4.10 (m, 2H), 3.47–3.38 (m, 2H), 3.36–3.24 (m, 2H), 3.11–2.97 (m, 2H), 2.79–2.74 (m, 3H), 2.37–2.32 (m, 3H). $^{13}$C NMR (101 MHz, DMSO-$d_6$) δ 177.1, 165.5, 162.1 (d, *J* = 246.4 Hz), 149.4 (d, *J* = 9.5 Hz), 148.3, 147.2 (d, *J* = 259.7 Hz), 145.7, 136.1 (d, *J* = 2.9 Hz), 129.8 (d, *J* = 9.2 Hz), 120.1 (d, *J* = 22.0 Hz), 116.1 (d, *J* = 23.2 Hz), 113.5 (d, *J* = 3.7 Hz), 108.7, 51.7, 43.7 (d, *J* = 7.8 Hz), 42.3, 30.8. HRMS (ESI) calcd for $[C_{20}H_{18}F_2N_4O_3 + H^+]$: 401.1425; found: 401.1421. Purity: 99.7%.

**Lithium 6-fluoro-7-(4-methylpiperazin-1-yl)-4-oxo-1-phenyl-1,4-dihydro-1,8-naphthyridine-3-carboxylate (7)**

Following representative procedure C but without neutralization, **7** was prepared in 92% yield from **31c** (480 mg, 1.17 mmol). $^1$H NMR (600 MHz, DMSO-$d_6$) δ 8.60 (s, 1H), 8.03 (d, *J* = 13.7 Hz, 1H), 7.61–7.47 (m, 5H), 3.50–3.44 (m, 4H), 2.29–2.26 (m, 4H), 2.13 (s, 3H). $^{13}$C NMR (101 MHz, DMSO-$d_6$) δ 176.6, 167.0, 149.1 (d, *J* = 8.8 Hz), 147.8, 146.4 (d, *J* = 277.7 Hz), 145.3, 140.8, 129.1, 128.7, 127.4, 120.2 (d, *J* = 21.2 Hz), 118.0, 114.3, 54.2, 46.4 (d, *J* = 7.5 Hz), 45.6. HRMS (ESI) calcd for $[C_{20}H_{19}FN_4O_3 + Na^+]$: 405.1339; found: 405.1341. Purity: 97.9%.

**1-(4,4-Difluorocyclohexyl)-6-fluoro-7-(4-methylpiperazin-1-yl)-4-oxo-1,4-dihydro-1,8-naphthyridine-3-carboxylic acid (8)**

Following representative procedure C and using general mesylate salt methods, **8•MsOH** was prepared in 60% yield) from **31d** (180 mg, 0.39 mmol). $^1$H NMR (400 MHz, DMSO-$d_6$) δ 15.08 (s, 1H), 9.84 (s, 1H), 8.77 (s, 1H), 8.23 (d, $J$ = 13.2 Hz, 1H), 4.63–4.54 (m, 2H), 3.61–3.11 (m, 8H), 2.85 (s, 3H), 2.31(s, 3H), 2.28–2.02 (m, 7H). $^{13}$C NMR (151 MHz, DMSO-$d_6$) δ 176.1, 165.7, 149.4 (d, $J$ = 9.5 Hz), 147.1 (d, $J$ = 259.5 Hz), 144.7, 124.7, 123.1, 121.5, 120.1 (d, $J$ = 21.5 Hz), 114.0, 108.3, 52.0, 44.0, 43.9, 42.4, 32.2 (dd, $J$ = 24.5, 24.5 Hz), 26.9. HRMS (ESI) calcd for [$C_{20}H_{23}F_3N_4O_3$ + $Na^+$]: 447.1619; found: 447.1632. Purity: 92.6%.

**1-(2,4-Dimethoxyphenyl)-6-fluoro-7-(4-methylpiperazin-1-yl)-4-oxo-1,4-dihydro-1,8-naphthyridine-3-carboxylic acid (9)**

Following representative procedure C and using general mesylate salt methods, **9•MsOH** was prepared in 20% yield from **26** (160 mg, 0.34 mmol). $^1$H NMR (600 MHz, DMSO-$d_6$) δ 14.95 (s, 1H), 9.88 (s, 1H), 8.62 (s, 1H), 8.23 (d, $J$ = 12.6 Hz, 1H), 7.44 (d, $J$ = 8.7 Hz, 1H), 6.80 (d, $J$ = 2.6 Hz, 1H), 6.69 (dd, $J$ = 8.7, 2.6 Hz, 1H), 4.16 (s, 2H), 3.86 (s, 3H), 3.73 (s, 3H), 3.49–3.22 (m, 4H), 3.14–2.93 (m, 2H), 2.77 (s, 3H), 2.34 (s, 3H). $^{13}$C NMR (151 MHz, DMSO-$d_6$) δ 177.0, 165.4, 161.4, 155.0, 149.4 (d, $J$ = 9.5 Hz), 149.2, 147.1 (d, $J$ = 259.8 Hz), 145.9, 129.4, 121.1, 120.0 (d, $J$ = 21.9 Hz), 113.3 (d, $J$ = 3.9 Hz), 108.5, 105.0, 99.1, 56.1, 55.8, 51.6, 43.5, 43.5, 42.2. HRMS (ESI) calcd for [$C_{22}H_{23}FN_4O_5$ + $H^+$]: 443.1731; found: 443.1734. Purity: 98.3%.

**1-(2,4-Difluorobenzyl)-6-fluoro-7-(4-methylpiperazin-1-yl)-4-oxo-1,4-dihydro-1,8-naphthyridine-3-carboxylic acid (10)**

Following representative procedure C and using general mesylate salt methods, **10•MsOH** was prepared in 92% yield) from **31f** (200 mg, 0.43 mmol). $^1$H NMR (600 MHz, DMSO-$d_6$) δ 15.03 (s, 1H), 9.81 (s, 1H), 9.19 (s, 1H), 8.21 (d, $J$ = 13.2 Hz, 1H), 7.36 (ddd, $J$ = 8.8, 8.8, 6.4 Hz, 1H), 7.30 (ddd, $J$ = 11.0, 8.9, 2.5 Hz, 1H), 7.06 (ddd, $J$ = 8.8, 8.6, 2.5 Hz, 1H), 5.76 (s, 2H), 4.53–4.40 (m, 2H), 3.57–3.45 (m, 2H), 3.44–3.29 (m, 2H), 3.12–3.02 (m, 2H), 2.81 (s, 3H), 2.30 (s, 3H). $^{13}$C NMR (151 MHz, DMSO-$d_6$) δ 176.8, 165.6, 161.9 (dd, $J$ = 247.1, 12.5 Hz), 160.1 (dd, $J$ = 248.1, 12.5 Hz), 149.5 (d, $J$ = 9.7 Hz), 149.1, 147.1 (d, $J$ = 259.4 Hz), 144.7, 131.1 (d, J = 5.7 Hz), 131.0 (d, J = 4.9 Hz), 120.1 (d, $J$ = 22.0 Hz), 119.6 (d, $J$ = 14.3 Hz), 113.7 (d, $J$ = 3.8 Hz), 111.9 (d, $J$ = 22.0 Hz), 108.5, 104.3 (dd, $J$ = 25.7, 25.7 Hz), 51.9, 48.9, 43.8 (d, $J$ = 7.9 Hz), 42.3. HRMS (ESI) calcd for [$C_{21}H_{19}F_3N_4O_3$ + $H^+$]: 433.1487; found: 433.1489. Purity: 96.5%.

**1-Benzyl-6-fluoro-7-(4-methylpiperazin-1-yl)-4-oxo-1,4-dihydro-1,8-naphthyridine-3-carboxylic acid (11)**

Following representative procedure C and using general mesylate salt methods, **11•MsOH** was prepared in 41% yield) from **31g** (200 mg, 0.47 mmol). $^1$H NMR (600 MHz, DMSO-$d_6$) δ 15.09 (s, 1H), 9.92 (s, 1H), 9.18 (s, 1H), 8.17 (d, $J$ = 13.1 Hz, 1H), 7.39–7.26 (m, 5H), 5.73 (s, 2H), 4.55–4.39 (m, 2H), 3.58–3.31 (m, 4H), 3.03 (s, 2H), 2.79 (s, 3H), 2.35 (s, 3H). $^{13}$C NMR (151 MHz, DMSO-$d_6$) δ 176.7, 165.7, 149.4 (d, $J$ = 9.0 Hz), 148.7, 147.0 (d, $J$ = 259.4 Hz), 144.8, 136.5, 128.8, 127.8, 127.1, 120.0 (d, $J$ = 21.8 Hz), 113.8 (d, $J$ = 3.7 Hz), 108.5, 54.4, 51.9, 43.8, 43.8, 42.3. HRMS (ESI) calcd for [$C_{21}H_{21}FN_4O_3$ + $H^+$]: 397.1675; found: 397.1676. Purity: 98.9%.

**Lithium 1-(2,4-dimethoxybenzyl)-6-fluoro-7-(4-methylpiperazin-1-yl)-4-oxo-1,4-dihydro-1,8-naphthyridine-3-carboxylate (12)**

Following representative procedure C but without neutralization, **12** was prepared in 96% yield from **27** (200 mg, 0.41 mmol). $^{1}H$ NMR (600 MHz, DMSO-$d_6$) δ 8.78 (s, 1H), 7.96 (d, *J* = 13.7 Hz, 1H), 6.99 (d, *J* = 8.3 Hz, 1H), 6.57 (d, *J* = 2.3 Hz, 1H), 6.46 (dd, *J* = 8.3, 2.3 Hz, 1H), 5.40 (s, 2H), 3.76 (s, 3H), 3.72 (s, 3H), 3.69–36.4 (m, 4H), 2.41–2.36 (m, 4H), 2.18 (s, 3H). $^{13}C$ NMR (151 MHz, DMSO-$d_6$) δ 176.0, 166.3, 160.4, 158.1, 149.2 (d, *J* = 9.0 Hz), 148.8, 146.3 (d, *J* = 256.0 Hz), 144.4, 129.8, 120.1 (d, *J* = 21.1 Hz), 117.4, 116.54 114.8, 104.7, 98.5, 55.5, 55.2, 54.4, 48.7, 46.6 (d, *J* = 7.3 Hz), 45.6. HRMS (ESI) calcd for [$C_{23}H_{25}FN_4O_5 + Na^+$]: 479.1706; found: 479.1703. Purity: 98.8%.

**Lithium 1-(2,4-difluorophenethyl)-6-fluoro-7-(4-methylpiperazin-1-yl)-4-oxo-1,4-dihydro-1,8-naphthyridine-3-carboxylate (13)**

Following representative procedure C but without neutralization, **13** was prepared in 89% yield from **31i** (100 mg, 0.21 mmol). $^{1}H$ NMR (600 MHz, DMSO-$d_6$) δ 8.64 (s, 1H), 7.96 (d, *J* = 13.7 Hz, 1H), 7.38–7.31 (m, 1H), 7.18–7.10 (m, 1H), 6.99 (ddd, *J* = 8.4, 8.4, 2.1 Hz, 1H), 4.52 (t, *J* = 7.2 Hz, 2H), 3.75–3.65 (m, 4H), 3.09 (t, *J* = 7.2 Hz, 2H), 2.47–2.42 (m, 4H), 2.22 (s, 3H). $^{13}C$ NMR (151 MHz, methanol-$d_4$) δ 178.1, 171.4, 163.6 (dd, *J* = 246.9, 12.1 Hz), 162.7 (dd, *J* = 246.9, 12.1 Hz), 151.4 (d, *J* = 9.4 Hz), 149.4, 148.5 (d, *J* = 256.5 Hz), 146.2, 133.5 (d, *J* = 8.0, 8.0 Hz), 122.0 (dd, *J* = 16.0, 3.7 Hz), 121.5 (d, *J* = 22.2 Hz), 117.4, 116.5, 112.5 (dd, *J* = 21.3, 3.6 Hz), 104.6 (dd, *J* = 26.1, 26.1 Hz), 55.8, 52.3, 47.7 (d, *J* = 8.0 Hz), 46.1, 29.9. HRMS (ESI) calcd for [$C_{22}H_{21}F_3N_4O_3 + Na^+$]: 469.1463; found: 469.1464. Purity: 98.0%.

**6-Fluoro-7-(4-methylpiperazin-1-yl)-4-oxo-1-phenethyl-1,4-dihydro-1,8-naphthyridine-3-carboxylic acid (14)**

Following representative procedure C and using general mesylate salt methods, **14•MsOH** was prepared in 50% yield) from **31j** (293 mg, 0.66 mmol). $^{1}H$ NMR (600 MHz, DMSO-$d_6$) δ 15.09 (s, 1H), 9.84 (s, 1H), 8.88 (s, 1H), 8.22 (d, *J* = 13.1 Hz, 1H), 7.32–7.17 (m, 5H), 4.71 (t, *J* = 7.5 Hz, 2H), 4.56 (s, 1H), 4.05 (s, 1H), 3.67–3.19 (m, 6H)3.11 (t, *J* = 7.5 Hz, 2H), 2.83 (s, 3H), 2.30 (s, 3H). $^{13}C$ NMR (151 MHz, DMSO-$d_6$) δ 176.5, 165.6, 149.8 (d, *J* = 9.5 Hz), 148.5, 147.1 (d, *J* = 259.3 Hz), 144.8, 137.6, 129.0, 128.6, 126.7, 120.0 (d, *J* = 21.7 Hz), 113.6, 108.0, 52.8, 52.5, 44.4, 44.3, 42.9, 35.0. HRMS (ESI) calcd for [$C_{22}H_{23}FN_4O_3 + H^+$]: 411.1832; found: 411.1828. Purity: 99.1%.

**Lithium 6-fluoro-7-(4-methylpiperazin-1-yl)-4-oxo-1-(3-phenylpropyl)-1,4-dihydro-1,8-naphthyridine-3-carboxylate (15)**

Following representative procedure C but without neutralization, **15** was prepared in 94% yield from **31k** (81.4 mg, 0.18 mmol). $^{1}H$ NMR (600 MHz, DMSO-$d_6$) δ 8.73 (s, 1H), 7.96 (d, *J* = 13.8 Hz, 1H), 7.31–7.25 (m, 2H), 7.22–7.15 (m, 3H), 4.32 (t, *J* = 7.5 Hz, 2H), 3.64–3.52 (m, 4H), 2.65 (t, *J* = 7.5 Hz, 2H), 2.42–2.36 (m, 4H), 2.19 (s, 3H), 2.09–2.00 (m, 2H). $^{13}C$ NMR (151 MHz, DMSO-$d_6$) δ 176.0, 166.0, 165.9, 149.3 (d, *J* = 8.8 Hz), 148.1, 146.4 (d, *J* = 256.1 Hz), 144.2, 141.0, 128.3 (d, *J* = 4.7 Hz), 125.9, 120.1 (d, *J* = 21.4 Hz), 118.0, 114.7, 54.3, 50.2, 46.5 (d, *J* = 7.6 Hz), 45.7, 32.3, 30.9. HRMS (ESI) calcd for [$C_{23}H_{25}FN_4O_3 + Na^+$]: 447.1808; found: 447.1808. Purity: 98.6%.

**Lithium 1-((4,4-difluorocyclohexyl)methyl)-6-fluoro-7-(4-methylpiperazin-1-yl)-4-oxo-1,4-dihydro-1,8-naphthyridine-3-carboxylate (24)**

Following representative procedure C but without neutralization, **24** was prepared in 32% yield from **25** (440 mg, 0.94 mmol). $^1$H NMR (600 MHz, DMSO-$d_6$) δ 8.69 (s, 1H), 7.98 (d, *J* = 13.7 Hz, 1H), 4.27 (d, *J* = 7.0 Hz, 2H), 3.76–3.68 (m, 4H), 2.47–2.43 (m, 4H), 2.21 (s, 3H), 2.06–1.94(m, 3H), 1.81– 1.66 (m, 2H), 1.65–1.56 (m, 2H), 1.38–1.27 (m, 2H). $^{13}$C NMR (151 MHz, DMSO-$d_6$) δ 176.0, 166.1, 149.4 (d, *J* = 9.2 Hz), 148.4, 146.5 (d, *J* = 256.4 Hz), 144.5, 124.2 (dd, *J* = 240.2, 240.2 Hz), 120.1 (d, *J* = 20.7 Hz), 117.5, 114.7, 54.7, 54.4, 46.7 (d, *J* = 7.4 Hz), 45.7, 35.1, 32.3 (dd, *J* = 23.6, 23.6 Hz), 26.1 (d, *J* = 9.8 Hz). HRMS (ESI) calcd for [$C_{21}H_{25}F_3N_4O_3$ + $Na^+$]: 461.1776; found: 461.1778. Purity: 97.5%.

**Lithium 1-(2,4-difluorophenethyl)-7-((2,4-difluorophenethyl)amino)-6-fluoro-4-oxo-1,4-dihydro-1,8-naphthyridine-3-carboxylate (21)**

Following representative procedure C but without neutralization, **21** was prepared in 81% yield from **32a** (70.9 mg, 0.13 mmol). $^1$H NMR (600 MHz, DMSO-$d_6$) δ 8.53 (s, 1H), 8.05 (t, *J* = 4.8 Hz, 1H), 7.85 (d, *J* = 10.7 Hz, 1H), 7.34–7.26 (m, 2H), 7.13 (ddd, *J* = 9.7, 2.6 Hz, 1H), 7.07 (ddd, *J* = 9.7, 2.6 Hz, 1H), 6.99–6.91 (m, 2H), 4.54 (t, *J* = 7.1 Hz, 2H), 3.72–3.64 (m, 2H), 3.09 (t, *J* = 7.1 Hz, 2H), 2.94 (t, *J* = 7.3 Hz, 2H). $^{13}$C NMR (151 MHz, DMSO-$d_6$) δ 176.1, 166.2, 161.2 (dd, *J* = 245.3, 12.3 Hz), 161.0 (dd, *J* = 244.6, 12.3 Hz), 160.7 (dd, *J* = 246.1, 11.9 Hz),160.6 (dd, *J* = 247.0, 11.9 Hz), 149.6 (d, *J* = 15.5 Hz), 147.3, 145.4, 144.9 (d, *J* = 257.4 Hz), 132.4 (dd, *J* = 7.6, 7.6 Hz), 132.1 (dd, *J* = 7.9, 7.9 Hz), 122.3 (d, *J* = 16.7 Hz), 121.0 (d, *J* = 16.7 Hz), 117.8, 116.1 (d, *J* = 16.0 Hz), 112.4, 111.3 (dd, *J* = 21.5, 21.5 Hz), 111.3 (dd, *J* = 21.5, 21.5 Hz), 103.7 (dd, *J* = 26.1, 4.6 Hz), 103.5 (dd, *J* = 26.0, 4.5 Hz), 50.3, 40.5, 28.0, 27.7. HRMS (ESI) calcd for [$C_{25}H_{18}F_5N_3O_3$ + $Na^+$]: 526.1166; found: 526.1157. Purity: 98.2%.

**1-Benzyl-6-fluoro-7-morpholino-4-oxo-1,4-dihydro-1,8-naphthyridine-3-carboxylic acid (22)**

Following representative procedure C, **22** was prepared in 58% yield from **32b** (150 mg, 0.36 mmol). $^1$H NMR (600 MHz, DMSO-$d_6$) δ 15.24 (s, 1H), 9.16 (s, 1H), 8.08 (d, *J* = 13.6 Hz, 1H), 7.35–7.31 (m, 2H), 7.29–7.25 (m, 3H), 5.69 (s, 2H), 3.75–3.70 (m, 4H), 3.64–3.59 (m, 4H). $^{13}$C NMR (151 MHz, DMSO-$d_6$) δ 176.6, 165.8, 149.8 (d, *J* = 9.1 Hz), 148.4, 146.9 (d, *J* = 259.5 Hz), 145.1, 136.5, 128.7, 127.8, 127.1, 119.6 (d, *J* = 21.9 Hz), 112.8, 108.3, 65.9, 54.3, 47.2 (d, *J* = 7.7 Hz). HRMS (ESI) calcd for [$C_{20}H_{18}FN_3O_4$ + $Na^+$]: 406.1179; found: 406.1174. Purity: 95.4%.

**6-Fluoro-7-morpholino-4-oxo-1,4-dihydro-1,8-naphthyridine-3-carboxylic acid (23)**

Following representative procedure C, **23** was prepared in 49% yield from **33** (40.9 mg, 0.12 mmol). $^1$H NMR (600 MHz, DMSO-$d_6$) δ 15.30 (s, 1H), 13.33 (s, 1H), 8.53 (d, *J* = 5.5 Hz, 1H), 8.05 (d, *J* = 13.6 Hz, 1H), 3.84–3.69 (m, 8H). $^{13}$C NMR (151 MHz, DMSO-$d_6$) δ 176.9, 166.0, 150. 8 (d, *J* = 9.0 Hz), 147.1 (d, *J* = 258.6 Hz), 144.0, 118.8 (d, *J* = 21.9 Hz), 111.9 (d, *J* = 3.7 Hz), 107.8, 65.9, 47.1, 47.0. HRMS (ESI) calcd for [$C_{13}H_{12}FN_3O_4$ + $Na^+$]: 316.0709; found: 316.0713. Purity: 93.7%.

**1-(2,4-Difluorophenyl)-6-fluoro-N-hydroxy-7-(4-methylpiperazin-1-yl)-4-oxo-1,4-dihydro-1,8-naphthyridine-3-carboxamide (19)**

To a solution of **1** (0.2 g, 0.5 mmol, 1.0 equiv.) in $CH_2Cl_2$ (2.5 mL) and $Et_3N$ (0.2 mL, 1.7 mmol, 3.5 equiv.) at 0°C was added ethyl chloroformate (93 µL, 1.0 mmol, 2.0 equiv.) and stirred for 30 minutes. The mixture was subsequently added hydroxylamine hydrochloride (50.7 mg, 0.8 mmol, 1.5 equiv.) and stirred at r.t. for 16 h. The resulting reaction directly extracted with $CH_2Cl_2$ and $H_2O$. The organic layer was dried over $MgSO_4$ and concentrated under reduced pressure. The residue was washed with ether and the precipitation was collected via filtration. The solid was washed with few $CH_2Cl_2$ again and collected the precipitation to give **19** in 41% yield. $^1$H NMR (600 MHz, DMSO-$d_6$) δ 11.57 (d, *J* = 1.8 Hz, 1H),

9.32 (d, *J* = 1.8 Hz, 1H), 8.60 (s, 1H), 8.08 (d, *J* = 13.5 Hz, 1H), 7.80 (ddd, J = 8.7, 8.7, 6.0 Hz, 1H), 7.59 (ddd, J = 11.0, 8.3, 2.7 Hz, 1H), 7.35–7.30 (m, 1H), 3.50 (s, 4H), 2.32 (s, 4H), 2.16 (s, 3H). $^{13}$C NMR (151 MHz, DMSO-$d_6$) δ 174.5, 162.4 (dd, *J* = 249.2, 11.7 Hz), 161.5, 157.3 (dd, *J* = 251.9, 13.2 Hz), 149.3 (d, *J* = 9.3 Hz), 146.7 (d, *J* = 257.9 Hz), 145.9, 144.9, 131.0 (d, *J* = 10.0 Hz), 124.4 (dd, *J* = 12.7, 3.3 Hz), 120.2 (d, *J* = 22.1 Hz), 113.1, 112.4 (dd, *J* = 19.9, 2.4 Hz), 112.3, 104.8 (dd, *J* = 26.9, 24.4 Hz), 53.9, 46.1 (d, *J* = 7.6 Hz), 45.2. HRMS (ESI) calcd for [$C_{20}H_{18}F_3N_5O_3$ + $Na^+$]: 456.1259; found: 456.1255. Purity: 97.5%.

**1-(2,4-Difluorophenyl)-6-fluoro-N-methoxy-N-methyl-7-(4-methylpiperazin-1-yl)-4-oxo-1,4-dihydro-1,8-naphthyridine-3-carboxamide (34)**

To a solution of **1** (1.8 g, 4.3 mmol, 1.0 equiv.) in $CH_2Cl_2$ (22 mL) and $Et_3N$ (2.4 mL, 17.5 mmol, 4.0 equiv.) at 0°C was added ethyl chloroformate (837 μL, 8.7 mmol, 2.0 equiv.) and stirred for 30 minutes. The mixture was subsequently added *N*, *O*-dimethylhydroxylamine hydrochloride (0.8 g, 8.7 mmol, 2.0 equiv.) and stirred at r.t. for 16 h. The resulting reaction directly extracted with $CH_2Cl_2$ and $H_2O$. The organic layer was dried over $MgSO_4$ and concentrated under reduced pressure to give **34** in 99% yield. This compound was directly used in the next step without further purification. $^1$H NMR (400 MHz, chloroform-*d*) δ 8.11 (d, *J* = 13.2 Hz, 1H), 7.84 (s, 1H), 7.45–7.30 (m, 1H), 7.10–6.96 (m, 2H), 3.77 (s, 3H), 3.69–3.55 (s, 4H), 3.34 (s, 3H), 2.65–2.45 (s, 4H), 2.38 (s, 3H). LRMS (ESI) *m/z:* 462.2 $[M+H]^+$.

**1-(2,4-Difluorophenyl)-6-fluoro-7-(4-methylpiperazin-1-yl)-4-oxo-1,4-dihydro-1,8-naphthyridine-3-carbaldehyde (18)**

To a solution of **19** (0.15 g, 0.3 mmol, 1.0 equiv.) in $CH_2Cl_2$ (6.5 mL) at -55°C was added 1.2 M DIBAL in toluene (570 μL, 16.5 mmol, 5.5 equiv.) and stirred at -55°C for 18 h. The resulting reaction was quenched with sat. $NH_4Cl$ (aq., 12 mL) and stirred at r.t. for 1 h. The mixture filtered through a pad of Celite and the filtrate was extracted with $CH_2Cl_2$ and $H_2O$. The organic layer was dried over $MgSO_4$ and concentrated under reduced pressure. The residue was purified by ACCQ Prep (30-95% methanol in $H_2O$) to give **18** in 6% yield. $^1$H NMR (400 MHz, chloroform-*d*) δ 10.41 (s, 1H), 8.25 (s, 1H), 8.16 (d, *J* = 13.2 Hz, 1H), 7.44–7.34 (m, 1H), 7.12–7.00 (m, 2H), 3.74–3.58 (m, 4H), 2.56–2.44 (m, 4H), 2.41–2.33 (s, 3H). $^{13}$C NMR (101 MHz, chloroform-*d*) δ 189.3 176.2 163.2 (dd, *J* = 253.6, 11.1 Hz), 157.9 (dd, *J* = 255.7, 12.4 Hz), 149.7 (d, *J* = 9.1 Hz), 147.3 (d, *J* = 259.0 Hz), 145.6, 145.0, 130.0 (d, *J* = 10.1 Hz), 124.4 (dd, *J* = 13.0, 4.1 Hz), 120.9 (d, *J* = 22.4 Hz), 118.4, 115.9, 112.2 (dd, *J* = 22.6, 3.8 Hz), 105.1 (dd, *J* = 26.6, 23.3 Hz), 54.5, 46.3 (d, *J* = 8.2 Hz), 45.6. HRMS (ESI) calcd for [$C_{20}H_{17}F_3N_4O_2$ + $Na^+$]: 425.1201; found: 425.1203. Purity: 98.6%.

**1-(2,4-Difluorophenyl)-6-fluoro-7-(4-methylpiperazin-1-yl)-1,8-naphthyridin-4(1H)-one (20)**

To a solution of **1** (0.2 g, 0.5 mmol, 1.0 equiv.) in DMSO (1 mL) was added potassium cyanide (0.03 g, 0.5 mmol, 1.0 equiv.) and stirred at 160 °C for 3 h. After cooling down, the reaction was added $H_2O$ (8 mL) and 1 N NaOH (0.8 mL). The mixture was extract with ethyl acetate and $H_2O$. The organic layer was dried over $MgSO_4$ and concentrated under reduced pressure. The residue was purified by column chromatography (MeOH/$CH_2Cl_2$ = 1:29) to give **20** in 50% yield. $^1$H NMR (400 MHz, DMSO-$d_6$) δ 9.87 (s, 1H), 8.08–7.99 (m, 2H), 7.70 (ddd, *J* = 8.8, 8.8, 6.0 Hz, 1H), 7.57 (ddd, *J* = 10.4, 9.0, 2.7 Hz, 1H), 7.35–7.26 (m, 1H), 6.17 (d, *J* = 7.9 Hz, 1H), 4.03 (s, 2H), 3.47–2.86 (m, 6H), 2.74 (s, 3H). $^{13}$C NMR (101 MHz, DMSO-$d_6$) δ 176.3, 162.0 (dd, *J* = 248.5, 11.7 Hz), 157.4 (dd, *J* = 251.6, 13.3 Hz), 148.4 (d, *J* = 9.6 Hz), 146.4 (d, *J* = 255.7 Hz), 145.1, 142.8, 131.2 (d, *J* = 10.3 Hz), 124.6 (dd, *J* = 12.9, 4.1 Hz),

120.4 (d, $J$ = 20.7 Hz), 114.3, 112.2 (dd, $J$ = 22.5, 3.6 Hz), 110.4, 104.8 (dd, $J$ = 25.8, 25.8 Hz), 51.6, 43.7, 43.6, 42.2. HRMS (ESI) calcd for [$C_{19}H_{17}F_3N_4O + Na^+$]: 397.1252; found: 397.1248. Purity: 99.8%.

$^{1}H$ NMR (400 MHz, DMSO-$d_6$) of **1**.

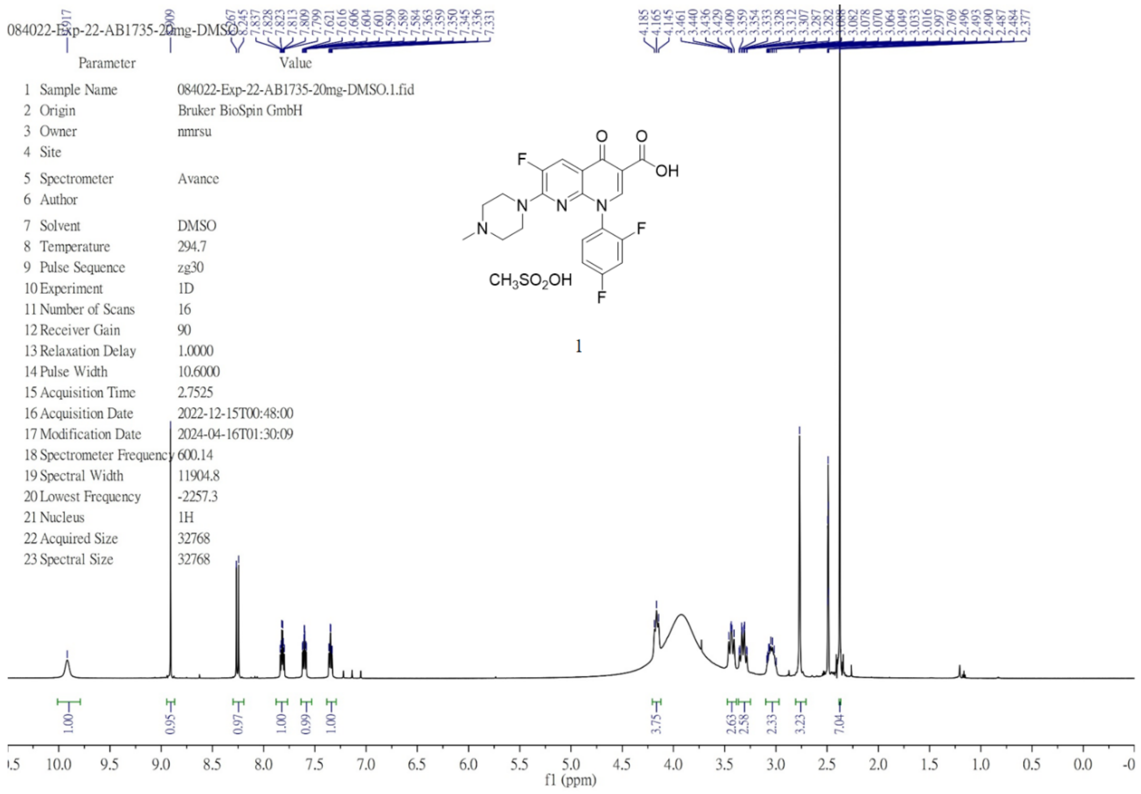


$^{13}C$ NMR (101 MHz, DMSO-$d_6$) of **1**.

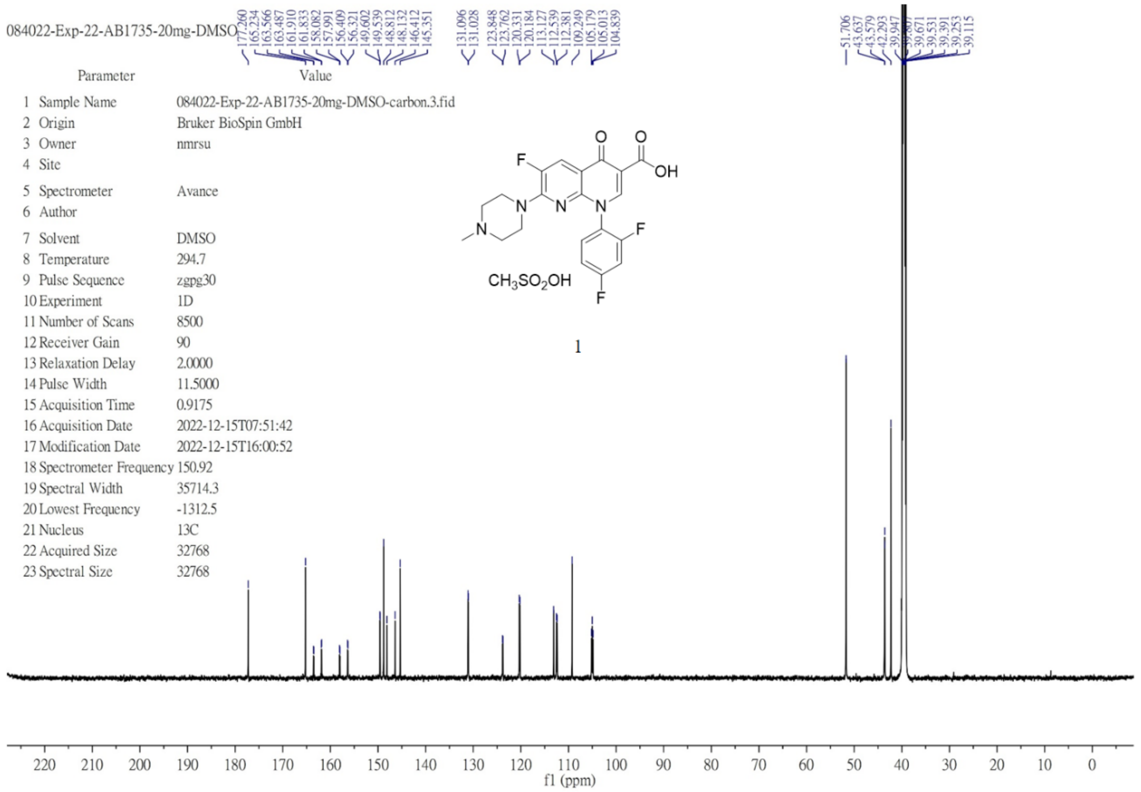

$^{1}H$ NMR (400 MHz, DMSO-$d_6$) of **2**.

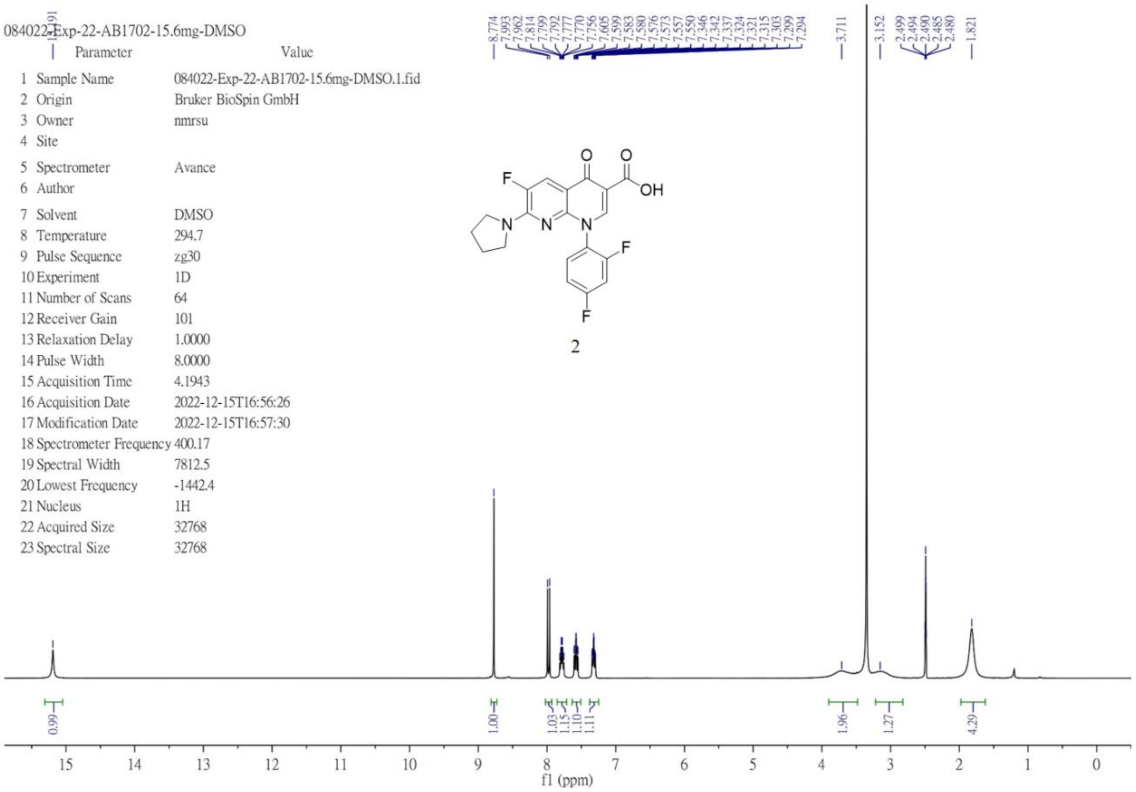


$^{13}C$ NMR (101 MHz, DMSO-$d_6$) of **2**.

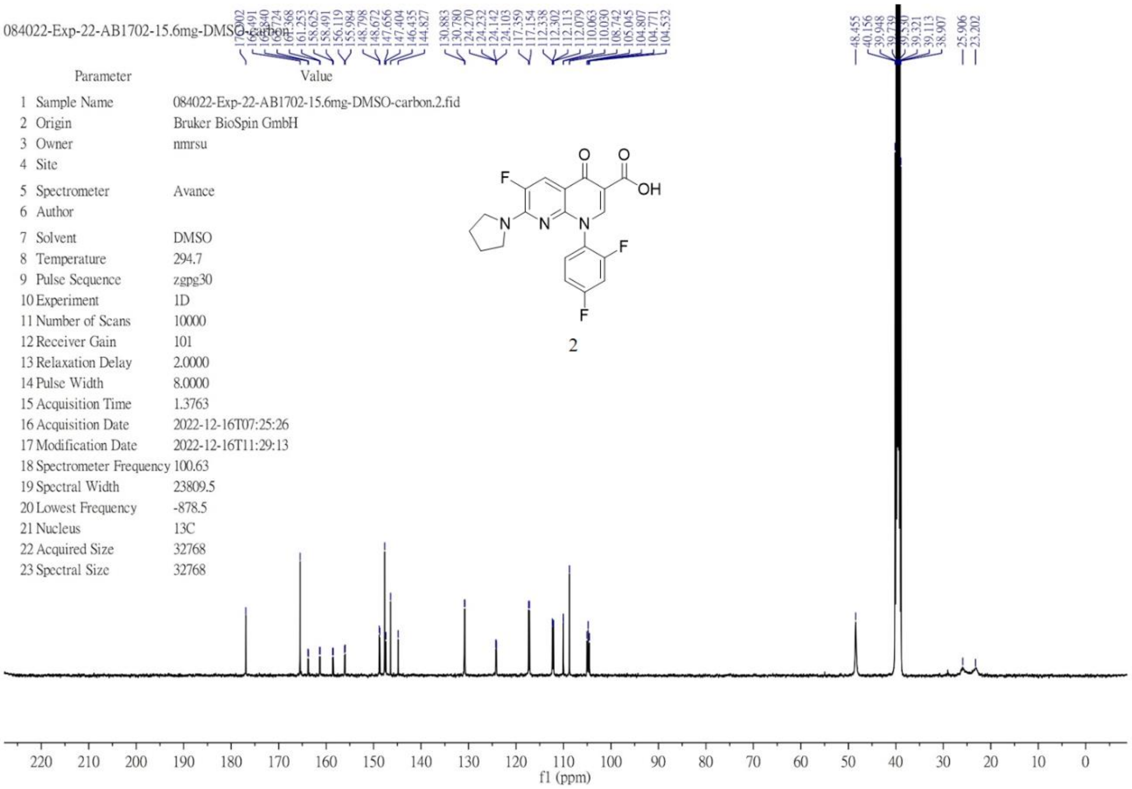

[1]H NMR (600 MHz, DMSO-$d_6$) of **3**.

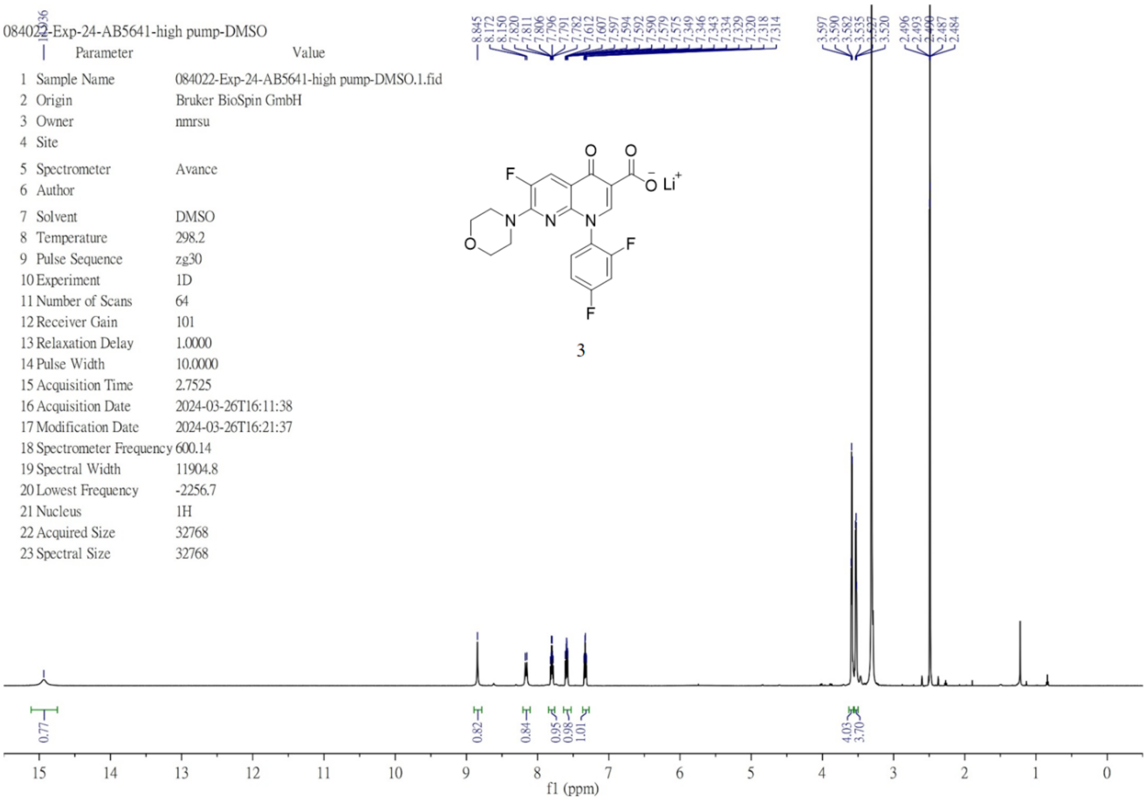


[13]C NMR (151 MHz, DMSO-$d_6$) of **3**.

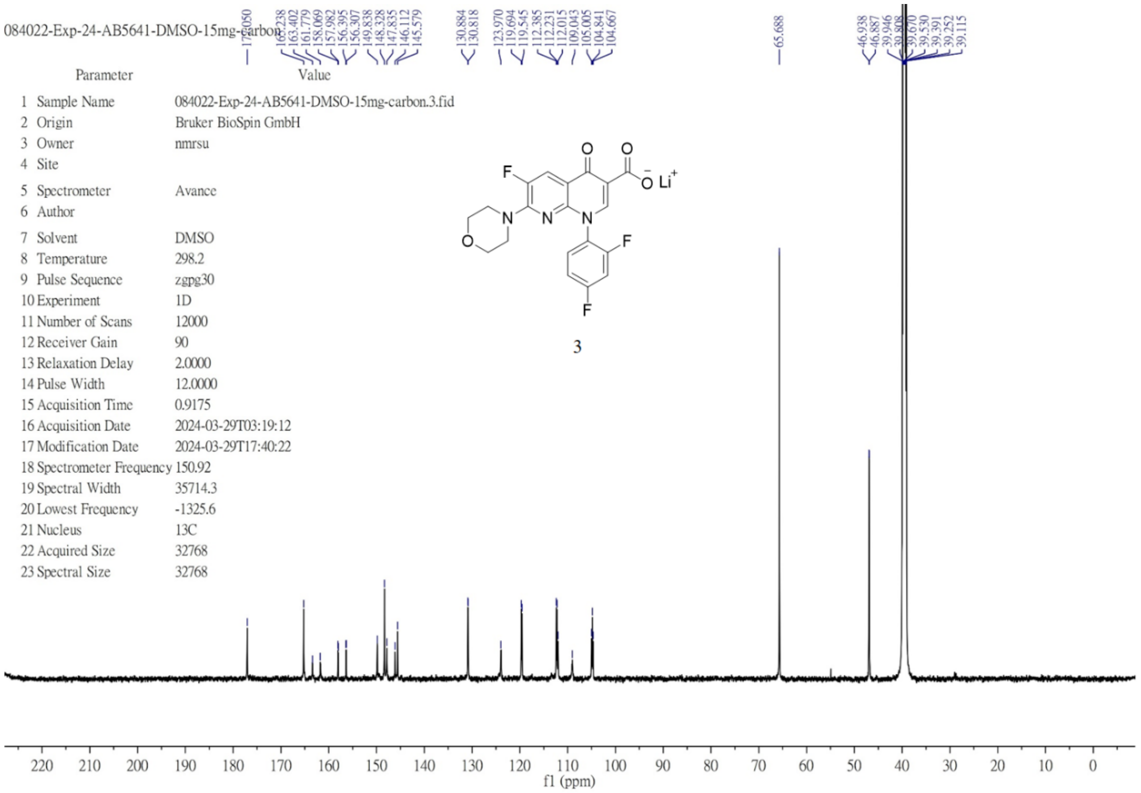

$^1H$ NMR (600 MHz, DMSO-$d_6$) of **4**.

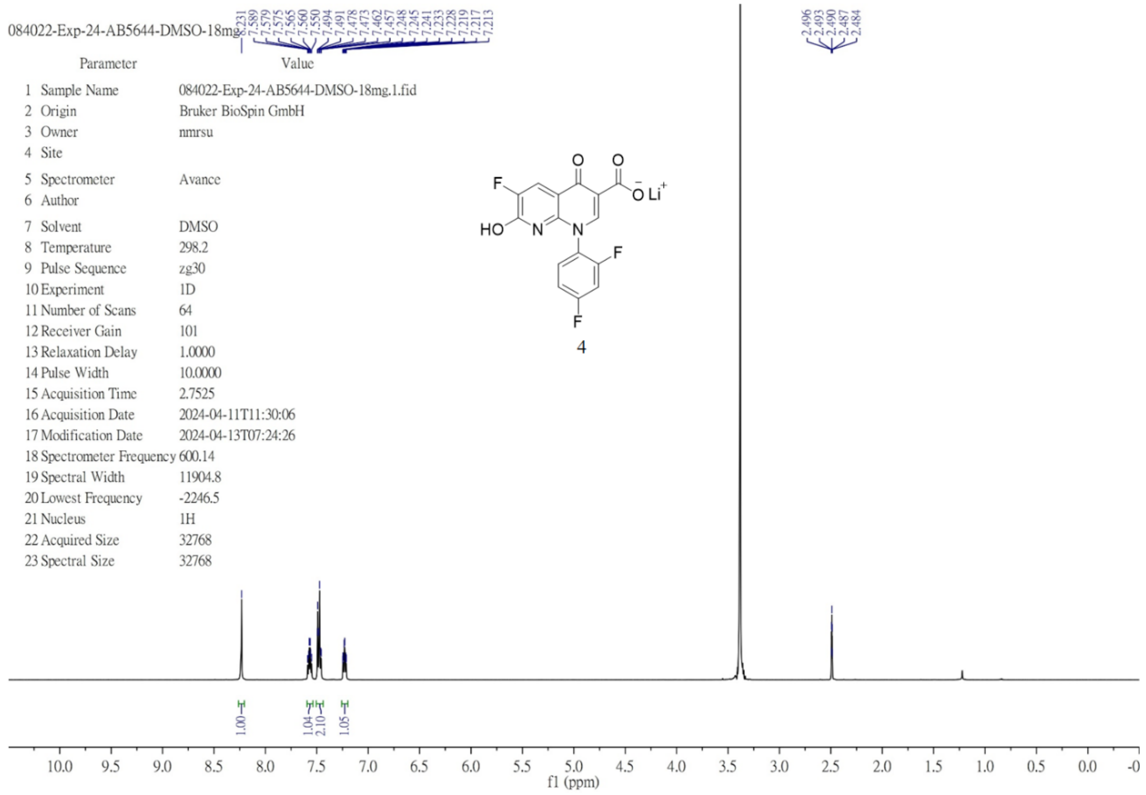


$^{13}C$ NMR (151 MHz, DMSO-$d_6$) of **4**.

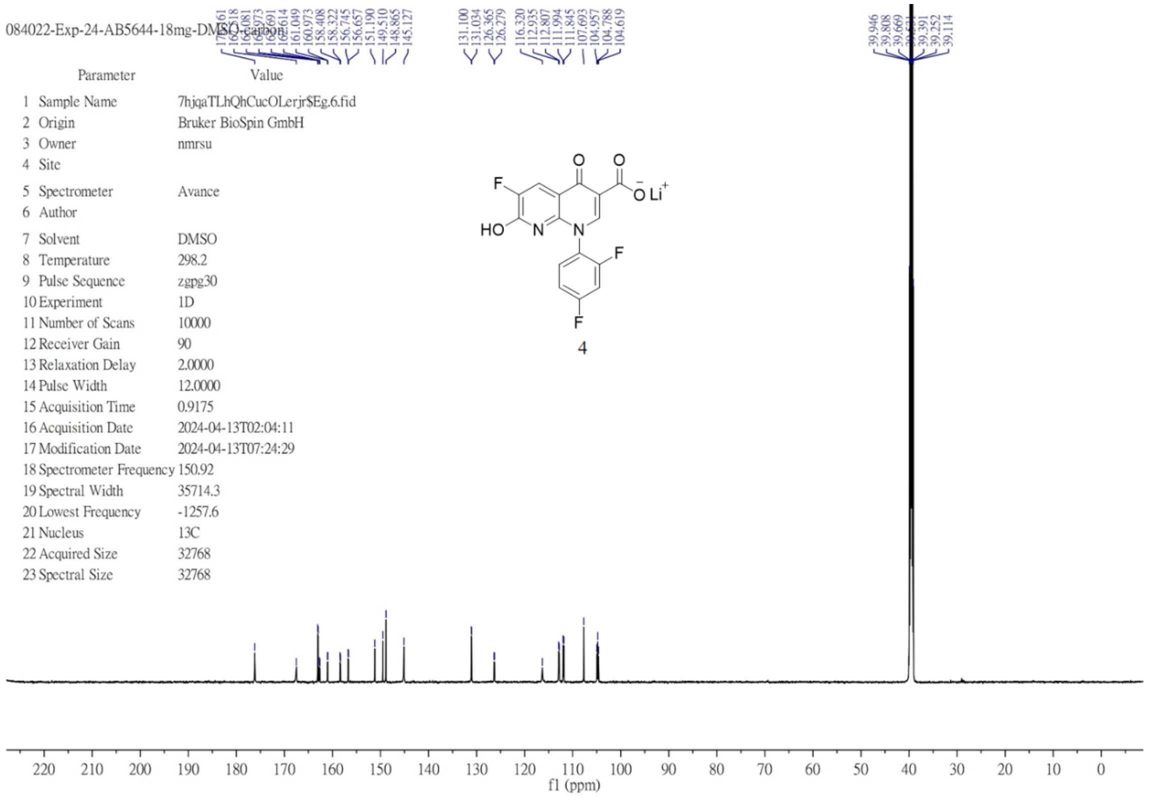

$^{1}H$ NMR (400 MHz, DMSO-$d_6$) of **5**.

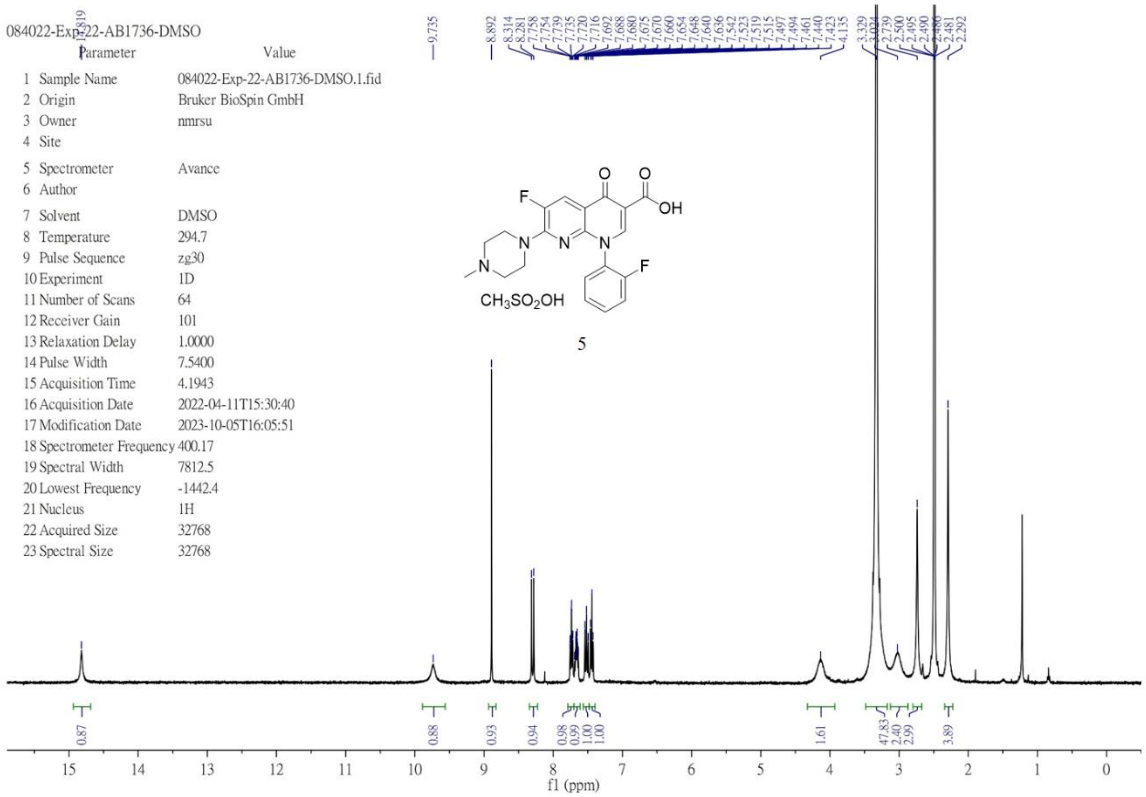


$^{13}C$ NMR (151 MHz, DMSO-$d_6$) of **5**.

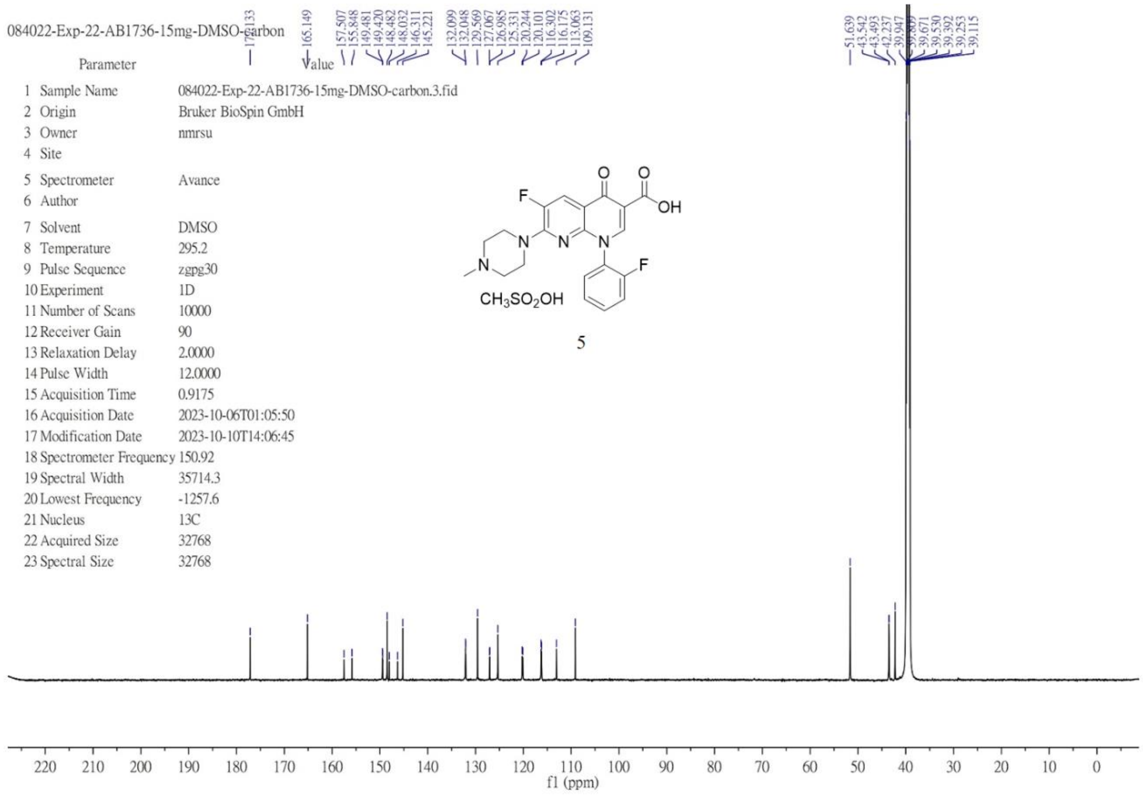

$^{1}$H NMR (400 MHz, DMSO-$d_6$) of **6**.

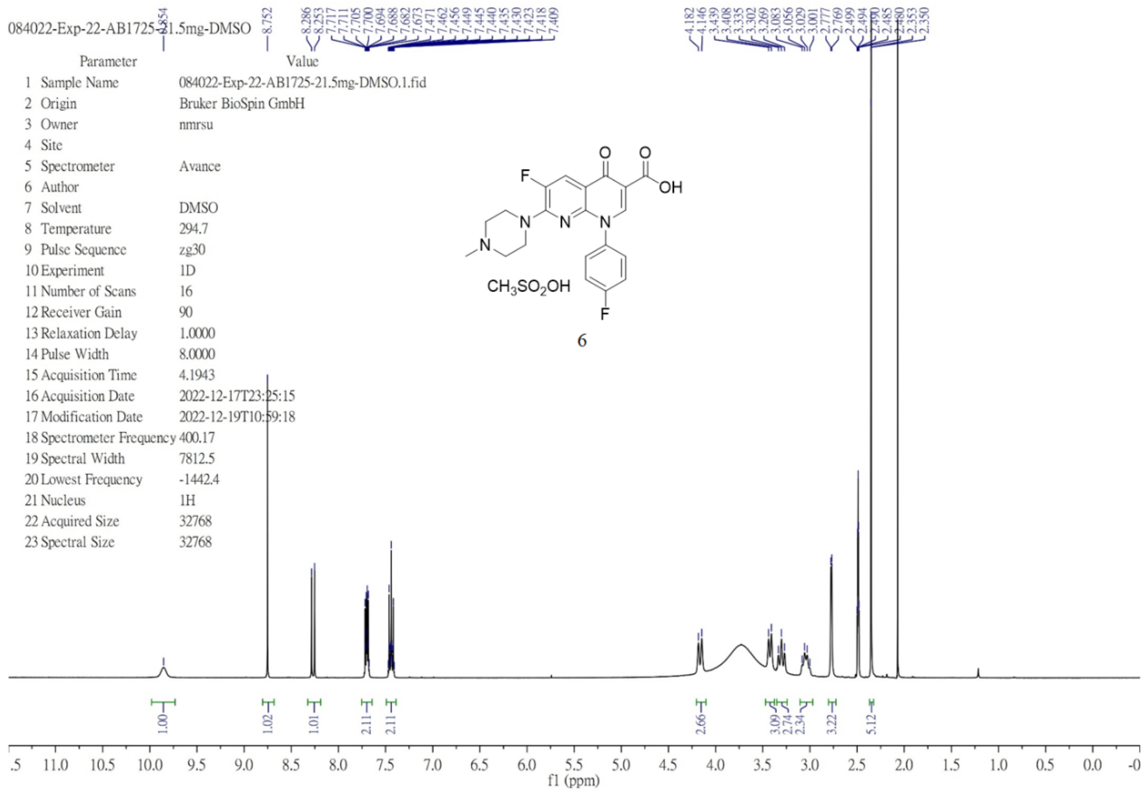


$^{13}$C NMR (101 MHz, DMSO-$d_6$) of **6**.

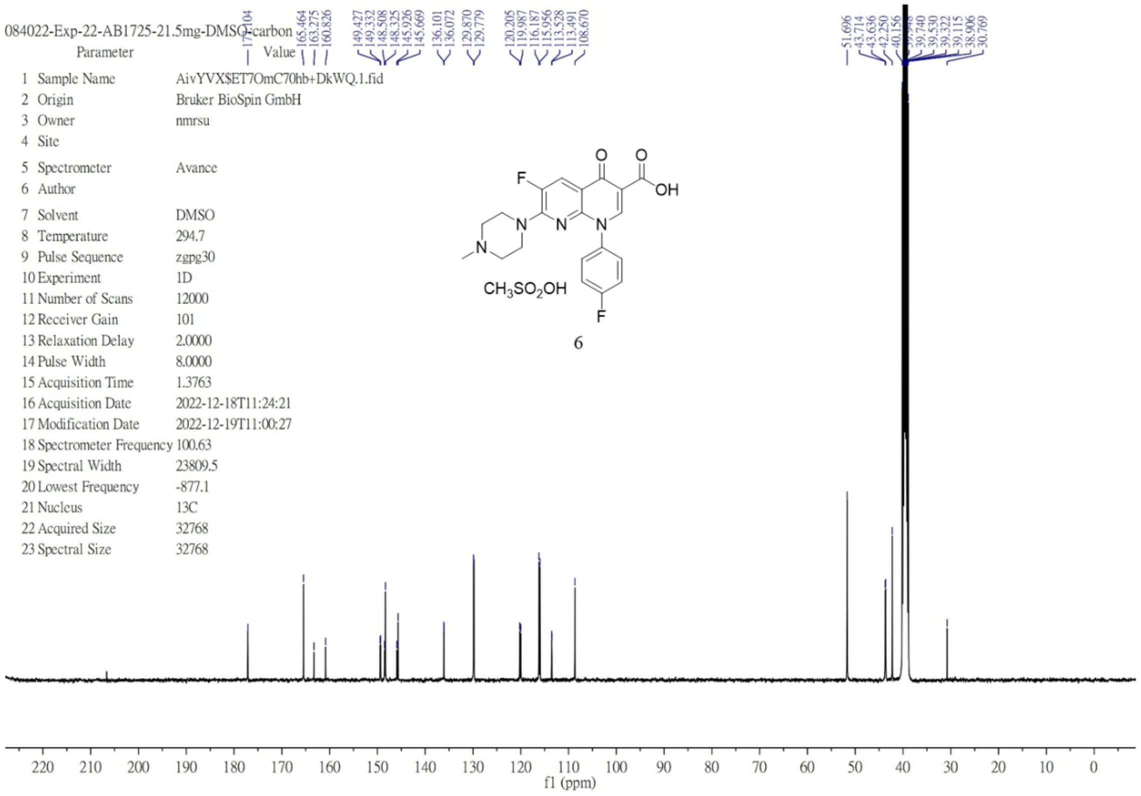

[1]H NMR (600 MHz, DMSO-$d_6$) of **7**.

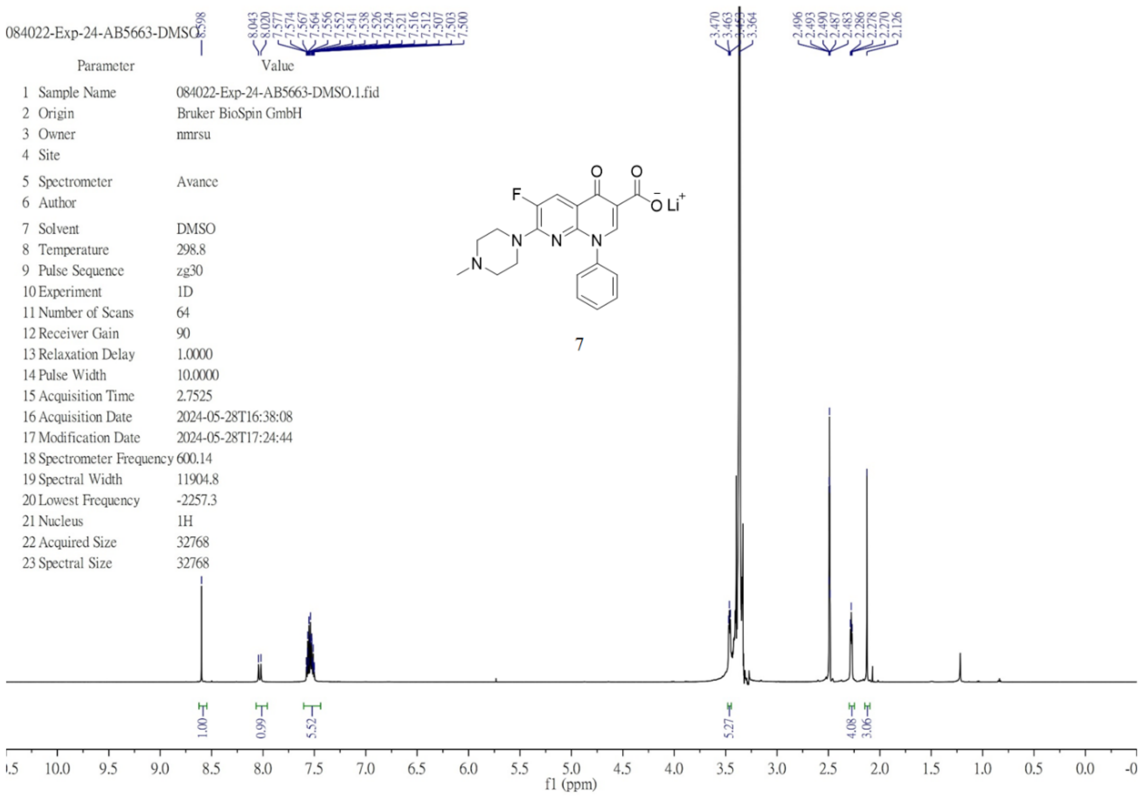

[13]C NMR (101 MHz, DMSO-$d_6$) of **7**.

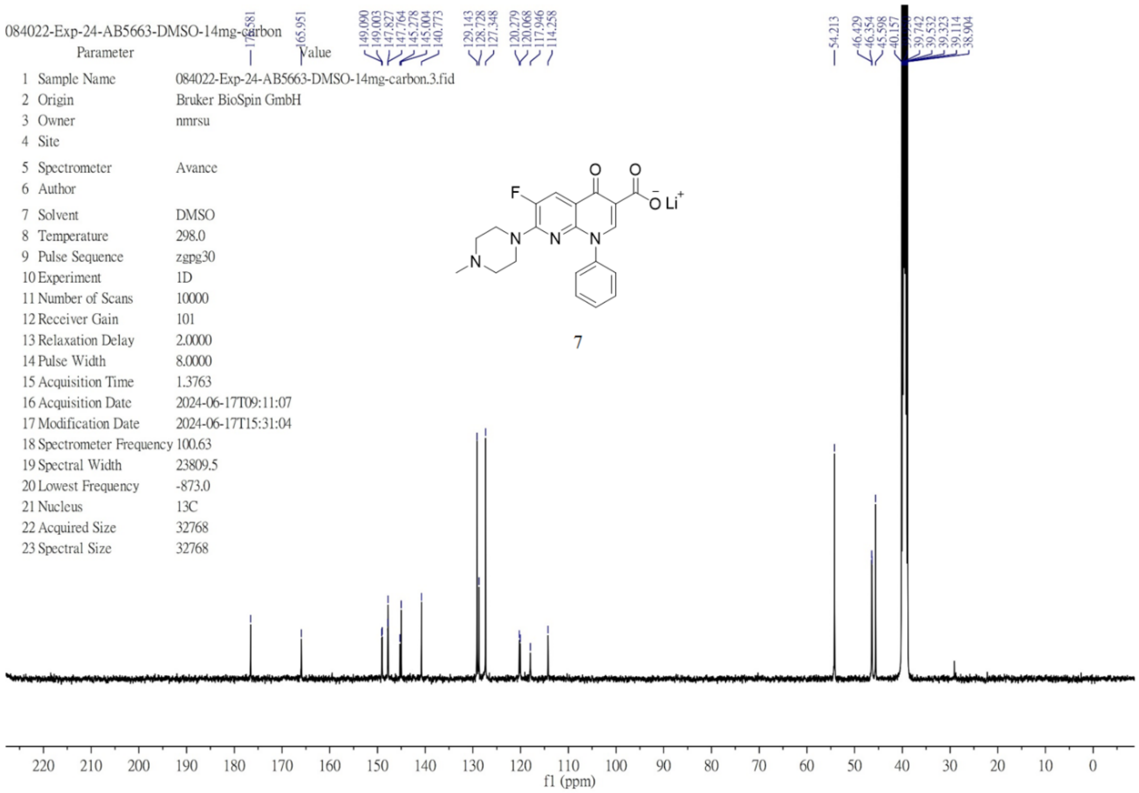

$^{1}H$ NMR (400 MHz, DMSO-$d_6$) of **8**.

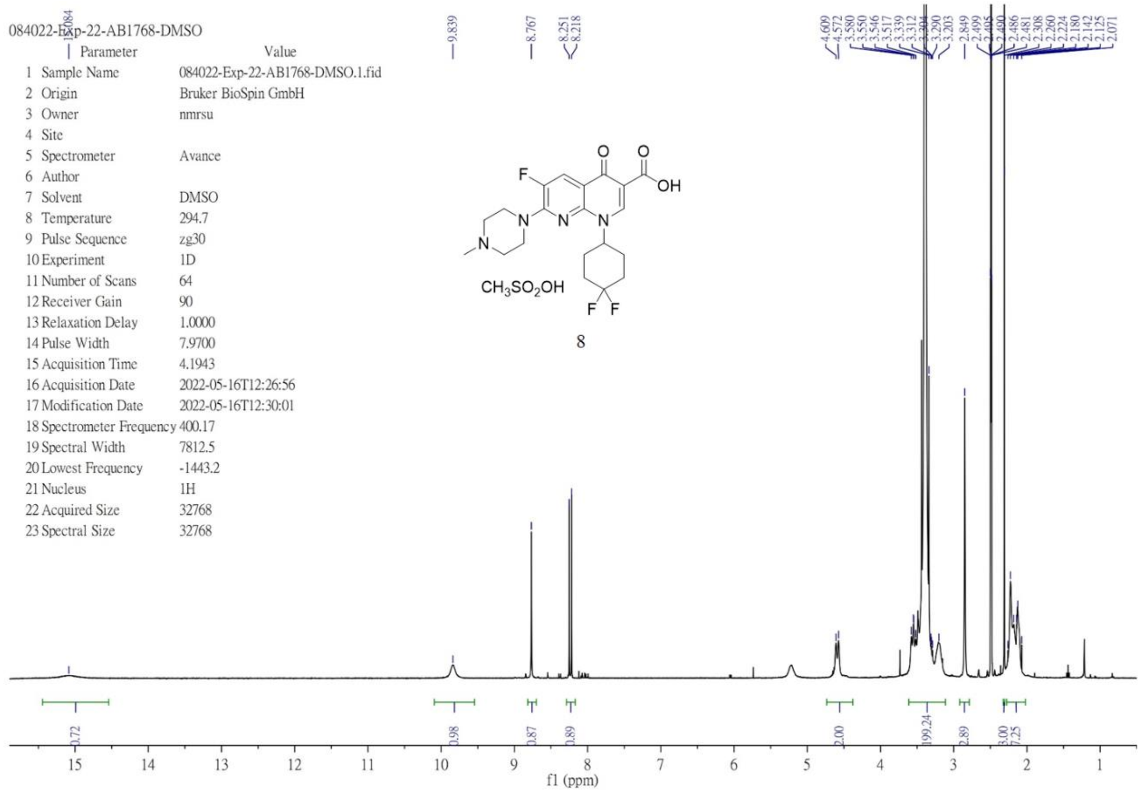

$^{13}C$ NMR (151 MHz, DMSO-$d_6$) of **8**.

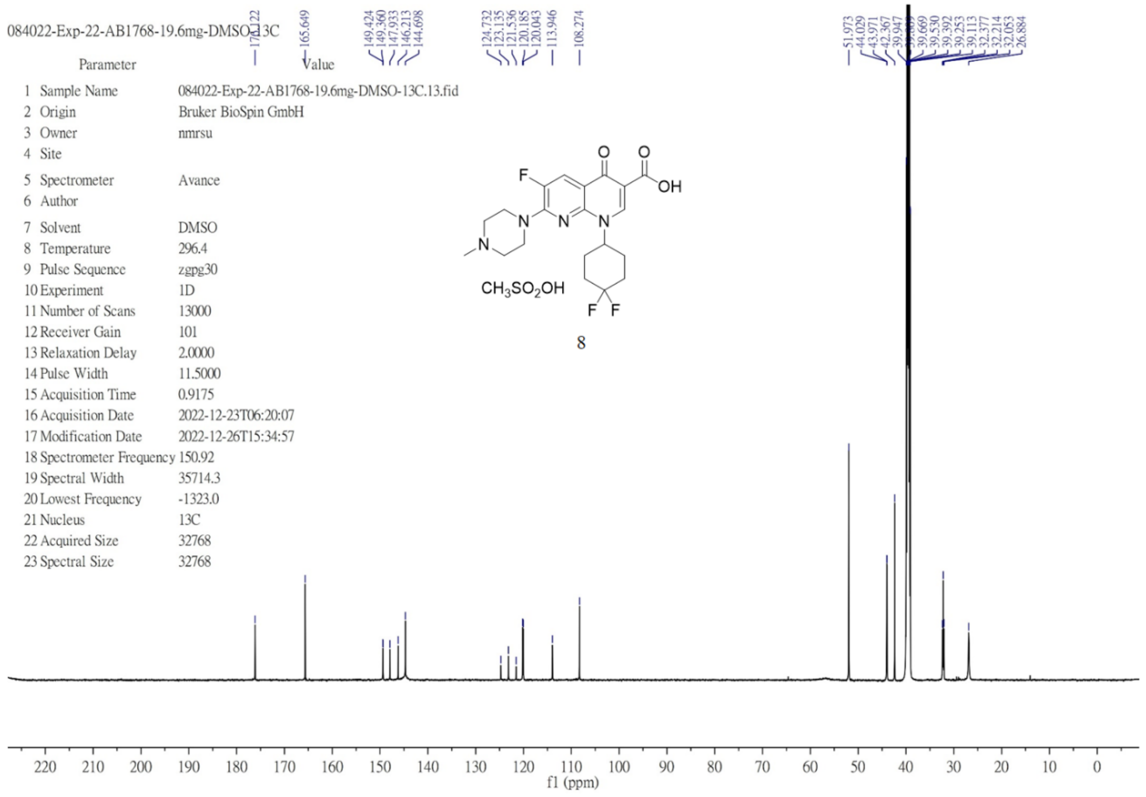

$^{1}$H NMR (600 MHz, DMSO-$d_6$) of **9**.

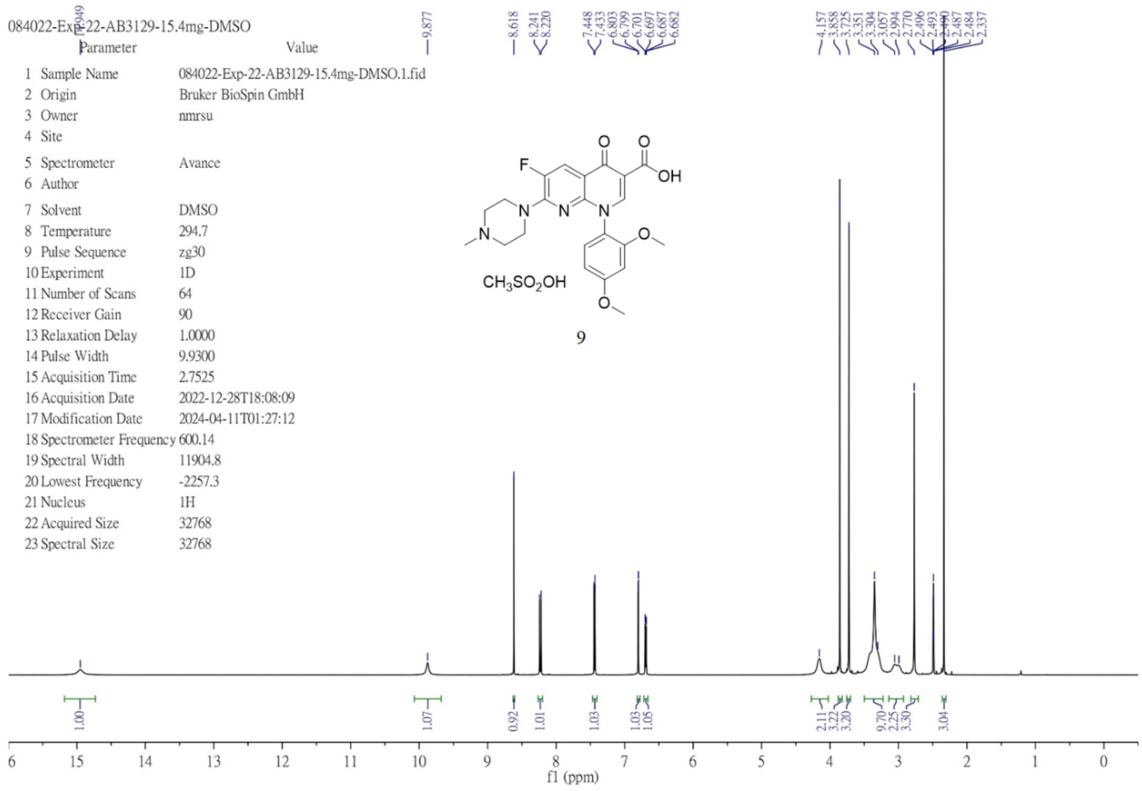


$^{13}$C NMR (151 MHz, DMSO-$d_6$) of **9**.

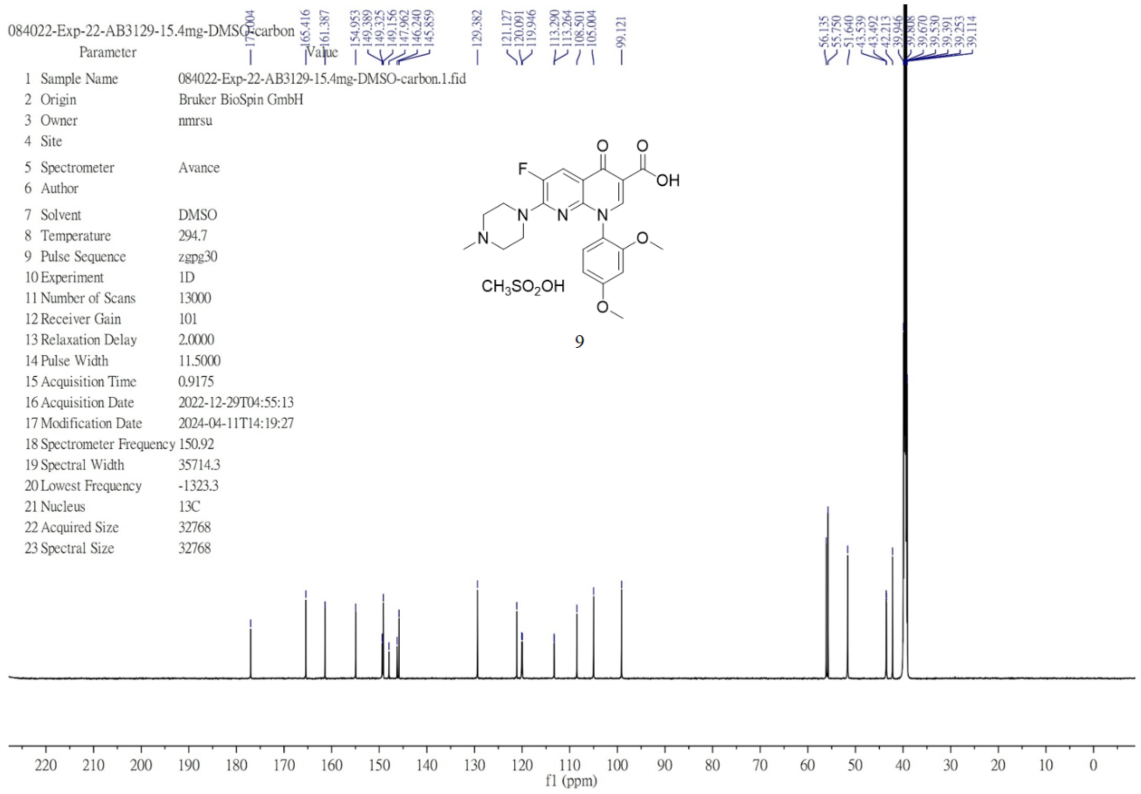

$^{1}H$ NMR (600 MHz, DMSO-$d_6$) of **10**.

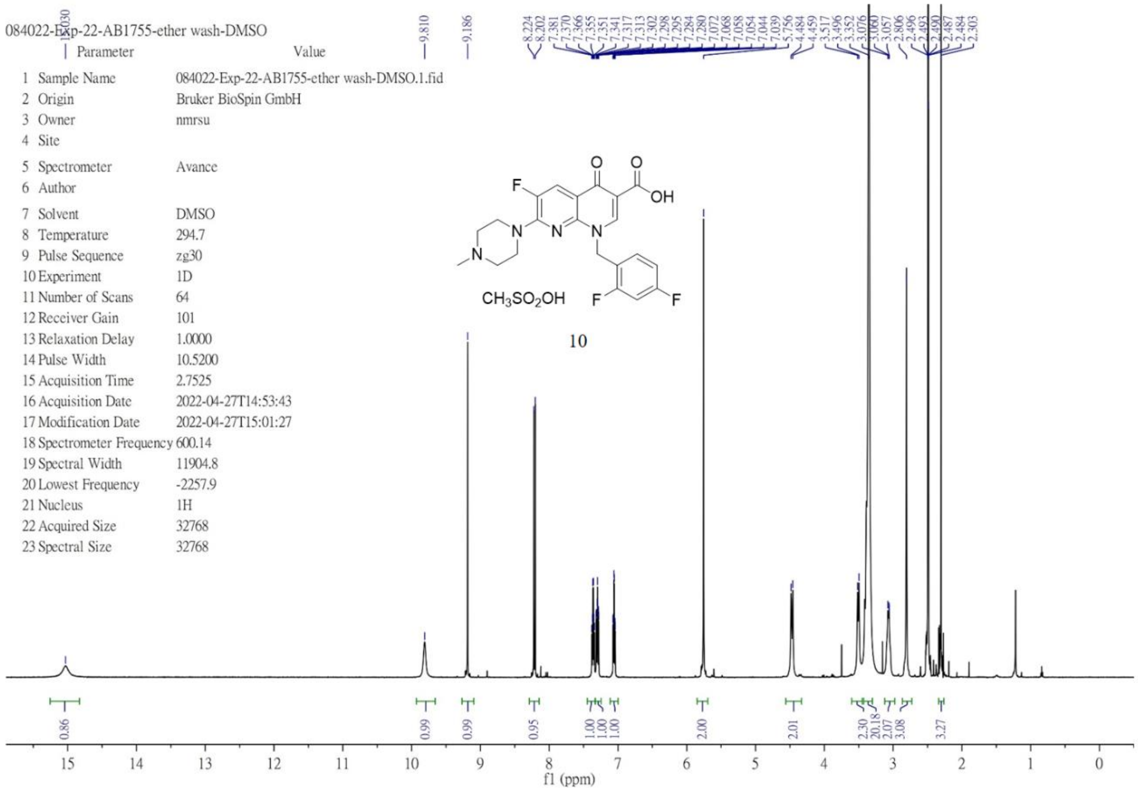


$^{13}C$ NMR (151 MHz, DMSO-$d_6$) of **10**.

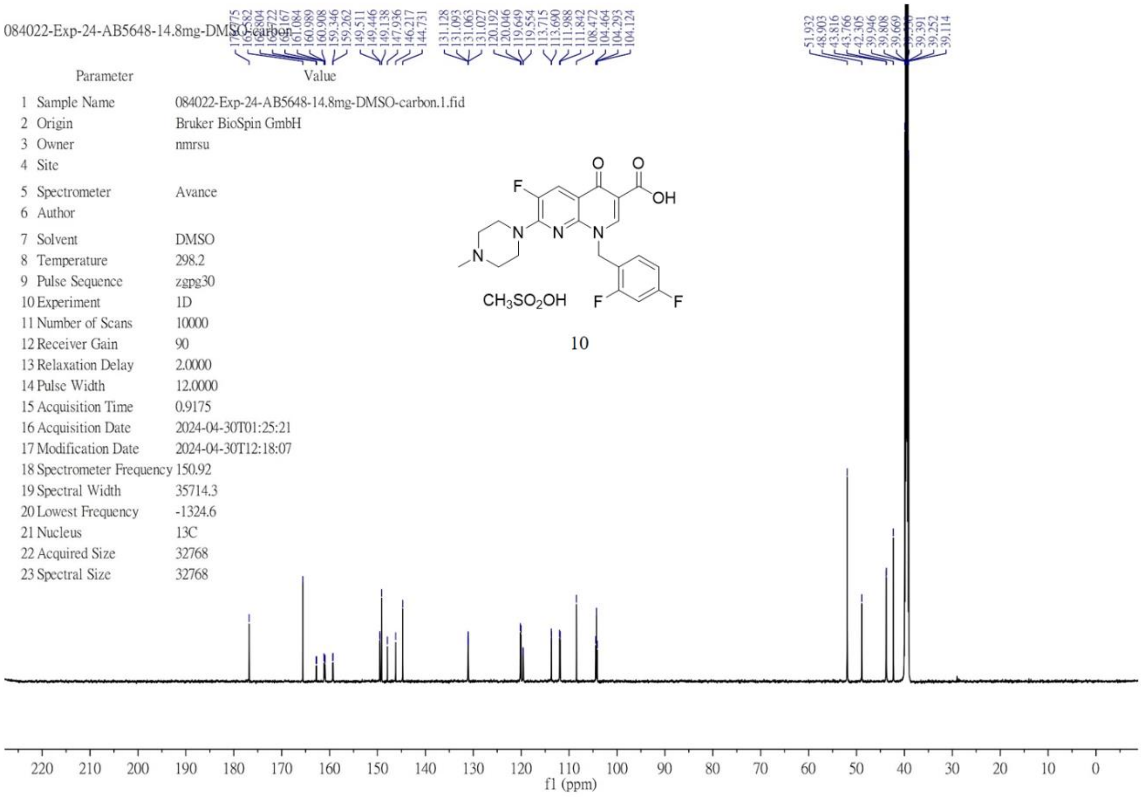

$^{1}H$ NMR (600 MHz, DMSO-$d_6$) of **11**.

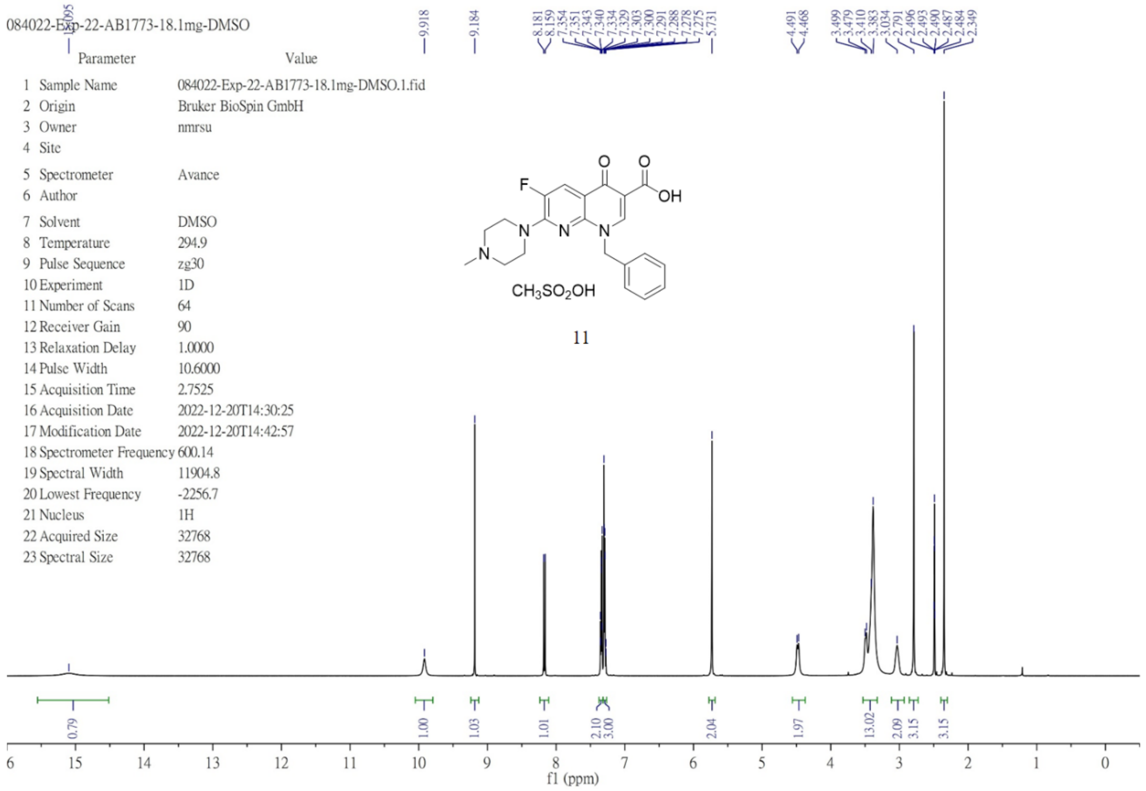


$^{13}C$ NMR (151 MHz, DMSO-$d_6$) of **11**.

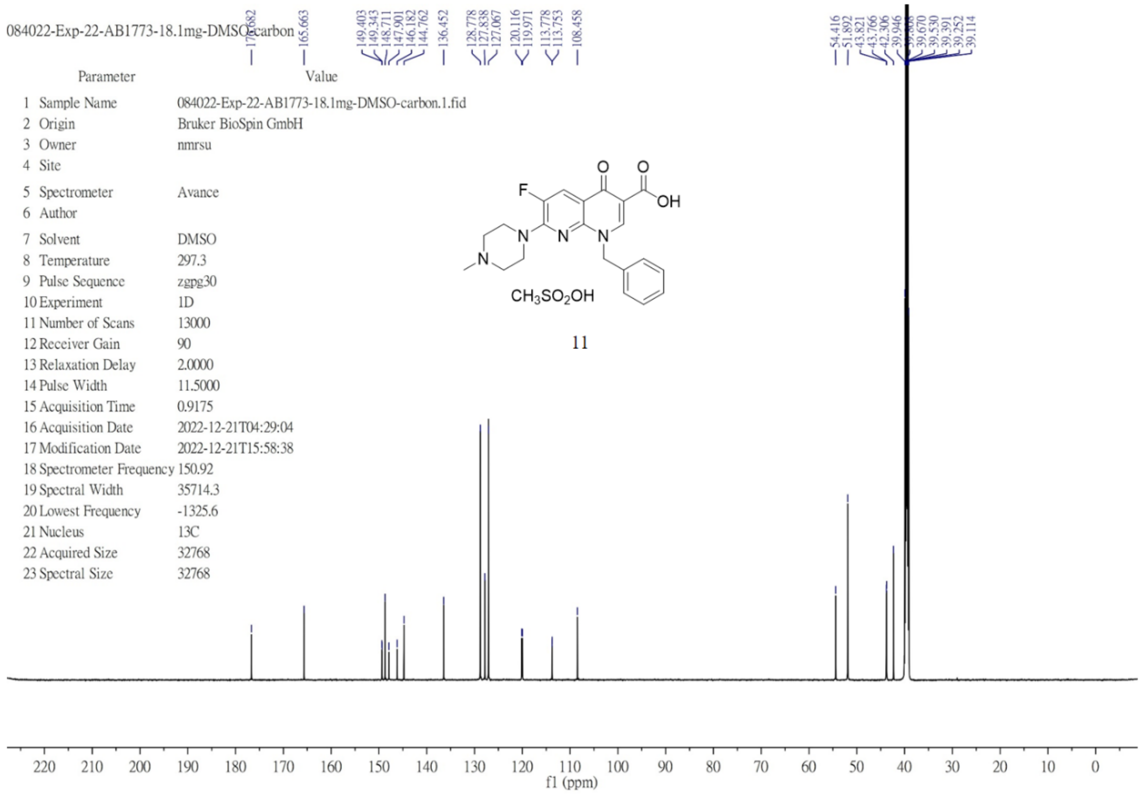

$^{1}H$ NMR (600 MHz, DMSO-$d_6$) of **12**.

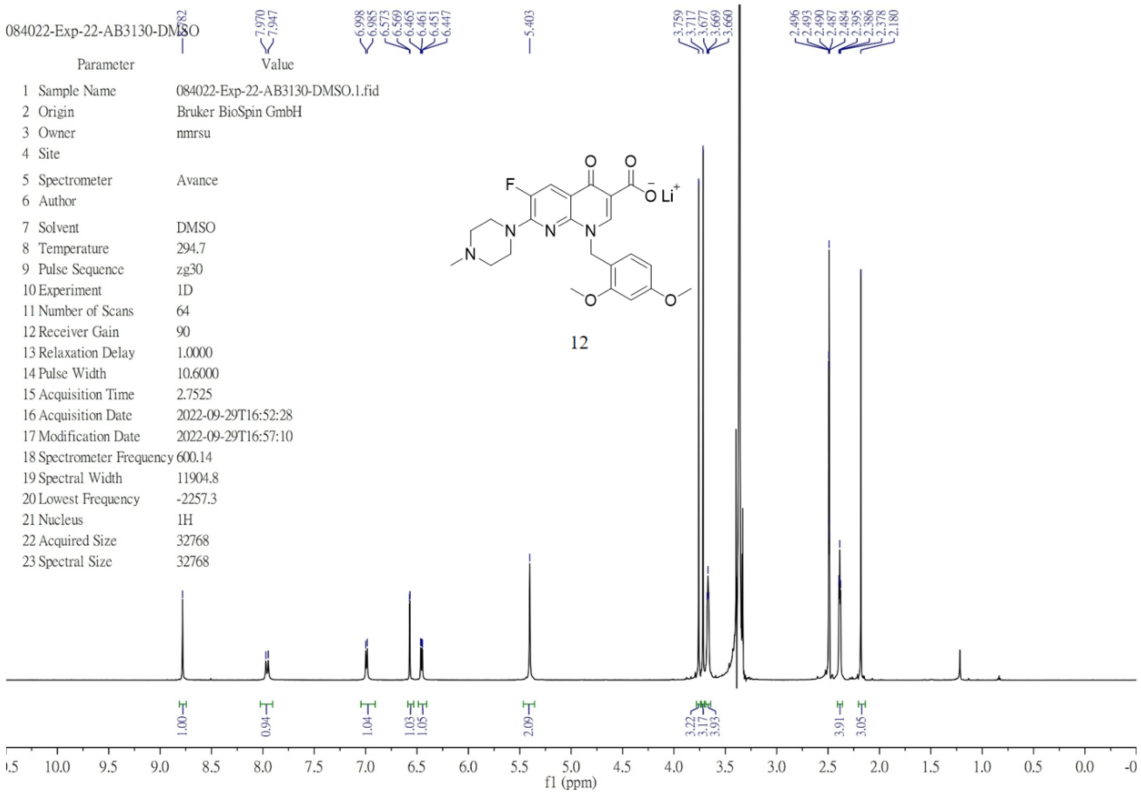


$^{13}C$ NMR (151 MHz, DMSO-$d_6$) of **12**.

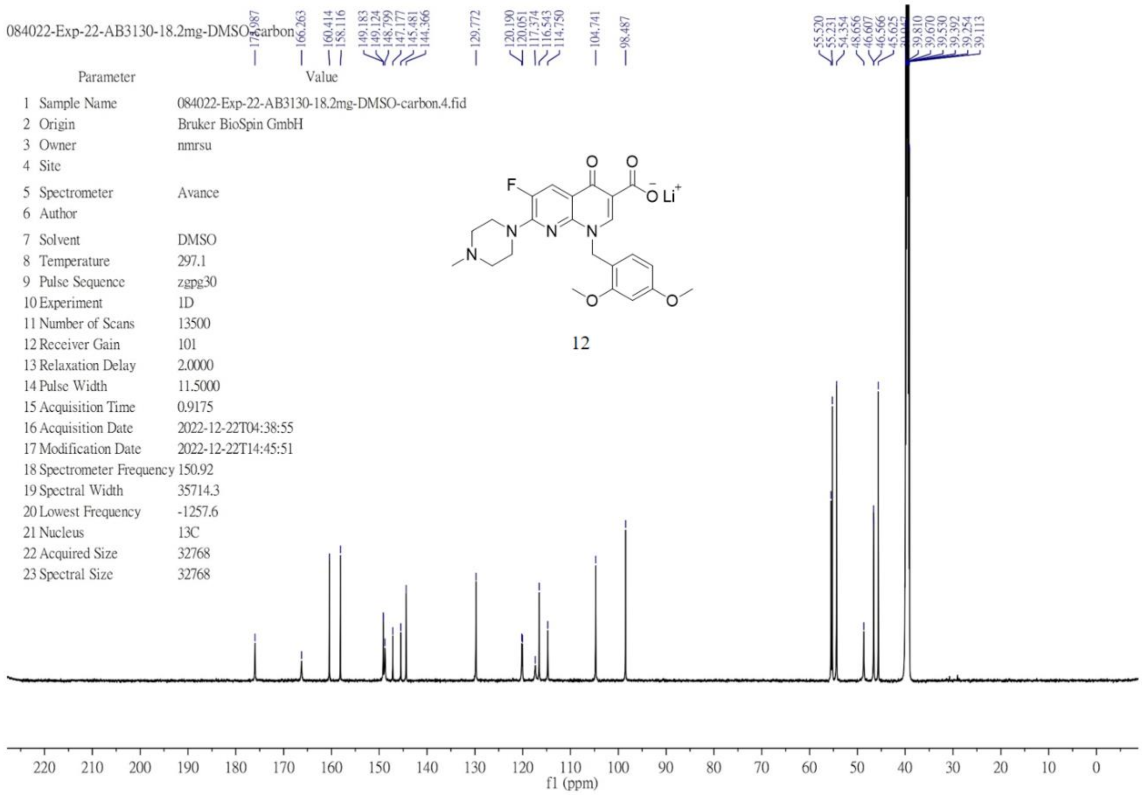

$^1H$ NMR (600 MHz, DMSO-$d_6$) of **13**.

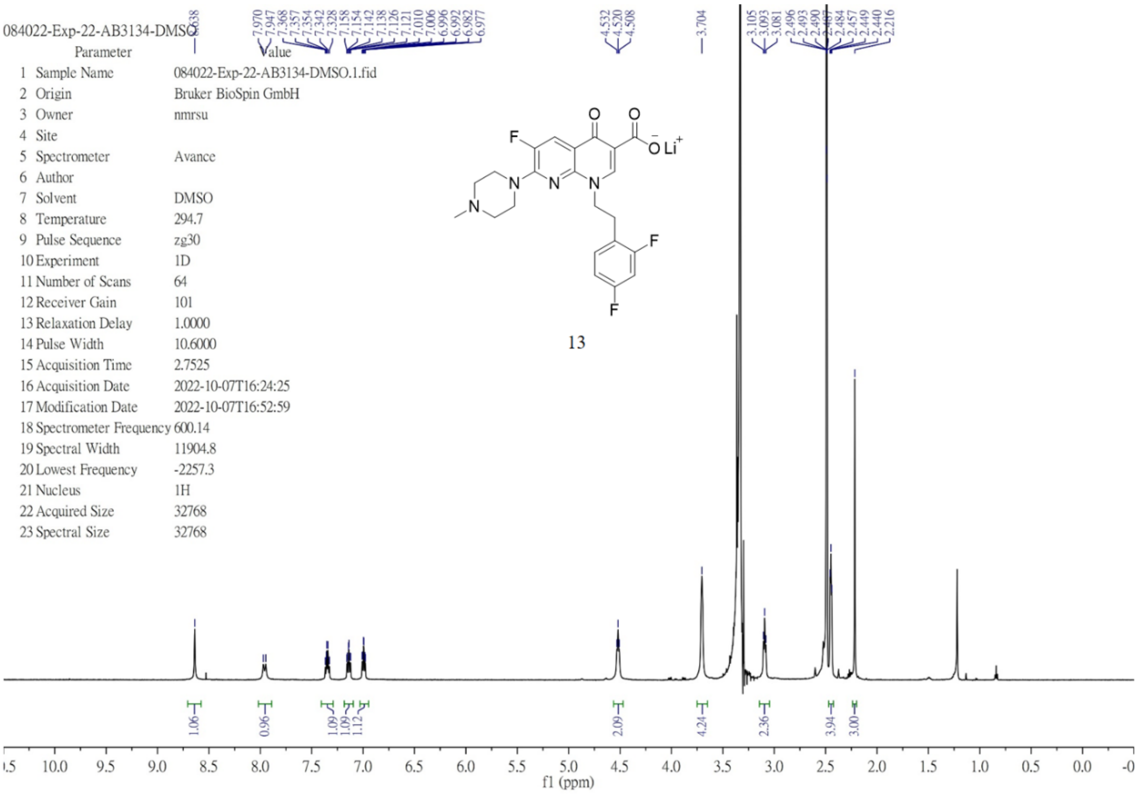


$^{13}C$ NMR (151 MHz, methanol-$d_4$) of **13**.

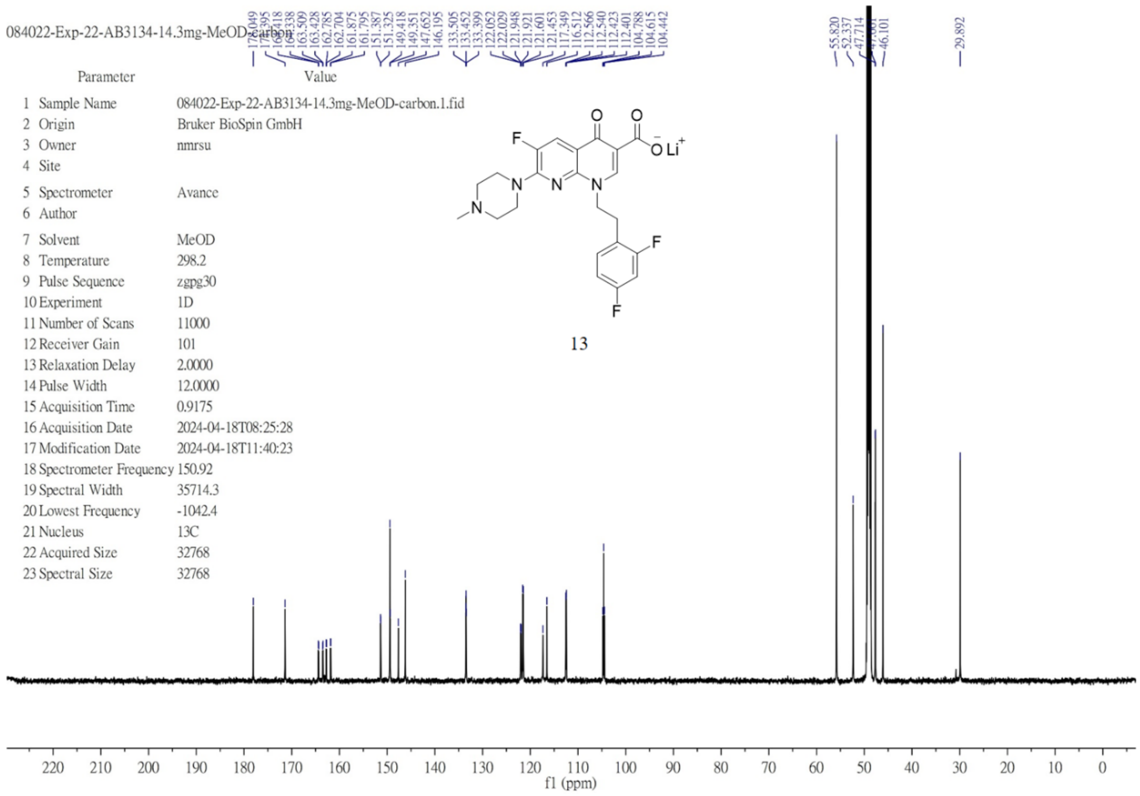

$^{1}H$ NMR (600 MHz, DMSO-$d_6$) of **14**.

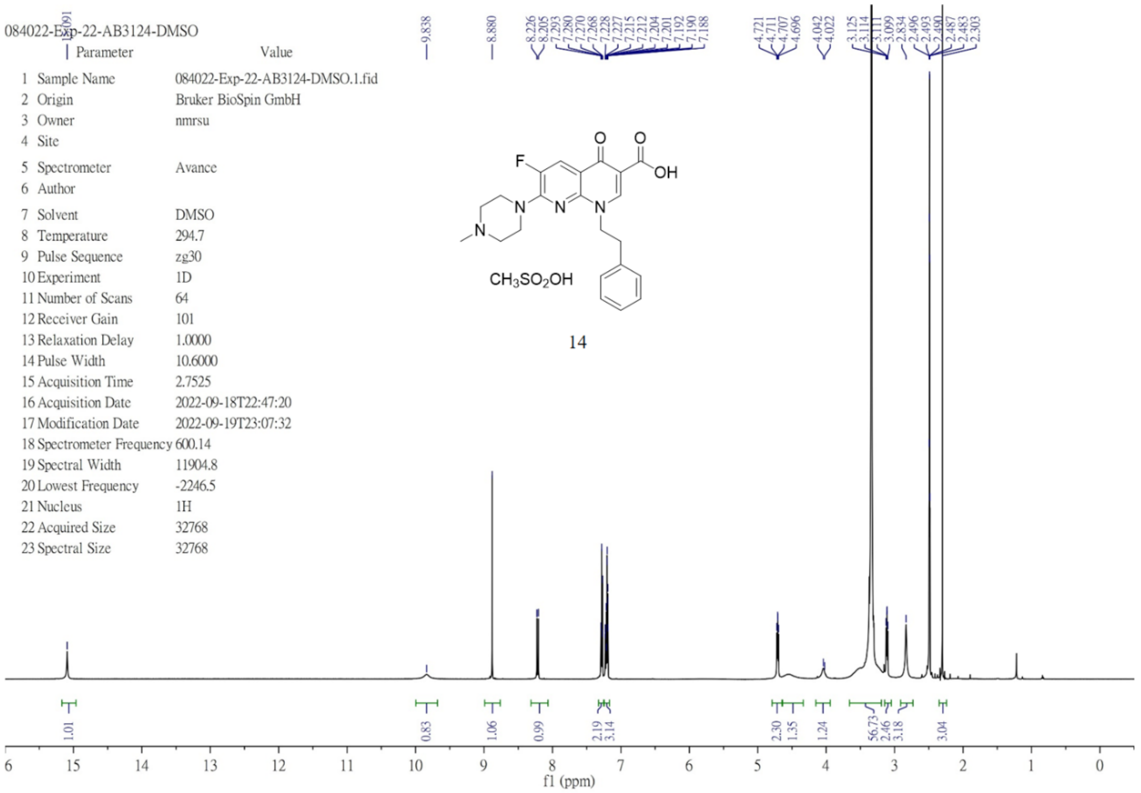

$^{13}C$ NMR (151 MHz, DMSO-$d_6$) of **14**.

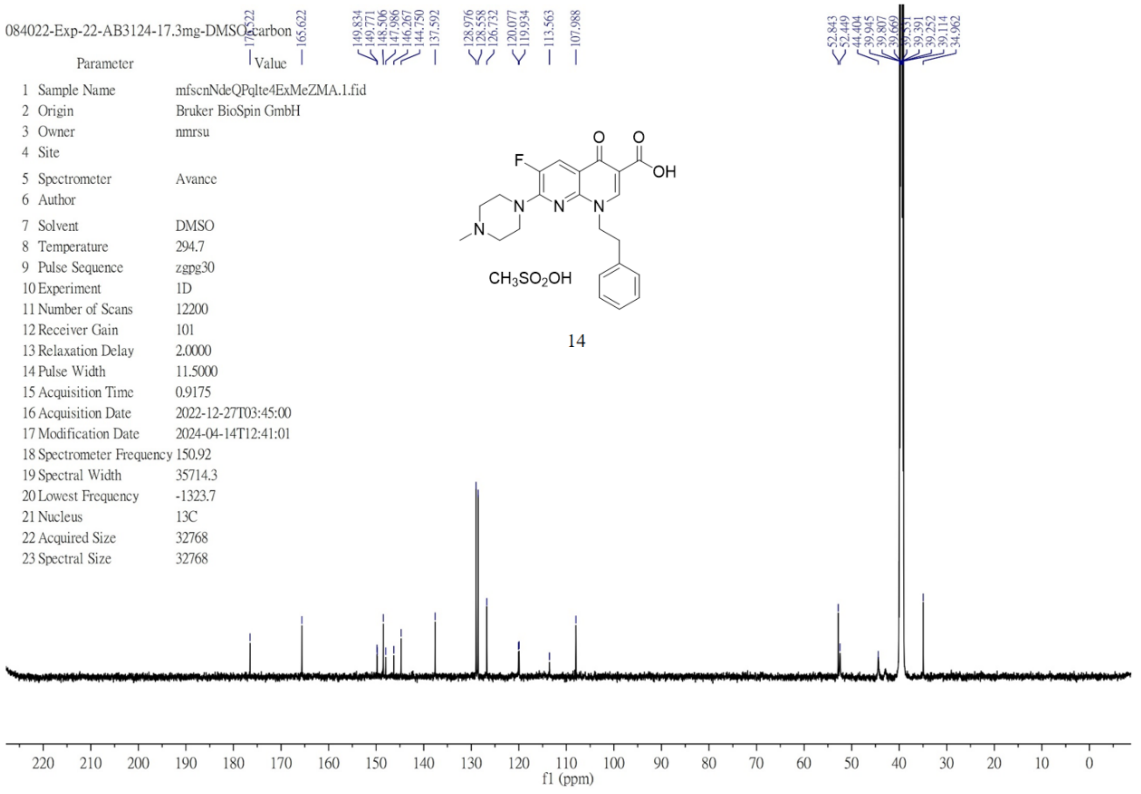

$^{1}$H NMR (600 MHz, DMSO-$d_6$) of **15**.

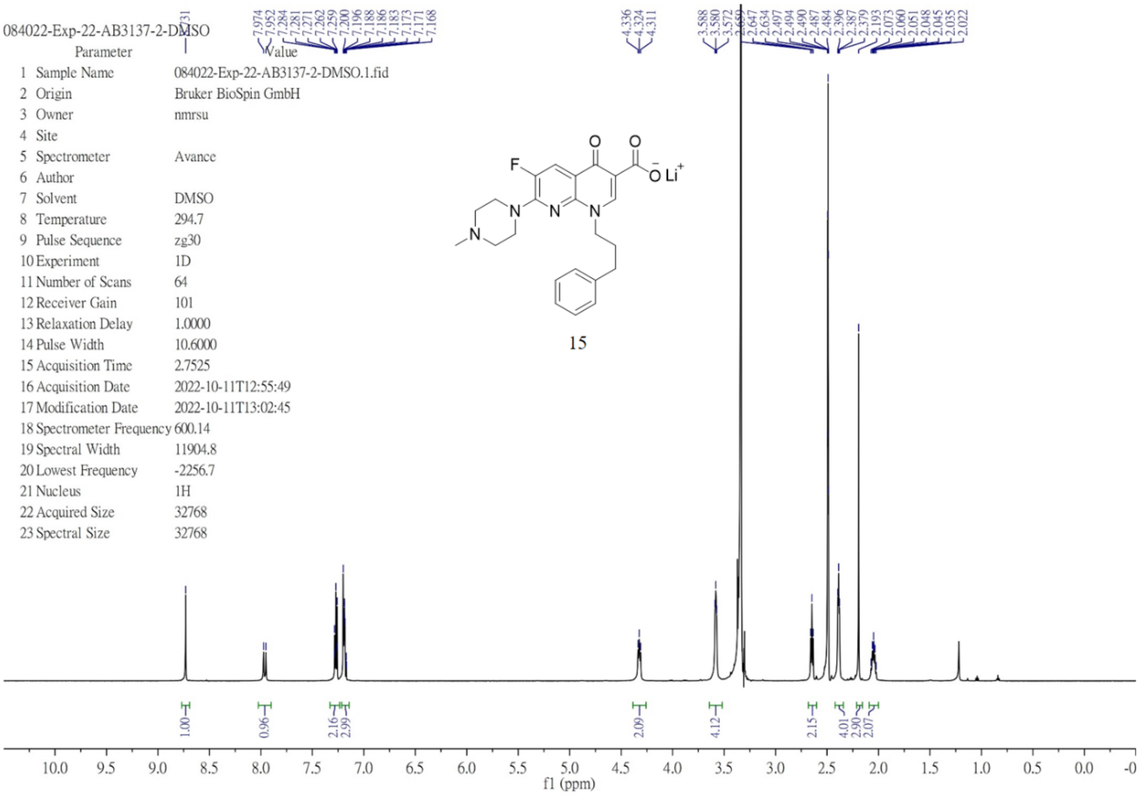


$^{13}$C NMR (151 MHz, DMSO-$d_6$) of **15**.

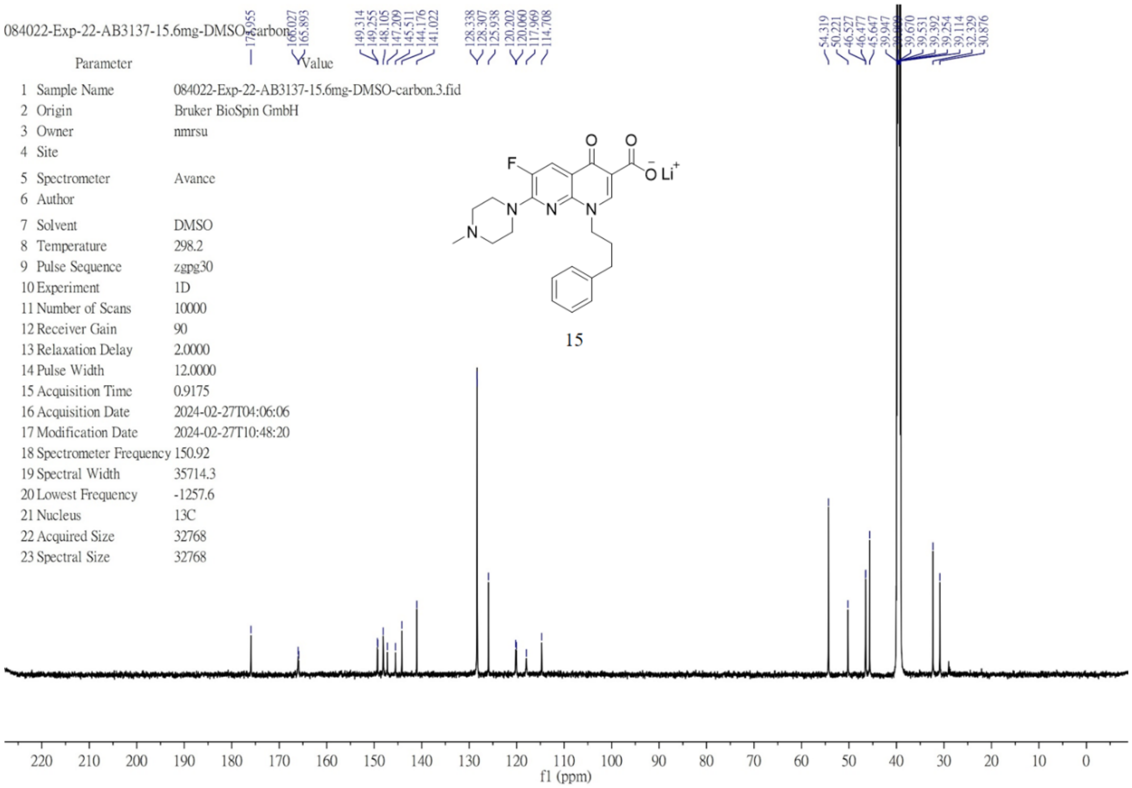

$^{1}H$ NMR (600 MHz, chloroform-*d*) of **16**.

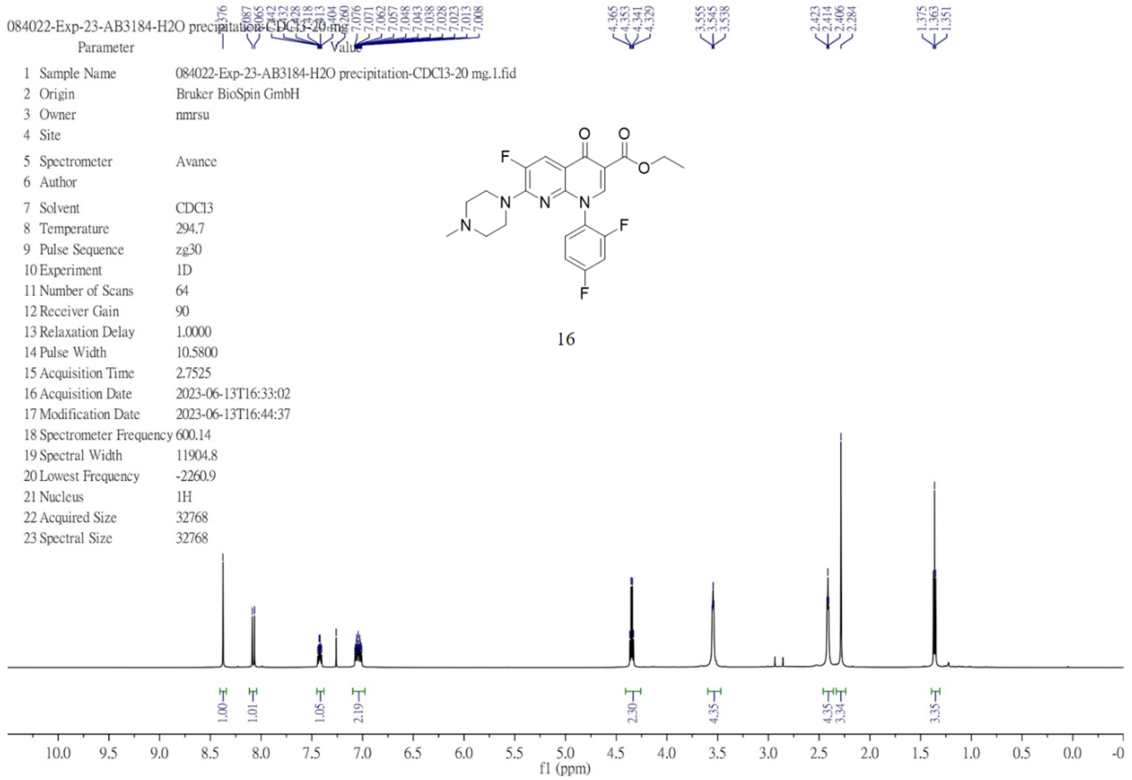

$^{13}C$ NMR (101 MHz, chloroform-*d*) of **16**.

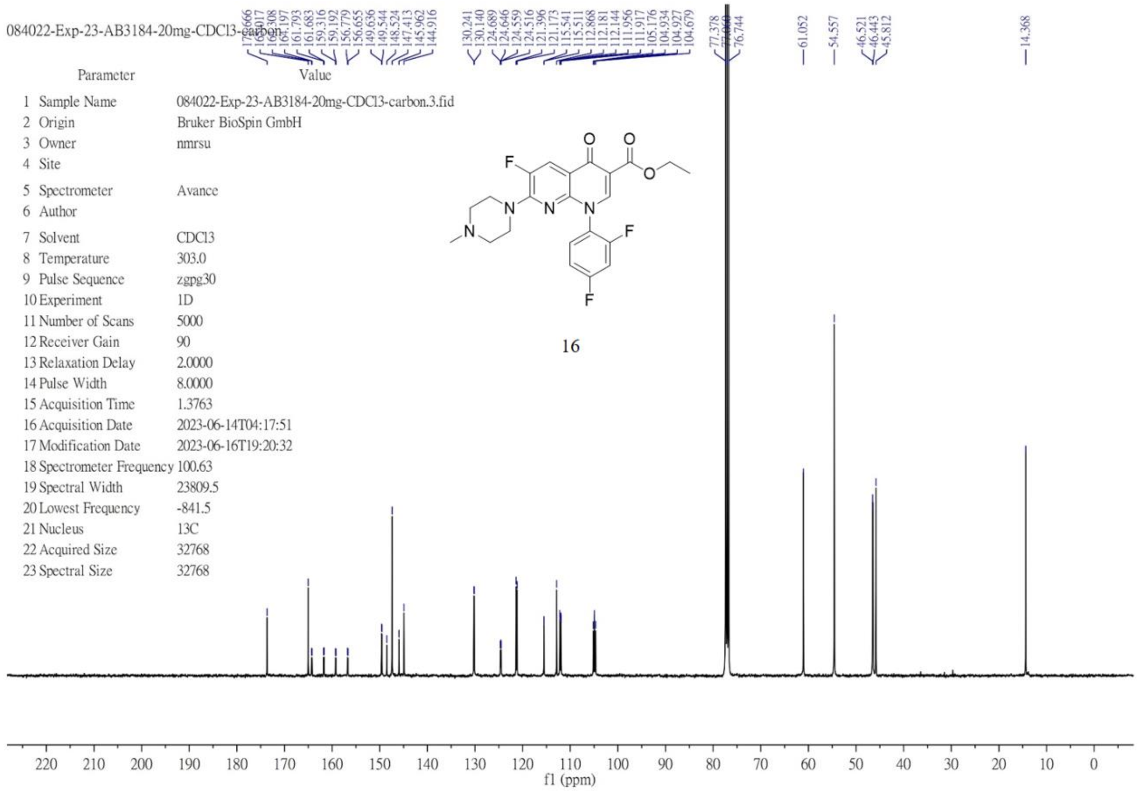

$^{1}$H NMR (400 MHz, DMSO-$d_6$) of **17**.

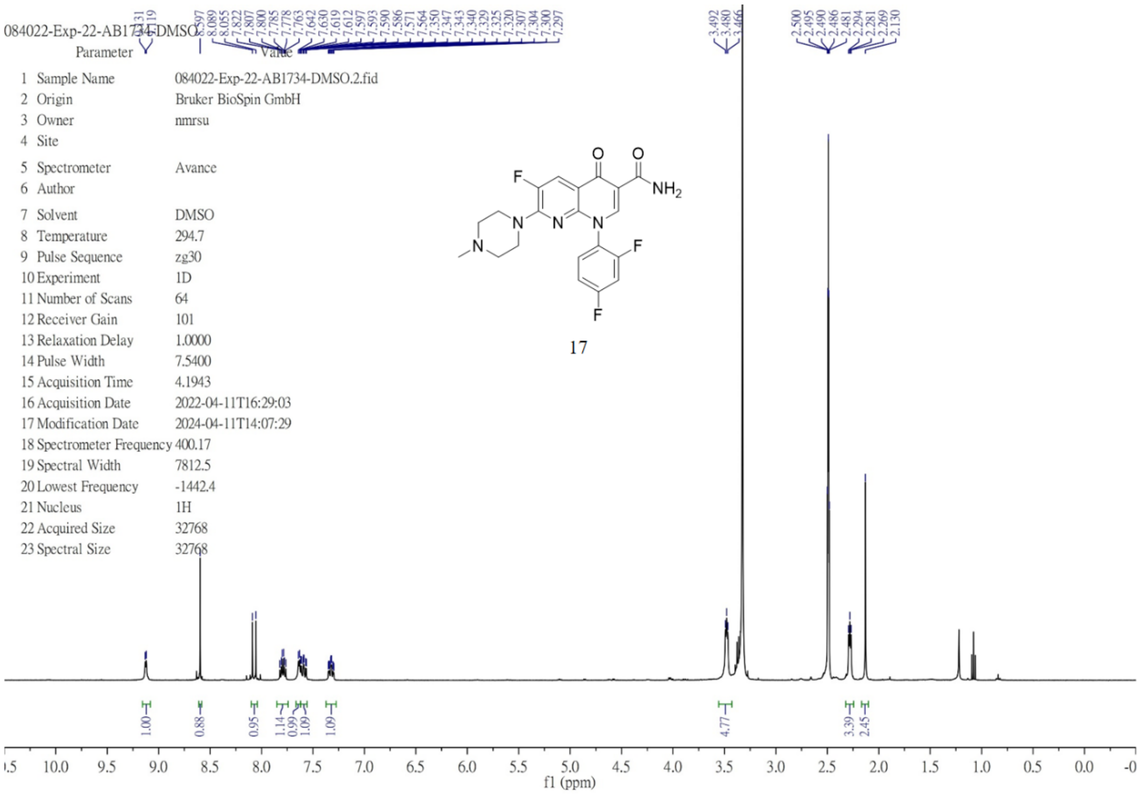


$^{13}$C NMR (151 MHz, DMSO-$d_6$) of **17**.

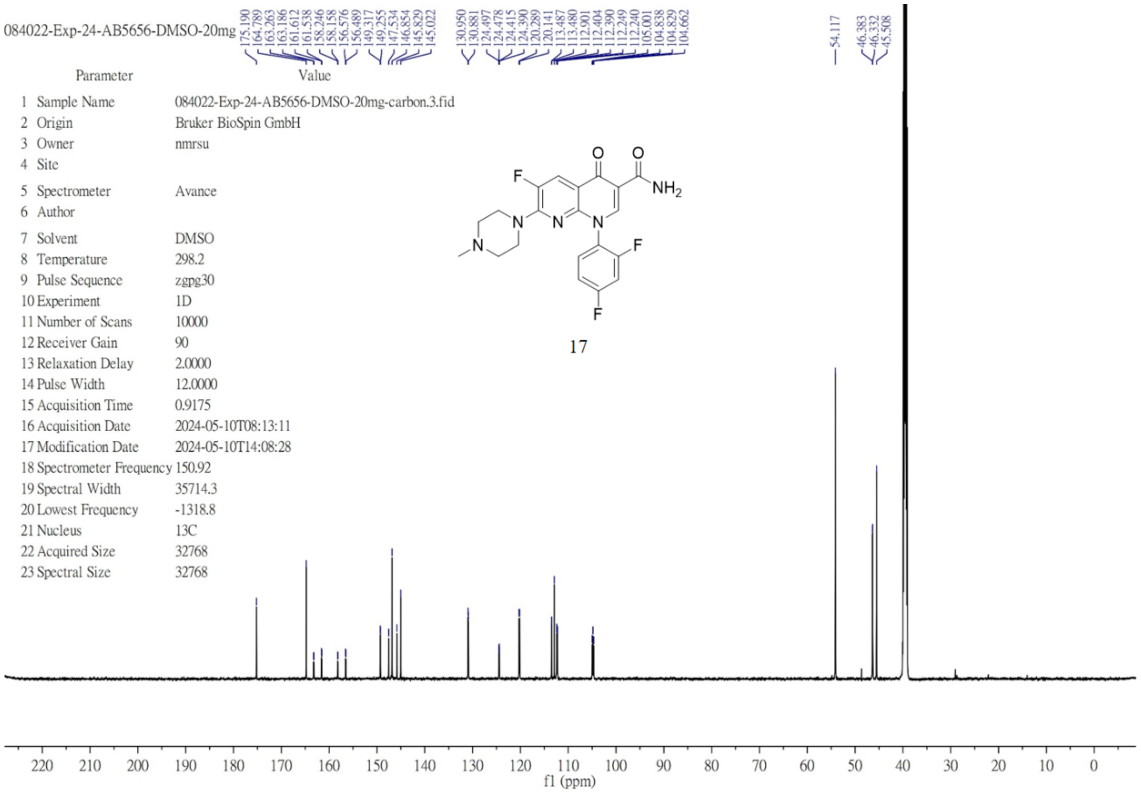

$^{1}H$ NMR (600 MHz, chloroform-*d*) of **18**.

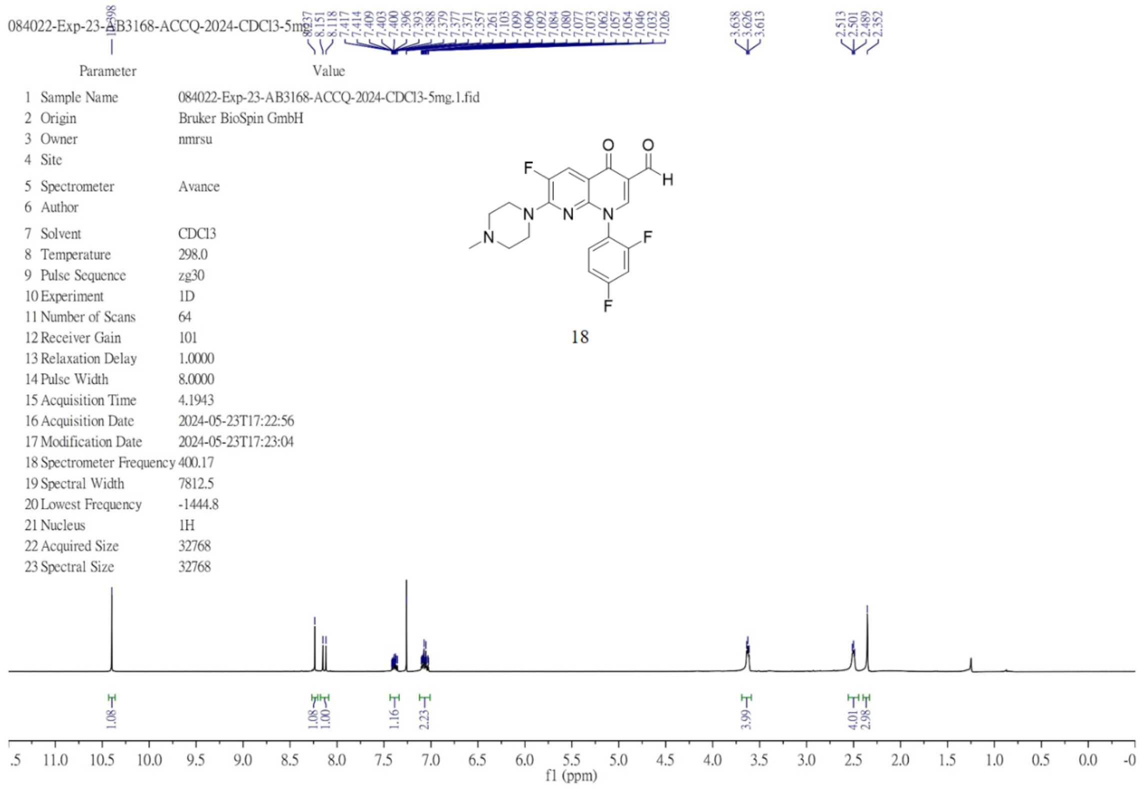

$^{13}C$ NMR (101 MHz, chloroform-*d*) of **18**.

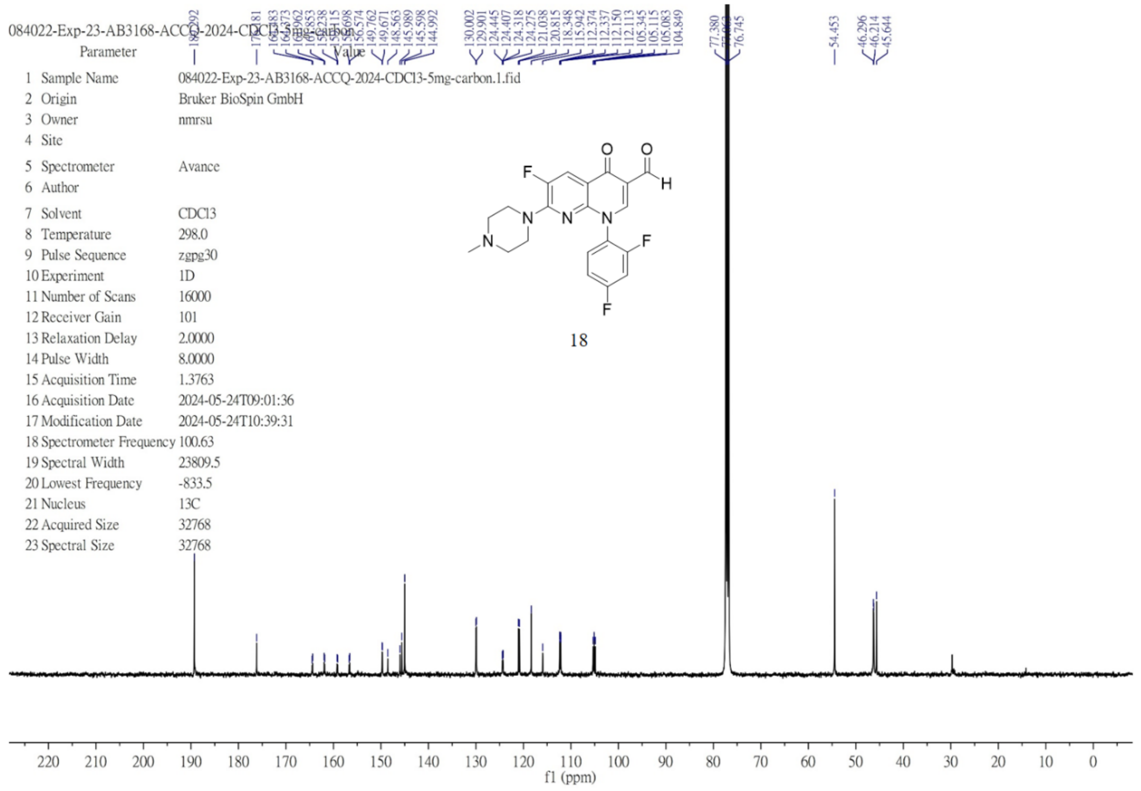

$^{1}$H NMR (600 MHz, DMSO-$d_6$) of **19**.

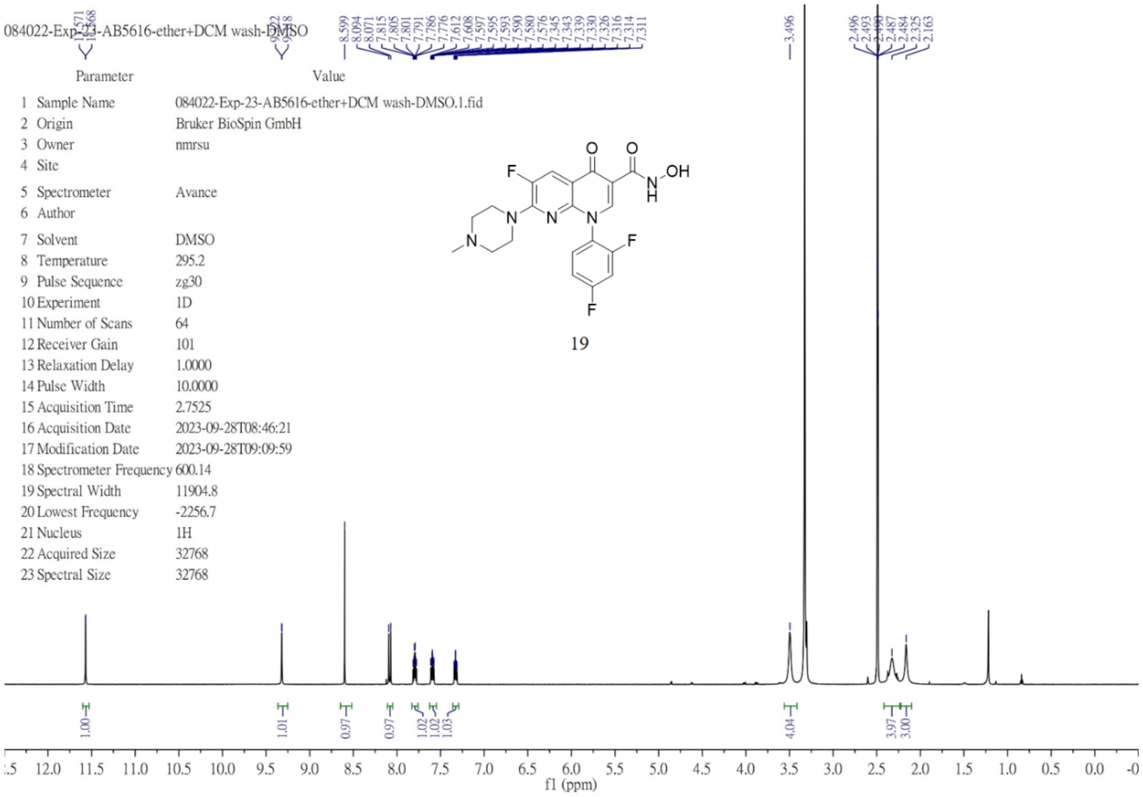


$^{13}$C NMR (151 MHz, DMSO-$d_6$) of **19**.

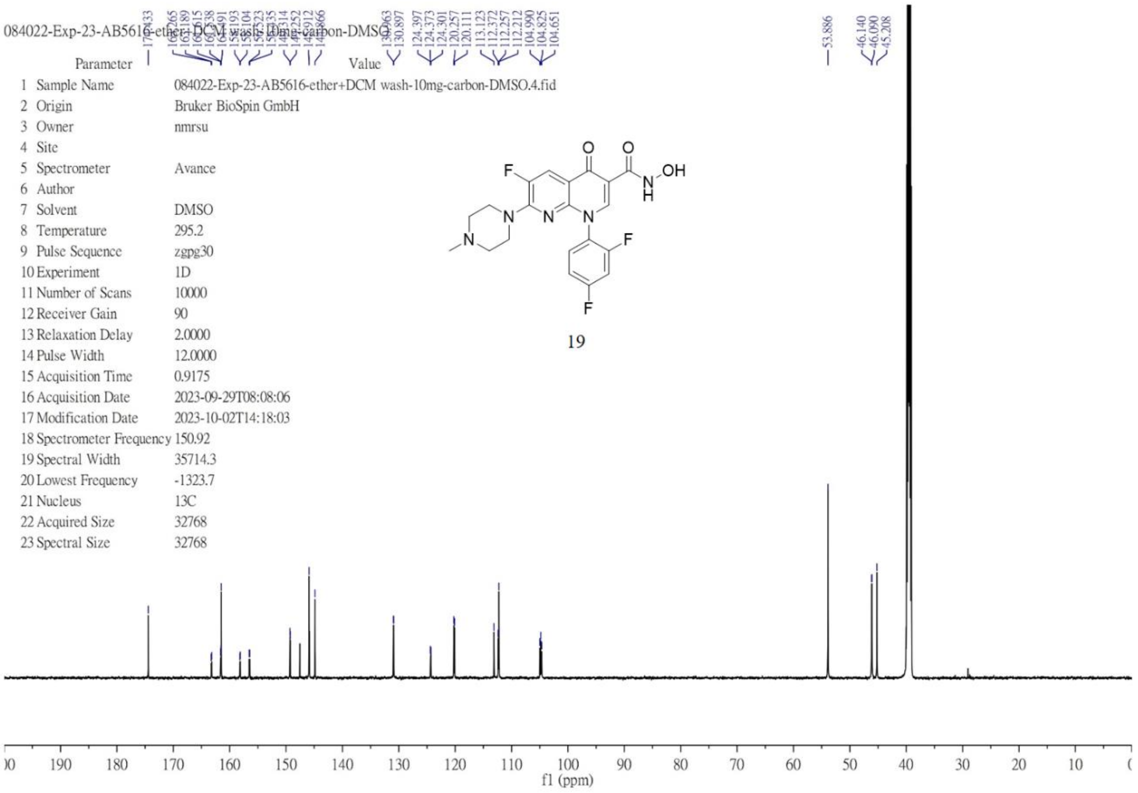

$^{1}$H NMR (400 MHz, DMSO-$d_6$) of **20**.

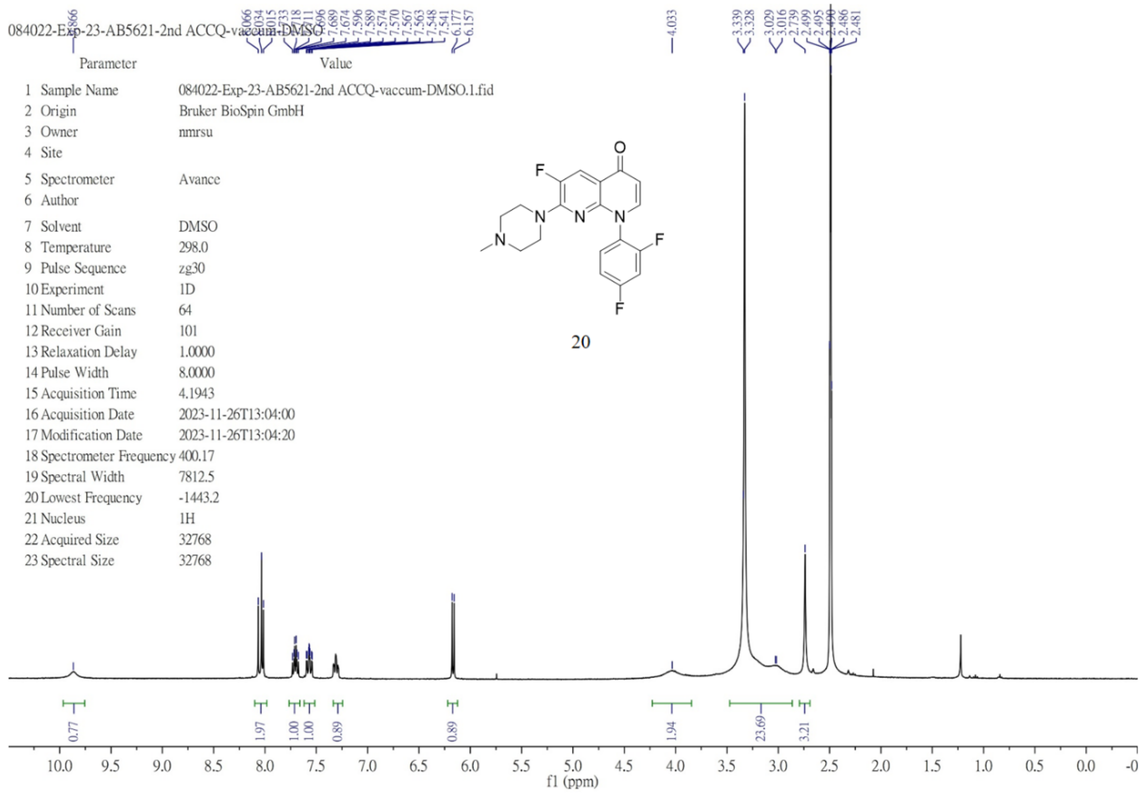


$^{13}$C NMR (151 MHz, DMSO-$d_6$) of **20**.

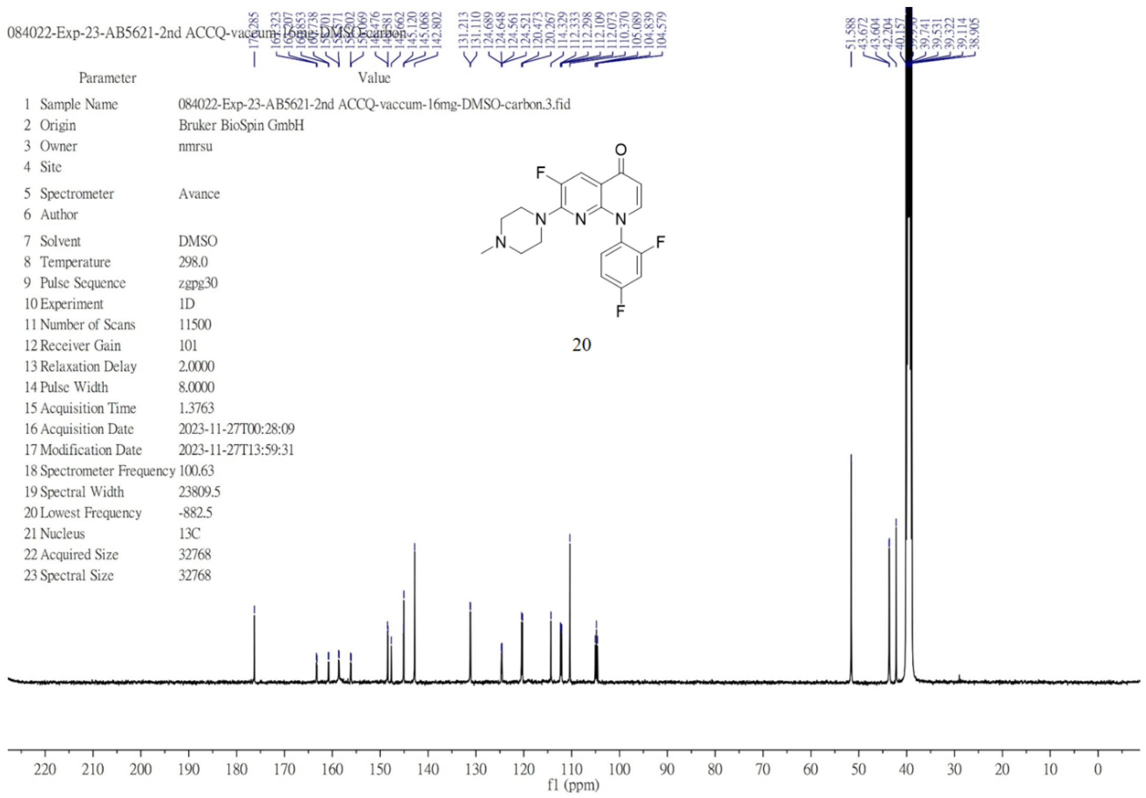

$^{1}H$ NMR (600 MHz, DMSO-$d_6$) of **21**.

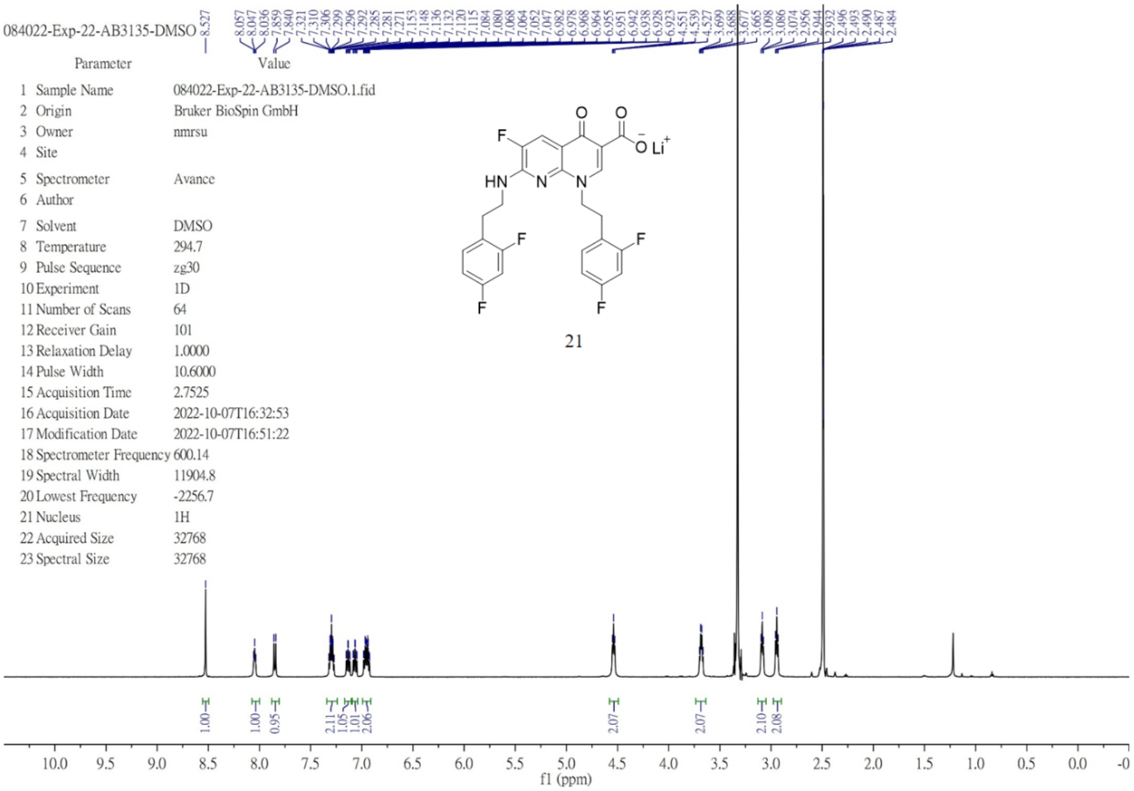


$^{13}C$ NMR (151 MHz, DMSO-$d_6$) of **21**.

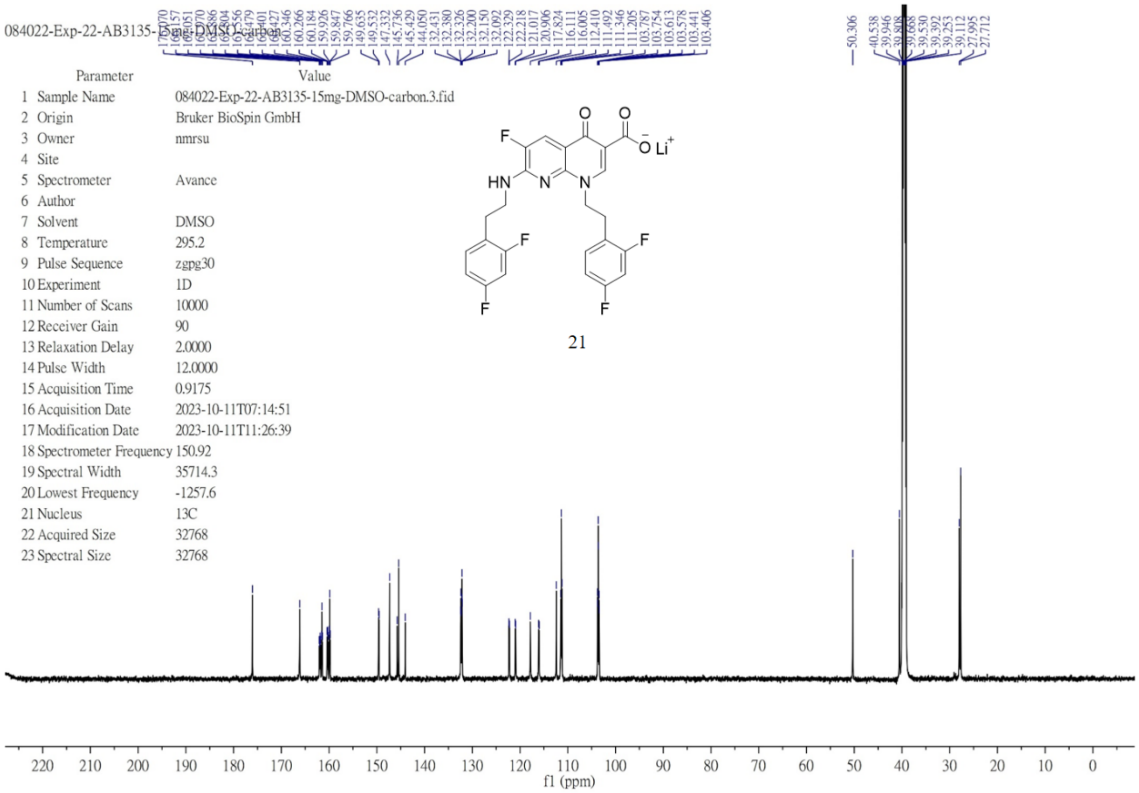

$^{1}$H NMR (600 MHz, DMSO-$d_6$) of **22**.

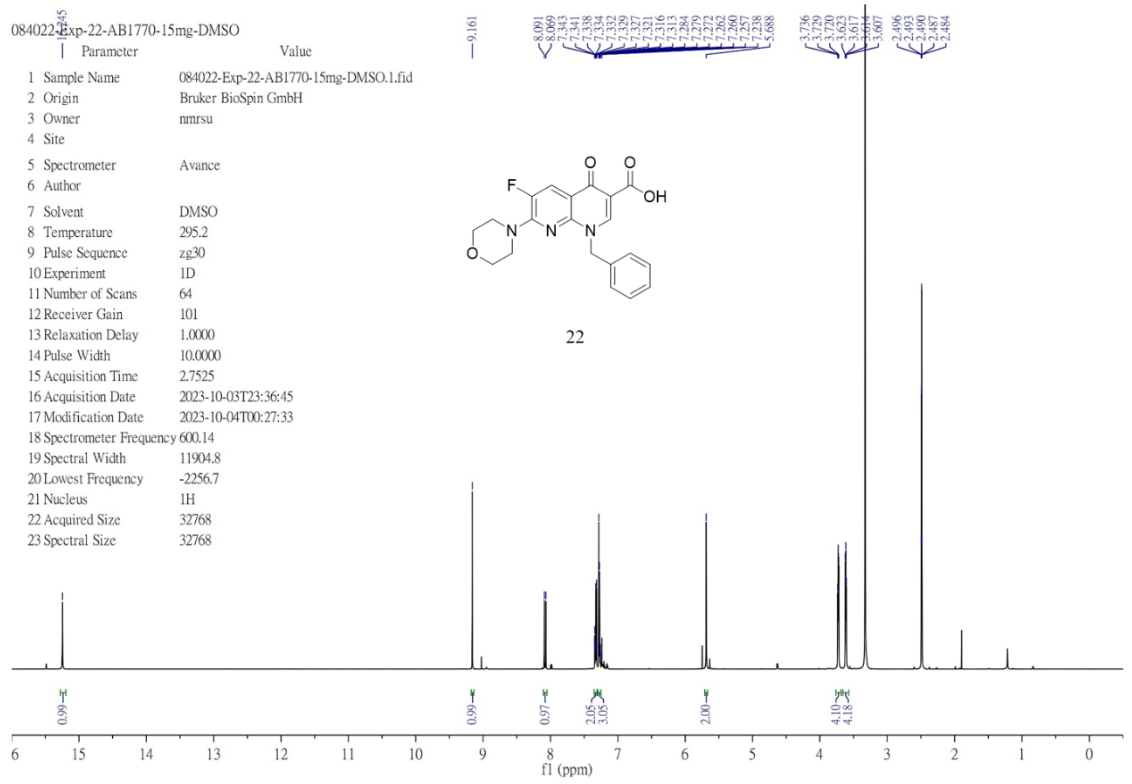

$^{13}$C NMR (151 MHz, DMSO-$d_6$) of **22**.

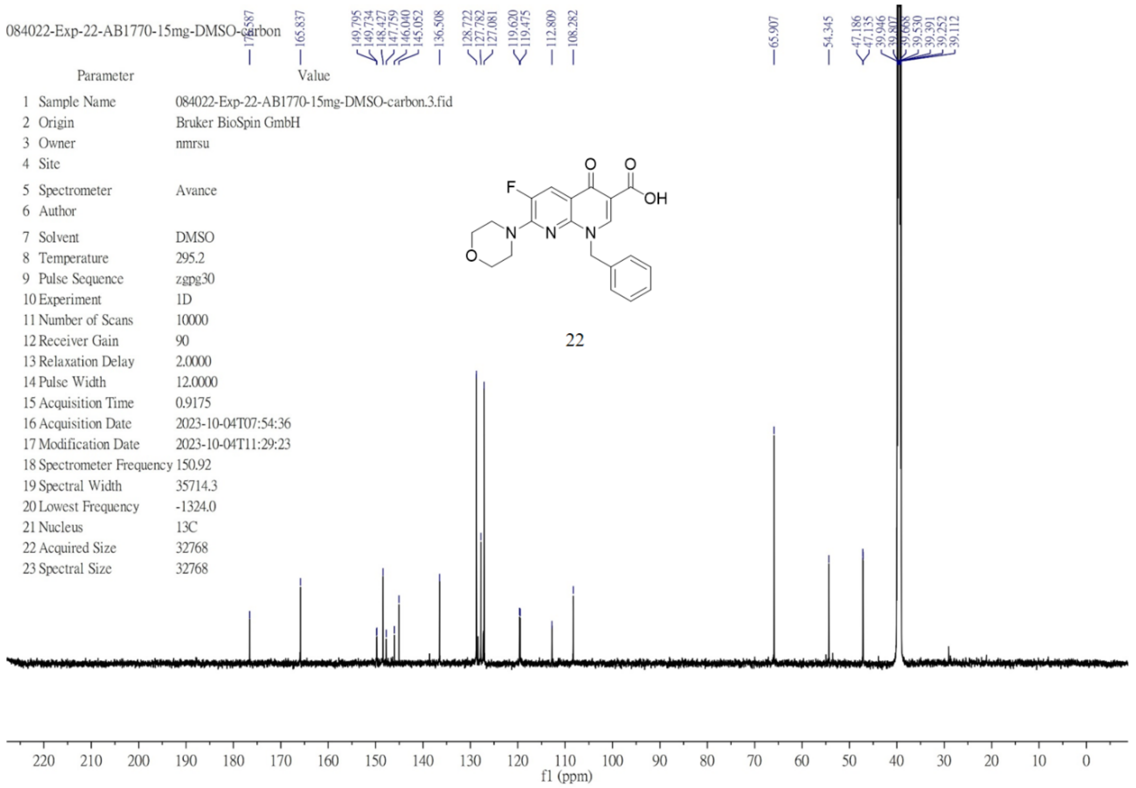

$^{1}H$ NMR (600 MHz, DMSO-$d_6$) of **23**.

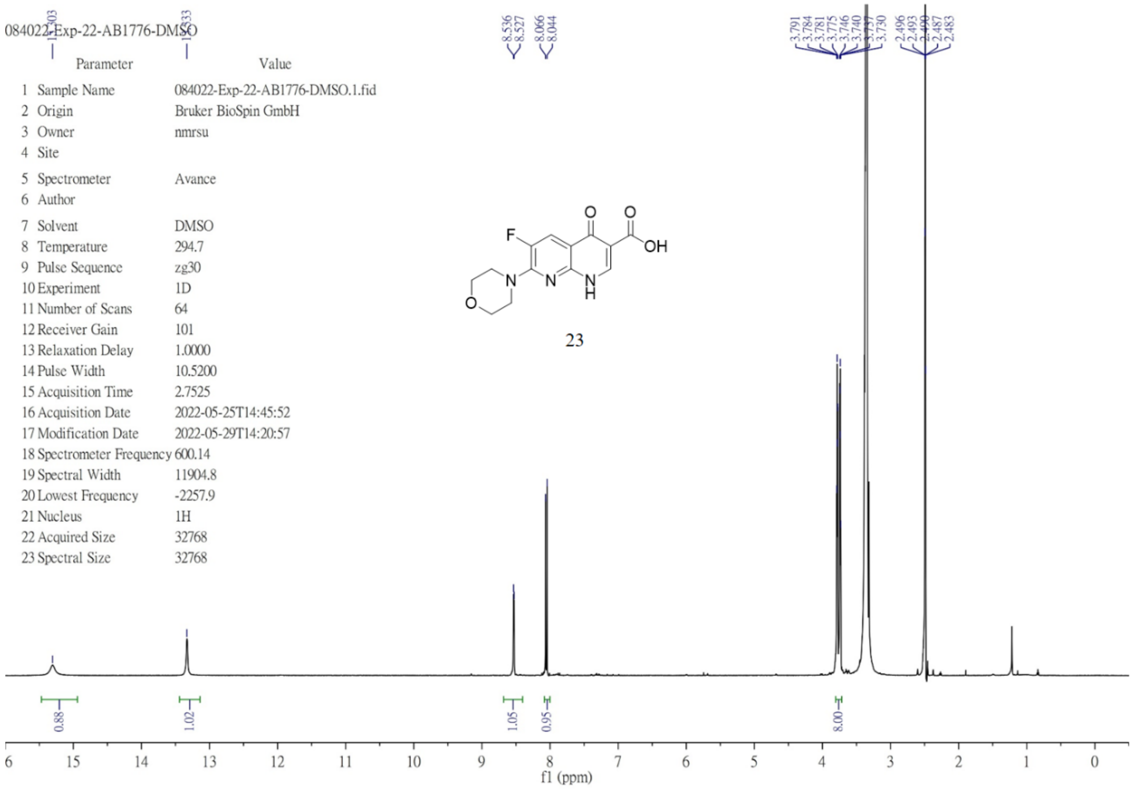


$^{13}C$ NMR (151 MHz, DMSO-$d_6$) of **23**.

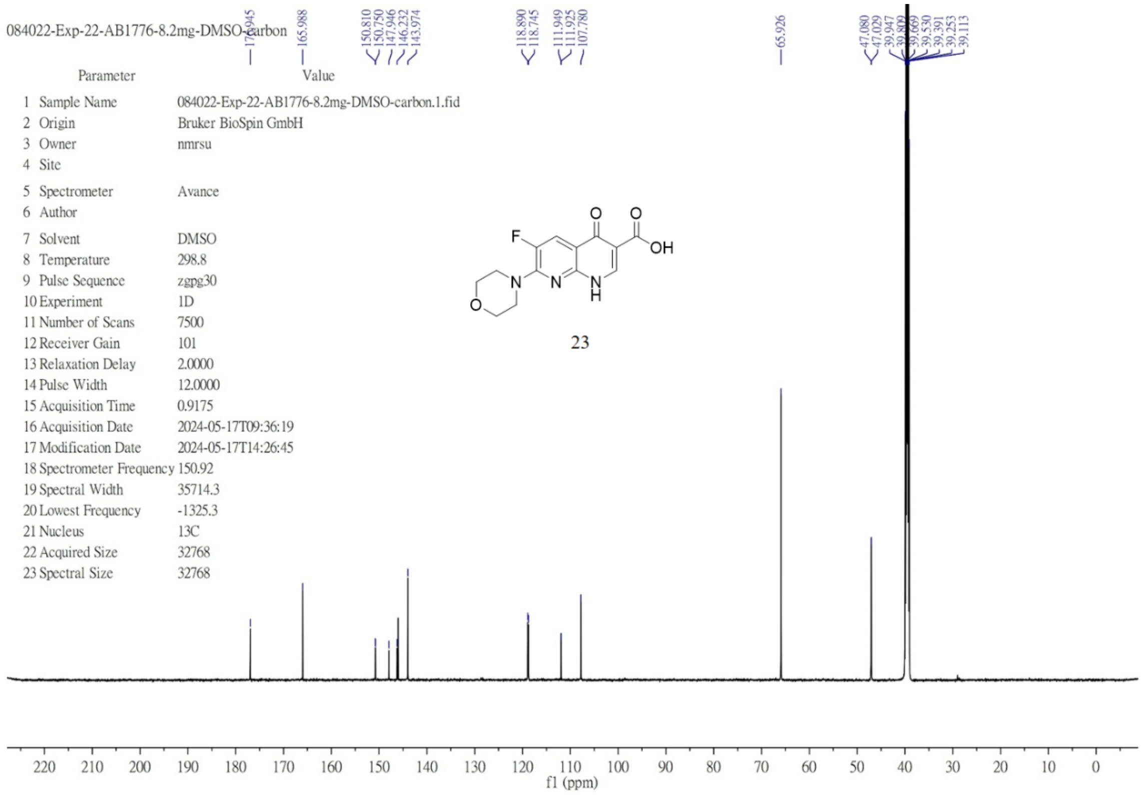

$^{1}H$ NMR (600 MHz, DMSO-$d_6$) of **24**.

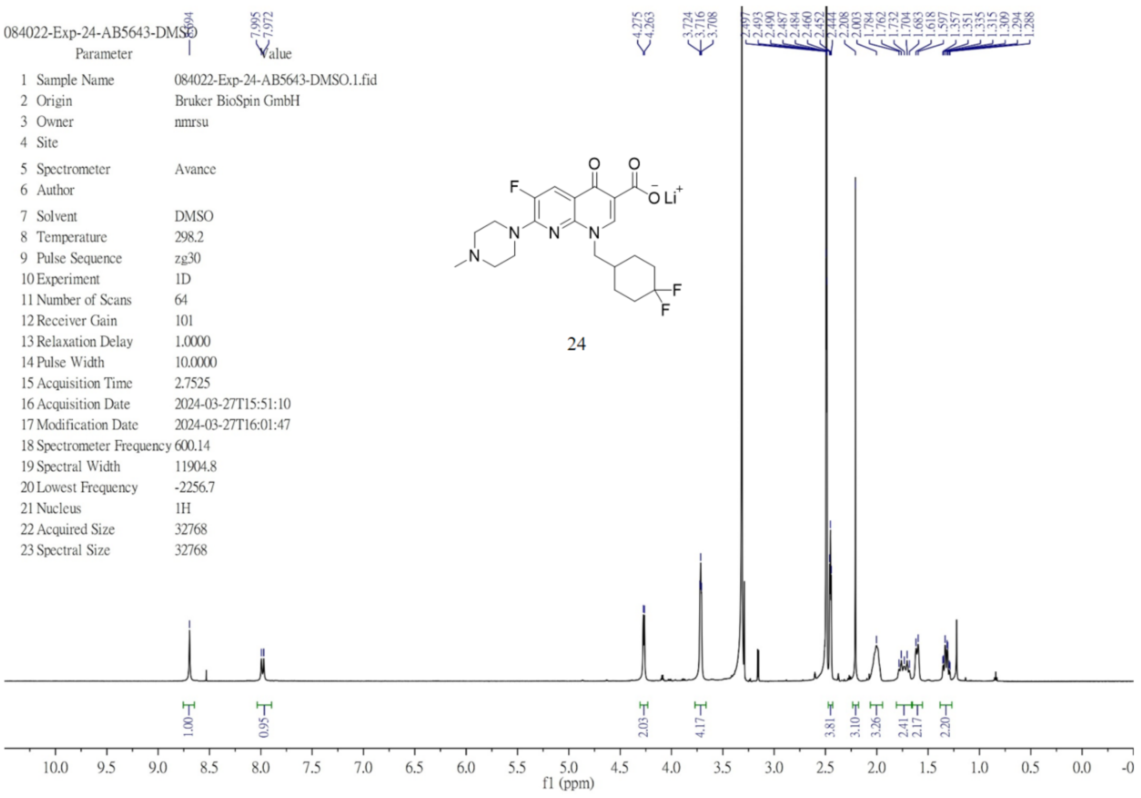

$^{13}C$ NMR (151 MHz, DMSO-$d_6$) of **24**.

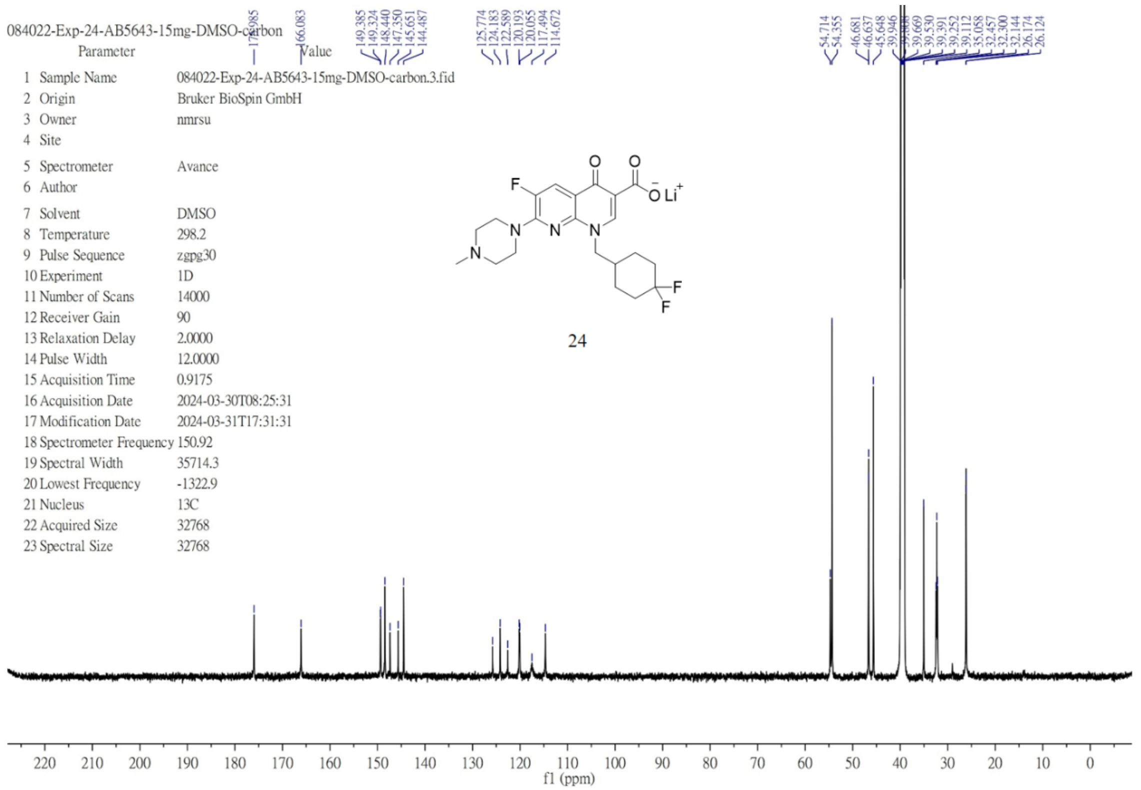

$^{1}$H NMR (400 MHz, chloroform-*d*) of **29**.

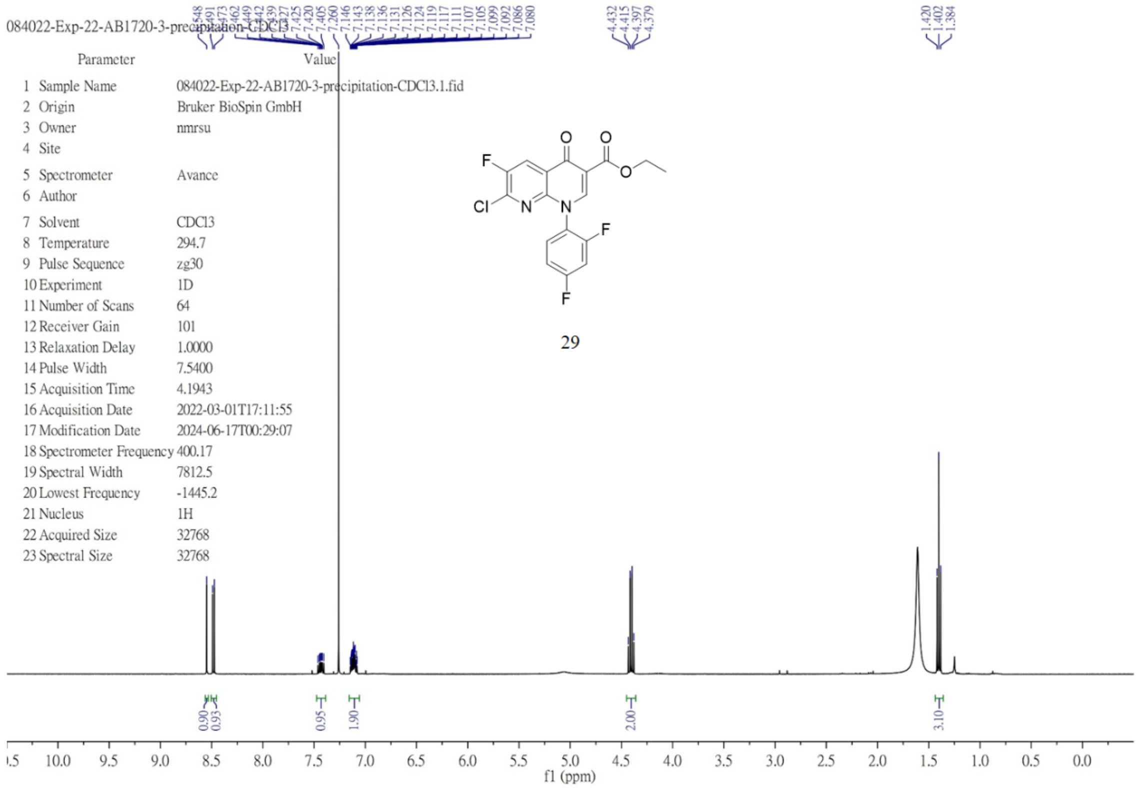

$^{1}$H NMR (600 MHz, chloroform-*d*) of **29’**.

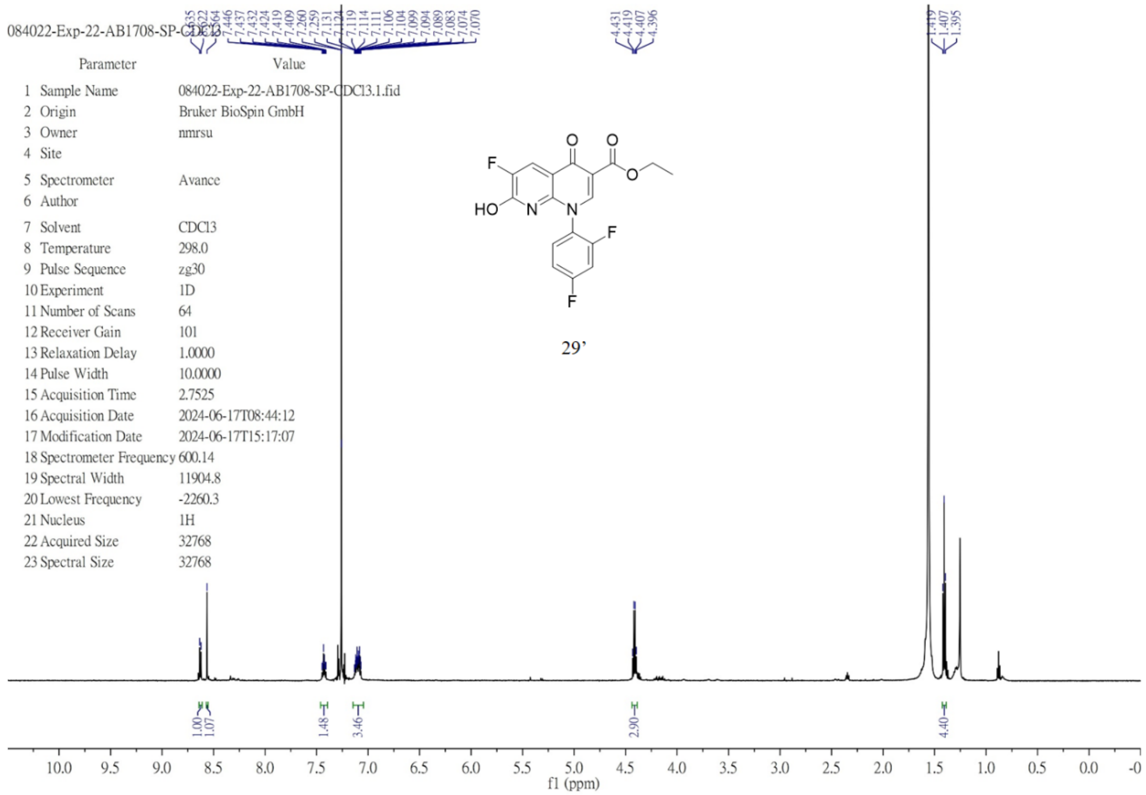

[1]H NMR (600 MHz, chloroform-*d*) of **29a**.

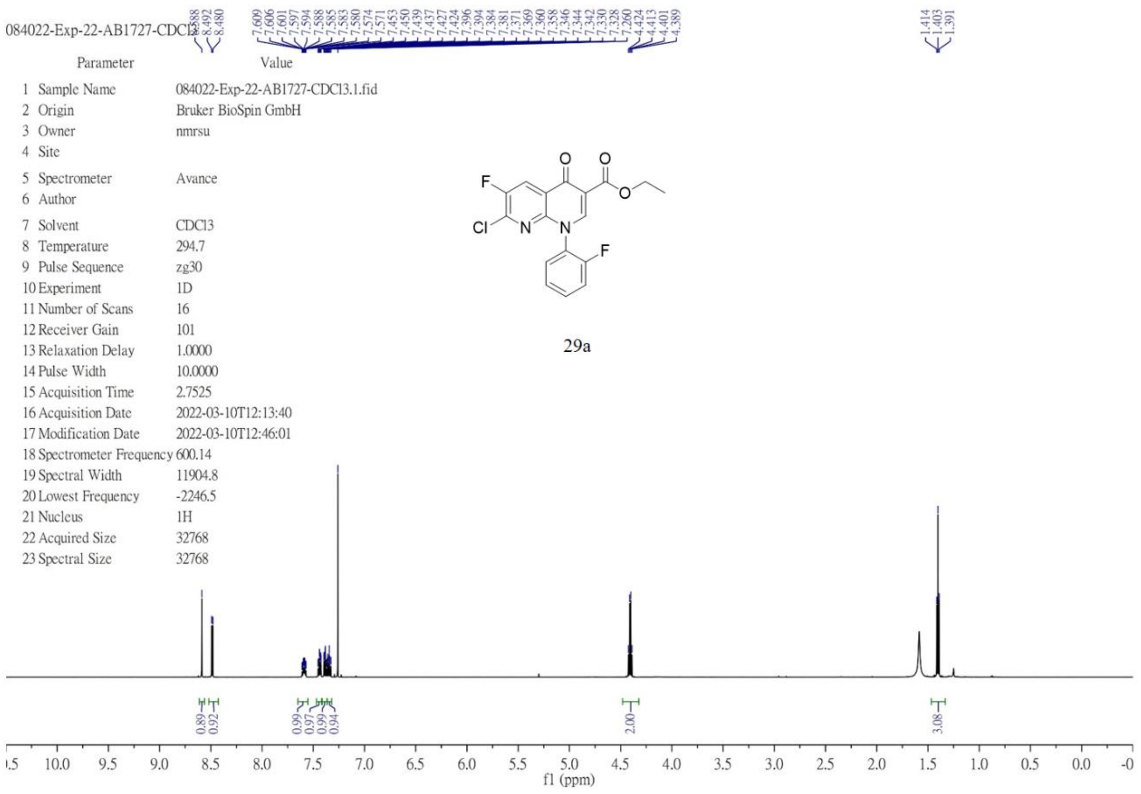


[1]H NMR (600 MHz, chloroform-*d*) of **29b**.

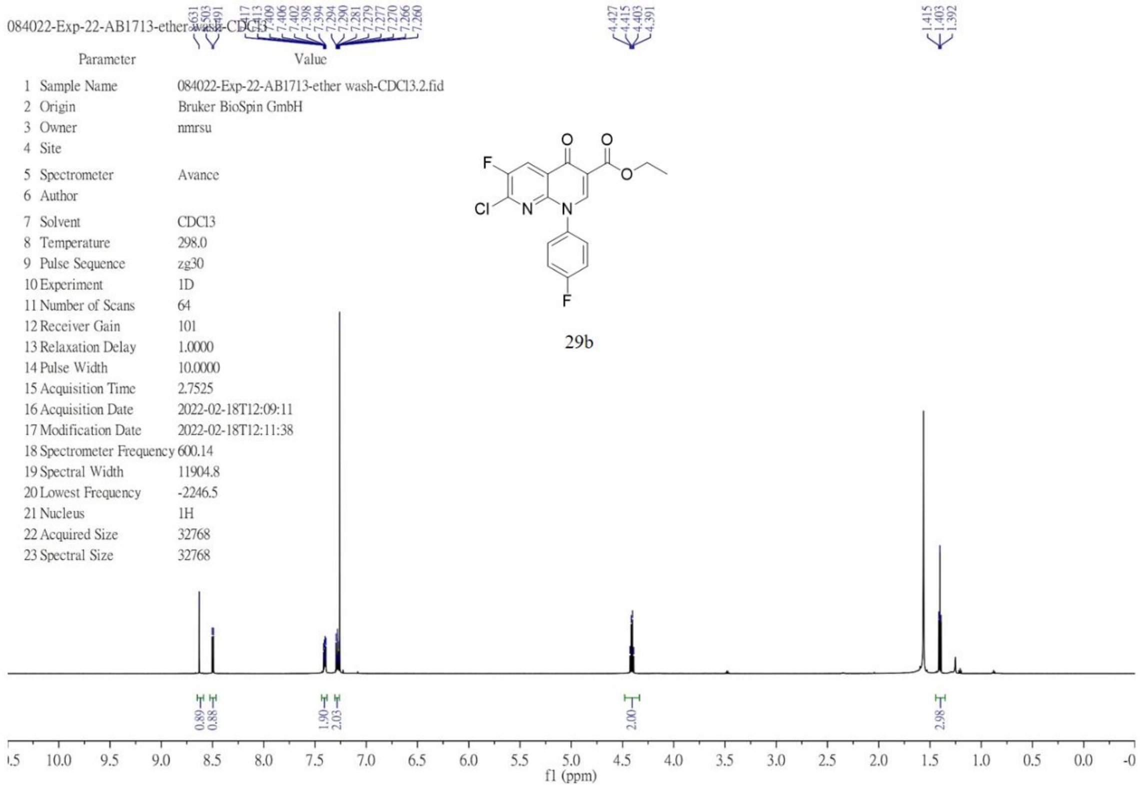

$^{1}$H NMR (600 MHz, chloroform-*d*) of **29c**.

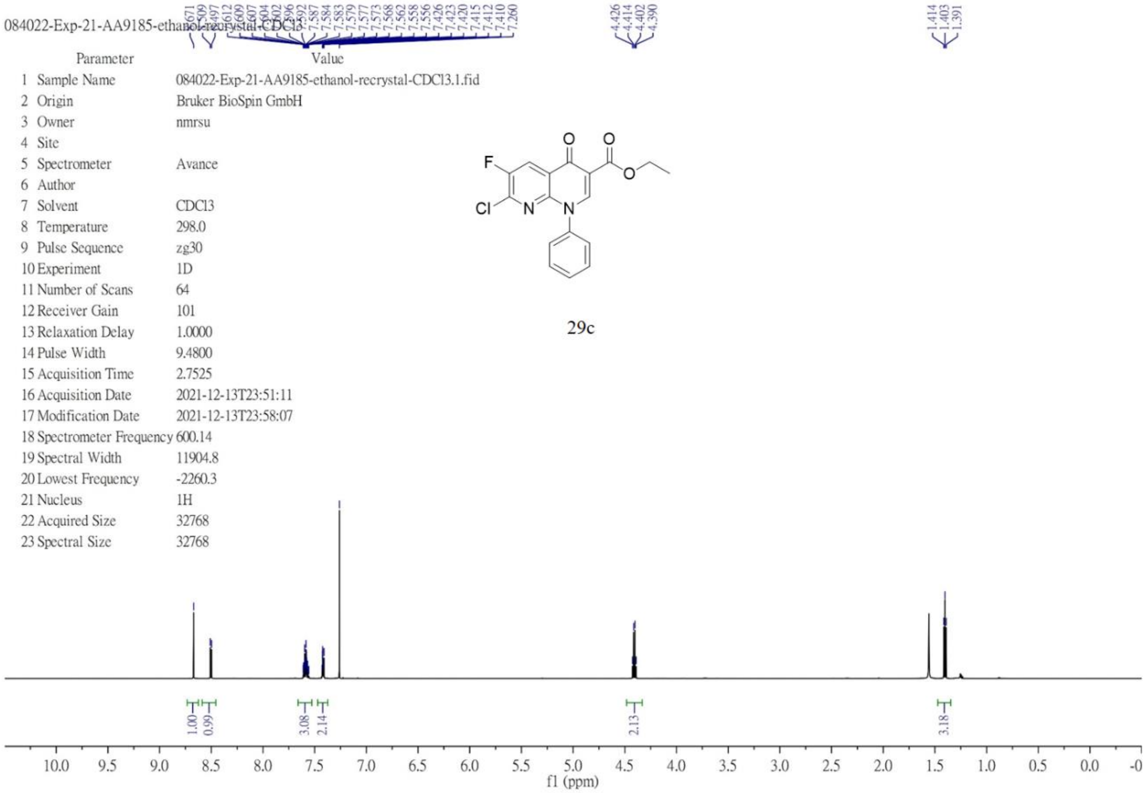


$^{1}$H NMR (600 MHz, chloroform-*d*) of **29d**.

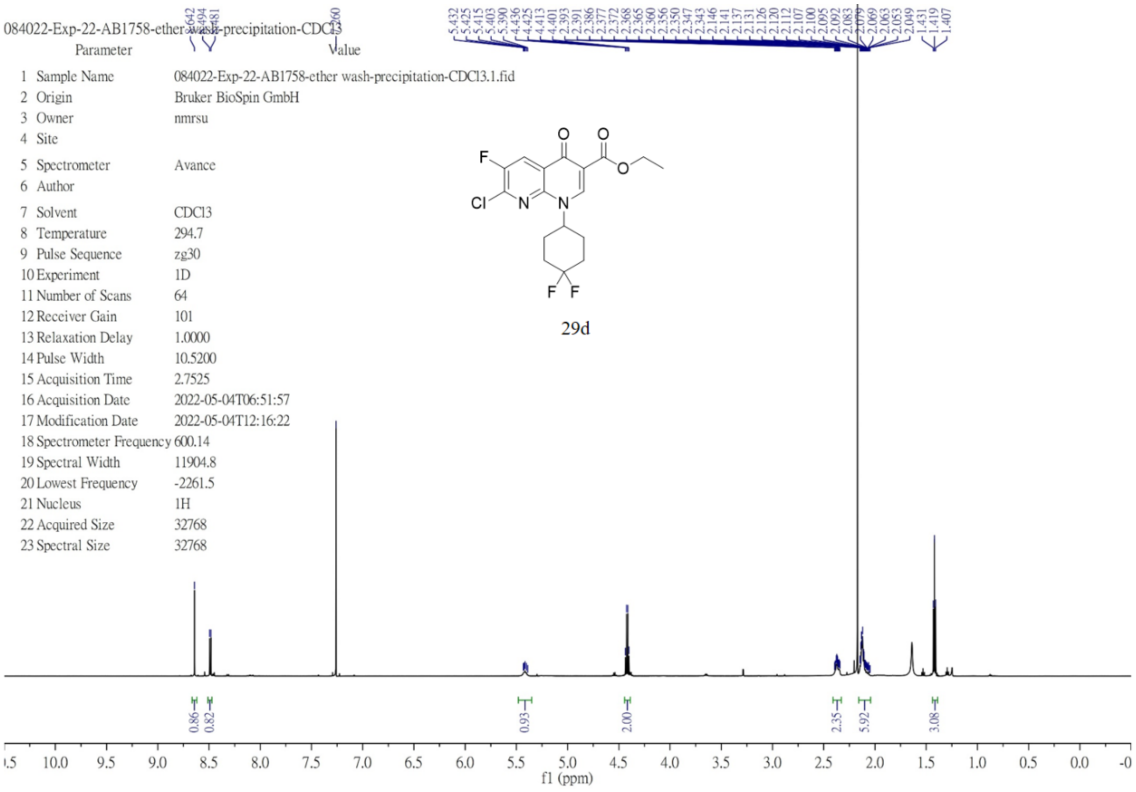

[1]H NMR (400 MHz, chloroform-*d*) of **29e**.

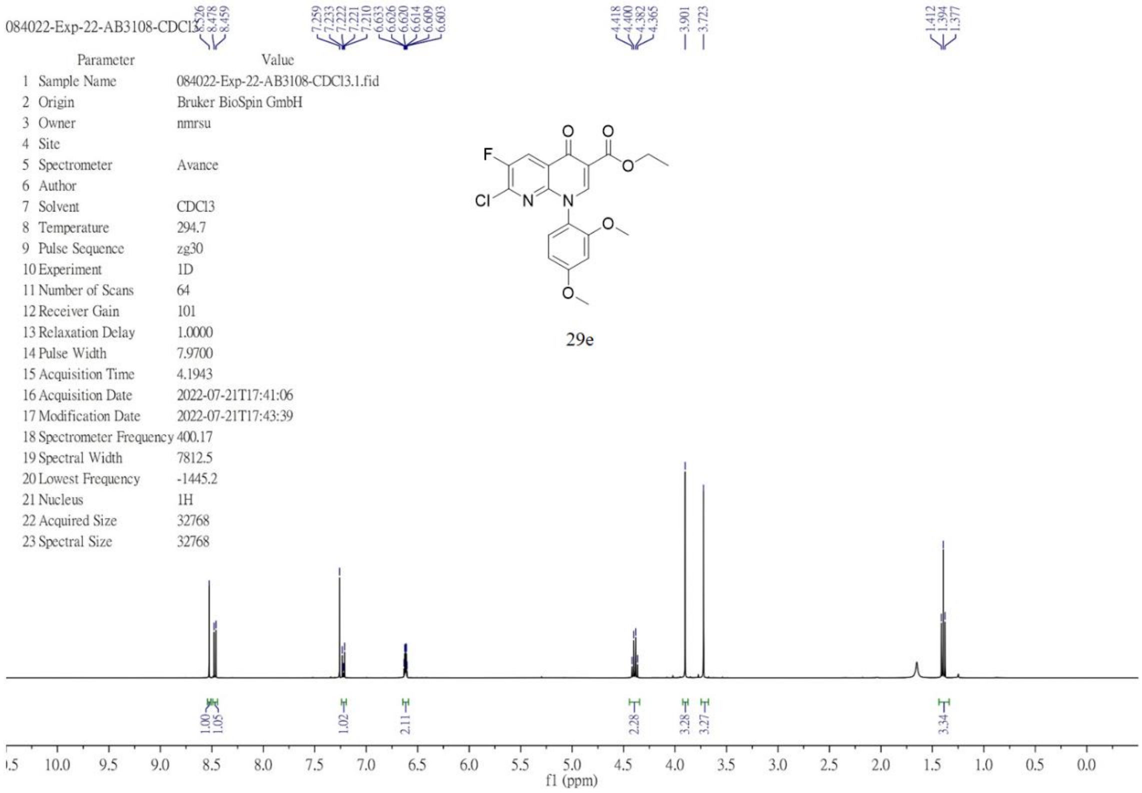


[1]H NMR (400 MHz, chloroform-*d*) of **29f**.

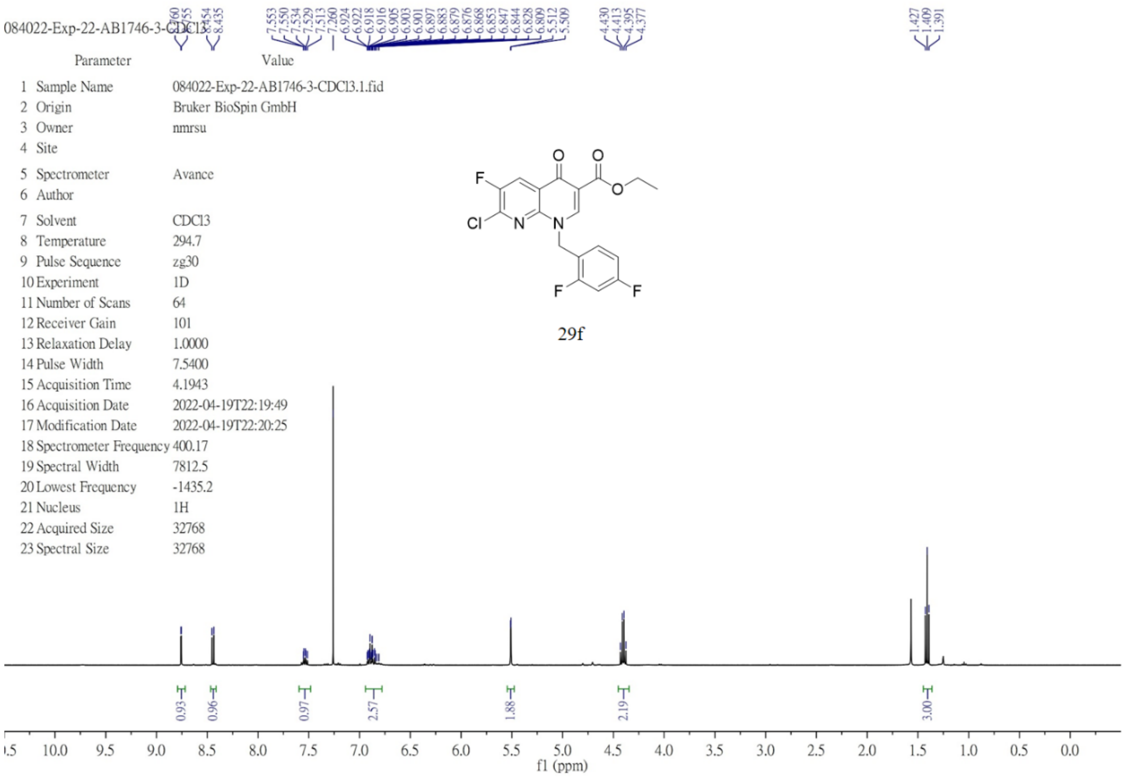

[1]H NMR (600 MHz, chloroform-*d*) of **29g**.

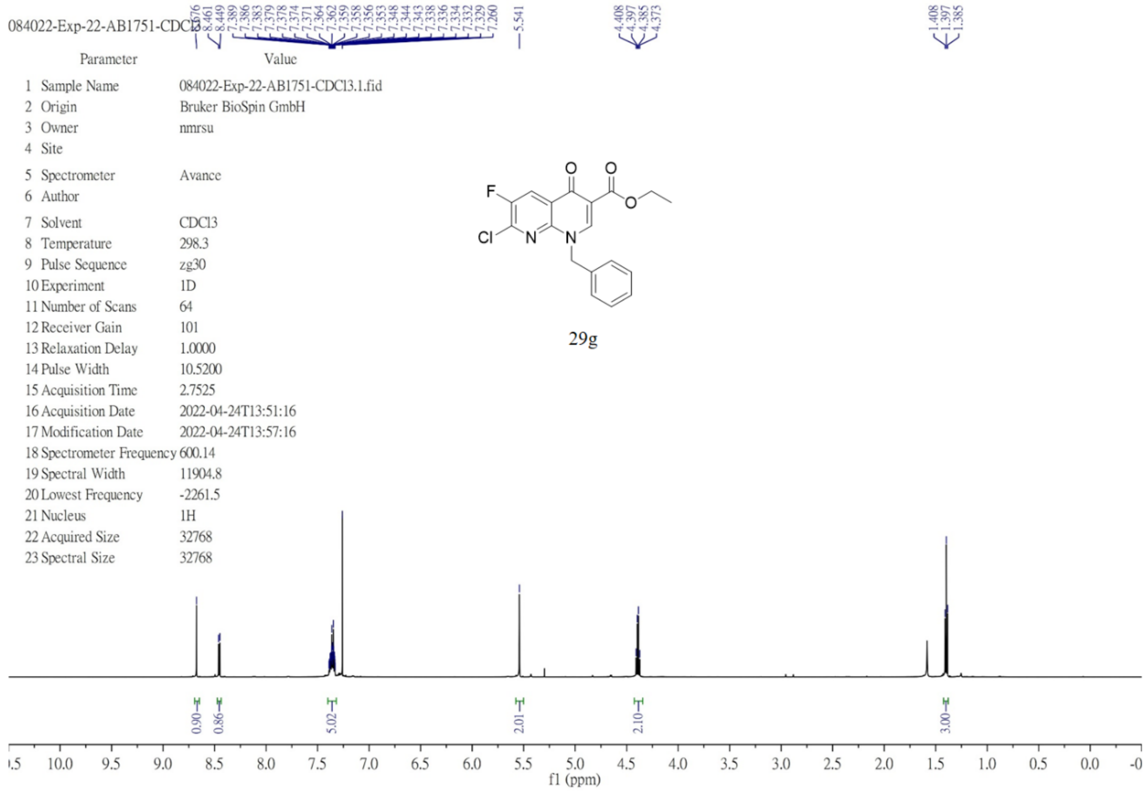


[1]H NMR (600 MHz, chloroform-*d*) of **29h**.

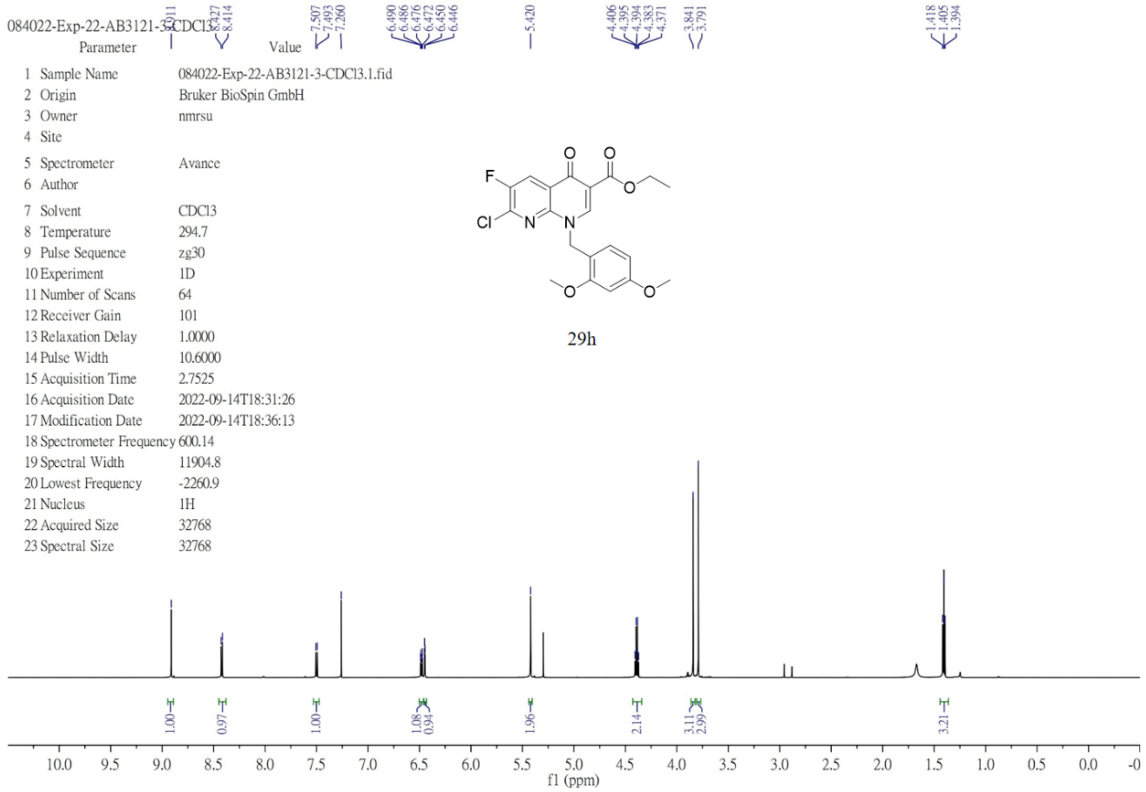

[1]H NMR (600 MHz, chloroform-*d*) of **29i**.

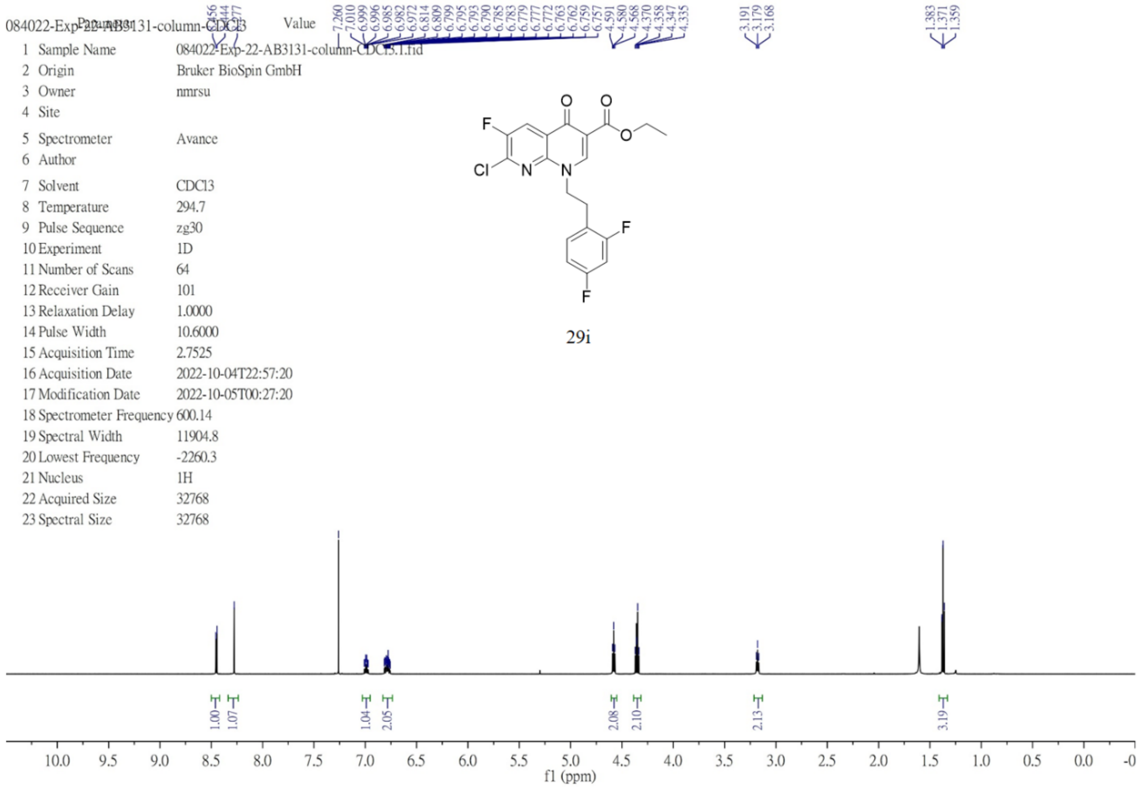


[1]H NMR (600 MHz, chloroform-*d*) of **29j**.

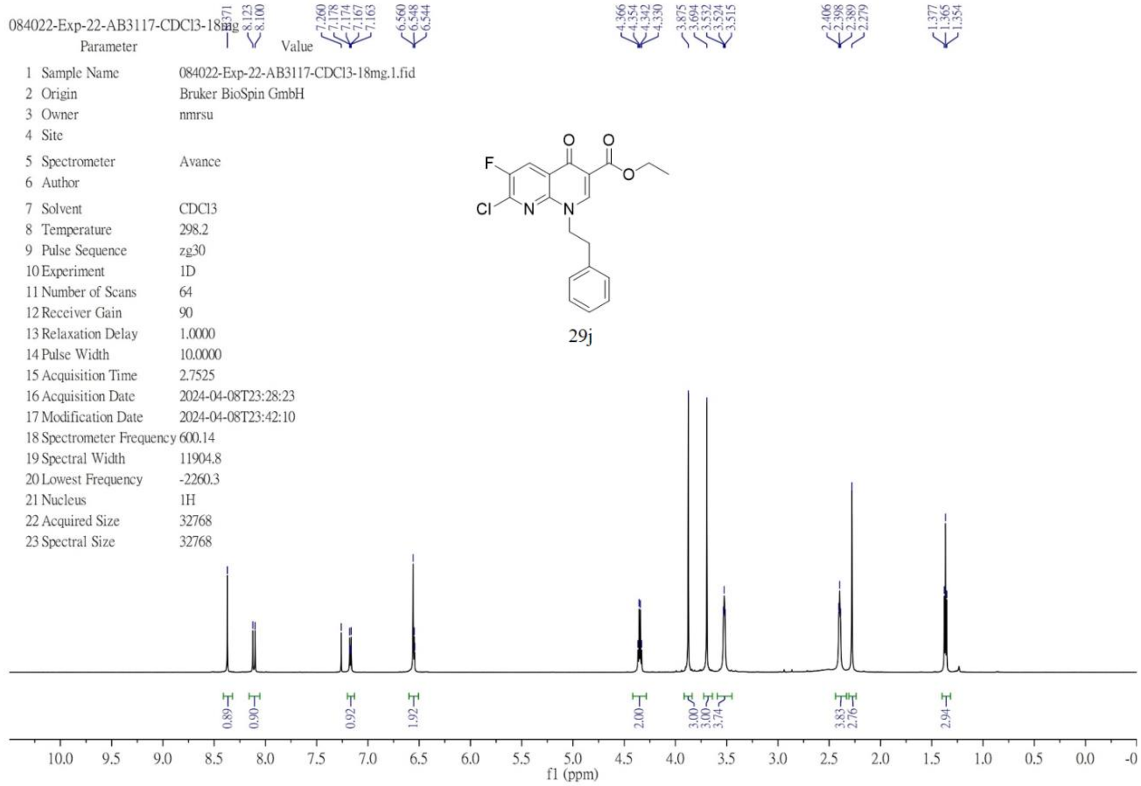

$^1$H NMR (400 MHz, chloroform-*d*) of **29k**.

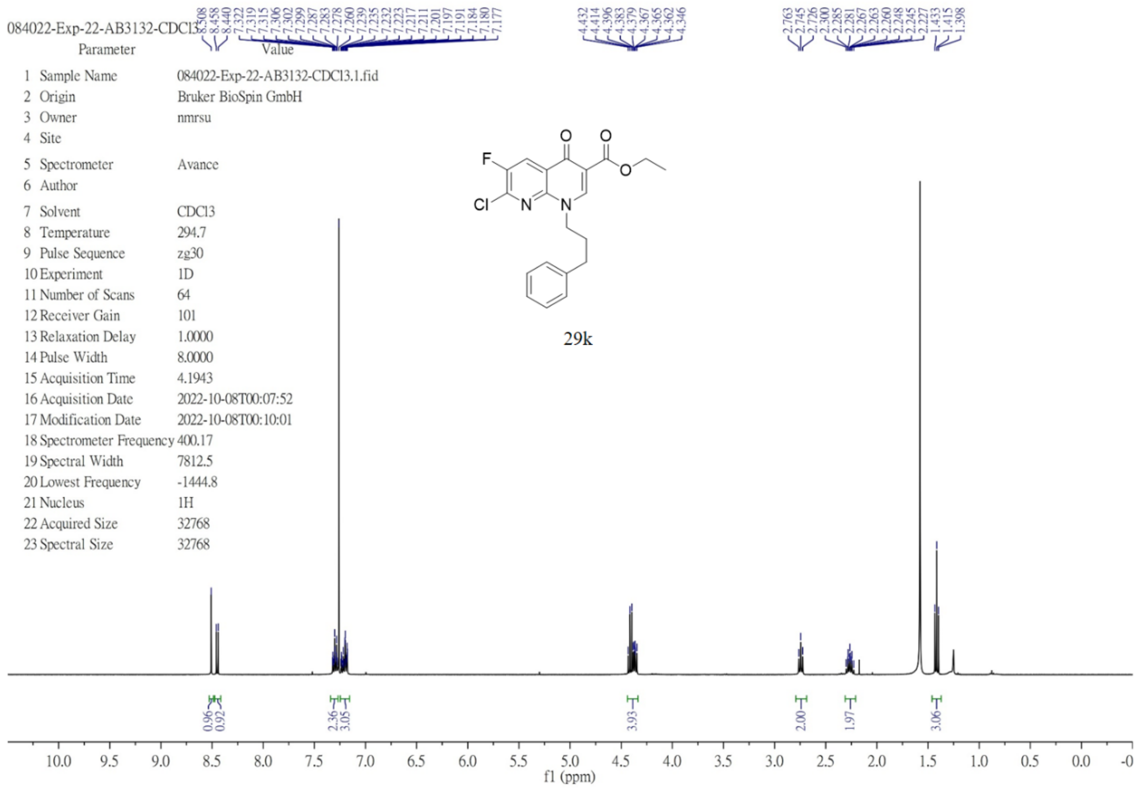


$^1$H NMR (400 MHz, chloroform-*d*) of **29l**.

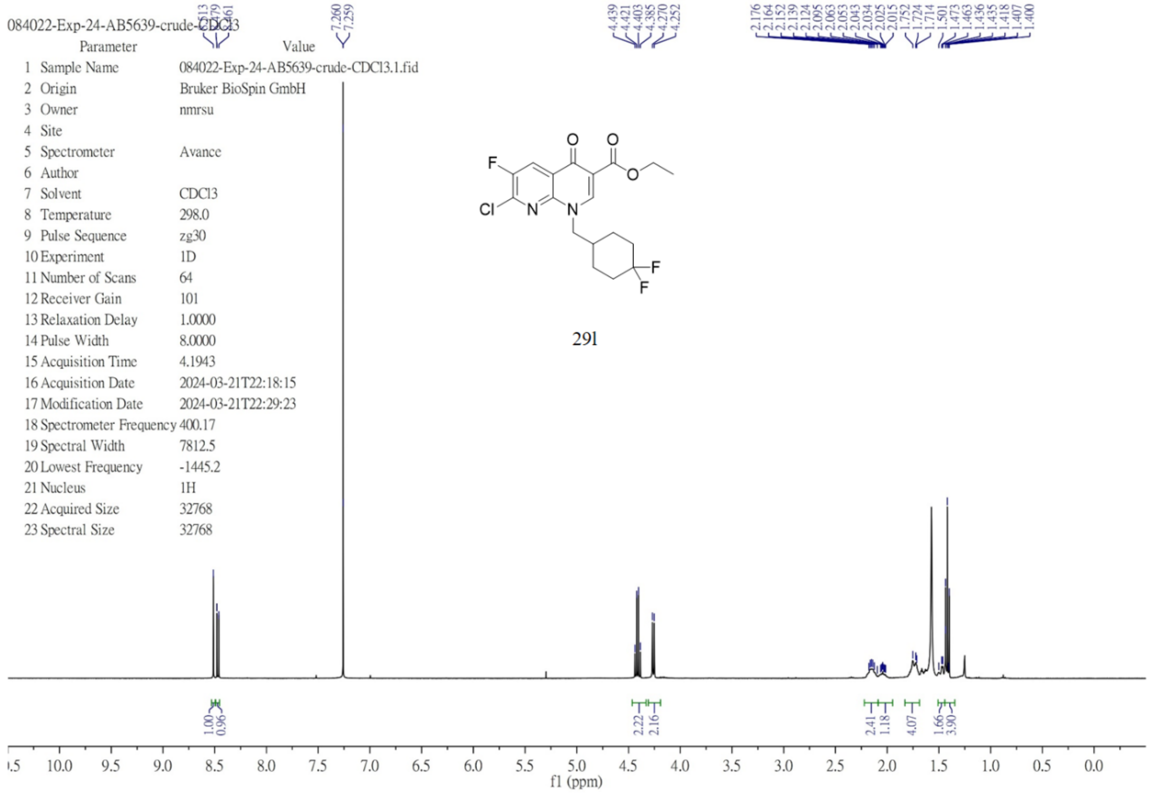

[1]H NMR (600 MHz, DMSO-$d_6$) of **30a**.

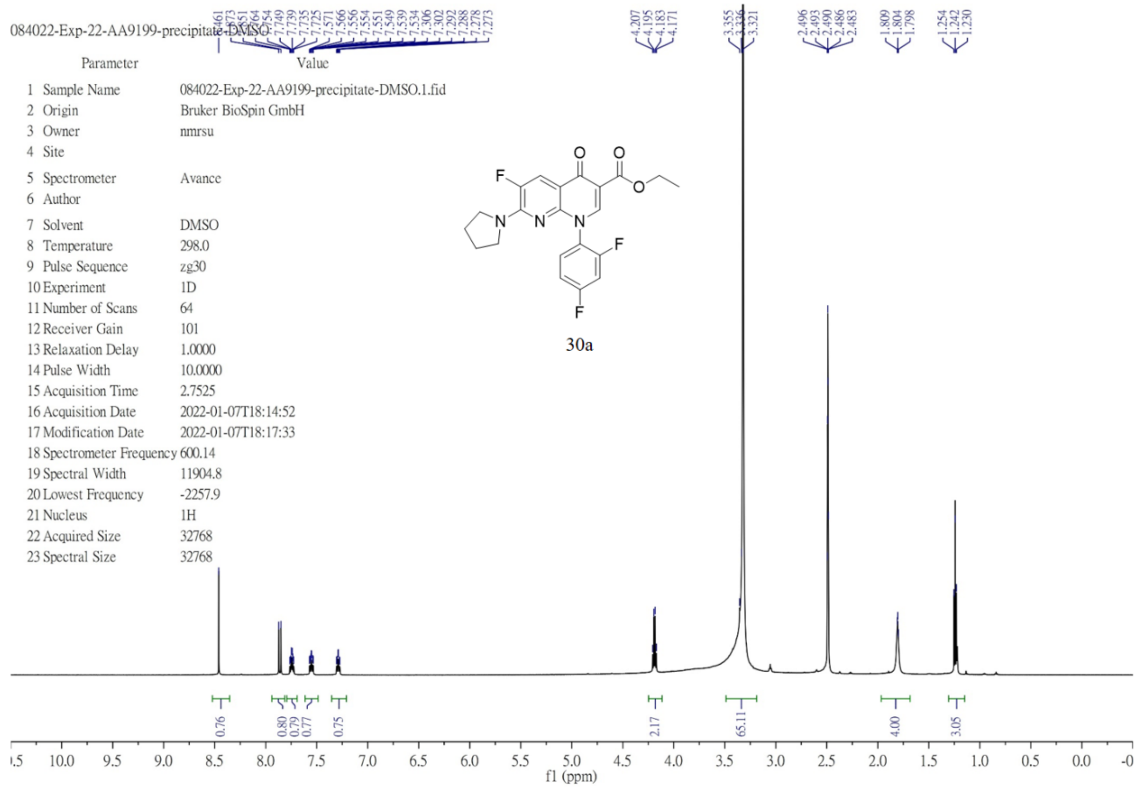

[1]H NMR (600 MHz, chloroform-$d$) of **30b**.

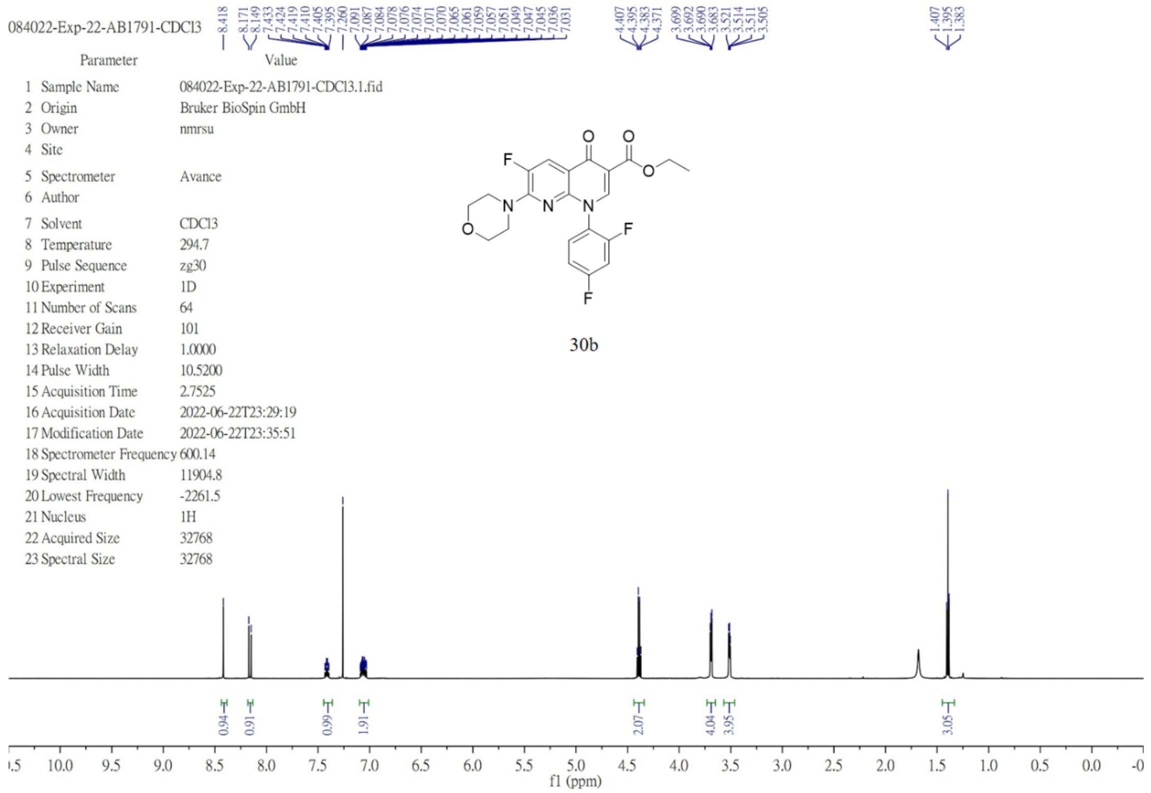

$^1$H NMR (400 MHz, chloroform-*d*) of **31a**.

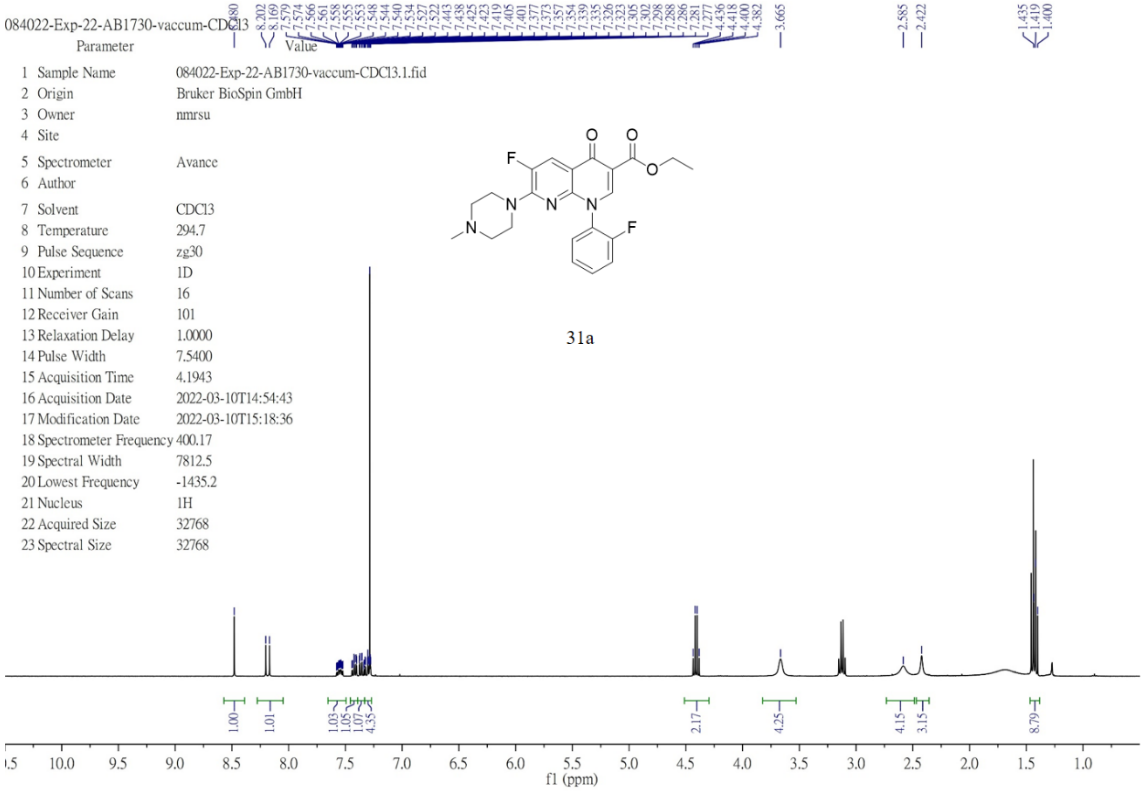

$^1$H NMR (400 MHz, chloroform-*d*) of **31b**.

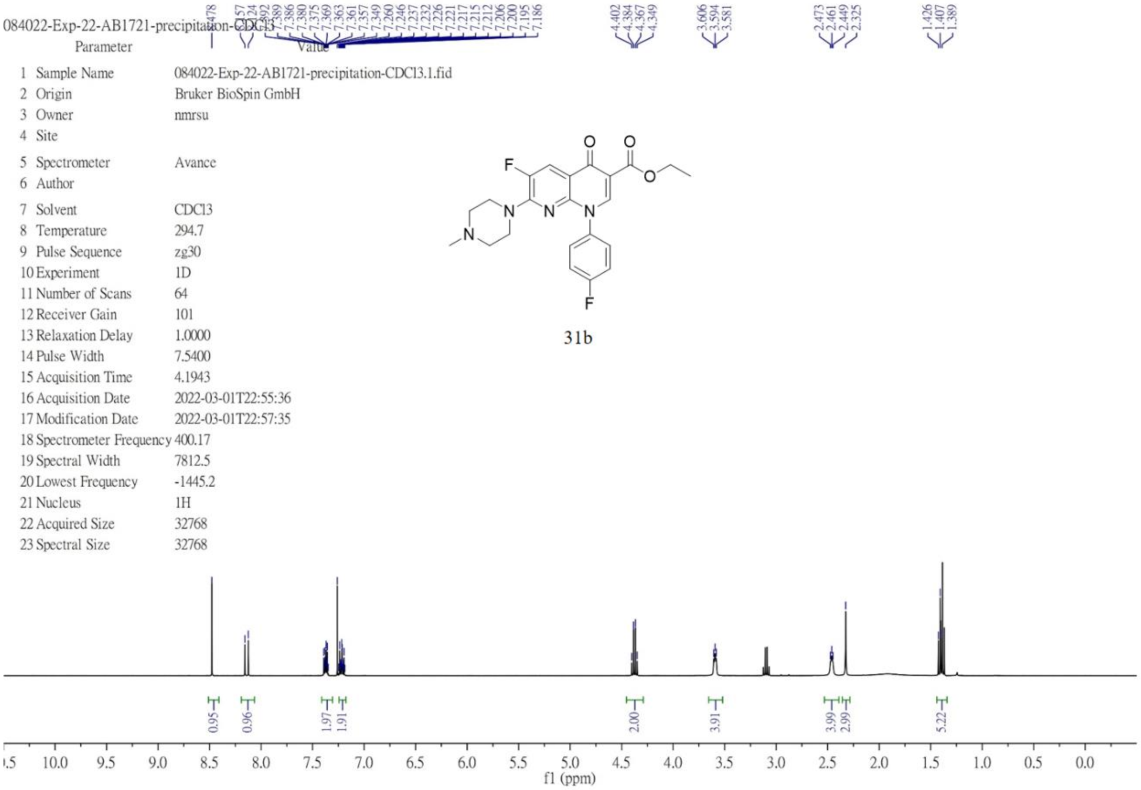

$^{1}$H NMR (400 MHz, chloroform-*d*) of **31c**.

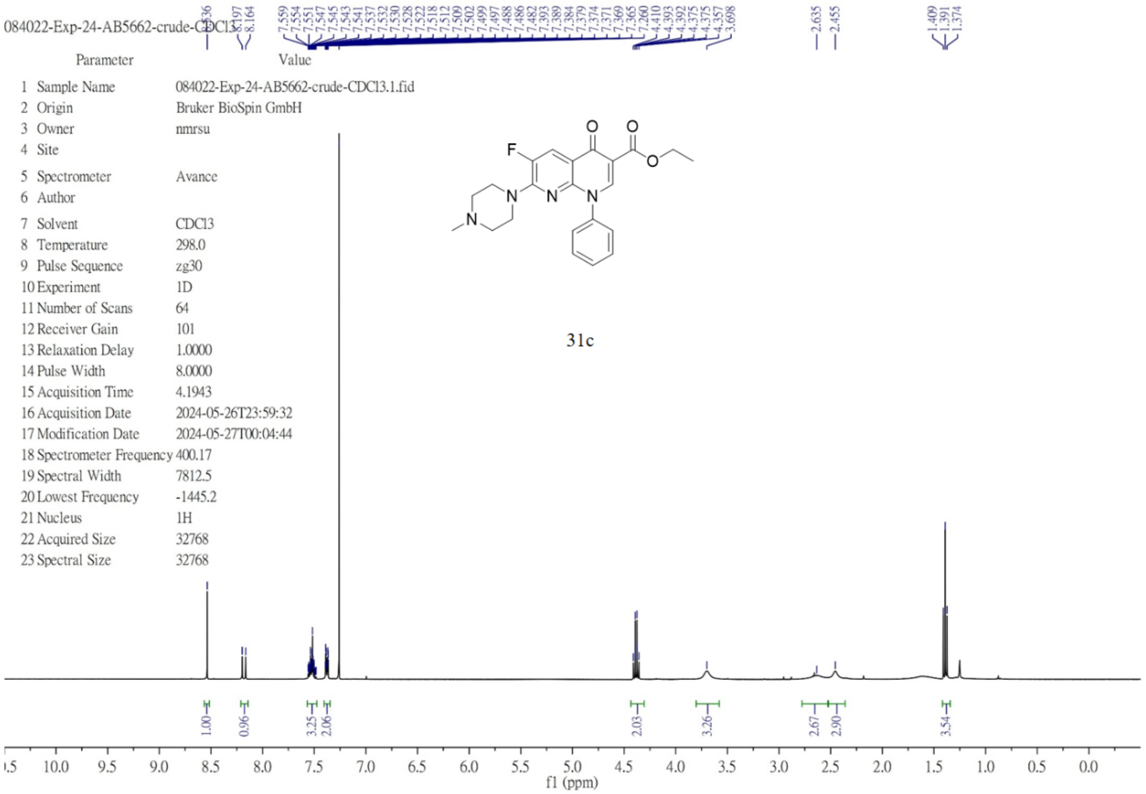


$^{1}$H NMR (600 MHz, chloroform-*d*) of **31d**.

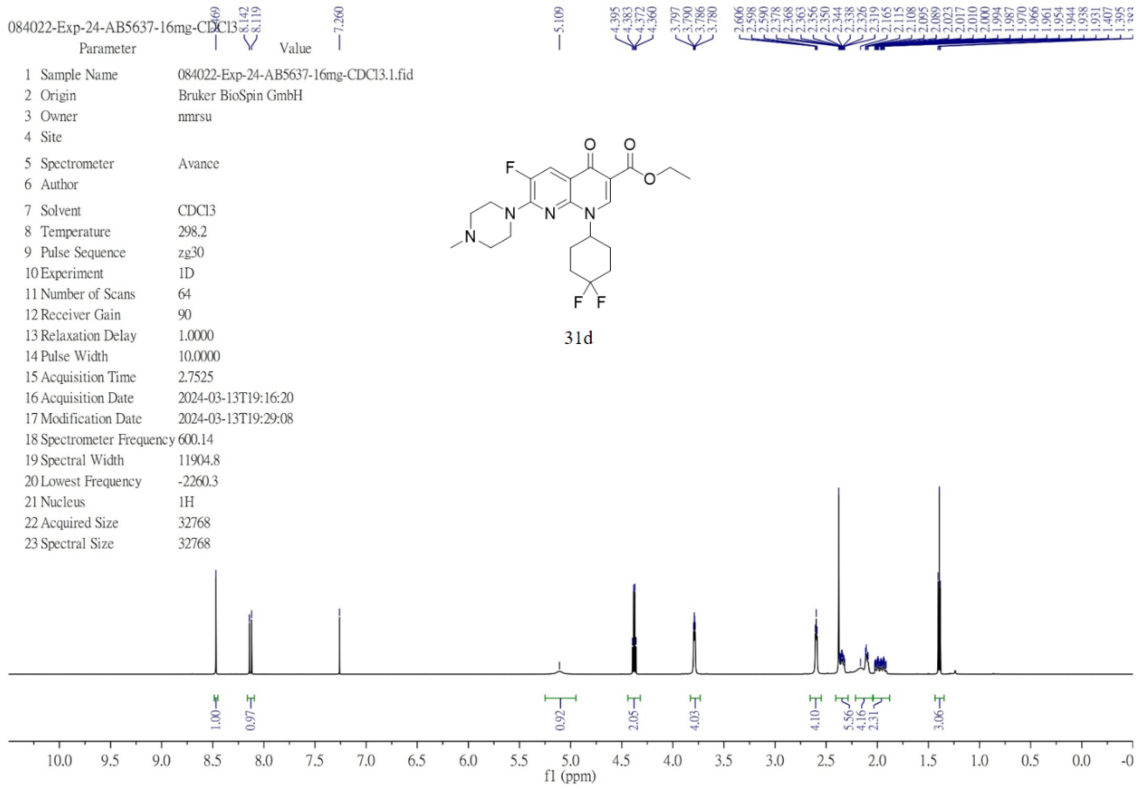

[1]H NMR (600 MHz, chloroform-*d*) of **31e (26 in the main text)**.

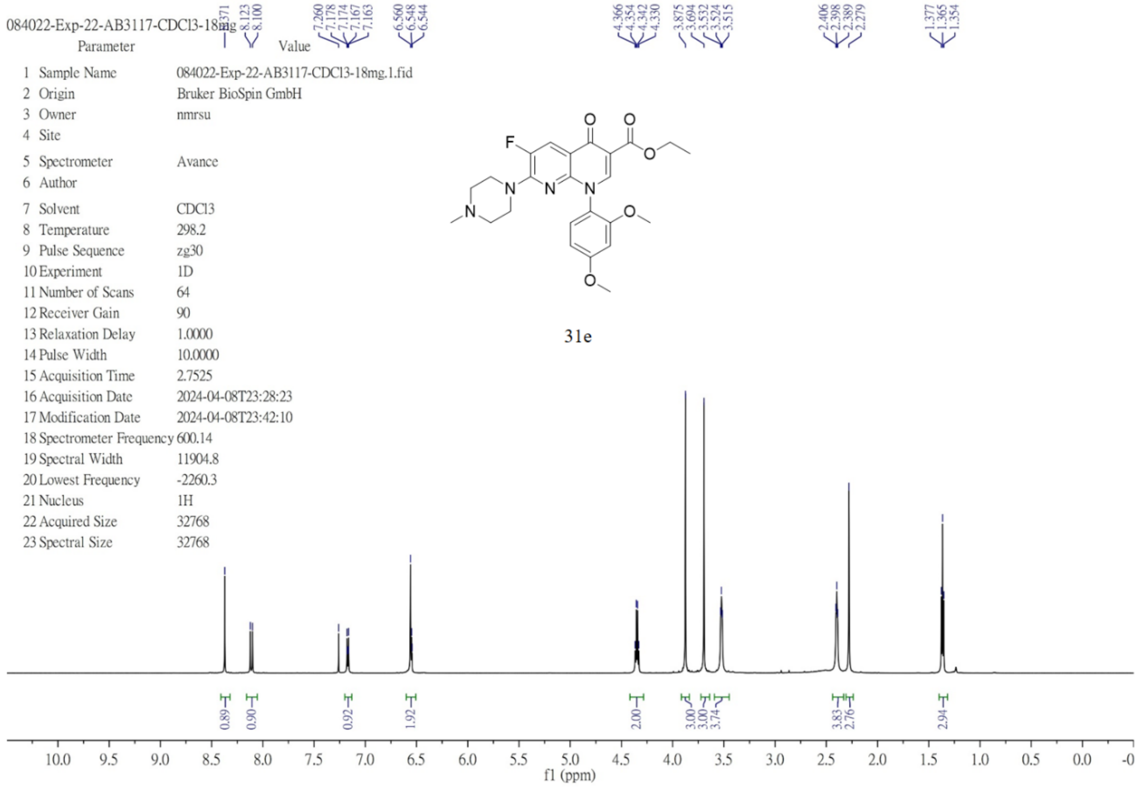


[13]C NMR (151 MHz, chloroform-*d*) of **31e (26 in the main text)**.

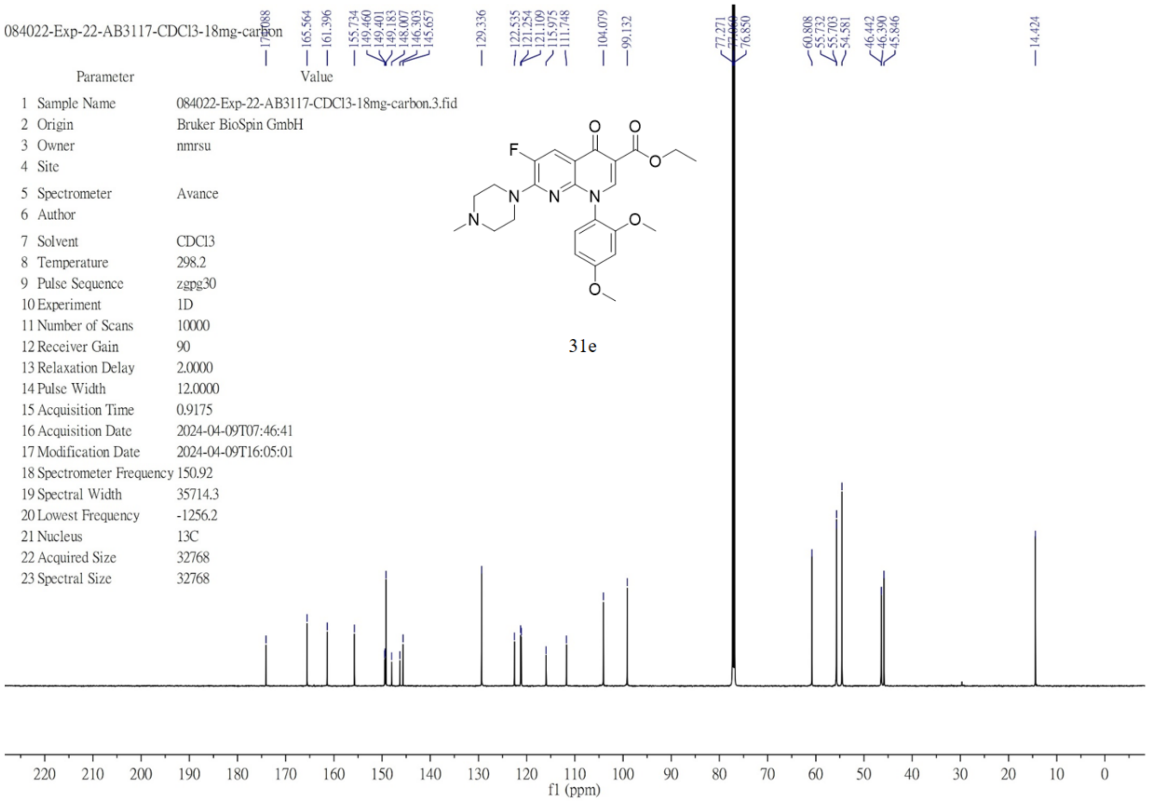

$^1$H NMR (400 MHz, chloroform-*d*) of **31f**.

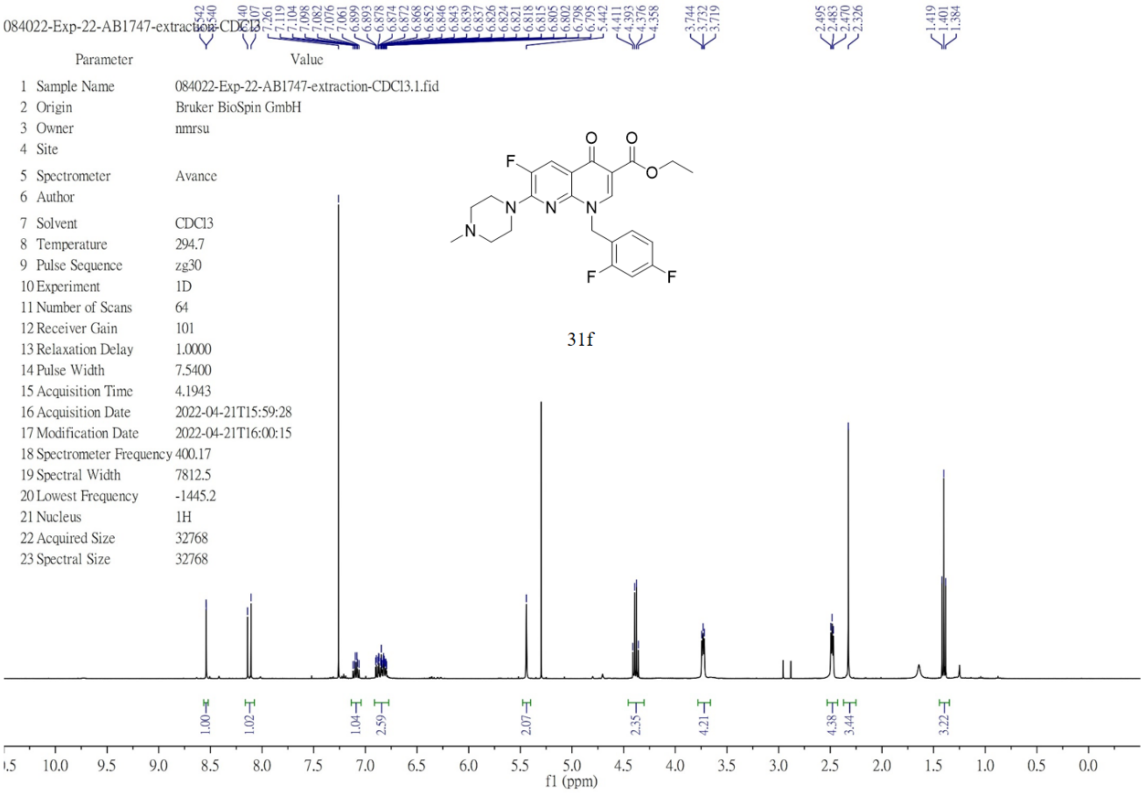

$^1$H NMR (600 MHz, chloroform-*d*) of **31g**.

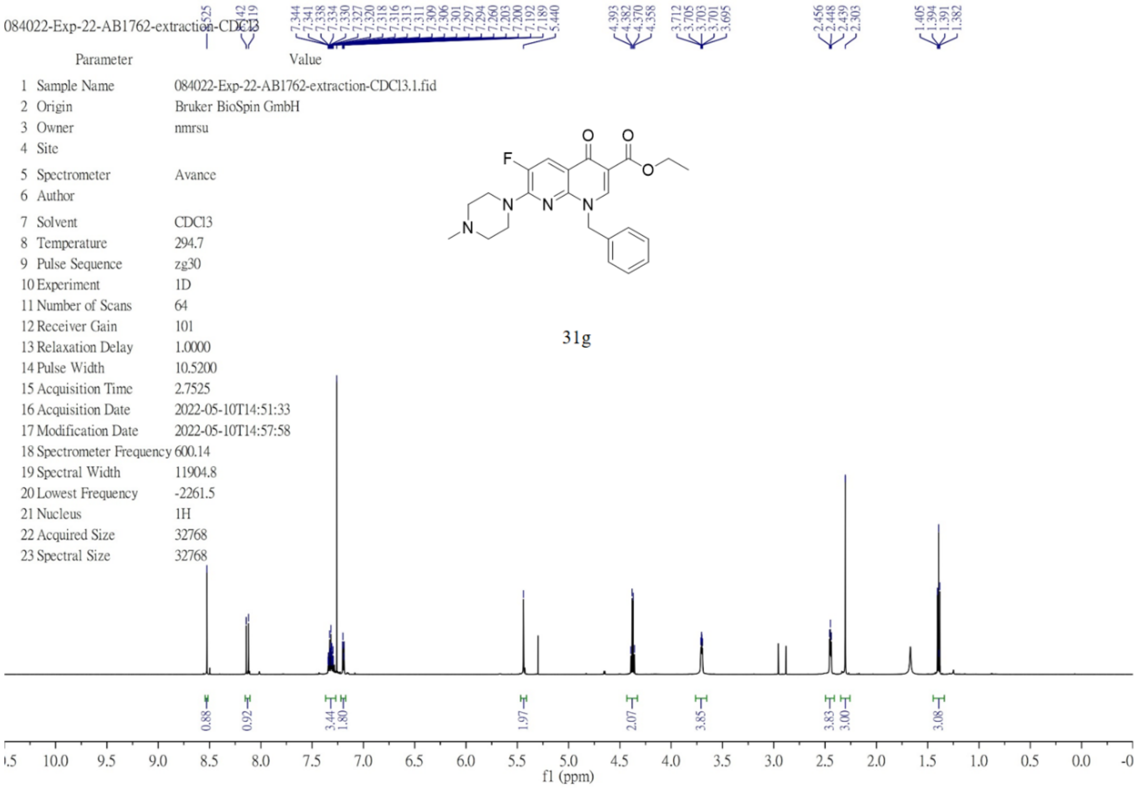

[1]H NMR (600 MHz, chloroform-*d*) of **31h (27 in the main text)**.

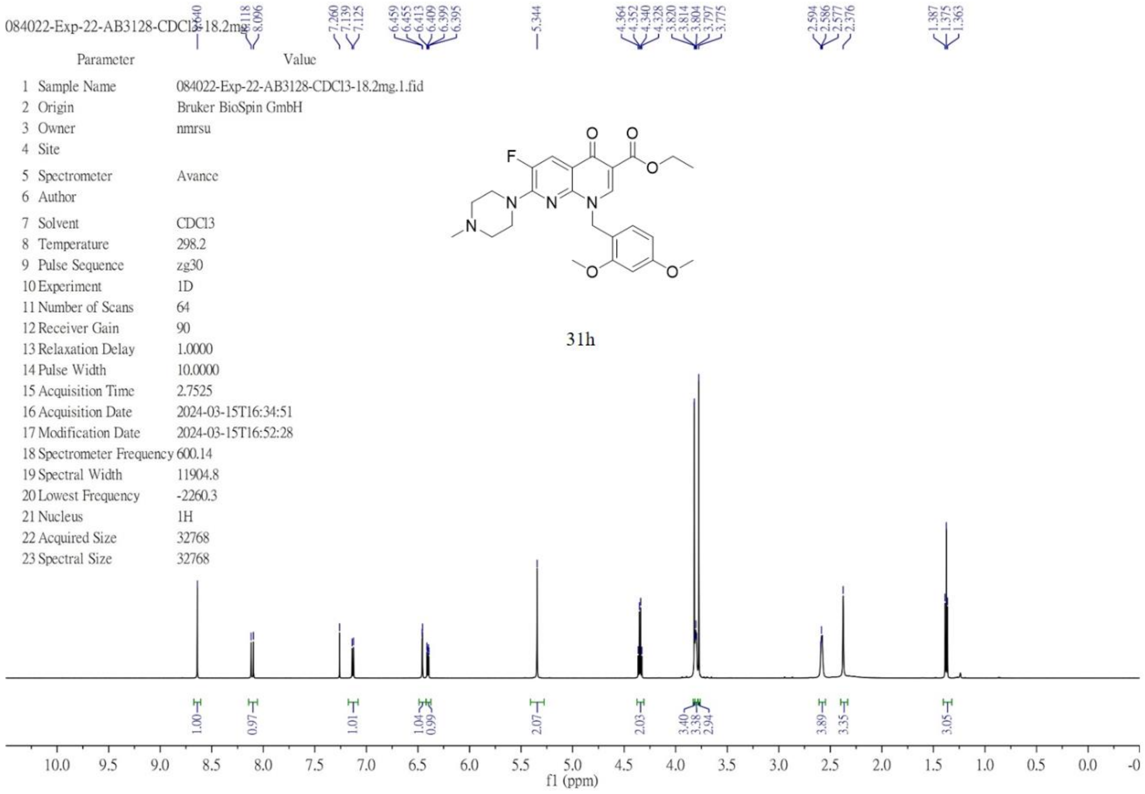


[13]C NMR (151 MHz, chloroform-*d*) of **31h (27 in the main text)**.

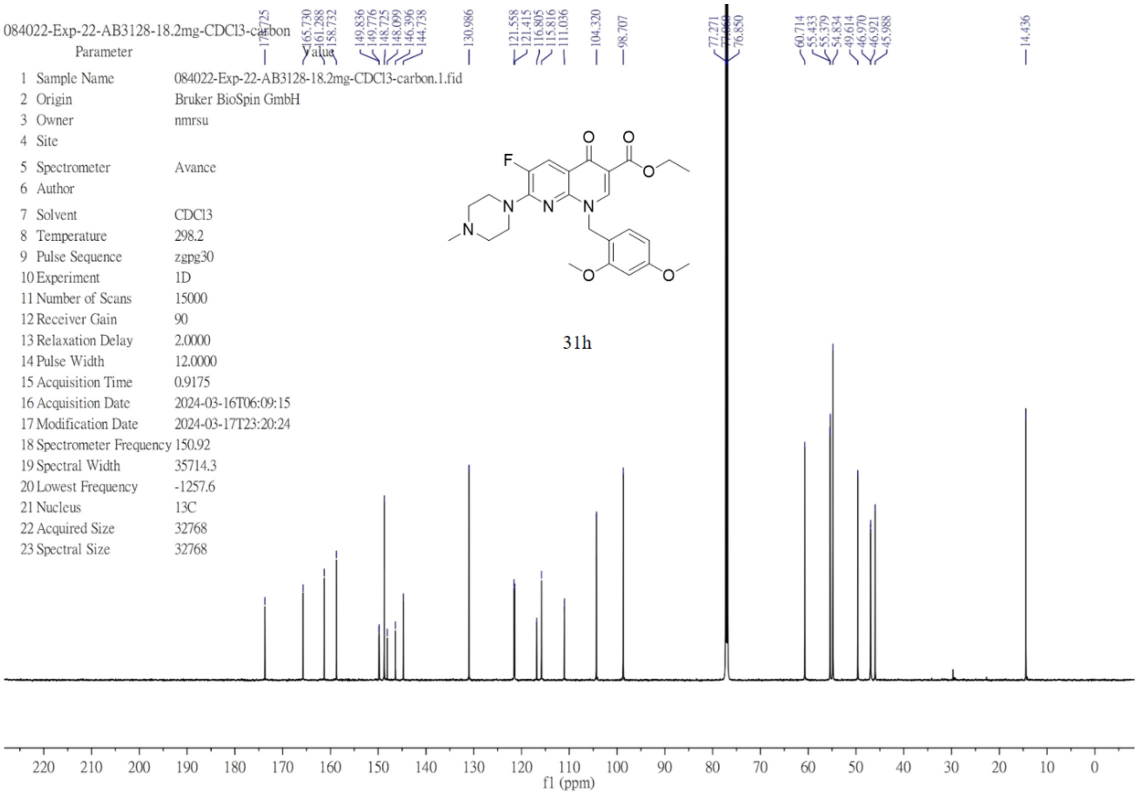

[1]H NMR (600 MHz, chloroform-*d*) of **31i**.

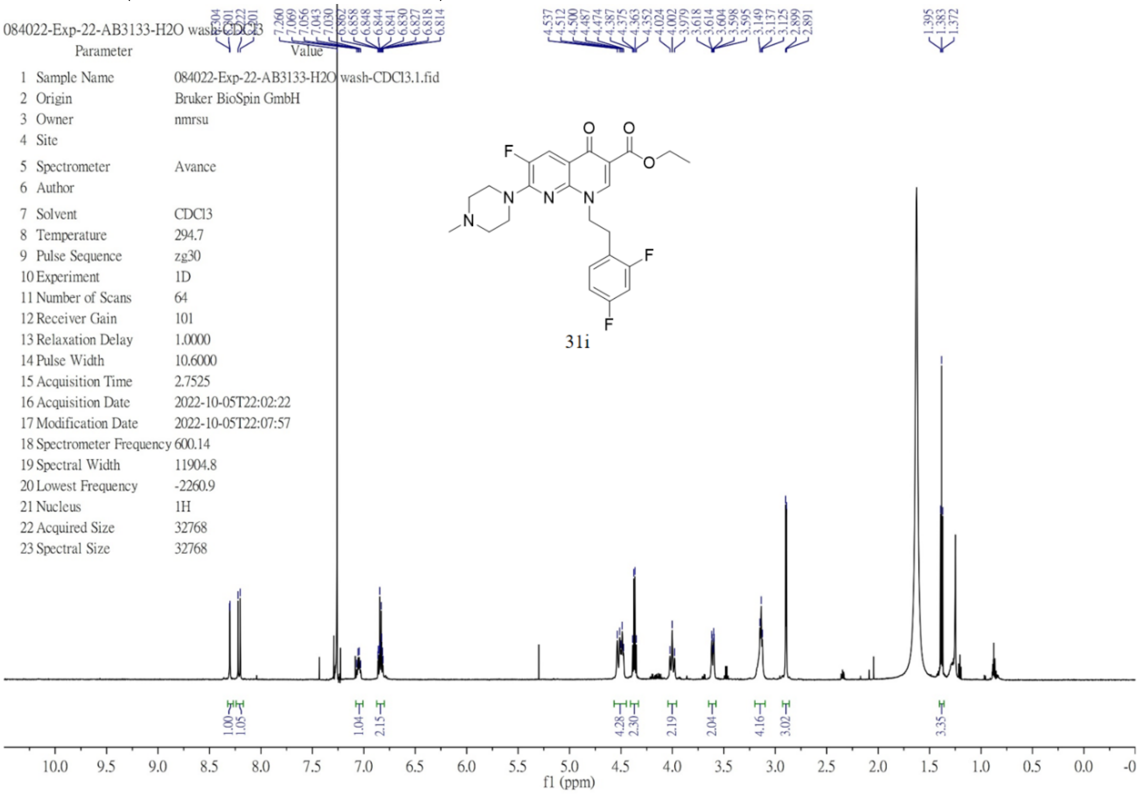


[1]H NMR (600 MHz, chloroform-*d*) of **31j**.

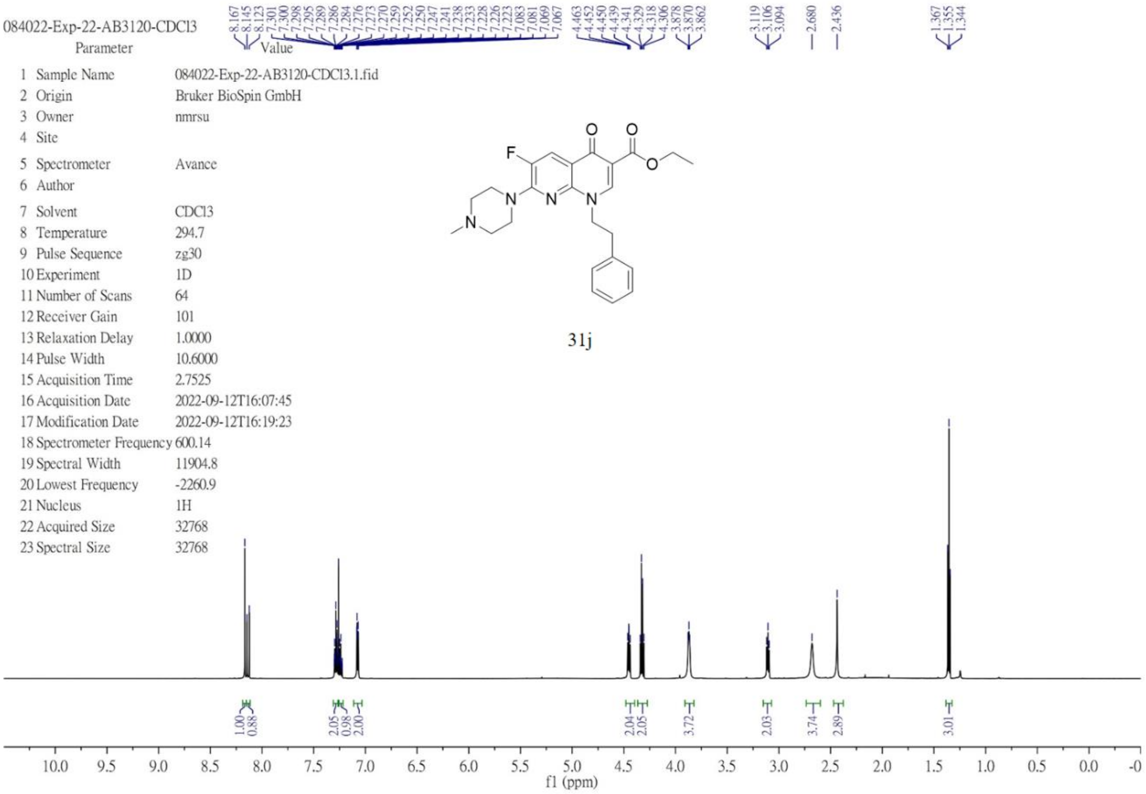

[1]H NMR (400 MHz, chloroform-*d*) of **31k**.

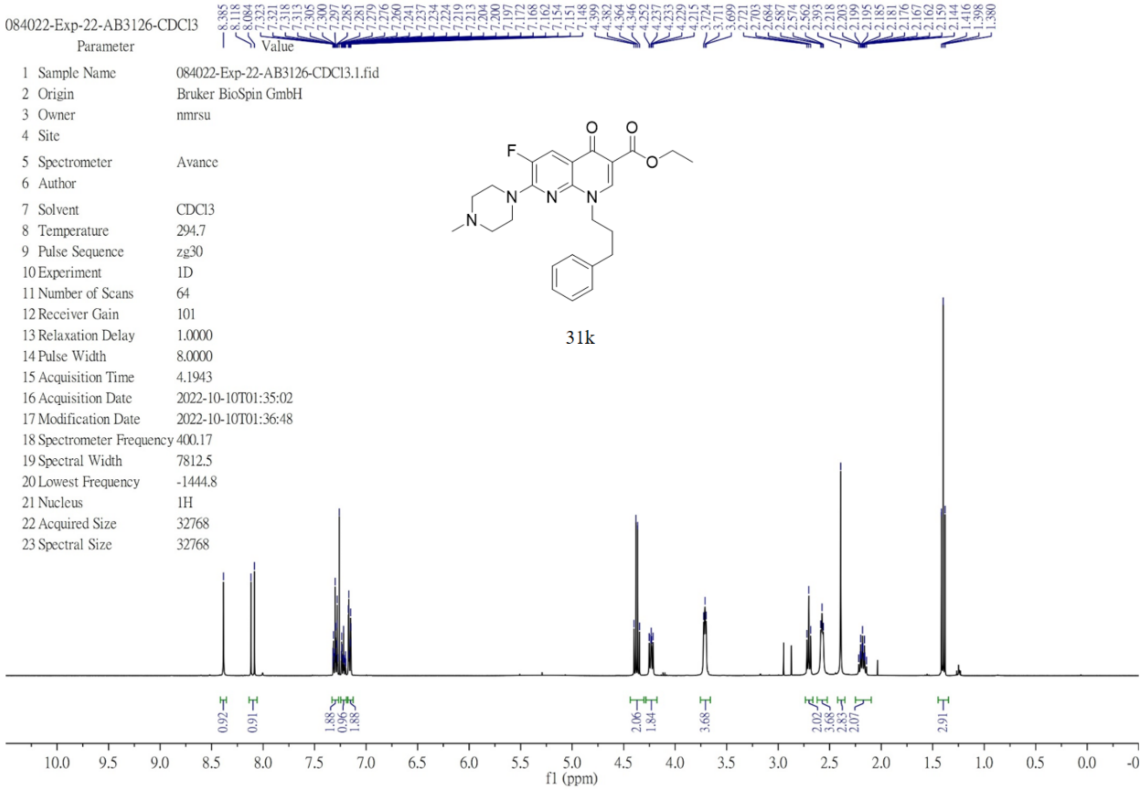


[1]H NMR (600 MHz, chloroform-*d*) of **31l (25 in the main text)**.

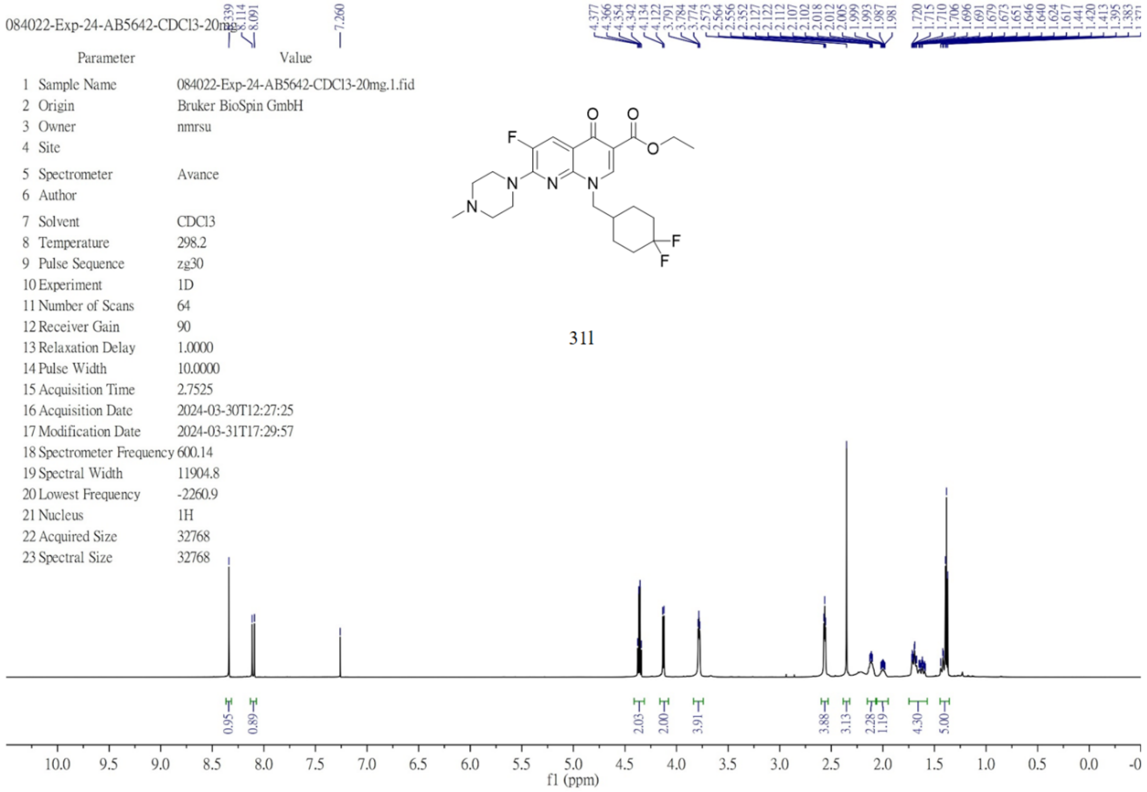

$^{13}C$ NMR (151 MHz, chloroform-*d*) of **31l (25 in the main text)**.

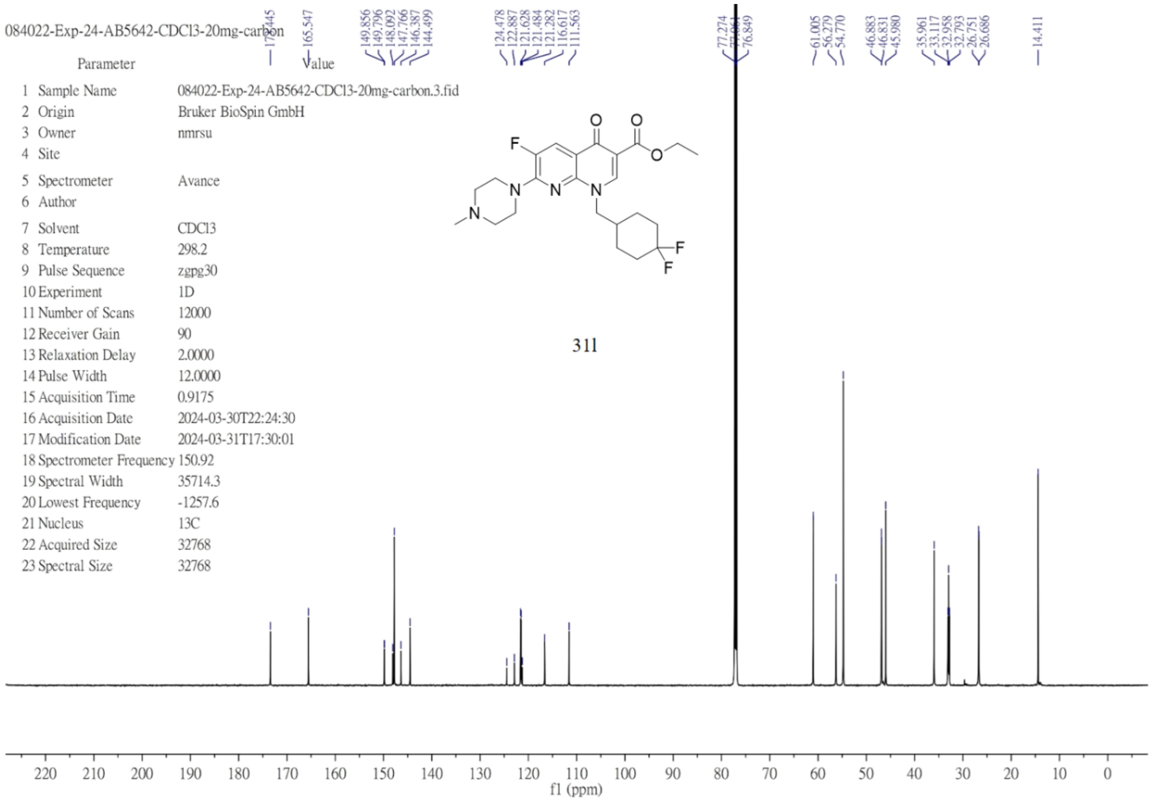

$^{1}H$ NMR (400 MHz, chloroform-*d*) of **32a**.

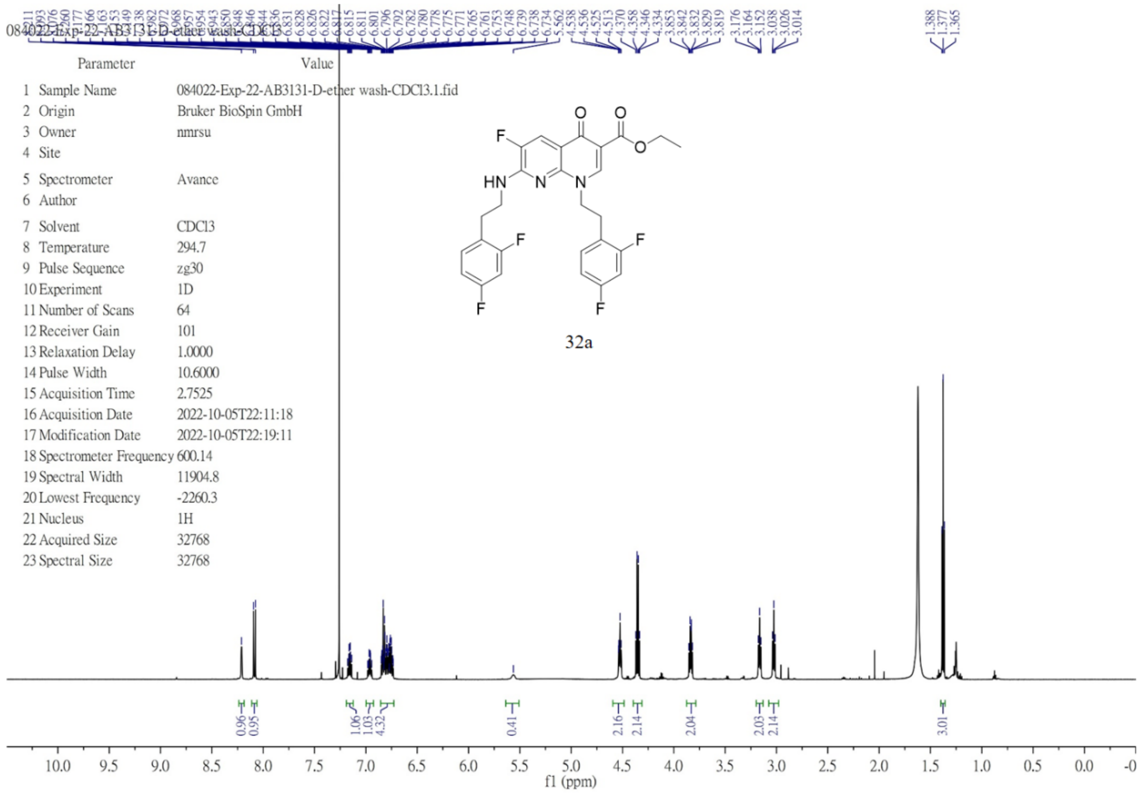

$^{1}$H NMR (400 MHz, chloroform-$d$) of **32b**.

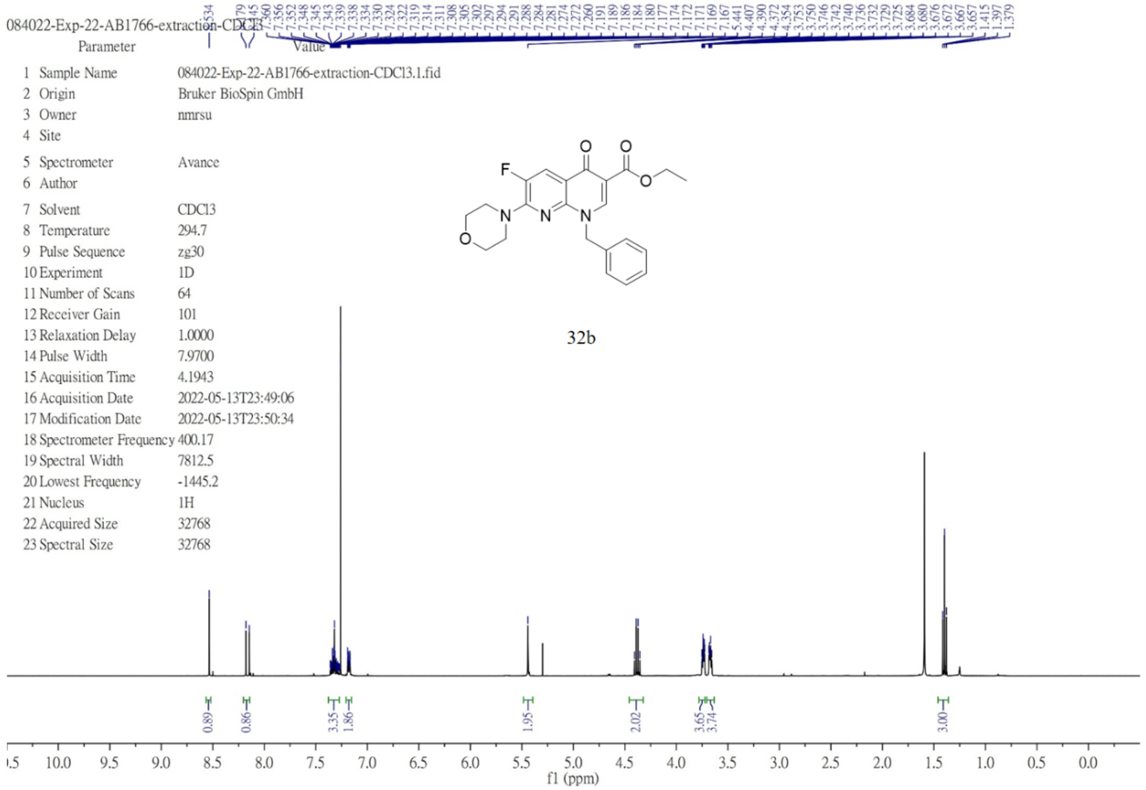


$^{1}$H NMR (600 MHz, DMSO-$d_6$) of **33**.

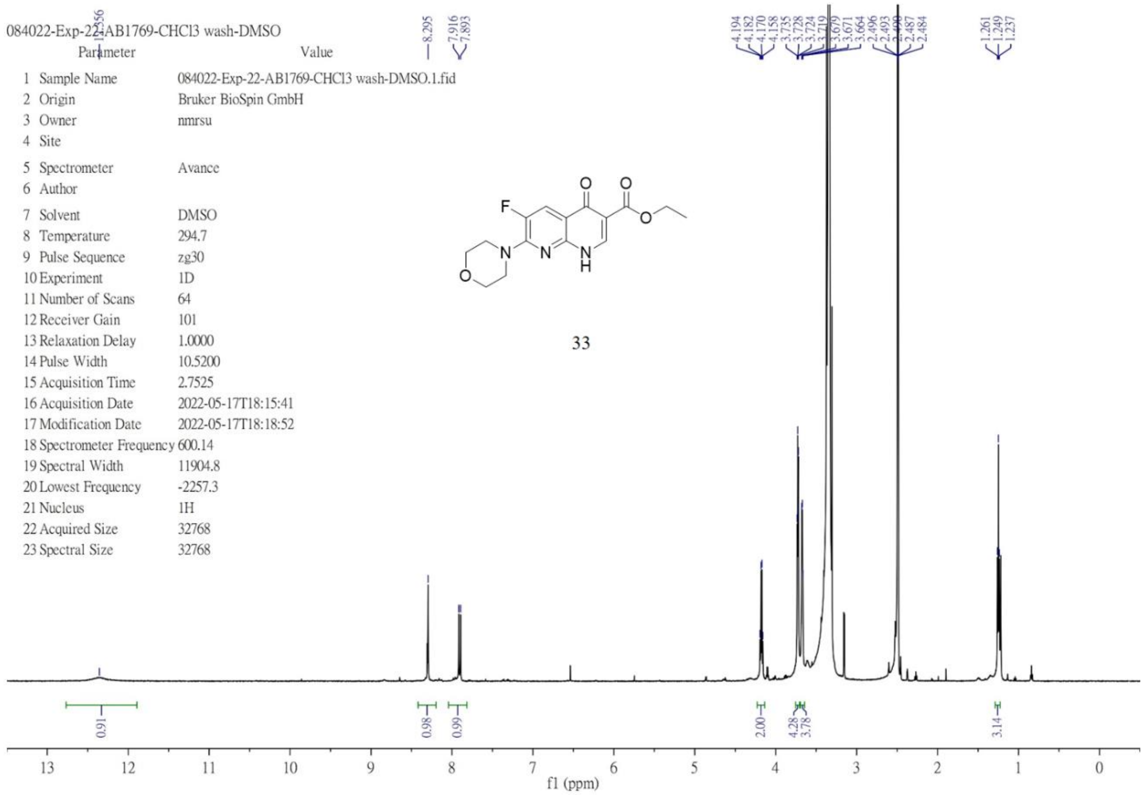

[1]H NMR (400 MHz, chloroform-*d*) of **34**.

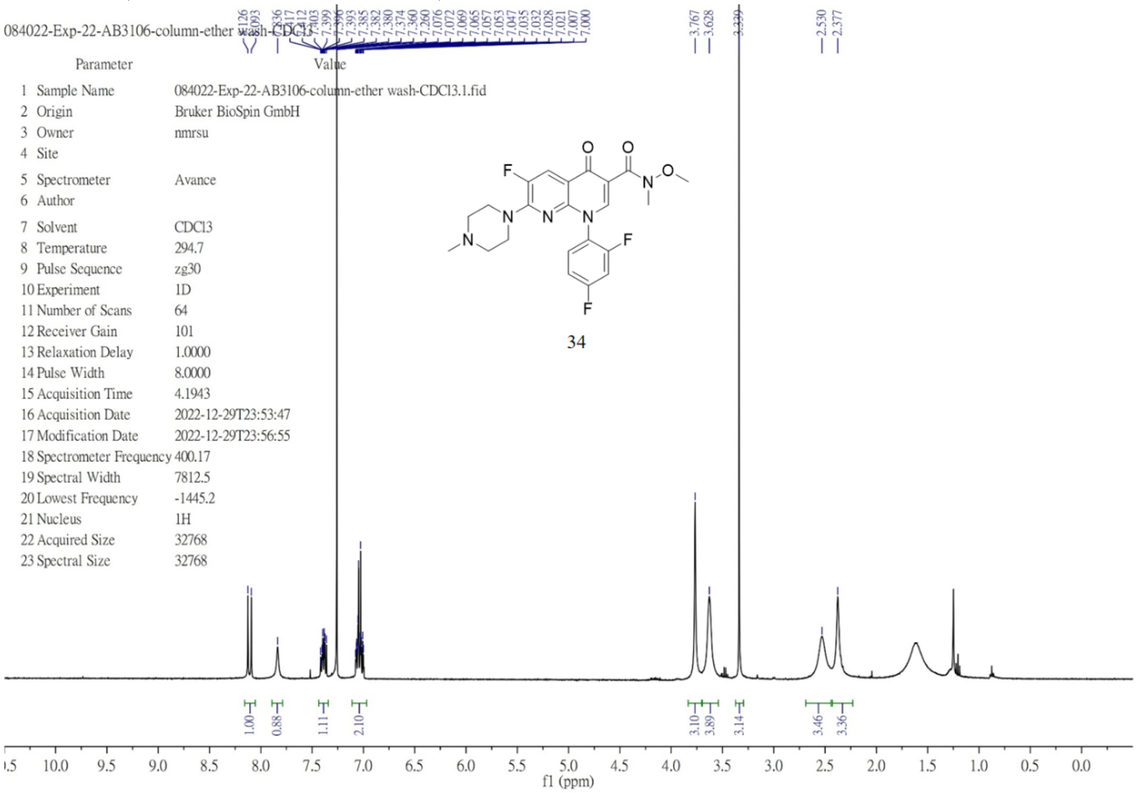


**Fig. S1.** **[1]H and [13]C spectra of synthesized compounds 1–34.**

Empower 3 SOFTWARE

Advanced Individual Report

### SAMPLE INFORMATION

| | | | |
|---|---|---|---|
| Sample Name: | EXP-22-AB3130-50 | Acquired By: | uplc |
| Sample Type: | Unknown | Sample Set Name | Purity_Test |
| Vial: | 2:B,2 | Acq. Method Set: | 5cm Std |
| Injection #: | 1 | Processing Method | Def_Processing_Method |
| Injection Volume: | 5.00 ul | Channel Name: | PDA Ch1 254nm@4.8nm |
| Run Time: | 6.5 Minutes | Proc. Chnl. Descr.: | PDA Ch1 254nm@4.8nm |
| | | eCord Name | ACQUITY UPLC?BEH C18 1.7瘐 |
| Date Acquired: | 2022/9/29 PM 06:50:23 CST | | |
| Date Processed: | 2022/9/30 AM 09:17:15 CST | | |

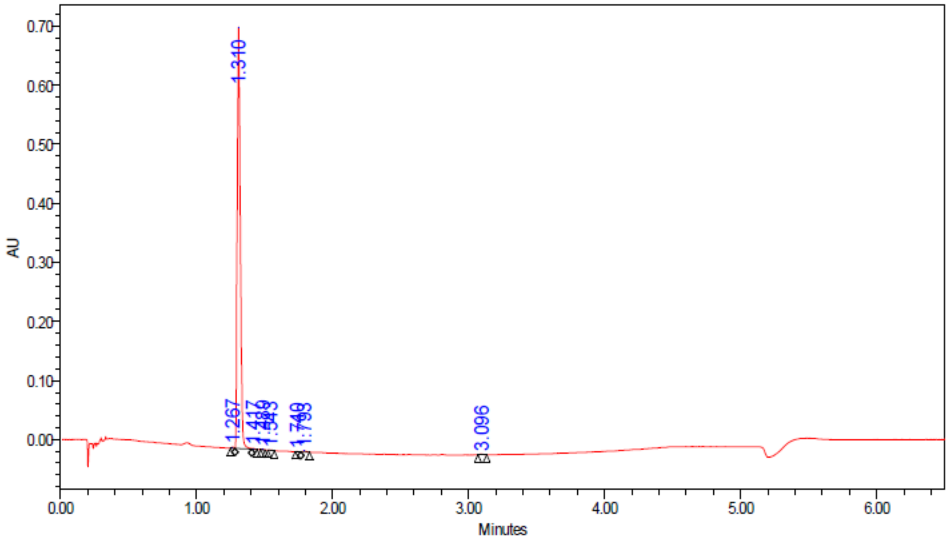


| | RT | Height | Area | % Area |
|---|---|---|---|---|
| 1 | 1.267 | 2121 | 2076 | 0.18 |
| 2 | 1.310 | 713992 | 1124821 | 98.89 |
| 3 | 1.417 | 1382 | 1732 | 0.15 |
| 4 | 1.489 | 1406 | 1532 | 0.13 |
| 5 | 1.543 | 526 | 796 | 0.07 |
| 6 | 1.740 | 577 | 742 | 0.07 |

| | RT | Height | Area | % Area |
|---|---|---|---|---|
| 7 | 1.793 | 1952 | 3316 | 0.29 |
| 8 | 3.096 | 1426 | 2458 | 0.22 |

**Fig. S2. HPLC trace of compound 12.**

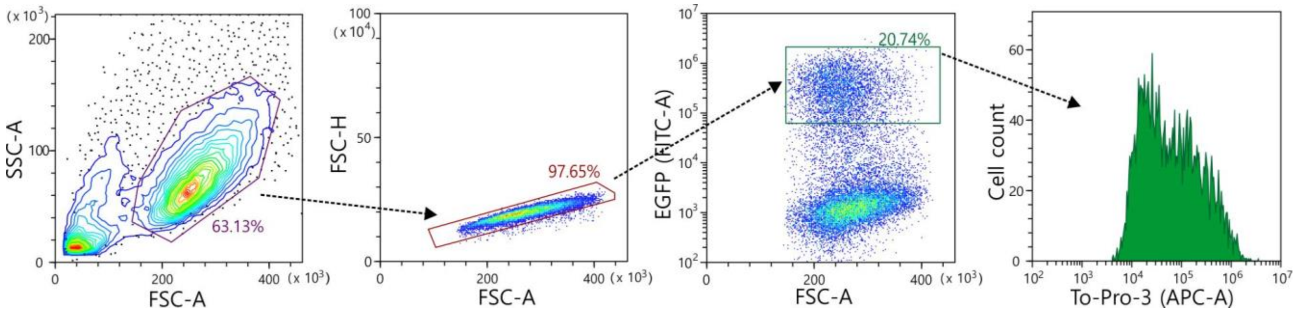


**Fig. S3. Electronic gating strategy for analyzing To-Pro-3 uptake in the HEK293T cells expressing different hPANX1 constructs.** Flow cytometry analyses showing electronic gating strategy used to determine the mean fluorescence intensity (MFI) of To-Pro-3 uptake from EGFP$^{High}$ HEK293T cells expressing either hPANX1-FL-EGPF (not shown) or hPANX1-CT-EGFP.

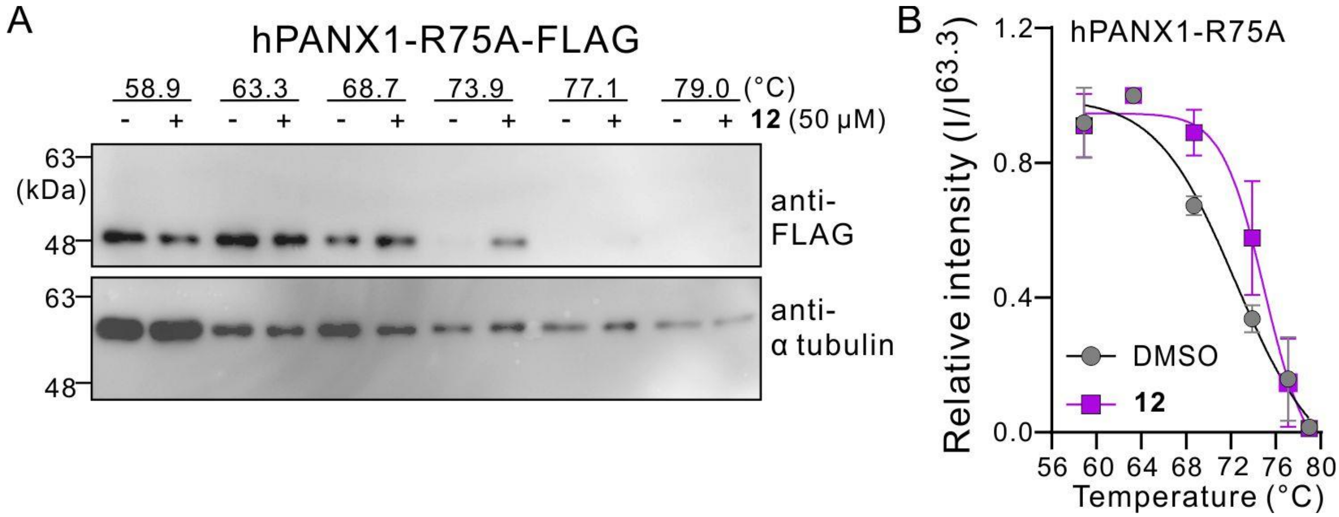


**Fig. S4. Application of compound 12 increased $T_{agg}$ of hPANX1-R75A. (A)** Representative immunoblots of cell thermal shift assays (CETSA) obtained from HEK293T cells expressing hPANX1-R75A-FLAG, with or without exposure to compound **12** (50 μM), at indicated temperatures. Anti-α-tubulin was used as a loading control. **(B)** Grouped results showing the intensity of immunoreactive signals from hPANX1-R75A at different temperatures, relative to that at 63.3℃. $T_{agg}$ of DMSO: 72.7℃; $T_{agg}$ of compound **12**: 75.1℃ (n=4 biologically independent experiments).

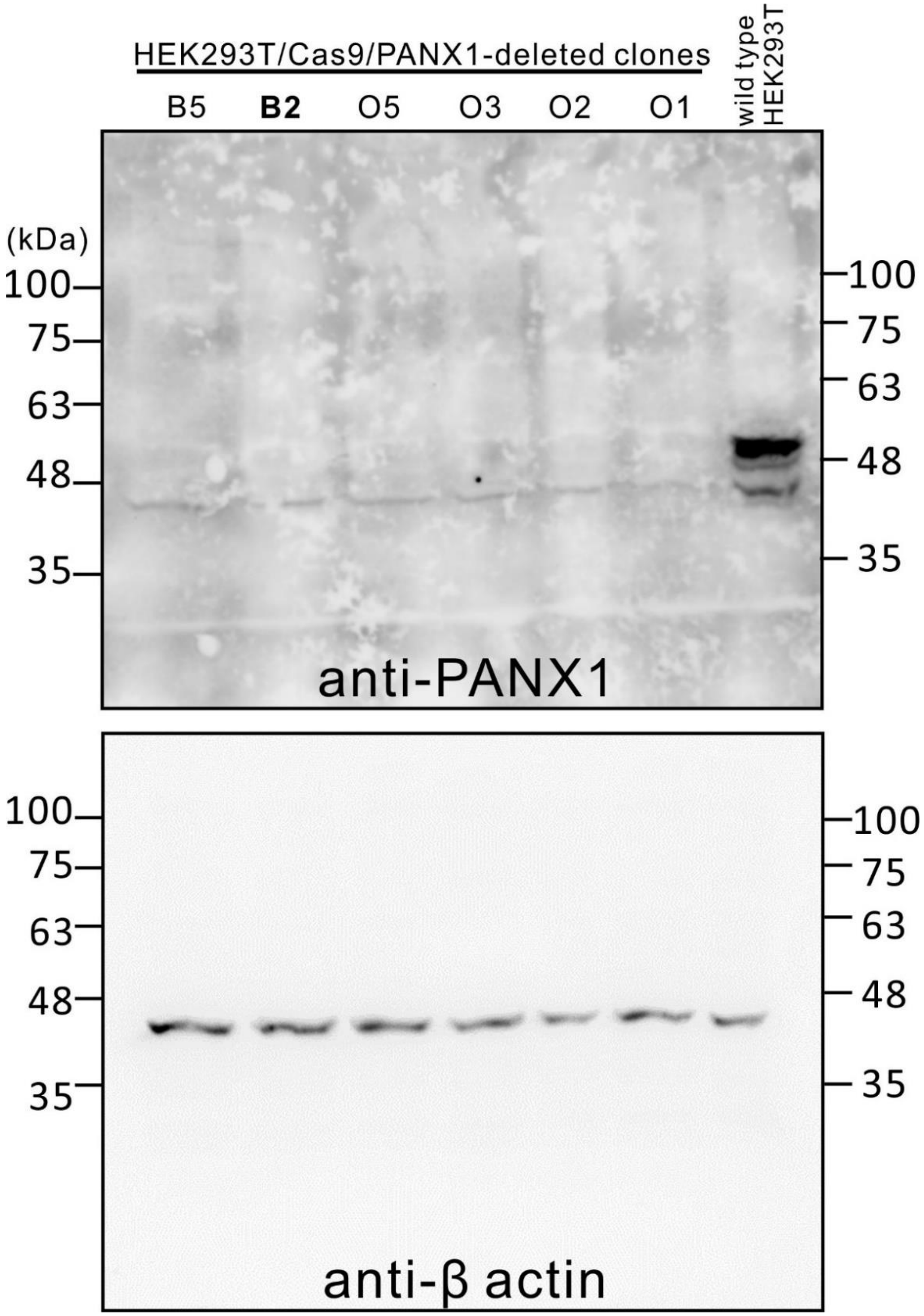


**Fig. S5. PANX1-deleted HEK293T cells.** Representative Western blots showed that expressions of PANX1 proteins in wild type HEK293T cells, but not in various clones of PANX1-deleted cells. An antibody to PANX1 (Cell Signaling; # 91137; 1:1000) was used to detect the endogenously expressed PANX1 proteins. Anti-β actin (Novus Biologicals; clone AC-15; # NB600-501; 1:5000) was used as a loading control. The current study used cells derived from the clone B2.

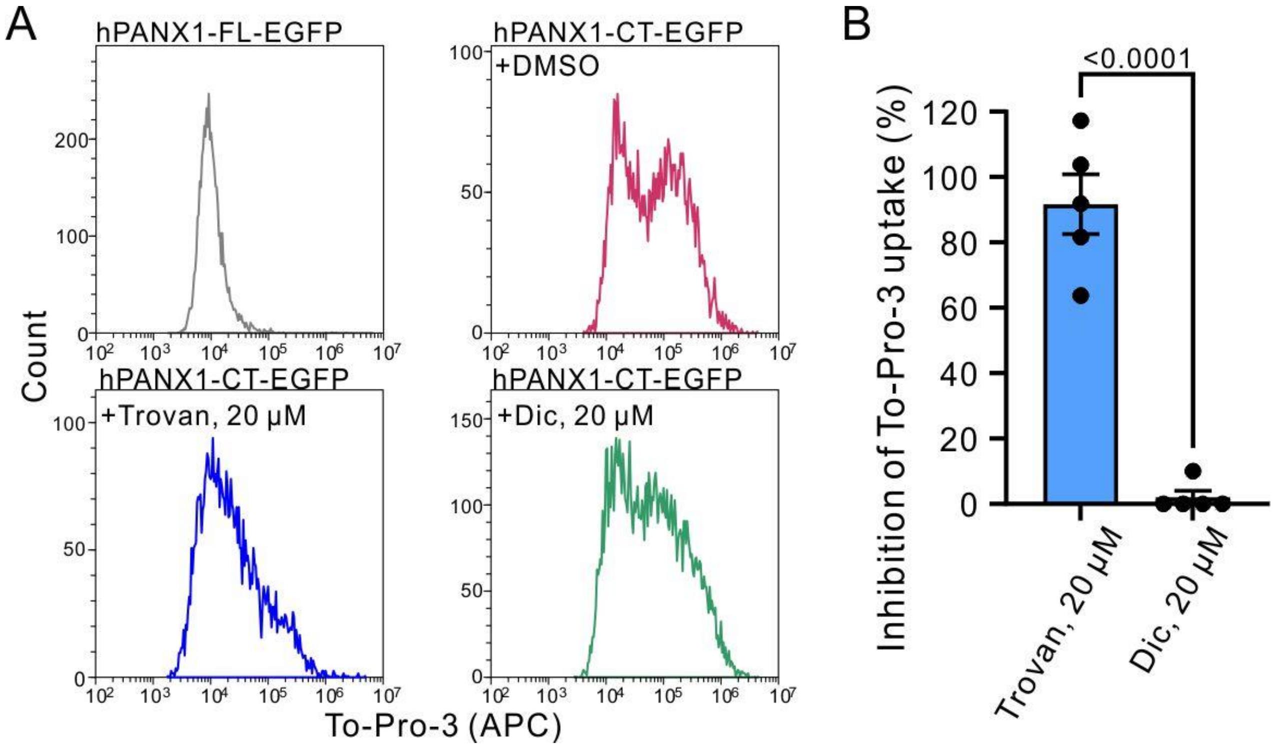


**Fig. S6. Dicoumarol does not inhibit To-Pro-3 uptake mediated by the C-terminally-cleaved hPANX1 channels. (A)** Exemplar histograms showing To-Pro-3 uptake of HEK293T cells expressing either hPANX1-FL-EGPF or hPANX1-CT-EGFP, with or without treatments of DMSO, trovafloxacin (Trovan; 20 μM), or dicoumarol (Dic, 20 μM). **(B)** Percent inhibition of To-Pro-3 uptake showing Trovan (91.7 ± 9.2%), but not Dic (2.0 ± 2.0%), inhibited C-terminally-cleaved hPANX1 channels. n=5 biologically independent experiments. *P*<0.0001 using two-tailed, unpaired t test.

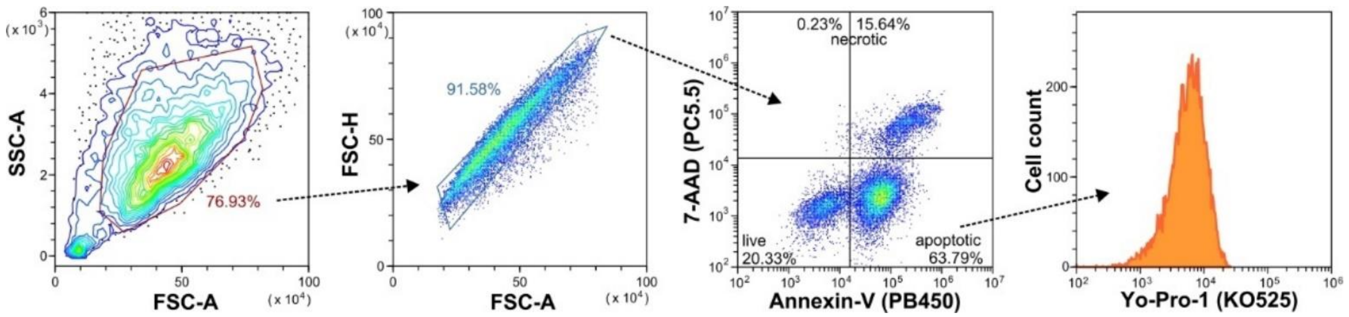


**Fig. S7. Electronic gating strategy for analyzing Yo-Pro-1 uptake in the apoptotic Jurkat cells.** Flow cytometry analyses showing electronic gating strategy used to distinguish live (Annexin $V^-$/7-$AAD^-$), apoptotic (Annexin $V^+$/7-$AAD^-$) and necrotic (7-$AAD^+$) subpopulations of Jurkat cells following 2 hours of UV irradiation (100 mJ $cm^{-2}$). Mean fluoresence intensity of Yo-Pro-1 uptake was further analyzed from the apoptotic cell population (Annexin $V^+$/7-$AAD^-$).

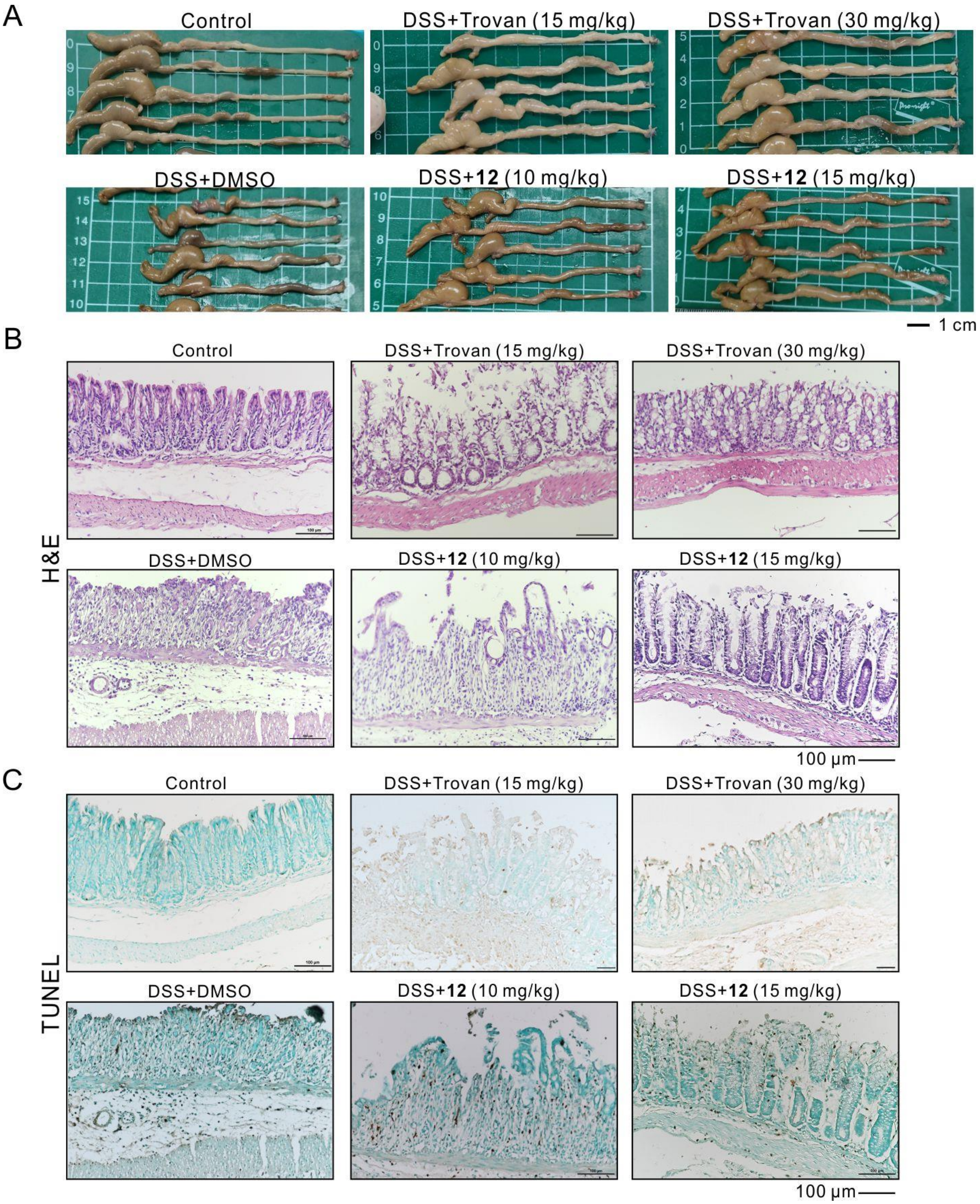


**Fig. S8. Macroscopic and histological effects of compound 12 on DSS-induced colitis mice. (A)** Exemplar macroscopic structures of control or DSS-fed mice, with or without treatments of Trovan or compound **12** as indicated. **(B)** Representative H&E-stained colon sections of control or DSS-treated mice, with or without treatments of Trovan or compound **12** as indicated. **(C)** Representative results of TUNEL assays performed using distal colon sections of control or DSS-induced colitis mice, with or without administration of Trovan or compound **12** as indicated.

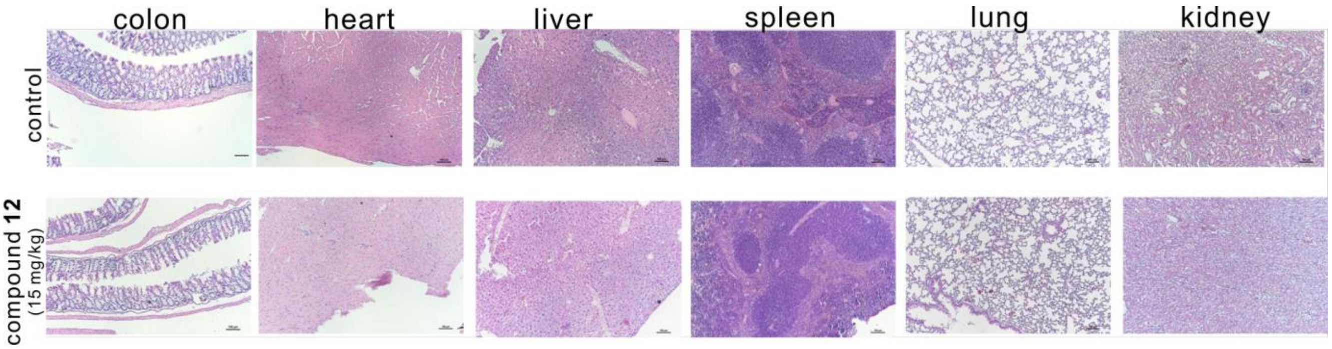


**Fig. S9. Histological examination of compound 12-treated mouse tissues showed no noticeable changes.** H&E-stained tissue samples showing that there was no obvious structural difference between the control mice (upper) and compound-**12**-treated (15 mg/kg) mice.
